# Overexpression of a G-protein coupled receptor-like gene affects encystment of *Acanthamoeba castellanii*

**DOI:** 10.1101/2021.01.07.425711

**Authors:** Steven Rolland, Anne Mercier, Luce Mengue, Yann Héchard, Ascel Samba-Louaka

## Abstract

*Acanthamoeba castellanii* is an amphizoïc free-living amoeba as it can be found in humans and in the environment. This amoeba represents an important reservoir of pathogenic microorganisms. Persistence of *A. castellanii* in the environment or in humans is allowed by the ability of the vegetative form to differentiate under cysts when surrounding conditions are unfavorable. In this study, we investigate the role of the *ACA1_383450* gene during encystment of *A. castellanii.* This gene encodes a putative G-protein coupled receptor, which shares homology with human GPR107 and murine GPR108. Expression of the *ACA1_383450* gene is transiently repressed at the early phase of encystment and its overexpression affects encystment of *A. castellanii.* This study reveals a new *Acanthamoeba* gene which could affect the encystment process.

**Highlights:**

- The *ACA1_383450* gene encodes for a putative G-protein coupled receptor (GPCR).
- The *ACA1_383450* mRNA levels are down-regulated during the early phase of encystment.
- Overexpression of the *ACA1_383450* gene affects formation of cysts.

## 1. Introduction

Species from the genus *Acanthamoeba* are free-living amoebae ubiquitous of natural and artificial aquatic environment (de Jonckheere, 1991; Marciano-Cabral and Cabral, 2003). In human, some strains are causative agents of granulomatous amoebic encephalitis, a rare fatal infection of the central nervous system, and of *Acanthamoeba* keratitis, a painful progressive eye disease (Marciano-Cabral and Cabral, 2003; Martinez and Visvesvara, 1997). Depending of the environmental conditions, *Acanthamoeba* spp. can adopt two states. The trophozoite state is a metabolically active form, which can move or feed on microorganisms (Bowers, 1977; Bowers and Olszewski, 1983). Under harsh conditions such as starvation, treatment with therapeutic agents, or change in pH or osmotic pressure, the trophozoite can differentiate into a cyst by a process called encystment (Fouque et al., 2012; Lloyd, 2014). Cysts represent resting and resistant forms of *Acanthamoeba.* The encystment process induces a rounding of the cell, a decreasing of the cell size, a growth arrest, changes in biochemical properties and the formation of a cell wall (Chávez-Munguía et al., 2013, 2005). Several genes associated with cell wall synthesis, such as glycogen phosphorylase, cellulose synthase and xylose isomerase have been identified and their repression prevents the formation of cysts (Aqeel et al., 2013; Lorenzo-Morales et al., 2008; Moon et al., 2014). The involvement of autophagy genes and of cysteine and serine proteases activities were also described (Dudley et al., 2008; Kim et al., 2015; Leitsch et al., 2010; Moon et al., 2015). However, signalling events involved in encystment remains still poorly understood.

The G-protein coupled receptors (GPCR) is one of the largest family of membrane receptors in mammalian cells. They are characterized by the presence of seven membrane-spanning α-helical segments separated by alternating intracellular and extracellular loop regions (Rosenbaum et al., 2009). They are involved in numerous cellular functions such as intracellular trafficking, cell cycle, mitosis, cytoskeletal reorganization, chemotaxis and phagocytosis, that make them an ideal targets for many therapeutics drugs (Hill, 2006; Lagerström and Schiöth, 2008). The analysis of *Acanthamoeba castellanii* str. Neff genome predicted 35 GPCR genes (Clarke et al., 2013). Recently, it has been described that the inhibition of β-adrenergic receptor (a class of GPCR) by propranolol affected the growth, the viability and the encystment process (Aqeel et al., 2015). The role of the other GPCR in *A. castellanii* encystment is still pending.

Here, we have investigated the importance of the *ACA1_383450* gene during encystment of *A. castellanii.* This gene encodes a putative G-protein coupled receptor. Its expression is transiently repressed during initial step of encystment. Overexpression of the *ACA1_383450* coding sequence does not alter the cell growth but affects the encystment process.

## 2. Materials and methods

### 2.1 Amoeba strains and cultural conditions

*A. castellanii* ATCC 30010 was grown at 30°C without shaking, in culture flask containing Peptone Yeast Glucose (PYG) medium (2 % proteose peptone, 0.1 % yeast extract, 0.1 % sodium citrate dehydrate, 0.1 M glucose, 4 mM MgSO_4_, 2.5 mM NaH_2_PO_3_, 2.5 mM K_2_HPO_3_, 0.4 mM CaCl_2_, 0.05 mM Fe(NH_4_)_2_(SO_4_)_2_ 6H_2_O, pH 6.5). For the transfected cells, G418 (Geneticin) was used at 50 μg/mL as selection pressure.

For the growth assay, *A. castellanii* trophozoites were seeded onto 24-well plates at a density of 5 x 10^4^ cells per well in 1 ml of Page’s Amoeba Saline solution (PAS) (0.1 % sodium citrate dehydrate, 4 mM MgSO_4_, 2.5 mM NaH_2_PO_3_, 2.5 mM K_2_HPO_3_, 0.4 mM CaCl_2_, 0.05 mM Fe(NH_4_)_2_(SO_4_)_2_ 6H_2_O, pH 6.5) and incubate at 30°C during 1 hour for cell adhesion. Then, PAS buffer was replaced by PYG growth medium (corresponding to time 0) and incubate at 30°C. Cells were harvested at 2 h, 24 h, 48 h and 72 h and counted using plastic counting slides FastRead 102® (Biosigma). All samples were counted three times and in three independent experiments.

### 2.2 Encystment assay

*A. castellanii* trophozoites were seeded onto 24-well plates at a density of 5 x 10^4^ cells per well in PAS buffer and incubated at 30°C during 1 hour for cell adhesion. PAS was then replaced by an encystment medium (0.1 M KCl, 8 mM MgSO_4_, 0.4 mM CaCl_2_, 1 mM NaHCO_3_ and 20 mM 2-amino-2-methyl-1,3-propanediol, pH 8.8) and incubated at 30°C (corresponding to time 0) up to 72 hours. At 24 h, 48 h, and 72 h, calcofluor white, a dye that binds to cellulose, was added into the wells following the manufacturer’s recommendations. The cysts were observed by fluorescence microscopy (Olympus IX51). More than 800 cells were counted per condition and per experiment. This experiment was done in three independent replicates.

### 2.3 Plasmid constructions and cloning

For the construction of the pTBPF-ACA1_383450 plasmid, the coding sequence of the *ACA1_383450* gene was amplified by PCR using total cDNA with flanking NdeI and SalI restriction sites, and finally ligated into the pTBPF-eGFP (Bateman, 2010). The pTBPF-eGFP was previously digested by NdeI and XhoI. The DNA digestion by SalI (insert) or XhoI (plasmid) generate compatible cohesive ends. PCR primers are listed in the Table 1. The pTBPF-empty (Rolland et al., 2020) was used as a control vector. All plasmids constructs were transformed in chemically-competent *Escherichia coli* DH5α and validated by SANGER sequencing. DNA sequencing was completed with the ABI Prism BigDye terminator v3.1 sequencing kit (Applied Biosystems) and then analysed by an automatic ABI Prism 3730 genetic analyser (Applied Biosystems).

**Table 1:**
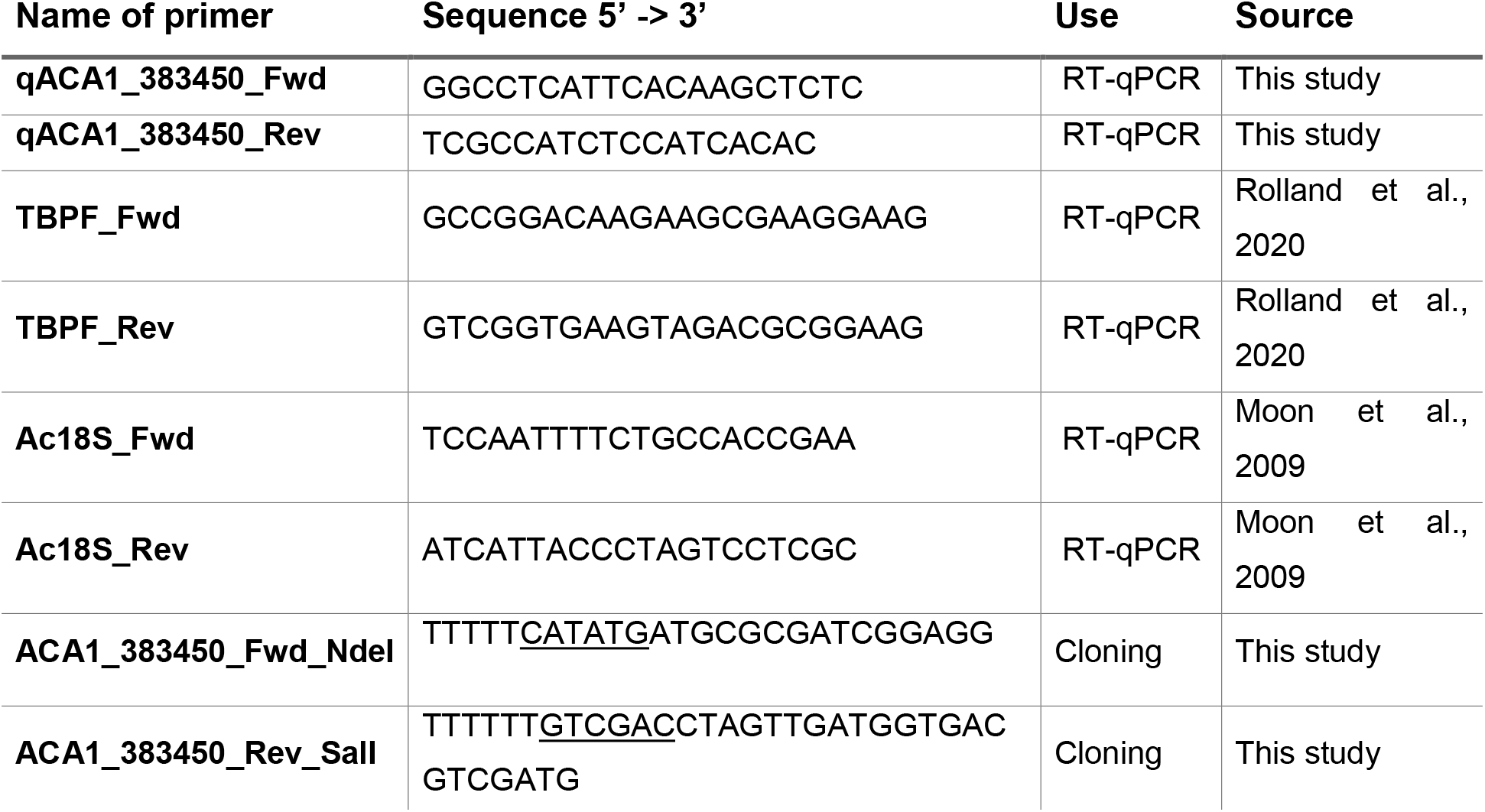
Primers used for RT-qPCR and plasmid constructions. “Fwd” for forward primer and “Rev” for reverse primer. The restriction site added was underlined.

### 2.4 Transfection of cells

Transfection of *A. castellanii* trophozoites were performed using the Viafect™ transfection reagent (Promega) as previously described (Rolland et al., 2020). Briefly, the cells were seeded into 24-well plates at a density of 1.25 x 10^5^ cells per well in encystment medium. The transfection medium containing reagent at a ratio of 5:1 (transfection reagent [μl]/plasmid [μg]) in the encystment medium was added directly on cells. After 3 h of incubation at 30°C, PYG medium was added in the wells and the plate were incubated during 24 hours. Then, the well content was transferred into a culture flask containing PYG medium supplemented with 20 μg/mL of G418 (Sigma) for two weeks. The concentration of G418 was then increased to 50 μg/mL to select transfected population.

### 2.5 Reverse transcription – quantitative PCR (RT-qPCR)

The RNeasy Mini Kit (Qiagen) was used to extract total RNA. The RNA samples were treated with RNase-free DNase I (Turbo DNA-free kit, Invitrogen) and reverse transcribed with the GoScript reverse transcriptase kit (Promega) according to the manufacturer’s recommendations. The reverse transcription products were used to carry out quantitative PCR. All primer sequences are shown in Table 1.

The LightCycler FastStart DNA Master plus SYBR Green I (Roche Applied Science) was used to perform all quantitative RT-PCR (RT-qPCR) as previously described (Rolland et al., 2020). Reactions were prepared in a total volume of 10 μl containing 5 μl of 2X SYBR mix, 2 μl of H_2_O, 2 μl of diluted cDNA template, and 0.5 μl of 10 μM primers.

The reactions were performed under the following conditions: an initial denaturation step of 95°C for 5 min, followed by a three-step thermal cycling profile comprising denaturation at 95°C for 10 s, primer annealing at 60°C for 10 s, and extension at 72°C for 10 s. This procedure was conducted for 45 cycles. To verify the specificity of the amplicon for each primer pair, a melting curve analysis was performed ranging from 65°C to 95°C.

The relative quantification method (2^-ΔΔCt^) was used to evaluate quantitative variation between replicates (Livak and Schmittgen, 2001). The relative expression of the *ACA1_383450* gene were normalized towards *tbpf* gene (NCBI accession: L46867.1). The overexpression of *ACA1_383450* in transfected amoebae was confirmed using the 18S rRNA gene as reference gene (Figure 1D). Primers used for RT-qPCR experiments were described in Table 1.

**Figure 1:**
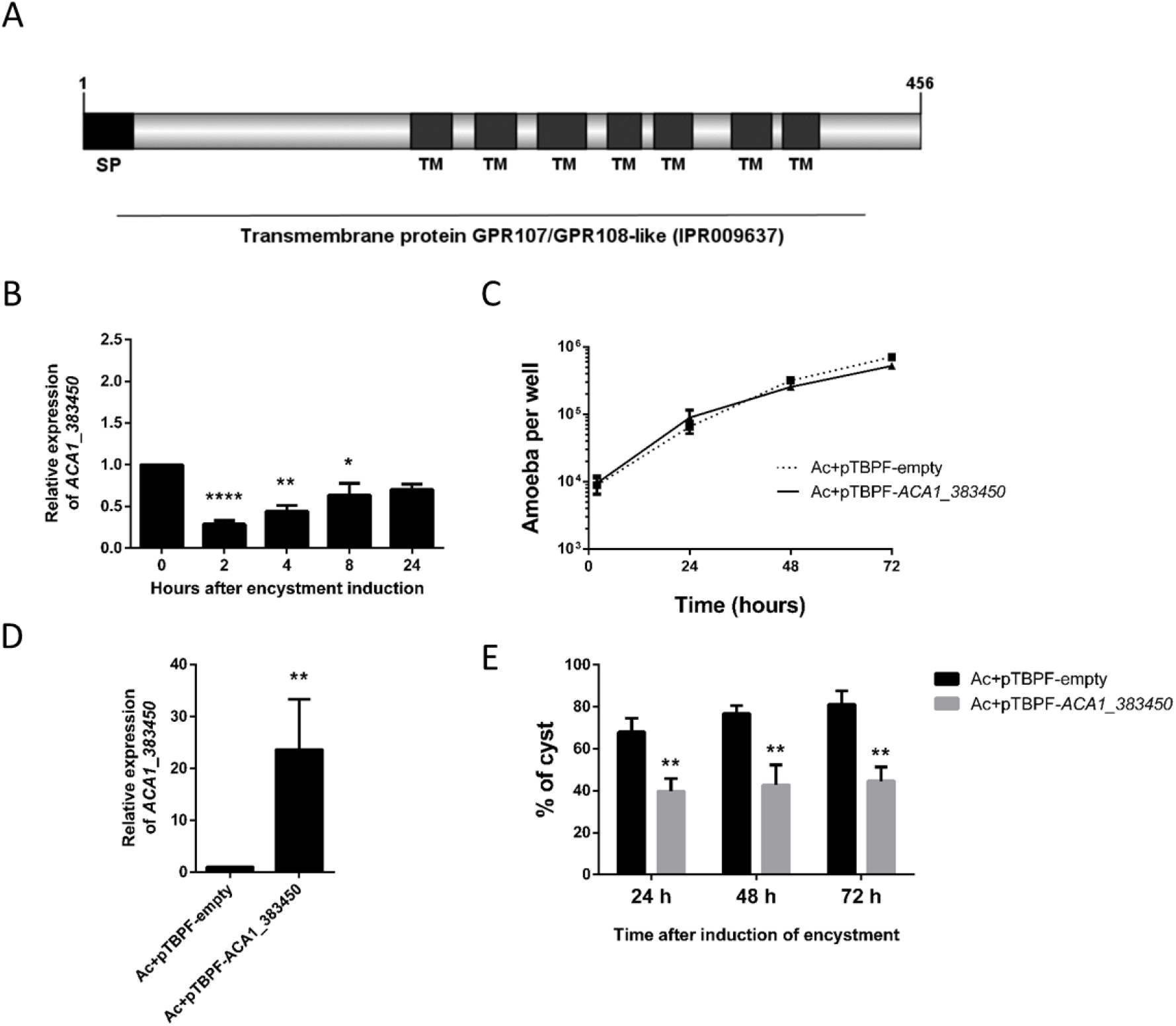
Expression and overexpression effect of *ACA1_383450* during *A. castellanii* encystment. (A) Domain prediction of the *ACA1_383450* gene using Interpro. Illustration made by IBS 1.0.3 software (Liu et al., 2015) (B) Relative expression of the ACA1_383450 gene during encystment performed by RT-qPCR. (C) Effect of the overexpression on *Acanthamoeba* growth. (D) The overexpression of *ACA1_383450* were analysed by RT-qPCR. (E) Effect of the overexpression on *Acanthamoeba* encystment. Results represent average values of the three independent experiments, and error bars represent the standard error of the mean (±SEM). Statistical analysis was performed sing the Student t-test (*p < 0.05, **p<0.01, ***p<0.001, ****p<0.0001). The ΔCt values were used for RT-qPCR for statistical analysis. Means were compared to the condition ‘0 hour’ (B) or Ac+pTBPF-empty’ (D-E).

### 2.6 Statistical analysis

All results are average of three independent experiments, and error bars represent the standard error of the mean (±SEM). Statistical analysis was performed using the unpaired Student t-test (GraphPad Prism 6). For RT-qPCR experiments, statistical analyses were performed on ΔCt values. Statistically significant differences were considered statistically significant when P values were < 0.05 (*p<0.05; **p<0.01; ***p<0.001;****p<0.0001).

## 3. Results

### 3.1. The *ACA1_383450* gene encodes for a putative G-protein coupled receptor (GPCR)

The analysis of *A. castellanii* genome exhibits 35 genes predicted to encode GPCR (Clarke et al., 2013). Inhibition of one G-protein coupled receptor (β adrenergic receptor) was reported to block encystment of *A. castellanii* (Aqeel et al., 2015). In this study, we were able to amplify and to clone another putative GPCR encoded by the *ACA1_383450* gene. This latter encodes for a protein of 456 amino-acids. As expected, the domain prediction using InterPro shows the presence of seven transmembrane helix, which are a characteristic of this protein family. A peptide signal is also predicted at the C-terminal position and serves, as many other proteins, for their addressing to their final cellular localization (Fig. 1A).

In order to find homologs of the ACA1_383450 protein, a Basic Local Alignment Search Tool (BLAST) analysis was performed against the NCBI non-redundant protein sequence database. The BLASTp analysis shows that the protein shares homologies with GPR107 and GPR108 proteins from multiple organisms (Supplementary data 1). Among them, we found the human GPR107 and the murine GPR108 which are described in literature (Supplementary data 2 and 3).

### 3.2. The *ACA1_383450* mRNA levels are down-regulated at the beginning of encystment

In order to study the expression of the *ACA1_383450* gene during encystment, a RT-qPCR was conducted at 0, 2, 4, 8 and 24 hours after the induction of the encystment. As observed on the Figure 1B, the *ACA1_383450* gene was transiently down-regulated after incubation of *A. castellanii* into the encystment medium. After 24 h its expression return to a level similar to the control.

### 3.3. Overexpression of the *ACA1_383450* gene affects the formation of cysts

To better characterize the function of the *ACA1_383450* gene, we examined if its overexpression could affect encystment. A plasmid with the *ACA1_383450* coding sequence (pTBPF-*ACA1_383450*) was constructed. In this plasmid, the *ACA1_383450* coding sequence was under the promoter of the TATA-Binding Protein Promoter Binding Factor (TPBF) for a constitutive expression even during the encystment (Bateman, 2010). Indeed, the mRNA encoding the TPBF was reported stable during the encystment (Orfeo and Bateman, 1998). As a control, we used the plasmid pTBPF-empty, which does not express of the coding sequence (Rolland et al., 2020). The growth of *A. castellanii* was not affected by the overexpression of the *ACA1_384820* gene compared cells transfected with the control plasmid pTBPF-empty (Fig. 1C). We incubated transfected amoebae within the encystment medium and evaluated the percentage of cysts using the calcofluor white stain at different time points (24 h, 48 h and 72 h). *A. castellanii* transfected with the pTBPF-empty was reported to encyst as efficiently as the untransfected amoebae (Rolland et al., 2020). Twenty-four hours after the induction of encystment, we observed a decrease of the percentage of cysts upon overexpression of ACA1_383450. This effect was more pronounced at 48 h and 72 h, and the cyst ratio is almost halved (Fig. 1E). The *ACA1_383450* expression of the transfected cells was assessed by RT-qPCR (Figure 1D).

Altogether, these results show that the overexpression of the *ACA1_383450* coding sequence did not disturb the growth of *A. castellanii* but affected formation of cysts.

## 4. Discussion

In this study, we have identified a new *Acanthamoeba* gene affecting encystment. In non-transfected cells, the *ACA1_383450* gene was transiently down-regulated during encystment. This gene encodes a putative G-protein coupled receptor (GPCR). GPCRs form a large superfamily of transmembrane receptors in Eukaryota. They are signal mediators that play a prominent role in most major physiological processes such as differentiation, cell cycle or vesicular trafficking. GPCRs are classified into different families including class A (rhodopsin-like receptors), class B (secretin receptors), class C (glutamate receptors), class D (fungal mating pheromone receptors), class E (cAMP receptors), class F (frizzled receptors), and other 7TM proteins (de Mendoza et al., 2014; Foord et al., 2005; Fredriksson and Schiöth, 2005; Kolakowski, 1994). It has been shown that, in *A. castellanii,* inhibition of the β-adrenergic receptor (a GPCR class A) by propranolol abolished its extracellular proteolytic activities, affected cell viability, blocked its growth and its encystation process (Aqeel et al., 2015). This result shows an involvement of the GPCRs during differentiation of *A. castellanii* into cyst, but the involvement of others GPCR, like ACA1_383450, was not reported.

BLAST analysis of the ACA1_383450 indicates that this protein shares homologies with GPR107 and GPR108 proteins. GPR107 in human and GPR108 in mice are members of the long extracellular domain and a carboxy-terminal seven transmembrane (LUSTR) family of proteins found in animals and plants; but absent from bacteria, archaea and viruses (Edgar, 2007). GPR107 belongs to “other 7TM proteins” class of GPCR. In mammals cells, GPR107 localizes to the trans-golgi network and could be involved in retrograde trafficking (Tafesse et al., 2014). This latter is essential for encystment of organisms such as *Entamoeba invadens* and *Giardia intestinalis* (Hehl and Marti, 2004; Herman et al., 2017). The role of ACA1_383450 in retrograde trafficking should be investigated.

We found that expression of *ACA1_383450* was transiently down-regulated in early phase of encystment. Ongoing work consists on determining the molecular targets and activities of ACA1_383450 protein. More generally, it will be also interesting to study the involvement of the different classes of GPCR during encystment of *A. castellanii.*

## 5. Conclusion

In conclusion, we describe a new *ACA1_383450* gene of which expression is transiently down-regulated during encystment of *A. castellanii.* Overexpression of *ACA1_383450* affects formation of cysts. This protein is a putative GPCR sharing homology with two members of the LUSTR family: GPR107 and GPR108. Further studies are needed to determine the activity of this protein and its specific role in encystment of *A. castellanii*.

## Supporting information

Supplementary 1 - BLASTp results

## Acknowledgements

We gratefully acknowledge Dr. Hong Yeonchul for providing the pTBPF-eGFP plasmid. We thank Dr. Willy Aucher for molecular biology advices. We thank Marie-Laure Mollichella and Daniel Guyonnet for their help on cytometer analysis, RT-qPCR and DNA sequencing.

## Author contributions

**Steven Rolland:** conceptualization, Data curation, Formal analysis, Investigation, Methodology, Validation, Visualization, Roles/Writing - original draft, Writing - review &editing

**Anne Mercier**: Data curation, Formal analysis, Writing - review &editing

**Luce Mengue**: Data curation, Formal analysis, Validation

**Yann Héchard**: conceptualization, Funding acquisition, Investigation, Methodology, Project administration, Resources, Supervision, Validation, Writing - review &editing

**Ascel Samba-Louaka**: conceptualization, Formal analysis, Funding acquisition, Investigation, Methodology, Project administration, Resources, Supervision, Validation, Visualization, Roles/Writing - original draft, Writing - review &editing

## References

Aqeel, Y., Siddiqui, R., Khan, N.A., 2013. Silencing of xylose isomerase and cellulose synthase by siRNA inhibits encystation in *Acanthamoeba castellanii*. Parasitol. Res. 112, 1221–1227. https://doi.org/10.1007/s00436-012-3254-6

Aqeel, Y., Siddiqui, R., Manan, Z., Khan, N.A., 2015. The role of G protein coupled receptor-mediated signaling in the biological properties of *Acanthamoeba castellanii* of the T4 genotype. Microb. Pathog. 81, 22–7. https://doi.org/10.1016/j.micpath.2015.03.006

Bateman, E., 2010. Expression plasmids and production of EGFP in stably transfected *Acanthamoeba*. Protein Expr. Purif. 70, 95–100. https://doi.org/10.1016/j.pep.2009.10.008

Bowers, B., 1977. Comparison of pinocytosis and phagocytosis in *Acanthamoeba castellanii*. Exp. Cell Res. 110, 409–417. https://doi.org/10.1016/0014-4827(77)90307-X

Bowers, B., Olszewski, T.E., 1983. *Acanthamoeba* discriminates internally between digestible and indigestible particles. J. Cell Biol. 97, 317–322. https://doi.org/10.1083/jcb.97.2.317

Chávez-Munguía, B., Omaña-Molina, M., González-Lázaro, M., González-Robles, A., Bonilla-Lemus, P., Martínez-Palomo, A., 2005. Ultrastructural study of encystation and excystation in *Acanthamoeba castellanii*. J. Eukaryot. Microbiol. 52, 153–158. https://doi.org/10.1111/j.1550-7408.2005.04-3273.x

Chávez-Munguía, B., Salazar-Villatoro, L., Lagunes-Guillén, A., Omaña-Molina, M., Espinosa-Cantellano, M., Martínez-Palomo, A., 2013. *Acanthamoeba castellanii* cysts: new ultrastructural findings. Parasitol. Res. 112, 1125–1130. https://doi.org/10.1007/s00436-012-3261-7

Clarke, M., Lohan, A.J., Liu, B., Lagkouvardos, I., Roy, S., Zafar, N., Bertelli, C., Schilde, C., Kianianmomeni, A., Bürglin, T.R., Frech, C., Turcotte, B., Kopec, K.O., Synnott, J.M., Choo, C., Paponov, I., Finkler, A., Heng Tan, C.S., Hutchins, A.P., Weinmeier, T., Rattei, T., Chu, J.S.C., Gimenez, G., Irimia, M., Rigden, D.J., Fitzpatrick, D.A., Lorenzo-Morales, J., Bateman, A., Chiu, C.H., Tang, P., Hegemann, P., Fromm, H., Raoult, D., Greub, G., Miranda-Saavedra, D., Chen, N., Nash, P., Ginger, M.L., Horn, M., Schaap, P., Caler, L., Loftus, B.J., 2013. Genome of *Acanthamoeba castellanii* highlights extensive lateral gene transfer and early evolution of tyrosine kinase signaling. Genome Biol. 14, R11. https://doi.org/10.1186/gb-2013-14-2-r11

de Jonckheere, J.F., 1991. Ecology of *Acanthamoeba.* Rev. Infect. Dis. 13, S385–S387. https://doi.org/10.1093/clind/13.Supplement_5.S385

de Mendoza, A., Sebé-Pedrós, A., Ruiz-Trillo, I., 2014. The evolution of the GPCR signaling system in eukaryotes: modularity, conservation, and the transition to metazoan multicellularity. Genome Biol. Evol. 6, 606–619. https://doi.org/10.1093/gbe/evu038

Dudley, R., Alsam, S., Khan, N.A., 2008. The role of proteases in the differentiation of *Acanthamoeba castellanii*. FEMS Microbiol. Lett. 286, 9–15. https://doi.org/10.1111/j.1574-6968.2008.01249.x

Edgar, A.J., 2007. Human GPR107 and murine Gpr108 are members of the LUSTR family of proteins found in both plants and animals, having similar topology to G-protein coupled receptors. DNA Seq.-J. DNA Seq. Mapp. 18, 235–241. https://doi.org/10.1080/10425170701207182

Foord, S.M., Bonner, T.I., Neubig, R.R., Rosser, E.M., Pin, J.-P., Davenport, A.P., Spedding, M., Harmar, A.J., 2005. International Union of Pharmacology. XLVI. G Protein-Coupled Receptor List. Pharmacol. Rev. 57, 279–288. https://doi.org/10.1124/pr.57.2.5

Fouque, E., Trouilhé, M.-C., Thomas, V., Hartemann, P., Rodier, M.-H., Héchard, Y., 2012. Cellular, biochemical, and molecular changes during encystment of free-living amoebae. Eukaryot. Cell 11, 382–387. https://doi.org/10.1128/EC.05301-11

Fredriksson, R., Schiöth, H.B., 2005. The repertoire of G–protein–coupled receptors in fully sequenced genomes. Mol. Pharmacol. 67, 1414–1425. https://doi.org/10.1124/mol.104.009001

Hehl, A.B., Marti, M., 2004. Secretory protein trafficking in *Giardia intestinalis*. Mol. Microbiol. 53, 19–28. https://doi.org/10.1111/j.1365-2958.2004.04115.x

Herman, E., Siegesmund, M.A., Bottery, M.J., Van Aerle, R., Shather, M.M., Caler, E., Dacks, J.B., Van Der Giezen, M., 2017. Membrane trafficking modulation during *Entamoeba* encystation. Sci. Rep. 7, 1–17. https://doi.org/10.1038/s41598-017-12875-6

Hill, S.J., 2006. G-protein-coupled receptors: past, present and future. Br. J. Pharmacol. https://doi.org/10.1038/sj.bjp.0706455

Kim, S.H., Moon, E.K., Hong, Y., Chung, D. Il, Kong, H.H., 2015. Autophagy protein 12 plays an essential role in *Acanthamoeba* encystation. Exp. Parasitol. 159, 46–52. https://doi.org/10.1016/j.exppara.2015.08.013

Kolakowski, L.F., 1994. GCRDb: a G-protein-coupled receptor database. Receptors Channels 2, 1–7.

Lagerström, M.C., Schiöth, H.B., 2008. Structural diversity of G protein-coupled receptors and significance for drug discovery. Nat. Rev. Drug Discov. 7, 339–357. https://doi.org/10.1038/nrd2518

Leitsch, D., Köhsler, M., Marchetti-Deschmann, M., Deutsch, A., Allmaier, G., Duchêne, M., Walochnik, J., 2010. Major role for cysteine proteases during the early phase of *Acanthamoeba castellanii* encystment. Eukaryot. Cell 9, 611–8. https://doi.org/10.1128/EC.00300-09

Liu, W., Xie, Y., Ma, J., Luo, X., Nie, P., Zuo, Z., Lahrmann, U., Zhao, Q., Zheng, Y., Zhao, Y., Xue, Y., Ren, J., 2015. IBS: an illustrator for the presentation and visualization of biological sequences: Fig. 1. Bioinformatics 31, 3359–3361. https://doi.org/10.1093/bioinformatics/btv362

Livak, K.J., Schmittgen, T.D., 2001. Analysis of relative gene expression data using real-time quantitative PCR and the 2-ΔΔCT method. Methods 25, 402–408. https://doi.org/10.1006/meth.2001.1262

Lloyd, D., 2014. Encystment in *Acanthamoeba castellanii:* A review. Exp. Parasitol. 145, S20–S27. https://doi.org/10.1016/j.exppara.2014.03.026

Lorenzo-Morales, J., Kliescikova, J., Martinez-Carretero, E., De Pablos, L.M., Profotova, B., Nohynkova, E., Osuna, A., Valladares, B., 2008. Glycogen phosphorylase in *Acanthamoeba* spp.: Determining the role of the enzyme during the encystment process using RNA interference. Eukaryot. Cell 7, 509–517. https://doi.org/10.1128/EC.00316-07

Marciano-Cabral, F., Cabral, G., 2003. *Acanthamoeba* spp. as agents of disease in humans. Clin. Microbiol. Rev. 16, 273–307. https://doi.org/10.1128/CMR.16.2.273

Martinez, A.J., Visvesvara, G.S., 1997. Free-living, amphizoic and opportunistic amebas. Brain Pathol. 583–598.

Moon, E.-K.K., Kim, S.-H.H., Hong, Y., Chung, D.-I. Il, Goo, Y.-K.K., Kong, H.-H.H., 2015. Autophagy inhibitors as a potential antiamoebic treatment for *Acanthamoeba* keratitis. Antimicrob. Agents Chemother. 59, 4020–4025. https://doi.org/10.1128/AAC.05165-14

Moon, E.K., Chung, D. Il, Hong, Y.C., Kong, H.H., 2009. Autophagy protein 8 mediating autophagosome in encysting *Acanthamoeba*. Mol. Biochem. Parasitol. 168, 43–48. https://doi.org/10.1016/j.molbiopara.2009.06.005

Moon, E.K., Hong, Y., Chung, D. Il, Goo, Y.K., Kong, H.H., 2014. Down-regulation of cellulose synthase inhibits the formation of endocysts in *Acanthamoeba*. Korean J. Parasitol. 52, 131–135. https://doi.org/10.3347/kjp.2014.52.2.131

Orfeo, T., Bateman, E., 1998. Transcription by RNA polymerase II during *Acanthamoeba* differentiation. Biochim. Biophys. Acta-Gene Struct. Expr. 1443, 297–304. https://doi.org/10.1016/S0167-4781(98)00227-9

Rolland, S., Mengue, L., Noël, C., Crapart, S., Mercier, A., Aucher, W., Héchard, Y., Samba-Louaka, A., 2020. Encystment induces down-regulation of an acetyltransferase-like gene in *Acanthamoeba castellanii*. Pathogens 9. https://doi.org/10.3390/pathogens9050321

Rosenbaum, D.M., Rasmussen, S.G.F., Kobilka, B.K., 2009. The structure and function of G-protein-coupled receptors. Nature 459, 356–363. https://doi.org/10.1038/nature08144

Tafesse, F.G., Guimaraes, C.P., Maruyama, T., Carette, J.E., Lory, S., Brummelkamp, T.R., Ploegh, H.L., 2014. GPR107, a G-protein-coupled receptor essential for intoxication by *Pseudomonas aeruginosa* exotoxin a, localizes to the Golgi and is cleaved by furin. J. Biol. Chem. 289, 24005–24018. https://doi.org/10.1074/jbc.M114.589275

