## Supplementary 1 - BLASTp results for "Overexpression of a G-protein coupled receptor-like gene affects encystment of *Acanthamoeba castellanii*"

RID: M0FKT0TS016
Job Title:Protein Sequence
Program: BLASTP
Query: None ID: lcl|Query_81164(amino acid) Length: 456
Database: nr All non-redundant GenBank CDS translations+PDB+SwissProt+PIR+PRF excluding environmental samples from WGS projects

Sequences producing significant alignments:

| Description | Max Score | Total Score | Query cover | E Value | Per. Ident | Accession |
|---|---|---|---|---|---|---|
| G protein coupled receptor, putative [Acanthamoeba castellanii... | 934 | 934 | 100% | 0.0 | 100.00 | XP_004338827.1 |
| seven transmembrane domain protein [Heterostelium album PN500] | 376 | 376 | 86% | 2e-123 | 47.82 | XP_020432478.1 |
| hypothetical protein SAMD00019534_116630 [Acytostelium... | 367 | 367 | 92% | 6e-120 | 46.59 | XP_012748526.1 |
| GPR108 protein [Thecamonas trahens ATCC 50062] | 357 | 357 | 91% | 1e-115 | 43.22 | XP_013762250.1 |
| seven transmembrane domain protein [Cavenderia fasciculata] | 356 | 356 | 87% | 1e-115 | 48.19 | XP_004357792.1 |
| seven transmembrane domain protein [Planoprotostelium fungivorum] | 350 | 350 | 89% | 7e-112 | 43.99 | PRP82284.1 |
| seven transmembrane domain protein [Tieghemostelium lacteum] | 337 | 337 | 87% | 3e-108 | 45.18 | KYQ89580.1 |
| seven transmembrane domain protein [Dictyostelium discoideum AX4] | 331 | 331 | 90% | 1e-105 | 40.05 | XP_637699.2 |
| hypothetical protein KFL_002230120 [Klebsormidium nitens] | 316 | 316 | 88% | 7e-100 | 43.06 | GAQ85191.1 |
| Transmembrane receptor, eukaryota [Nannochloropsis gaditana] | 317 | 317 | 90% | 1e-99 | 41.34 | EWM24893.1 |
| PREDICTED: protein GPR107 [Ochotona princeps] | 313 | 313 | 89% | 2e-98 | 40.79 | XP_004593744.1 |
| PREDICTED: protein GPR107 isoform X1 [Galeopterus variegatus] | 310 | 310 | 87% | 4e-96 | 37.63 | XP_008577101.1 |
| hypothetical protein NGA_0212400 [Nannochloropsis gaditana... | 306 | 306 | 90% | 2e-95 | 41.11 | XP_005854159.1 |
| hypothetical protein CEUSTIGMA_g1453.t1 [Chlamydomonas eustigma] | 301 | 301 | 90% | 1e-94 | 40.50 | GAX74003.1 |
| protein GPR107 [Microcaecilia unicolor] | 303 | 303 | 88% | 3e-94 | 38.27 | XP_030063706.1 |
| G-protein coupled receptor [Chloropicon primus] | 300 | 300 | 84% | 1e-93 | 42.68 | QDZ20040.1 |
| hypothetical protein CYY_001089 [Polysphondylium violaceum] | 299 | 299 | 87% | 2e-93 | 38.94 | KAF2077626.1 |
| hypothetical protein COCSUDRAFT_53024 [Coccomyxa subellipsoide... | 298 | 298 | 83% | 3e-93 | 40.85 | XP_005649187.1 |
| hypothetical protein MARPO_0114s0018 [Marchantia polymorpha] | 296 | 296 | 85% | 3e-92 | 40.94 | PTQ31198.1 |
| PREDICTED: protein GPR107 [Callorhinchus milii] | 299 | 299 | 91% | 4e-92 | 36.49 | XP_007901015.1 |
| predicted protein [Ostreococcus lucimarinus CCE9901] | 295 | 295 | 89% | 5e-92 | 38.88 | XP_001419678.1 |

uncharacterized protein HaLaN_07828 [Haematococcus lacustris] 292 292
61% 1e-91 50.00 GFH12190.1
protein GPR107 [Ricinus communis] 293 293
84% 3e-91 39.61 XP_025013062.1
hypothetical protein AXG93_1838s1160 [Marchantia polymorpha... 296 296
83% 7e-91 41.21 OAE31806.1
protein GPR107-like [Physcomitrium patens] 291 291
86% 1e-90 42.16 XP_024386861.1
protein GPR107 isoform X2 [Cimex lectularius] 291 291
87% 3e-90 39.48 XP_014261704.1
PREDICTED: protein GPR107 isoform X2 [Galeopterus variegatus] 292 292
64% 3e-90 47.92 XP_008577102.1
protein GPR107 isoform X1 [Rhincodon typus] 294 337
90% 4e-90 42.40 XP_020386814.1
PREDICTED: protein GPR107 isoform X2 [Latimeria chalumnae] 293 293
64% 5e-90 47.12 XP_014351767.1
hypothetical protein NSK_002311 [Nannochloropsis salina CCMP1776] 290 290
81% 7e-90 42.28 TFJ86657.1
protein GPR107 [Echinops telfairi] 285 285
64% 2e-89 46.96 XP_012862854.1
Transmembrane receptor, eukaryota [Nannochloropsis gaditana] 291 291
82% 2e-89 41.92 EWM24894.1
protein GPR107 isoform X1 [Amblyraja radiata] 292 292
64% 2e-89 47.44 XP_032904977.1
protein GPR108-like [Oncorhynchus mykiss] 285 285
64% 2e-89 47.57 XP_021412141.1
hypothetical protein CBR_g49976 [Chara braunii] 286 286
63% 3e-89 49.01 GBG65182.1
protein GPR107 isoform X3 [Balaenoptera acutorostrata scammoni] 287 287
66% 3e-89 46.42 XP_028017752.1
predicted protein [Micromonas pusilla CCMP1545] 288 288
89% 4e-89 39.95 XP_003056720.1
PREDICTED: protein GPR107 isoform X1 [Latimeria chalumnae] 291 291
64% 5e-89 47.12 XP_006008808.1
protein GPR107 isoform X4 [Pongo abelii] 284 284
64% 5e-89 47.76 XP_009243232.1
protein GPR107 isoform X2 [Nomascus leucogenys] 285 285
66% 8e-89 47.32 XP_030673890.1
protein GPR107-like [Durio zibethinus] 287 287
89% 9e-89 38.69 XP_022735937.1
PREDICTED: protein GPR107 isoform X1 [Saimiri boliviensis... 283 283
64% 9e-89 47.44 XP_010349023.1
unknown [Picea sitchensis] 287 287
91% 1e-88 39.55 ABR17223.1
protein GPR108-like isoform X1 [Etheostoma cragini] 290 290
64% 1e-88 47.57 XP_034715705.1
hypothetical protein GPECTOR_8g123 [Gonium pectorale] 286 286
85% 1e-88 40.77 KXZ52730.1
protein GPR108-like [Oncorhynchus mykiss] 282 282
63% 1e-88 48.03 XP_021412142.1
protein GPR107 isoform 4 [Homo sapiens] 283 283
64% 1e-88 47.44 NP_001274275.1
G protein-coupled receptor 107 [Molossus molossus] 284 284
67% 1e-88 45.68 KAF6433526.1

protein GPR107 isoform X3 [Rhinopithecus roxellana] 283 283
64% 1e-88 47.44 XP_010359244.1
protein GPR107 isoform X2 [Pteropus vampyrus] 283 283
64% 2e-88 46.65 XP_011362432.1
protein GPR107 isoform X2 [Rhincodon typus] 290 333
90% 2e-88 42.40 XP_020386815.1
protein CANDIDATE G-PROTEIN COUPLED RECEPTOR 7 [Cucumis sativus] 286 286
88% 2e-88 38.52 XP_011650293.1
protein GPR107 isoform X2 [Papio anubis] 283 283
66% 3e-88 47.00 XP_021782822.2
G protein-coupled receptor 107 [Molossus molossus] 285 285
66% 3e-88 45.79 KAF6433525.1
hypothetical protein [Chiloscyllium punctatum] 289 332
83% 3e-88 46.98 GCC30454.1
PREDICTED: protein GPR107 [Daucus carota subsp. sativus] 285 285
89% 4e-88 38.86 XP_017248308.1
protein GPR107-like [Cynara cardunculus var. scolymus] 285 285
95% 5e-88 38.75 XP_024985659.1
PREDICTED: protein GPR107 [Charadrius vociferus] 281 281
65% 9e-88 45.86 XP_009886300.1
hypothetical protein [Gossypium davidsonii] 283 283
92% 3e-87 36.88 MBA0621736.1
hypothetical protein [Gossypium raimondii] 283 283
88% 3e-87 38.22 MBA0588979.1
Lung seven transmembrane receptor-like [Parasponia andersonii] 283 283
88% 4e-87 38.19 PON73519.1
hypothetical protein B456_007G019200 [Gossypium raimondii] 283 283
88% 4e-87 38.22 KJB39560.1
PREDICTED: protein GPR107 [Beta vulgaris subsp. vulgaris] 283 283
89% 6e-87 40.19 XP_010684174.1
hypothetical protein F3Y22_tig00113156pilonHSYRG00094 [Hibiscu... 282 282
89% 6e-87 37.38 KAE8662714.1
Lung_7-TM_R domain-containing protein [Cephalotus follicularis] 282 282
86% 6e-87 38.76 GAV88895.1
protein GPR107 [Carex littledalei] 283 283
87% 7e-87 38.37 KAF3332350.1
PREDICTED: protein GPR107 [Cucumis melo] 282 282
88% 8e-87 38.52 XP_008448554.1
protein GPR107-like [Physcomitrium patens] 281 281
89% 1e-86 40.14 XP_024394221.1
hypothetical protein F3Y22_tig00110785pilonHSYRG00198 [Hibiscu... 281 281
89% 2e-86 37.50 KAE8694305.1
hypothetical protein C1H46_025649 [Malus baccata] 281 281
84% 2e-86 39.36 TQD88760.1
hypothetical protein E2562_016196 [Oryza meyeriana var.... 282 282
85% 3e-86 38.63 KAF0902345.1
beta-carotene isomerase D27 [Hibiscus syriacus] 280 280
89% 3e-86 36.58 KAE8727010.1
G protein-coupled seven transmembrane receptor [Chlamydomonas... 280 280
85% 3e-86 40.83 XP_001696905.1
unnamed protein product [Digitaria exilis] 281 281
91% 3e-86 36.55 CAB3469365.1
protein GPR107 [Momordica charantia] 280 280
84% 3e-86 38.78 XP_022145456.1

hypothetical protein SO13M23_000006 [Saccharum officinarum]       281    281
89%   4e-86  37.70  AWA44644.1
unnamed protein product [Digitaria exilis]                        281    281
91%   5e-86  36.78  CAB3466863.1
protein GPR107-like [Malus domestica]                             280    280
84%   7e-86  39.11  XP_008356560.2
protein GPR107-like [Cucurbita pepo subsp. pepo]                  279    279
88%   9e-86  38.05  XP_023551801.1
hypothetical protein GOBAR_AA25214 [Gossypium barbadense]         279    279
84%   1e-85  38.94  PPR95457.1
hypothetical protein ES319_A11G019500v1 [Gossypium barbadense]    279    279
84%   1e-85  38.94  KAB2055199.1
protein GPR108 precursor [Zea mays]                               280    280
89%   1e-85  37.47  ACG34585.1
protein GPR107-like [Cucurbita moschata]                          279    279
88%   1e-85  38.50  XP_022923496.1
hypothetical protein [Colocasia esculenta]                        283    283
85%   2e-85  38.17  MQL86691.1
protein GPR107-like [Cucurbita maxima]                            278    278
88%   2e-85  38.50  XP_022965377.1
protein GPR107 [Populus trichocarpa]                              278    278
84%   2e-85  38.33  XP_024453523.1
Lung seven transmembrane receptor family protein [Tripterygium... 278    278
89%   2e-85  38.39  KAF5734950.1
PREDICTED: protein GPR107 [Populus euphratica]                    278    278
84%   3e-85  37.80  XP_011022403.1
hypothetical protein EJD97_007360 [Solanum chilense]              278    278
86%   3e-85  36.73  TMW81933.1
Transmembrane receptor, eukaryota [Corchorus olitorius]           278    278
89%   3e-85  37.68  OMP02814.1
hypothetical protein ES319_A02G106800v1 [Gossypium barbadense]    278    278
89%   4e-85  37.44  KAB2093650.1
PREDICTED: protein GPR107-like [Gossypium hirsutum]               278    278
89%   5e-85  37.44  XP_016741571.1
hypothetical protein EE612_031732 [Oryza sativa]                  278    278
85%   5e-85  38.39  KAB8101064.1
hypothetical protein EE612_031732 [Oryza sativa]                  278    278
85%   5e-85  38.39  KAB8101063.1
protein GPR107 [Oryza sativa Japonica Group]                      278    278
85%   5e-85  38.39  XP_015643947.1
hypothetical protein [Gossypium harknessii]                       277    277
89%   6e-85  37.83  MBA0819879.1
Lung seven transmembrane receptor-like [Trema orientale]          277    277
86%   7e-85  38.20  PON38231.1
protein GPR107-like [Lactuca sativa]                              277    277
88%   7e-85  38.04  XP_023754948.1
hypothetical protein E1A91_A11G019800v1 [Gossypium mustelinum]    277    277
84%   8e-85  38.69  TYJ07649.1
protein GPR107 [Carica papaya]                                    277    277
90%   9e-85  37.39  XP_021905116.1

Alignments:
>G protein coupled receptor, putative [Acanthamoeba castellanii str. Neff]
Sequence ID: XP_004338827.1 Length: 456

```
>G protein coupled receptor, putative [Acanthamoeba castellanii str. Neff]
Sequence ID: ELR16814.1 Length: 456
Range 1: 1 to 456

Score:934 bits(2414), Expect:0.0,
Method:Compositional matrix adjust.,
Identities:456/456(100%), Positives:456/456(100%), Gaps:0/456(0%)

Query  1    MMRDRRQTLMGCFLIFCLVAFLAPTANGLIHKLSIKNDRRLAFRIETFGFFTGGVMEMAI  60
            MMRDRRQTLMGCFLIFCLVAFLAPTANGLIHKLSIKNDRRLAFRIETFGFFTGGVMEMAI
Sbjct  1    MMRDRRQTLMGCFLIFCLVAFLAPTANGLIHKLSIKNDRRLAFRIETFGFFTGGVMEMAI  60

Query  61   ENFKVVDDKGSLWDDLSAGFIIKHIETDSGSFIEETDASKCVSLLDEPERGDTIAVKVTK  120
            ENFKVVDDKGSLWDDLSAGFIIKHIETDSGSFIEETDASKCVSLLDEPERGDTIAVKVTK
Sbjct  61   ENFKVVDDKGSLWDDLSAGFIIKHIETDSGSFIEETDASKCVSLLDEPERGDTIAVKVTK  120

Query  121  PSKENEKQSKVITIDKTRPEGFYTVVFLNCQPGTYVSFDLTLTNYNPGPNYLSAGLTALP  180
            PSKENEKQSKVITIDKTRPEGFYTVVFLNCQPGTYVSFDLTLTNYNPGPNYLSAGLTALP
Sbjct  121  PSKENEKQSKVITIDKTRPEGFYTVVFLNCQPGTYVSFDLTLTNYNPGPNYLSAGLTALP  180

Query  181  TLYAMLFVVWTVILGVWLFHFMRGQGKRIFRIHHLVTGIILLKLLTLLFEAIEFHYKKTT  240
            TLYAMLFVVWTVILGVWLFHFMRGQGKRIFRIHHLVTGIILLKLLTLLFEAIEFHYKKTT
Sbjct  181  TLYAMLFVVWTVILGVWLFHFMRGQGKRIFRIHHLVTGIILLKLLTLLFEAIEFHYKKTT  240

Query  241  GHPGGWVIAYYIFSGLKGTMMFVVIALIGTGWAFIKPFLGEKDKNIFLVVIPLQILANIA  300
            GHPGGWVIAYYIFSGLKGTMMFVVIALIGTGWAFIKPFLGEKDKNIFLVVIPLQILANIA
Sbjct  241  GHPGGWVIAYYIFSGLKGTMMFVVIALIGTGWAFIKPFLGEKDKNIFLVVIPLQILANIA  300

Query  301  IIVLEETAPEVLRLVDIICCGAILVPIIWSIKHLRDAAAIDGKAKRNMEKLKLFREFYLL  360
            IIVLEETAPEVLRLVDIICCGAILVPIIWSIKHLRDAAAIDGKAKRNMEKLKLFREFYLL
Sbjct  301  IIVLEETAPEVLRLVDIICCGAILVPIIWSIKHLRDAAAIDGKAKRNMEKLKLFREFYLL  360

Query  361  VVTYIYFTRIIVFLLDATLHYQYVWLGEFFTELATLIFWGLTGYKFRPVADNPYLKLDDE  420
            VVTYIYFTRIIVFLLDATLHYQYVWLGEFFTELATLIFWGLTGYKFRPVADNPYLKLDDE
Sbjct  361  VVTYIYFTRIIVFLLDATLHYQYVWLGEFFTELATLIFWGLTGYKFRPVADNPYLKLDDE  420

Query  421  EEDAEREEAQRQSRTEAGETIVMEPLEPKNIDVTIN  456
            EEDAEREEAQRQSRTEAGETIVMEPLEPKNIDVTIN
Sbjct  421  EEDAEREEAQRQSRTEAGETIVMEPLEPKNIDVTIN  456



>seven transmembrane domain protein [Heterostelium album PN500]
Sequence ID: XP_020432478.1 Length: 442
>seven transmembrane domain protein [Heterostelium album PN500]
Sequence ID: EFA80358.1 Length: 442
Range 1: 23 to 429

Score:376 bits(965), Expect:2e-123,
Method:Compositional matrix adjust.,
Identities:197/412(48%), Positives:271/412(65%), Gaps:21/412(5%)

Query  27   NGLIHKLSIKNDRRLAFRIETFGFFTGGVMEMAIENFKVVDDKGSLWDDLSAGFIIKHIE  86
            NG IH LS+K D R  F +ETFGF  GGVM + I N+K+ +   S     S GF I+  E
```

```
Sbjct  23   NGFIHHLSVKEDARTLFLVETFGFGMGGVMNLNITNWKINNQPISDETKSSVGFYIQISE  82

Query  87   TDSGSFIEETD--ASKCVSLLDEPERGDTIAVKVTKPSKENEKQSKV-ITIDKTRPEGFY  143
            TD+ +F++E    +  C  ++++     T    +   S +N   ++    I     EG++
Sbjct  83   TDASAFVDEATVASGNCQDMINK-----TFDYTIQYDSAKNPNITEFNFKIKPGFKEGYH  137

Query  144  TVVFLNCQPGTYVSFDLTLTNYNPGPN----YLSAGLTALPTLYAMLFVVWTVILGVWLF  199
             + FL+C   T VSFDL L  YN   N    YLS G T LPTLY +  + + VIL +WLF
Sbjct  138  NLYFLSCNRDTTVSFDLFLQEYNVDSNGQISYLSIGDTPLPTLYFVFAMAFFVILLLWLF  197

Query  200  HFMRGQGKRIFRIHHLVTGIILLKLLTLLFEAIEFHYKKTTGHPGGWVIAYYIFSGLKGT  259
             F+RG+GKR+ +IH+L    +L+  ++LLFE+IE HY K TG   GW IAYYIF+ ++G+
Sbjct  198  VFLRGEGKRVNKIHYLCAAYLLVLAISLLFESIEKHYIKKTGSAHGWNIAYYIFACIQGS  257

Query  260  MMFVVIALIGTGWAFIKPFLGEKDKNIFLVVIPLQILANIAIIVLEETAP---------E  310
               ++IALIG+GWAFIKPFL +KDK IF+VVIPLQIL NIA+++++E AP
Sbjct  258  FFVILIALIGSGWAFIKPFLSDKDKTIFMVVIPLQILDNIALVMIDEEAPGSIGSLSWRH  317

Query  311  VLRLVDIICCGAILVPIIWSIKHLRDAAAIDGKAKRNMEKLKLFREFYLLVVTYIYFTRI  370
            +  +VDIICC AI+VPIIWSIKHL+DA+ +D KA +NM+KLKLFR FYL+V++Y+YFTRI
Sbjct  318  IFTVVDIICCIAIIVPIIWSIKHLKDASQVDDKAAQNMQKLKLFRHFYLMVISYLYFTRI  377

Query  371  IVFLLDATLHYQYVWLGEFFTELATLIFWGLTGYKFRPVADNPYLKLDDEEE  422
               LL A+L ++Y W+G+F   LA+LIF+  TGY+FRP  DNPY  L  +E+
Sbjct  378  FFALLRASLPFRYAWIGDFSLLLASLIFYCSTGYQFRPSLDNPYFNLPKDED  429
```

>hypothetical protein SAMD00019534_116630 [Acytostelium subglobosum LB1]
Sequence ID: XP_012748526.1 Length: 442
>hypothetical protein SAMD00019534_116630 [Acytostelium subglobosum LB1]
Sequence ID: GAM28487.1 Length: 442
Range 1: 3 to 432

Score:367 bits(942), Expect:6e-120,
Method:Compositional matrix adjust.,
Identities:205/440(47%), Positives:280/440(63%), Gaps:27/440(6%)

```
Query  6    RQT--LMGCFLIFCLVAFLAPTANGLIHKLSIKNDRRLAFRIETFGFFTGGVMEMAIENF  63
            RQT   +   L+ CLV  L  T N  IH L+IK+D R  F IE+FG    GVM+ ++ N+
Sbjct  3    RQTSLFLSATLLICLV--LISTTNAFIHHLNIKDDIRQVFLIESFGLGKNGVMKCSVANW  60

Query  64   KVVDDKGSLWDDLSAGFIIKHIETDSGSFIEETDASK-CVSLLDEPERGDTIAVKVTKPS  122
            K+             GF IK  +TD+ + ++E    K CV   +      T
Sbjct  61   KLNGQPIDSSMSNGTGFFIKVTDTDASALVDEIILQKDCVHFFENQSFFFTY--------  112

Query  123  KENEKQSKVITIDKTR-PEGFYTVVFLNCQPGTYVSFDLTLTNYNPGPN----YLSAGLT  177
            K  +  +  +T+D     EGFY + F+NC  G  VSFDL+L  YN   N    YLS G T
Sbjct  113  KSGDTTNFDLTVDAPGFKEGFYNLHFINCNHGDKVSFDLSLEQYNVESNGQKSYLSIGDT  172

Query  178  ALPTLYAMLFVVWTVILGVWLFHFMRGQGKRIFRIHHLVTGIILLKLLTLLFEAIEFHYK  237
             LPTLY +   V+  +L +W+F F++G+GKR+ +IHHL T  +L++ ++LLFEAIE HY
Sbjct  173  PLPTLYGVFSAVFFGLLLLWVFFFLKGEGKRVNKIHHLCTAYLLIQSISLLFEAIEKHYI  232
```

```
Query  238  KTTGHPGGWVIAYYIFSGLKGTMMFVVIALIGTGWAFIKPFLGEKDKNIFLVVIPLQILA  297
            KTTG   GW +AYYIF+ L+G+   V+IALIG+GWAFIKP+L +KDK IF+VVIPLQIL
Sbjct  233  KTTGSAHGWNVAYYIFACLQGSFFIVLIALIGSGWAFIKPYLSDKDKTIFMVVIPLQILD  292

Query  298  NIAIIVLEETAP---------EVLRLVDIICCGAILVPIIWSIKHLRDAAAIDGKAKRNM  348
            NIA+++++E AP          +  +VDIICC AIL+PIIWSIKHL+DA+ +D KA +N+
Sbjct  293  NIALVMVDEEAPGSVGSIGWKHIFTVVDIICCIAILIPIIWSIKHLKDASQVDDKAAQNL  352

Query  349  EKLKLFREFYLLVVTYIYFTRIIVFLLDATLHYQYVWLGEFFTELATLIFWGLTGYKFRP  408
            +KL LFR+FYL V++YIYFTRII+ LL +TL +++VWLG+ F  +A LIF+  TGY+FRP
Sbjct  353  QKLTLFRQFYLFVISYIYFTRIIITLLRSTLPFKWVWLGDCFQLIAALIFYSTTGYQFRP  412

Query  409  VADNPYLKLDDEEEDAEREE  428
              DNPY  L   E+  +  E
Sbjct  413  SLDNPYFHLPQNEDGVQMSE  432
```

>GPR108 protein [Thecamonas trahens ATCC 50062]
Sequence ID: XP_013762250.1 Length: 451
>GPR108 protein [Thecamonas trahens ATCC 50062]
Sequence ID: KNC52248.1 Length: 451
Range 1: 10 to 436

Score:357 bits(915), Expect:1e-115,
Method:Compositional matrix adjust.,
Identities:188/435(43%), Positives:277/435(63%), Gaps:24/435(5%)

```
Query  10   MGCFLIFCLVAFLAPTANGLIHKLSIKNDRRLAFRIETFGFFTGGVMEMAIE-NFKVVDD  68
            +G F++  LVA  A   NGL    ++K D R AF +E FG+  GG +E+ +    ++  +
Sbjct  10   VGLFVVTALVAESAALRNGL----TMKQDGRAAFFVENFGYEAGGHLEVIVHPGVRMWVE  65

Query  69   KGSLWDDLSA-GFIIKHIETDSGSFIEETDASKCVSLLDEPER-GDTIAVKVTKPSKENE  126
            KG+        GF+I+  ++DS  +++E   S    ++ E  R  D +++       +
Sbjct  66   KGAYVSHTHPMGFVIRKTKSDSAEYVDEYGTSCLADIVPEDARERDIVSLANETTWIDGV  125

Query  127  KQSKVITIDKTRPEGFYTVVFLNCQPGTYVSFDLTLTNYNPGPNYLSAGLTALPTLYAML  186
               +VI   +   EG Y + + NC P T VS  + +  YNPGPNYLS+G + LPTLY +
Sbjct  126  HYKRVI---QPGEEGLYNLYWFNCAPATQVSLTVDVVMYNPGPNYLSSGKSPLPTLYVLF  182

Query  187  FVVWTVILGVWLFHFMRGQGKRIFRIHHLVTGIILLKLLTLLFEAIEFHYKKTTGHPGGW  246
                  I+GV L  FMR  G  ++++H+L+  + +LK LTLL  ++ +HY KTTGH  GW
Sbjct  183  TFAHFAIVGV-LVMFMRRAGAVVYKVHYLMAVVGVLKALTLLTTSLRYHYMKTTGHGSGW  241

Query  247  VIAYYIFSGLKGTMMFVVIALIGTGWAFIKPFLGEKDKNIFLVVIPLQILANIAIIVLEE  306
             + +YI  GL G M+F +IALIGTGWAF+KP+L ++DK +FLVVIPLQ+++NIA+I+++E
Sbjct  242  NVVFYILKGLTGLMLFTLIALIGTGWAFVKPYLTDRDKKVFLVVIPLQLISNIALIMVDE  301

Query  307  TAP---------EVLRLVDIICCGAILVPIIWSIKHLRDAAAIDGKAKRNMEKLKLFREF  357
              P         ++LR+VDI+CCG IL PI+WSIKHL+DA+  DGK + +  KLK+FR+F
Sbjct  302  MQPGSQSYLTWYDILRIVDIVCCGVILFPIVWSIKHLQDASGTDGKGRSSASKLKIFRQF  361

Query  358  YLLVVTYIYFTRIIVFLLDATLHYQYVWLGEFFTELATLIFWGLTGYKFRPVADNPYLKL  417
            Y+LVV +IYFTRIIV++L+AT+ ++Y+WL  F T  +TL F+ + GYKFRPV +NPYL L
```

```
Sbjct  362  YMLVVAWIYFTRIIVYMLEATVPFRYIWLSYFATLASTLAFYIVAGYKFRPVPNNPYLGL  421

Query  418  ----DDEEEDAEREE  428
                D E ED +  E
Sbjct  422  AGGSDVELEDFDGGE  436
```

>seven transmembrane domain protein [Cavenderia fasciculata]
Sequence ID: XP_004357792.1 Length: 433
>seven transmembrane domain protein [Cavenderia fasciculata]
Sequence ID: EGG19498.1 Length: 433
Range 1: 20 to 420

Score:356 bits(913), Expect:1e-115,
Method:Compositional matrix adjust.,
Identities:200/415(48%), Positives:278/415(66%), Gaps:32/415(7%)

```
Query  28   GLIHKLSIKNDRRLAFRIETFGFFTGGVMEMAIENFKVVDDKGSLWDDLSAGFIIKHIET  87
             L H L I+NDRR +F IE FGF +GGV+  ++ N+ +V+   S  +D +A F+IK  +T
Sbjct  20   SLKHHLKIENDRRQSFLIEGFGFSSGGVLNASVSNW-IVNGIPSTKNDNTA-FVIKISDT  77

Query  88   DSGSFIEETDASKCVSLLDEPERGDTIAVKVTKPSKENEKQSKVITI---DKTRPEGFYT  144
            D   ++E+ D   C S+ D         +  T  S+EN   S VI I    K  PEGFY
Sbjct  78   DGTLYLEDIDYQNC-SIPD---------IYFTNSSEENPF-SFVINIANDPKKYPEGFYN  126

Query  145  VVFLNCQPGTYVSFDLTLTNYNPGPN----YLSAGLTALPTLYAMLFVVWTVILGVWLFH  200
            + F+NC+    VSF L L  YN   N    YLS G   LPT+Y +  +++ V+L +W+F
Sbjct  127  LFFVNCK-AVPVSFTLDLVEYNIDSNGNINYLSVGDNPLPTVYGLFSIIFAVLLVLWIFV  185

Query  201  FMRGQGKRIFRIHHLVTGIILLKLLTLLFEAIEFHYKKTTGHPGGWVIAYYIFSGLKGTM  260
            F+RG+GKR+ +IHHL +  ++L+ + LLF +IE HY KTTG   GW IAYYIF+ L+G+
Sbjct  186  FLRGEGKRVNKIHHLCSVYLVLQAVELLFRSIEMHYIKTTGSANGWDIAYYIFACLQGSF  245

Query  261  MFVVIALIGTGWAFIKPFLGEKDKNIFLVVIPLQILANIAIIVLEETAP---------EV  311
              V+IALIGTGW FIKPFL +KDK IF++VIPLQIL NIA+I+++E AP         +
Sbjct  246  FIVLIALIGTGWTFIKPFLSDKDKTIFMIVIPLQILDNIALIMIDERAPGSESWVSWKHI  305

Query  312  LRLVDIICCGAILVPIIWSIKHLRDAAIDGKAKRNMEKLKLFREFYLLVVTYIYFTRII  371
              +VDIICCGAI+VPIIWSI HL+DAA+ D KA +++ KL+LFR+FY+ V++Y+YFTRI+
Sbjct  306  FTVVDIICCGAIIVPIIWSINHLKDAASADDKAAKSLAKLQLFRQFYVFVISYVYFTRIV  365

Query  372  VFLLDATLHYQYVWLGEFFTELATLIFWGLTGYKFRPVADNPYLKL--DDEEEDA  424
            VFL+ A+L +  +WLG+F T +A+L+F+  TGY+FRP  DNPY  +  DD+ E A
Sbjct  366  VFLVKASLEFHLIWLGDFSTLVASLVFYSATGYQFRPSLDNPYFNVPADDDLEAA  420
```

>seven transmembrane domain protein [Planoprotostelium fungivorum]
Sequence ID: PRP82284.1 Length: 558
Range 1: 113 to 536

Score:350 bits(899), Expect:7e-112,
Method:Compositional matrix adjust.,

```
Identities:194/441(44%), Positives:271/441(61%), Gaps:49/441(11%)

Query  26   ANGLIHKLSIKNDRRLAFRIETFGFFTGGVMEMAIENFKVVDDKGSLWDDL-SAGFIIKH  84
            + GLIH L I+ND R  F I++FGF  GG M M +  FK  D      D +   GF++ H
Sbjct  113  SEGLIHNLKIRNDPRQRFYIQSFGFGRGGFMTMKMSGFKASDGSSISEDAMKQTGFLMPH  172

Query  85   IETDSGSFIEETDASKCVSLLDEPERGDTIAVKVTKPSKENEKQSKVITIDKTRPEGFYT  144
             ETDS +  EE   SK   L       D   V  TK   E       ITI K + EG Y+
Sbjct  173  TETDSETLSEENIVSKDGCL-------DGATVGATKVYSEE------ITIAKGK-EGLYS  218

Query  145  VVFLNC--QPGTYVSFDLTLTNYN----PGPNYLSAGLTALPTLYAMLFVVWTVILGVWL  198
            + F++C  +P   + FDL L  YN       NYLS GL+ LPT+Y +  ++ + IL +W+
Sbjct  219  LYFVSCIKKP---IDFDLELIQYNVDKNNNKNYLSIGLSNLPTIYGVFTIIHSAILILWI  275

Query  199  FHFMRGQGKRIFRIHHLVTGIILLKLLTLLFEAIEFHYKKTTGHPGGWVIAYYIFSGLKG  258
            F F+   G  + ++H+ +T ++ LK+++L F+ I+ +Y KT G   GW I +YIF+ LKG
Sbjct  276  FGFLLPNGNEVNKLHYAMTLLVFLKIMSLFFQTIDQYYVKTKGSASGWNIPFYIFTSLKG  335

Query  259  TMMFVVIALIGTGWAFIKPFLGEKDKNIFLVVIPLQILANIAIIVLEETAP---------  309
              +FV+IALIG+G+ FIKPFL +++K IF +VI LQ+L NIA+I++EETAP
Sbjct  336  IALFVIIALIGSGYQFIKPFLHDREKKIFTIVITLQVLDNIALIIVEETAPGSVGWLSWQ  395

Query  310  EVLRLVDIICCGAILVPIIWSIKHLRDAAAIDGKAKRNMEKLKLFREFYLLVVTYIYFTR  369
             +L++VD+ICCGAI+VPI WSI+HLR+AA IDGKA  ++ KL LFR+FYL V++Y+YFTR
Sbjct  396  HILKIVDVICCGAIIVPIFWSIRHLREAAQIDGKAAVSVRKLSLFRQFYLTVISYVYFTR  455

Query  370  IIVFLLDATLHYQYVWLGEFFTELATLIFWGLTGYKFRP-VADNPYLKLD----------  418
            II++L++ATL YQ VWLG   +E ATL+F+ +TGY FRP V  N  L +D
Sbjct  456  IIIYLVEATLPYQMVWLGHMSSEAATLLFFVITGYNFRPNVELNKDLAIDLDEIEELGVV  515

Query  419  -----DEEEDAEREEAQRQSR  434
                 DEEE  +  E ++  R
Sbjct  516  IPLDSDEEEKEDHREDEKVGR  536
```

>seven transmembrane domain protein [Tieghemostelium lacteum]
Sequence ID: KYQ89580.1 Length: 442
Range 1: 21 to 436

Score:337 bits(865), Expect:3e-108,
Method:Compositional matrix adjust.,
Identities:192/425(45%), Positives:263/425(61%), Gaps:33/425(7%)

```
Query  28   GLIHKLSIKNDRRLAFRIETFGFFTGGVMEMAIENFKVVDDKGSLWDDL---SAGFIIKH  84
            G  H LSI+ND R  F IE FGF  GGV +  I N+K+  ++  L D     S GF IK
Sbjct  21   GYKHHLSIENDIRPNFHIEGFGFGKGGVFQATITNWKINGEQLDLNDSAMVASVGFEIKV  80

Query  85   IETDSGSFIEETDASKCVSLLDEPERGDTIAVKVTKPSKENEKQSKVITIDKT-RPEGFY  143
             + +    IE+   + C   L   +       ++T         +K I ID T   EG Y
Sbjct  81   TKIEDNQNIEDFTYTNCSERLKAADYTIFYTGEIT---------NKTIEIDGTDNIEGIY  131

Query  144  TVVFLNCQPGTYVSFDLTLTNYNPGP----NYLSAGLTALPTLYAMLFVVWTVILGVWLF  199
             + ++NC P    +F+L L  YN       +YL  GL+ LPT+Y +  + +  +L  W+
```

```
Sbjct  132  GLYYINCNPSATTTFNLVLEEYNVNSKGEISYLPIGLSPLPTVYVIFSIAFFALLAFWVL  191

Query  200  HFMRG---QGKRIFRIHHLVTGIILLKLLTLLFEAIEFHYKKTTGHPGGWVIAYYIFSGL  256
             F+RG    GKRI R+H L    +L++ +++LFE IE+HY KTTG P GW IAYYIF+ L
Sbjct  192  GFLRGGNLDGKRINRVHWLCAAYLLIQSISILFEGIEYHYIKTTGSPNGWNIAYYIFATL  251

Query  257  KGTMMFVVIALIGTGWAFIKPFLGEKDKNIFLVVIPLQILANIAIIVLEETAP-------  309
            +G    V+IALIGTGW FIKPFL ++DK IF++VIPLQIL NIA+I+++E +P
Sbjct  252  QGLFFIVLIALIGTGWYFIKPFLSDRDKTIFMIVIPLQILDNIALIMVDENSPGSIGWVS  311

Query  310  --EVLRLVDIICCGAILVPIIWSIKHLRDAAAIDGKAKRNMEKLKLFREFYLLVVTYIYF  367
               +L +VDI+CC AI  PI+WSIKHLRD +  D KA +N++KLKLFR+FYL VV Y+YF
Sbjct  312  WYHLLTVVDILCCIAITCPIVWSIKHLRDGSQGDDKAAQNLQKLKLFRQFYLFVVFYLYF  371

Query  368  TRIIVFLLDATLHYQYVWLGEFFTELATLIFWGLTGYKFRPVADNPYLKLDDEEED----  423
            TRIIV LL ATL ++ +W+G+F   LA++IF+   GY+FRP ADNPY +L   EE+
Sbjct  372  TRIIVILLKATLPFKLLWVGDFCLNLASVIFYAAVGYQFRPSADNPYFRLPTAEEEGVIL  431

Query  424  AEREE  428
            AER++
Sbjct  432  AERDK  436



>seven transmembrane domain protein [Dictyostelium discoideum AX4]
Sequence ID: XP_637699.2 Length: 449
>seven transmembrane domain protein [Dictyostelium discoideum AX4]
Sequence ID: EAL64194.2 Length: 449
Range 1: 20 to 444

Score:331 bits(848), Expect:1e-105,
Method:Compositional matrix adjust.,
Identities:173/432(40%), Positives:260/432(60%), Gaps:27/432(6%)

Query  27   NGLIHKLSIKNDRRLAFRIETFGFFTGGVMEMAIENFKVV--DDKGSLWDDLSAGFIIKH  84
            N   H L + ND R  F IE FGF   G   + + N+ +   D K +  DD    F ++
Sbjct  20   NAYKHHLEVNNDDRKLFAIEGFGFGDKGTFSIKVTNWTLNNGDGKKTKVDD-KVAFFLRI  78

Query  85   IETDSGSFIEETDASKCVSLLDEPE---RGDTIAVKVTKPSKENEKQSKVITIDKTRPEG  141
             ++D+   ++E   +KC    + P    RGD+  +       ++  Q   I+ +   PEG
Sbjct  79   TDSDTARHVDEYSVTKCDFESESPSLLFRGDSNVID------DSFSQQFKISAENDLPEG  132

Query  142  FYTVVFLNCQPG-TYVSFDLTLTNYN----PGPNYLSAGLTALPTLYAMLFVVWTVILGV  196
            FYT+ + NC P  +  +F L L  YN     G ++L  G   LPTLY    ++      V
Sbjct  133  FYTLYYKNCNPSPSKTTFKLVLEEYNIDSKGGISWLPIGNAPLPTLYTFFSLLLFATALV  192

Query  197  WLFHFMRGQGKRIFRIHHLVTGIILLKLLTLLFEAIEFHYKKTTGHPGGWVIAYYIFSGL  256
            W+   +RG+GKR+  +HHL T  ++++ + LLFEAIE HY K TG   GW +AYYIF+
Sbjct  193  WVLFCLRGEGKRVNLLHHLCTAYLVIQAIELLFEAIEQHYIKLTGSANGWNVAYYIFAFA  252

Query  257  KGTMMFVVIALIGTGWAFIKPFLGEKDKNIFLVVIPLQILANIAIIVLEETA--------  308
            +G+   +++A+IG+GW FIKPFL +K+K IF++VIPLQIL NIA+IV++E +
Sbjct  253  QGSFFIILLAMIGSGWYFIKPFLSDKEKKIFMIVIPLQILDNIALIVVDEESRGAANWIS  312
```

```
Query  309  -PEVLRLVDIICCGAILVPIIWSIKHLRDAA-AIDGKAKRNMEKLKLFREFYLLVVTYIY  366
               +   VD+ICC AI+VPI+WS+ HL+D+  + + K  +N++KL+LFR FYL+V+ Y+Y
Sbjct  313  WSNIFIFVDLICCLAIIVPIVWSMNHLKDSVDSNNDKVVKNIQKLRLFRSFYLVVICYLY  372

Query  367  FTRIIVFLLDATLHYQYVWLGEFFTELATLIFWGLTGYKFRPVADNPYLKLDDEEEDAER  426
            FTR+I++LL ATL Y+Y+WLGEFF+  A+  F+ +TGY+FRP  DNPY  L  +++D +
Sbjct  373  FTRVIIYLLKATLPYKYIWLGEFFSLTASFAFYCITGYQFRPSLDNPYFYLPTQDDDHDL  432

Query  427  EEAQRQSRTEAG  438
             E + Q   E G
Sbjct  433  NELRAQLEAEDG  444
```

>hypothetical protein KFL_002230120 [Klebsormidium nitens]
Sequence ID: GAQ85191.1 Length: 438
Range 1: 12 to 421

Score:316 bits(809), Expect:7e-100,
Method:Compositional matrix adjust.,
Identities:180/418(43%), Positives:260/418(62%), Gaps:24/418(5%)

```
Query  18   LVAFLAPTANGLIHKLSIKNDRRLAFRIETFGFFTGGVMEMAIENFKVVDDKGSLWDDLS  77
            L+++LA +   + H   ++ D R     ETFGF + G ++M + +  V   KG+   D S
Sbjct  12   LLSWLACSMAEISHT-HVERDPRPIILFETFGFDSTGHIDMNVSDDMVYIPKGAPDADQS  70

Query  78   A-GFIIKHIETDSGSFIEETDASKCVSLLDEP--ERGDTIAVKVTKPSKENEKQSKVITI  134
              GF I   E ++   IE  +   CV  LD P   +  T+A   T  +++    +    +
Sbjct  71   VMGFFITTAEAETQLQIELRNGG-CV--LDNPNIHKLFTLADVETGKNRDTGVYTHYYDV  127

Query  135  DKTRPEGFYTVVFLNCQPGTYVSFDLTLTNYNPGP----NYLSAGLTALPTLYAMLFVVW  190
               R    Y++ F NCQP + VS  + L  YN  P    +YL AG T LP L+ + F+++
Sbjct  128  PDARE---YSMFFANCQPNSVVSMKVRLAFYNIEPGGKKDYLPAGKTQLPKLFFLCFLIY  184

Query  191  TVILGVWLFHFMRGQGKRIFRIHHLVTGIILLKLLTLLFEAIEFHYKKTTGHPGGWVIAY  250
             +  GVW++   R +   + +IH L+ G++ LK LT L +A  + + K+TG P GW IA+
Sbjct  185  AIAAGVWIYICYRAKAT-VHKIHWLMGGLVFLKALTCLTQAGMYSWIKSTGQPDGWNIAF  243

Query  251  YIFSGLKGTMMFVVIALIGTGWAFIKPFLGEKDKNIFLVVIPLQILANIAIIVLEETAP-  309
            Y+FS  +G M+FVVI LIGTGW+F+KPFL +K+K + ++VIPLQ+LANIA+++L+ET P
Sbjct  244  YVFSFFRGIMLFVVIVLIGTGWSFLKPFLQDKEKKVMMIVIPLQVLANIAVVILDETGPS  303

Query  310  --------EVLRLVDIICCGAILVPIIWSIKHLRDAAAIDGKAKRNMEKLKLFREFYLLV  361
                    ++L LVDI+CC AIL PI+WSIKHLR+AA+ DGKA RN+ KL LFR+FY++V
Sbjct  304  NREWFTWRDILHLVDIVCCCAILFPIVWSIKHLREAASTDGKAARNLIKLTLFRQFYVMV  363

Query  362  VTYIYFTRIIVFLLDATLHYQYVWLGEFFTELATLIFWGLTGYKFRPVADNPYLKLDD  419
            V+YIYFTRI+V+LL +T  Y Y W  +   + A L+F+ LTGYKFRPV  NPY  LDD
Sbjct  364  VSYIYFTRIVVYLLKSTTPYHYAWTADLADQAAALLFYVLTGYKFRPVEQNPYFVLDD  421
```

>Transmembrane receptor, eukaryota [Nannochloropsis gaditana]
Sequence ID: EWM24893.1 Length: 505

Range 1: 23 to 451

Score:317 bits(812), Expect:1e-99,
Method:Compositional matrix adjust.,
Identities:179/433(41%), Positives:266/433(61%), Gaps:26/433(6%)

```
Query  18   LVAFLAPTANGLIHKLSIKNDRRLAFRIETFGFFTGGVMEMAIENFKVVDDKGSLWDDLS  77
            L+  LA  A+G++     +++    F++ETFGF  GGVME+   +F V   + +     +
Sbjct  23   LLGVLARVADGMVQNFHYEDETSPMFQLETFGFRPGGVMELNFNHFVVKVPEKTKDATVR  82

Query  78   AGFIIKHIETDSGS------FIEETDASKCVSLLDEPERGDTIAVKVTKPSKENEKQSKV  131
            AGF++   E+++ +       +E   A +    LD     D + + ++ P    EKQ +
Sbjct  83   AGFLMHLTESETTARQDLEEALERLAADQEDCWLDHRGPNDEV-IDLSDPESWAEKQVQH  141

Query  132  ITIDKTRPEGFYTVVFLNCQPG-TYVSFDLTLTNYNPGPNYLSAGLTALPTLYAMLFVVW  190
                    EG Y+++F+ CQP  + VSF +    YNPGPNYLSAG   LPTLY + F+ +
Sbjct  142  TVAPGE--EGLYSLIFVRCQPAPSAVSFKIHAKFYNPGPNYLSAGEAPLPTLYFVFFLCY  199

Query  191  TVILGVWLFHFMRGQGKRIFRIHHLVTGIILLKLLTLLFEAIEFH-YKKT--TGHPGGWV  247
             V +  W+    R + + + RIH+++  +++ K L+LLFEA+ F  Y+K   TGH  GW
Sbjct  200  LVAMAAWV-AVCRRRKEHVHRIHNMMLALLVFKTLSLLFEAVRFQAYQKHGLTGHADGWA  258

Query  248  IAYYIFSGLKGTMMFVVIALIGTGWAFIKPFLGEKDKNIFLVVIPLQILANIAIIVLEE-  306
            + YY+F+ +KG M FVVI LIGTGW+ +KP+L +++K + LVV+ LQ++ N A++VL+E
Sbjct  259  VVYYVFAFVKGIMRFVVILLIGTGWSLLKPYLSDREKKVVLVVLALQVIDNTAMVVLDEL  318

Query  307  --TAP---------EVLRLVDIICCGAILVPIIWSIKHLRDAAAIDGKAKRNMEKLKLFR  355
              T+P         ++  L+DI+CC AIL PI+WSI+HLR AAA DGK +  + KL LFR
Sbjct  319  SQTSPGSASWLTWRDIFHLIDIVCCCAILFPIVWSIRHLRQAAAADGKMEHILRKLTLFR  378

Query  356  EFYLLVVTYIYFTRIIVFLLDATLHYQYVWLGEFFTELATLIFWGLTGYKFRPVADNPYL  415
            +FYL+VV YIYFTRI+VFL+ ATL Y  +WL  FF E AT +F+ +TG+KF P   NPYL
Sbjct  379  QFYLMVVAYIYFTRIVVFLVKATLPYDLLWLQAFFDEGATFLFYTVTGWKFCPADANPYL  438

Query  416  KLDDEEEDAEREE  428
             +D +E DA   E
Sbjct  439  AVDTDERDAAELE  451
```

>PREDICTED: protein GPR107 [Ochotona princeps]
Sequence ID: XP_004593744.1 Length: 488
Range 1: 39 to 461

Score:313 bits(803), Expect:2e-98,
Method:Compositional matrix adjust.,
Identities:175/429(41%), Positives:254/429(59%), Gaps:29/429(6%)

```
Query  28   GLIHKLSIKNDRRLAFRIETFGFFTGGVMEMAIENFKVVDDKGSLWDDLSAGFIIKHIET  87
            G +H L++K+D R    + TFGFF  G M + + +  V + +G+   D + GF ++   +
Sbjct  39   GRVHHLALKDDVRHKVHLNTFGFFKDGYMAVNVSSLSVSEPEGATDKDAAIGFSLERTKN  98

Query  88   DSGSFIEETDASKCVSLLDEPERGDTIAVKVTKPSKENEKQSKV-----ITIDKTRPEGF  142
            D  S   + D + C+  L +     T+ + +  P  E      +     + I     EG
```

```
Sbjct  99   DGFSSYLDEDVNYCI--LKKQSVSVTLLI-LDIPRSEVPHFVPLHFQFFVNISTDDQEGQ  155

Query  143  YTVVFLNCQPGTYV--------SFDLTLTNYNPGPNYLSAGLTALPTLYAMLFVVWTVIL  194
            Y++ F  C PG  +        S D+ +T  NP  +YLSAG   LP LY  +   + +
Sbjct  156  YSLYFHKC-PGKELPAGGQFSFSLDIEITEKNPD-SYLSAGEIPLPKLYISMAFFFLLAG  213

Query  195  GVWLFHFMRGQGKRIFRIHHLVTGIILLKLLTLLFEAIEFHYKKTTGHP-GGWVIAYYIF  253
             VW+ H +R +   +F+IH L+  +   K L+L+F AI++HY  + G P  GW + YYI
Sbjct  214  TVWV-HILRKRRNDVFKIHWLMAALPFTKSLSLVFHAIDYHYISSQGFPIEGWAVVYYIT  272

Query  254  SGLKGTMMFVVIALIGTGWAFIKPFLGEKDKNIFLVVIPLQILANIAIIVLEETAP----  309
              LKG ++F+ IALIGTGWAFIK  L +KDK IF++VIPLQ+LAN+A I++E T
Sbjct  273  HLLKGALLFITIALIGTGWAFIKHILSDKDKKIFMIVIPLQVLANVAYIIIESTEAGTTE  332

Query  310  -----EVLRLVDIICCGAILVPIIWSIKHLRDAAAIDGKAKRNMEKLKLFREFYLLVVTY  364
                 + L LVD++CCGAIL P++WSI+HL++A+A DGKA  N+ KLKLFR +Y+L+V Y
Sbjct  333  YGLWKDSLFLVDLLCCGAILFPVVWSIRHLQEASATDGKAAINLAKLKLFRHYYVLIVCY  392

Query  365  IYFTRIIVFLLDATLHYQYVWLGEFFTELATLIFWGLTGYKFRPVADNPYLKLDDEEEDA  424
            IYFTRII FLL   + +Q+ WL +   E ATL+F+ LTGYKFRP +DNPYL+L  E++D
Sbjct  393  IYFTRIIAFLLKLAVPFQWKWLYQLLDETATLVFFILTGYKFRPASDNPYLQLSQEDDDL  452

Query  425  EREEAQRQS  433
            E E     S
Sbjct  453  EMESVVTAS  461
```

```
>PREDICTED: protein GPR107 isoform X1 [Galeopterus variegatus]
Sequence ID: XP_008577101.1 Length: 556
Range 1: 39 to 524

Score:310 bits(794), Expect:4e-96,
Method:Compositional matrix adjust.,
Identities:184/489(38%), Positives:261/489(53%), Gaps:91/489(18%)

Query  28   GLIHKLSIKNDRRLAFRIETFGFFTGGVMEMAIENFKVVDDKGSLWDDLSAGFIIKHIET  87
            G +H L++K+D R    + TFGFF  G M + + +  V + +G+   D + GF +    +
Sbjct  39   GRVHHLALKDDVRHKVHLNTFGFFKDGYMVVNVSSLSVHEPEGATHKDATIGFSLDRTKN  98

Query  88   DSGSFIEETDASKC--------VSLL----------------------------------  105
            D  S   + D + C        V+LL
Sbjct  99   DGFSSYLDEDVNYCILQKKAVSVTLLILDISRSEVLIRSPPEAGTQLPKIIFSKDEKVLG  158

Query  106  --DEPERGDTIAVKVTKPSKENEKQSKVITIDKTR-------------------------  138
               EP     +A   T+  +E+ K SK  T+D
Sbjct  159  QSQEPNVNPVLADNQTQKKQESGK-SKRSTVDSKAMGEKSFSVHNNDGKVSFQFFFNIST  217

Query  139  --PEGFYTVVFLNC-----QPGTYVSF--DLTLTNYNPGPNYLSAGLTALPTLYAMLFVV  189
               EG Y++ F  C     +PG   SF  D+ +T  NP  +YLSAG   LP LY  +
Sbjct  218  DDQEGLYSLYFHKCLRKELRPGDKYSFSLDIDITEKNPD-SYLSAGEIPLPKLYISMAFF  276

Query  190  WTVILGVWLFHFMRGQGKRIFRIHHLVTGIILLKLLTLLFEAIEFHYKKTTGHP-GGWVI  248
            + V   +W+ H +R +   +F+IH L+  +   K L+L+F AI++HY  + G P  GW +
```

```
Sbjct  277  FFVSGTIWI-HILRKRRNDVFKIHWLMAALPFTKSLSLVFHAIDYHYISSQGFPIEGWAV  335

Query  249  AYYIFSGLKGTMMFVVIALIGTGWAFIKPFLGEKDKNIFLVVIPLQILANIAIIVLEETA  308
             YYI   LKG ++F+ IALIGTGWAFIK  L +KDK IF++VIPLQ+LAN+A I++E T
Sbjct  336  VYYITHLLKGALLFITIALIGTGWAFIKHILSDKDKRIFMIVIPLQVLANVAYIIIESTE  395

Query  309  ---------PEVLRLVDIICCGAILVPIIWSIKHLRDAAAIDGKAKRNMEKLKLFREFYL  359
                      + L LVD++CCGAIL P++WSI+HL++A+A DGKA  N+ KLKLFR +Y+
Sbjct  396  EGTTEYGLWKDSLFLVDLLCCGAILFPVVWSIRHLQEASATDGKAAINLAKLKLFRHYYV  455

Query  360  LVVTYIYFTRIIVFLLDATLHYQYVWLGEFFTELATLIFWGLTGYKFRPVADNPYLKLDD  419
            L+V YIYFTRII FLL   + +Q+ WL +   E+ATL+F+ LTGYKFRP +DNPYL+L
Sbjct  456  LIVCYIYFTRIIAFLLKLAVPFQWKWLYQLLNEMATLVFFVLTGYKFRPASDNPYLQLSQ  515

Query  420  EEEDAEREE  428
            EE+D E E
Sbjct  516  EEDDLEMES  524



>hypothetical protein NGA_0212400 [Nannochloropsis gaditana CCMP526]
Sequence ID: XP_005854159.1 Length: 493
>hypothetical protein NGA_0212400 [Nannochloropsis gaditana CCMP526]
Sequence ID: EKU22201.1 Length: 493
Range 1: 23 to 439

Score:306 bits(784), Expect:2e-95,
Method:Compositional matrix adjust.,
Identities:178/433(41%), Positives:261/433(60%), Gaps:38/433(8%)

Query  18   LVAFLAPTANGLIHKLSIKNDRRLAFRIETFGFFTGGVMEMAIENFKVVDDKGSLWDDLS  77
            L+  LA  A+G++     +++    F++ETFGF  GGV E      K  D        +
Sbjct  23   LLGVLARVADGMVQNFHYEDETSPMFQLETFGFRPGGVPE------KTKDAT------VR  70

Query  78   AGFIIKHIETDSGS------FIEETDASKCVSLLDEPERGDTIAVKVTKPSKENEKQSKV  131
            AGF++   E+++ +       +E   A +    LD     D + + ++ P    EKQ +
Sbjct  71   AGFLMHLTESETTARQDLEEALERLAADQEDCWLDHRGPNDEV-IDLSDPESWAEKQVQH  129

Query  132  ITIDKTRPEGFYTVVFLNCQPG-TYVSFDLTLTNYNPGPNYLSAGLTALPTLYAMLFVVW  190
                    EG Y+++F+ CQP  + VSF +    YNPGPNYLSAG   LPTLY + F+ +
Sbjct  130  TVAPGE--EGLYSLIFVRCQPAPSAVSFKIHAKFYNPGPNYLSAGEAPLPTLYFVFFLCY  187

Query  191  TVILGVWLFHFMRGQGKRIFRIHHLVTGIILLKLLTLLFEAIEFH-YKKT--TGHPGGWV  247
             V +  W+    R + + + RIH+++  +++ K L+LLFEA+ F  Y+K   TGH  GW
Sbjct  188  LVAMAAWV-AVCRRRKEHVHRIHNMMLALLVFKTLSLLFEAVRFQAYQKHGLTGHADGWA  246

Query  248  IAYYIFSGLKGTMMFVVIALIGTGWAFIKPFLGEKDKNIFLVVIPLQILANIAIIVLEE-  306
            + YY+F+ +KG M FVVI LIGTGW+ +KP+L +++K + LVV+ LQ++ N A++VL+E
Sbjct  247  VVYYVFAFVKGIMRFVVILLIGTGWSLLKPYLSDREKKVVLVVLALQVIDNTAMVVLDEL  306

Query  307  --TAP---------EVLRLVDIICCGAILVPIIWSIKHLRDAAAIDGKAKRNMEKLKLFR  355
              T+P         ++  L+DI+CC AIL PI+WSI+HLR AAA DGK +  + KL LFR
Sbjct  307  SQTSPGSASWLTWRDIFHLIDIVCCCAILFPIVWSIRHLRQAAAADGKMEHILRKLTLFR  366
```

```
Query  356  EFYLLVVTYIYFTRIIVFLLDATLHYQYVWLGEFFTELATLIFWGLTGYKFRPVADNPYL  415
            +FYL+VV YIYFTRI+VFL+ ATL Y  +WL  FF E AT +F+ +TG+KF P   NPYL
Sbjct  367  QFYLMVVAYIYFTRIVVFLVKATLPYDLLWLQAFFDEGATFLFYTVTGWKFCPADANPYL  426

Query  416  KLDDEEEDAEREE  428
             +D +E DA   E
Sbjct  427  AVDTDERDAAELE  439
```

>hypothetical protein CEUSTIGMA_g1453.t1 [Chlamydomonas eustigma]
Sequence ID: GAX74003.1 Length: 431
Range 1: 7 to 430

Score:301 bits(772), Expect:1e-94,
Method:Compositional matrix adjust.,
Identities:177/437(41%), Positives:249/437(56%), Gaps:39/437(8%)

```
Query  15   IFCLVAFLAPTANGLI-HKLSIKNDRRLAFRIETFGFFTGGVMEMAIENFKVVDDKGSL-  72
            I     F  PT +  I H     +DR+L    E FGF  GG +   I +  +     +
Sbjct  7    ILVCALFQLPTLDAKITHSFVSVDDRKLIPLSEAFGFAVGGKLTFKISSIDIYQQHTTEV  66

Query  73   ---WDDLSAGFIIKHIETDSGSFIEETDASKCVSLLDEPERGDTIAVKVTKPSKENEKQS  129
               +D++  GF +  +E D+    +  D ++C+  L+  ER        + P   +
Sbjct  67   EPNYDNM--GFFLSPMEADTALEADLADDTQCI--LNTIERN-------SLPMFSDSAIR  115

Query  130  KVITIDKTRPE--------GFYTVVFLNCQPGTYVSFDLTLTNYNPGP----NYLSAGLT  177
            KVI    T  +        G + + F NC   T VSFD+ +  YN       NYLS G
Sbjct  116  KVIKRTATEADFEFTLENGGLFYLYFANCDKDTPVSFDMHIEMYNLDKDGHRNYLSVGEI  175

Query  178  ALPTLYAMLFVVWTVILGVWLFHFMRGQGKRIFRIHHLVTGIILLKLLTLLFEAIEFHYK  237
             L  LY ++F ++TVI G W   ++  Q +   RIH L+  +   K LT+L +AI  H+
Sbjct  176  ELEPLYWVMFSLFTVITGAWC-AYVWTQREHAQRIHFLMGVLCFFKSLTVLSQAIMTHHI  234

Query  238  KTTGHPGGWVIAYYIFSGLKGTMMFVVIALIGTGWAFIKPFLGEKDKNIFLVVIPLQILA  297
              TGH   GW IAYYIF+  +G + F VI LIGTGW+++KPFLG+++K I +VVIPLQ+ A
Sbjct  235  SLTGHADGWNIAYYIFTFFRGVLFFTVIVLIGTGWSYMKPFLGDREKRILMVVIPLQVFA  294

Query  298  NIAIIVLEETAP---------EVLRLVDIICCGAILVPIIWSIKHLRDAAAIDGKAKRNM  348
            NIAII+++E +P         +V  L+DI+CC AIL PI+WSIKHLR+A+  DGKA RN+
Sbjct  295  NIAIIIMDEESPSVRDWFTWRDVFHLIDIVCCCAILFPIVWSIKHLREASQTDGKAARNL  354

Query  349  EKLKLFREFYLLVVTYIYFTRIIVFLLDATLHYQYVWLGEFFTELATLIFWGLTGYKFRP  408
            EKL+LFR+FY++VV YIYFTRI+V+LL +T+  +Y WL E   +LATL F+  T   FRP
Sbjct  355  EKLQLFRQFYIMVVVYIYFTRIVVYLLRSTVS-EYEWLSEAADQLATLAFYVWTAVSFRP  413

Query  409  VADNPYLKLDDEEEDAE  425
             A NPYL+L   E + E
Sbjct  414  HAANPYLRLQTTESEIE  430
```

>protein GPR107 [Microcaecilia unicolor]
Sequence ID: XP_030063706.1 Length: 517

Range 1: 22 to 484

Score:303 bits(777), Expect:3e-94,
Method:Compositional matrix adjust.,
Identities:181/473(38%), Positives:255/473(53%), Gaps:81/473(17%)

```
Query  26   ANGLIHKLSIKNDRRLAFRIETFGFFTGGVMEMAIENFKVVDDKGSLWDDLSAGFIIKHI  85
            A   IH LS+K+D R    + TFGFF  G M + + +  +  D          GF +
Sbjct  22   AQARIHHLSLKDDVRRKVHLNTFGFFKNGFMVVNVNSLSI--DPA-----FQCGFSLDRT  74

Query  86   ETDSGSFIEETDASKCVSLLDEPERGDTIA----------VKVTK---------------  120
            + +     ++ D   C  L   PE+ ++I           VKVTK
Sbjct  75   KNEGFPTYQDEDVDYC-GLNRIPEQDESITFLKVDLTTNNVKVTKHNGKAATLPDITFKQ  133

Query  121  ---------PSKENEKQSKVITIDKTR-----------------------PEGFYTVVFL  148
                          E QSK I   KT                         EG Y+++F
Sbjct  134  PGNLSSTAKADASGETQSKSIIEGKTDNYSLSKAGGPVSFEFYFNVSSDDQEGLYSLLFH  193

Query  149  NCQPGT----YVSFDLTLTNYNPGPNYLSAGLTALPTLYAMLFVVWTVILGVWLFHFMRG  204
             CQ  T      S D+ +T  NP  +YLSA    LP LY  +  ++  + G+   H +R
Sbjct  194  ACQGNTGSRLLFSLDILITEKNP-ESYLSADEIPLPKLYISM-ALFFFLSGIIWAHILRK  251

Query  205  QGKRIFRIHHLVTGIILLKLLTLLFEAIEFHYKKTTGHP-GGWVIAYYIFSGLKGTMMFV  263
            +   +F+IH L+  +   K L+L+F AI++HY  + G+P  GW + YYI   LKG ++F+
Sbjct  252  RRHDVFKIHWLMAALPFTKSLSLVFHAIDYHYISSQGYPMEGWAVVYYIAHLLKGALLFI  311

Query  264  VIALIGTGWAFIKPFLGEKDKNIFLVVIPLQILANIAIIVLEETA---------PEVLRL  314
             IALIGTGWAF+K  L +KDK IF++VIPLQ+LAN+A I+LE T           E+L L
Sbjct  312  TIALIGTGWAFVKHILSDKDKKIFMIVIPLQVLANVAYIILESTEEGTTEYGLWKEILFL  371

Query  315  VDIICCGAILVPIIWSIKHLRDAAAIDGKAKRNMEKLKLFREFYLLVVTYIYFTRIIVFL  374
            VD++CCGAIL P++WSI+HL++A+A DGKA  N+ KLKLFR +Y+++V YIYFTRII  L
Sbjct  372  VDLLCCGAILFPVVWSIRHLQEASATDGKAAINLAKLKLFRHYYVMIVCYIYFTRIIAIL  431

Query  375  LDATLHYQYVWLGEFFTELATLIFWGLTGYKFRPVADNPYLKLDDEEEDAERE  427
            +   + +Q+ WL +  TE+ATL+F+ LTGYKFRP +DNPYL+L  EE+D E E
Sbjct  432  IKLAVPFQWKWLYQLLTEMATLLFFVLTGYKFRPTSDNPYLQLSQEEDDLEME  484
```

>G-protein coupled receptor [Chloropicon primus]
Sequence ID: QDZ20040.1 Length: 430
Range 1: 32 to 426

Score:300 bits(767), Expect:1e-93,
Method:Compositional matrix adjust.,
Identities:172/403(43%), Positives:237/403(58%), Gaps:26/403(6%)

```
Query  37   NDRRLAFRIETFGFFTGGVMEMAIENFKVVDDKGS-LWDDLSAGFIIKHIETDSGSFIEE  95
            +DR +    E FGF    GV+EM I    V   +GS   D    G ++    + D+
Sbjct  32   DDRSVILVAEPFGFNENGVIEMEISKATVFLPEGSPQADKTRIGVVLVTDDADAEVMRPS  91

Query  96   TDASKCVSLLDEPERGDTIAVKVTKPSKENEKQSKVITIDKTR----PEGFYTVVFLNCQ  151
            TD S C  LLDE    +   V  T    EN +    I   +        G Y++ F NC
```

```
Sbjct  92   TD-SGC--LLDE----ENARVLYTLEEVENNRAKDGIFYYRNSITEGSGGIYSLFFANCV  144

Query  152  PGTYVSFDLTLTNYNP----GPNYLSAGLTALPTLYAMLFVVWTVILGVWLFHFMRGQGK  207
              + V+  +  + YN     G +YLSAG   LP +Y  +F+    +LGVW    +  +
Sbjct  145  EDSDVTLSMATSLYNALPGGGKDYLSAGKKLLPGVYMFMFICSLAMLGVW-GSLLSKRRN  203

Query  208  RIFRIHHLVTGIILLKLLTLLFEAIEFHYKKTTGHPGGWVIAYYIFSGLKGTMMFVVIAL  267
             + RIH L+  +I+ K LTLL +    HY + TG P GW IAYY+F+ L+G M F VI L
Sbjct  204  HVQRIHFLMLVLIIFKALTLLSQYGMNHYIQKTGDPEGWNIAYYVFTFLRGLMFFTVIIL  263

Query  268  IGTGWAFIKPFLGEKDKNIFLVVIPLQILANIAIIVLEETAP---------EVLRLVDII  318
            +GTG+++ KPFL + +K + +VVIPLQ+LANIAII+++E +P         ++  L+DII
Sbjct  264  LGTGYSYFKPFLSDNEKKLLMVVIPLQVLANIAIIIMDEDSPADRDWFTWRDIFHLLDII  323

Query  319  CCGAILVPIIWSIKHLRDAAAIDGKAKRNMEKLKLFREFYLLVVTYIYFTRIIVFLLDAT  378
            CC A+L PI+WSIK LR+AA  DGKA RNM+KL LF++FY++VV Y+YFTRIIV+LL AT
Sbjct  324  CCCAVLFPIVWSIKRLREAATTDGKAARNMQKLALFKQFYIMVVAYVYFTRIIVYLLQAT  383

Query  379  LHYQYVWLGEFFTELATLIFWGLTGYKFRPVADNPYLKLDDEE  421
            L YQY WLG+   E+ATL F+  TG KF+P   NPYL L++ E
Sbjct  384  LPYQYTWLGDAAGEVATLAFYLTTGIKFQPAQHNPYLSLEEAE  426


>hypothetical protein CYY_001089 [Polysphondylium violaceum]
Sequence ID: KAF2077626.1 Length: 433
Range 1: 24 to 423

Score:299 bits(765), Expect:2e-93,
Method:Compositional matrix adjust.,
Identities:162/416(39%), Positives:243/416(58%), Gaps:31/416(7%)

Query  28   GLIHKLSIKNDRRLAFRIETFGFFTGGVMEMAIENFKVVDDKGSLWDDLSAGFIIKHIET  87
            G  H+L I+ D R  F IE+FGF  GGV ++ + N+K+    G+  +  + G
Sbjct  24   GFKHRLVIEKDNRDNFLIESFGFGEGGVFKLNVTNWKI---NGNPINQDTVG-------V  73

Query  88   DSGSFIEETDASKCVSLLDEPERGDTIAVKVTKPSKENEKQSKVITIDKTRPEGFYTVVF  147
             +G F ++T  ++ + +++     + I +  T  + + E  + V       PEG Y++ +
Sbjct  74   KAGFFTKKTLYAQEIPVMNCSVIENAIPLSSTGAAYDFEIDNDV------YPEGLYSLYY  127

Query  148  LNCQPGTYVSFDLTLTNYN---PGPNYLSAGLTALPTLYAMLFVVWTVILGVWLFHFMRG  204
             NC P    SF++ L  YN      +YL  G T LP+LY     ++  +L  W+F F+RG
Sbjct  128  YNCGPPAITSFEMVLEEYNIINGQISYLPLGSTNLPSLYGTFSAIFLALLLFWVFVFLRG  187

Query  205  QGKRIFRIHHLVTGIILLKLLTLLFEAIEFHYKKTTGHPGGWVIAYYIFSGLKGTMMFVV  264
            + KR   IHHL +  +L+  + L FEA ++H+ KTTG   GW +AYYIF+ ++GT   ++
Sbjct  188  EEKRTNIIHHLCSAYLLIHSIELFFEAFDYHFIKTTGSANGWNVAYYIFAIIQGTFFIIL  247

Query  265  IALIGTGWAFIKPFLGEKDKNIFLVVIPLQILANIAIIVLEETAP---------EVLRLV  315
             ALIGTGW F+KPFL + DK IF+++IPLQ+L NIA+++++E +P          +  +V
Sbjct  248  FALIGTGWKFVKPFLNDNDKIIFMIIIPLQVLDNIALVMIDEKSPGSIGWVSWYHLFIIV  307

Query  316  DIICCGAILVPIIWSIKHLRDAAAIDGKAKRNMEKLKLFREFYLLVVTYIYFTRIIVFLL  375
            D ICC +IL  I  SI HLR A  ++ K  +N++KL LFR++YL+V +Y+YFTRIIV L
```

```
Sbjct  308  DFICCLSILYLIYRSINHLRQAVDVNDKVNQNVQKLNLFRKYYLIVFSYVYFTRIIVALF  367

Query  376  DATLHYQYVWLGEFFTELATLIFWGLTGYKFRPVADNPYLKL---DDEEEDAEREE  428
              TL Y  VW+ +    +ATLIF+ +TGY FRP  DNPY  L    DD EE     E
Sbjct  368  RNTLQYTSVWISDVIFLVATLIFYAVTGYCFRPSLDNPYFNLPQDDDMEEGTNSVE  423
```

>hypothetical protein COCSUDRAFT_53024 [Coccomyxa subellipsoidea C-169]
Sequence ID: XP_005649187.1 Length: 414
>hypothetical protein COCSUDRAFT_53024 [Coccomyxa subellipsoidea C-169]
Sequence ID: EIE24643.1 Length: 414
Range 1: 18 to 407

Score:298 bits(762), Expect:3e-93,
Method:Compositional matrix adjust.,
Identities:163/399(41%), Positives:243/399(60%), Gaps:28/399(7%)

```
Query  46   ETFGFFTGGVMEMAIENFKVVDDKGSLWDDLSAGFII------KHIETDSGSFIEETDAS  99
             TFG  + G +++A+++F     KGS+ D    GF +        +E D        +A
Sbjct  18   NTFG--SDGYIDIAVKDFNYWH-KGSI-DLTQIGFFVTTSEAEAQLEVDLAQGTCAFEAE  73

Query  100  KCVSLLDEPERGDTIAVKVTKPSKENEKQSKVITIDKTRPEGFYTVVFLNCQPGTYVSFD  159
              + LL      D +       S+  +  +++  + K    G +++ + NCQ    VSFD
Sbjct  74   NVIKLLT----FDAVESNKAAGSELTQLSARLADLIKDYRGGEFSLFYANCQKLAVVSFD  129

Query  160  LTLTNYNPGPN----YLSAGLTALPTLYAMLFVVWTVILGVWLFHFMRGQGKRIFRIHHL  215
            + +  YN   N    +LS G   LPT++ ++FV+++V   +W     R Q K+  R+H+L
Sbjct  130  IRVALYNVRGNGIKDFLSVGEDMLPTVFMIMFVLFSVAAALWGMVLFR-QRKQAHRLHYL  188

Query  216  VTGIILLKLLTLLFEAIEFHYKKTTGHPGGWVIAYYIFSGLKGTMMFVVIALIGTGWAFI  275
            +  ++  K LTLL +A  +H  +TTGHP GW +A+Y+F+ L+G + F V+ L+GTGW+++
Sbjct  189  MMVLVAFKALTLLSQAGMYHLIRTTGHPEGWNVAFYVFNFLRGILFFTVVVLVGTGWSYM  248

Query  276  KPFLGEKDKNIFLVVIPLQILANIAIIVLEETAP---------EVLRLVDIICCGAILVP  326
             PFL +++K + +VVIPLQ+ A +AI++L+E  P         ++L LVDIICC AIL P
Sbjct  249  TPFLRDREKRLLMVVIPLQVCAEVAIVILDENTPASRSWFTWRDILHLVDIICCCAILFP  308

Query  327  IIWSIKHLRDAAAIDGKAKRNMEKLKLFREFYLLVVTYIYFTRIIVFLLDATLHYQYVWL  386
            I+WSIKHLR+AA  DGKA RN+ KL+LFR FY++VV YIYFTRI+V+LL +T+ YQYVWL
Sbjct  309  IVWSIKHLREAAGTDGKAARNLLKLQLFRHFYVMVVIYIYFTRIVVYLLRSTMPYQYVWL  368

Query  387  GEFFTELATLIFWGLTGYKFRPVADNPYLKLDDEEEDAE  425
             +   ELATL F+  T   FRP ADNPYL L D+E D +
Sbjct  369  SDGAGELATLAFYVTTAVSFRPTADNPYLHLADDEIDLQ  407
```

>hypothetical protein MARPO_0114s0018 [Marchantia polymorpha]
Sequence ID: PTQ31198.1 Length: 442
>hypothetical protein Mp_4g06350 [Marchantia polymorpha subsp. ruderalis]
Sequence ID: BBN07774.1 Length: 442
Range 1: 27 to 421

```
Score:296 bits(758), Expect:3e-92,
Method:Compositional matrix adjust.,
Identities:165/403(41%), Positives:229/403(56%), Gaps:23/403(5%)

Query  30   IHKLSIKNDRRLAFRIETFGFFTGGVMEMAIENFKVVDDKGSLWDDLSAGFIIKHIETDS  89
            I +  I++D R     E FGF   G + + + N +V  D  S  D    GF +   E
Sbjct  27   IRQSEIRDDNRQMIMFEKFGFDRNGKINITVSNVQVTMDDASEADYDMLGFFLTTEEDLV  86

Query  90   GSFIEETDASKCVSLLDEPERGDTIAVKVTKPSKENEKQSKVITIDKTRPEGF-YTVVFL  148
               +E  +      +L   +      +K         K ++  T   + P+   YT+ F
Sbjct  87   AVLVETEEQMSQTCVLKNSQIKTLFTLK-------EVKLTETFTRSLSPPDANEYTLFFA  139

Query  149  NCQPGTYVSFDLTLTNYN-----PGPNYLSAGLTALPTLYAMLFVVWTVILGVWLFHFMR  203
            NC   + VS D+    YN       P+YL AG T LP L+   FV++  +LGVW++  ++
Sbjct  140  NCLRRSQVSMDVRTAMYNLEGKSNTPDYLPAGQTQLPKLFFSFFVIYVFLLGVWIYTCVK  199

Query  204  GQGKRIFRIHHLVTGIILLKLLTLLFEAIEFHYKKTTGHPGGWVIAYYIFSGLKGTMMFV  263
             +   + RIH L+  ++ LK L L+ EA E  Y K TG   GW IA+YIF  L+G M+F
Sbjct  200  HRDT-VHRIHILMGVLVFLKALNLVAEATEKSYIKKTGLAHGWDIAFYIFGFLRGVMLFT  258

Query  264  VIALIGTGWAFIKPFLGEKDKNIFLVVIPLQILANIAIIVLEETAP---------EVLRL  314
            VI LIGTGW+F+KP+L EK+K + +VVIPLQ+ AN+A IV++ET P         +V  L
Sbjct  259  VIVLIGTGWSFLKPYLQEKEKKVLMVVIPLQVFANVASIVIDETGPSTKDWFTWKQVFLL  318

Query  315  VDIICCGAILVPIIWSIKHLRDAAAIDGKAKRNMEKLKLFREFYLLVVTYIYFTRIIVFL  374
            +DIICC A+L PI+WSIKHLR+A+  DGKA RN+ KL LFR +Y++VV+YIYFTRI+VF
Sbjct  319  LDIICCCAVLFPIVWSIKHLREASHTDGKAARNVVKLTLFRHYYMVVVSYIYFTRIVVFA  378

Query  375  LDATLHYQYVWLGEFFTELATLIFWGLTGYKFRPVADNPYLKL  417
            +     Y Y W  +   ELA+L F+  TGYKFRPV  NPY  L
Sbjct  379  VTTVTAYHYRWTSDLAEELASLAFYLFTGYKFRPVVHNPYFVL  421


>PREDICTED: protein GPR107 [Callorhinchus milii]
Sequence ID: XP_007901015.1 Length: 548
Range 1: 12 to 516

Score:299 bits(766), Expect:4e-92,
Method:Compositional matrix adjust.,
Identities:185/507(36%), Positives:264/507(52%), Gaps:94/507(18%)

Query  14   LIFCLVAFLAPTANGLIHKLSIKNDRRLAFRIETFGFFTGGVM--EMAIENFKVVDDK--  69
             +  L  F +     G IH L +KND R    + TFGFF  G M  EM     K +D K
Sbjct  12   FLLLLCGFFSVPVWGRIHHLILKNDVRHKVHLNTFGFFKHGSMTVEMTSLTLKGIDLKTI  71

Query  70   ------------------GSLWDDLSAGFIIKHIETD-------------------SGSF  92
                                L DD+    + K   +D                   SGS
Sbjct  72   DNSELGLSLDRTKNDGFSSYLEDDVDYCILKKKPPSDVPVTLLLFDFGKQRVTIQSSGSN  131

Query  93   IEETDASKCVSLLDEPERGDTIAVKVTKPSKENE---------------KQSKV------  131
              +   ++      E ++G+      + P+ E +               +QSK
Sbjct  132  QPQIGDARNAQSKPEEDKGNQKDKPGSGPADETQAANVQNISGKTKRSVEQSKSEAGSIP  191
```

```
Query  132  -------------ITIDKTRPEGFYTVVFLNC-----QPGTYVSFDL--TLTNYNPGPNY  171
                         ++I+ T  EG Y++ F NC     Q      FD+  T+   NPG ++
Sbjct  192  MQKKNSSYSVKFSLSINGTDEEGLYSLYFHNCYGKGQQNSELPRFDIDITIVEQNPG-SF  250

Query  172  LSAGLTALPTLYAMLFVVWTVILGVWLFHFMRGQGKRIFRIHHLVTGIILLKLLTLLFEA  231
            LSAG   LP LY  +   + +   +W++  +R +   +F+IH L+  +   K  +L+F A
Sbjct  251  LSAGEIPLPKLYISMAFFFFLAAVLWVY-ILRKRSNDVFKIHWLMGALAFTKSFSLVFHA  309

Query  232  IEFHYKKTTGHP-GGWVIAYYIFSGLKGTMMFVVIALIGTGWAFIKPFLGEKDKNIFLVV  290
            I++HY  T G P  GW + YYI   LKG ++F+ IALIGTGWAF+K  L +KDK IF++V
Sbjct  310  IDYHYISTQGFPIEGWAVVYYITHLLKGALLFITIALIGTGWAFVKHILSDKDKKIFMIV  369

Query  291  IPLQILANIAIIVLEETAP---------EVLRLVDIICCGAILVPIIWSIKHLRDAAAID  341
            IPLQ+LAN+A I++E T           E+L LVD++CCGAIL P++WSI+HL++A+A D
Sbjct  370  IPLQVLANVAYIIIESTEAGTSEYGLWKEILFLVDLLCCGAILFPVVWSIRHLQEASATD  429

Query  342  GKAKRNMEKLKLFREFYLLVVTYIYFTRIIVFLLDATLHYQYVWLGEFFTELATLIFWGL  401
            GKA  N+ KLKLFR +Y+++V YIYFTRII  L+  T+ +Q+ WL +F  ELATL+F+ L
Sbjct  430  GKAAINLAKLKLFRHYYVMIVCYIYFTRIIAILIKITVPFQWKWLYQFLDELATLVFFCL  489

Query  402  TGYKFRPVADNPYLKLDDEEEDAEREE  428
            TGYKFRP +DNPYL+L   EED E +E
Sbjct  490  TGYKFRPASDNPYLQLSQNEEDLEMDE  516
```

>predicted protein [Ostreococcus lucimarinus CCE9901]
Sequence ID: XP_001419678.1 Length: 433
>predicted protein [Ostreococcus lucimarinus CCE9901]
Sequence ID: XP_001422864.1 Length: 433 >predicted protein [Ostreococcus lucimarinus CCE9901]
Sequence ID: ABO97971.1 Length: 433 >predicted protein [Ostreococcus lucimarinus CCE9901]
Sequence ID: ABP01223.1 Length: 433
Range 1: 3 to 428

Score:295 bits(755), Expect:5e-92,
Method:Compositional matrix adjust.,
Identities:166/427(39%), Positives:243/427(56%), Gaps:19/427(4%)

```
Query  23   APTANGLIHKLSI-KNDRRLAFRIETFGFFTGGVMEMAIENFKV-VDDKGSLWDDLSAGF  80
            A TA   I    + K+DR L    E FGF  GG ME+A++  +V + +     D    GF
Sbjct  3    AATAEARIATYEVSKDDRSLILLREPFGFAVGGTMEIALKESRVYLPEHAPPLDKTKLGF  62

Query  81   IIKHIETDSGSFIEETDASKCVSLLDEPERGDTIAVKVTKPSKENEKQSKVITIDKTRPE  140
             I   + D G+     +   CV  +D  ++  T A    +  K   + S    + + +
Sbjct  63   FITEAK-DEGALDAALEDGACVLDVDFVDQAFTFADMDAQLKKGEREVSSYEFMKEIKVP  121

Query  141  GFYTVVFLNCQPGTYVSFDLTLTNYNPGP----NYLSAGLTALPTLYAMLFVVWTVILGV  196
            G Y + F +C P T VSF++T T  N       +YL AG  +LP++Y + F++ T
Sbjct  122  GDYMLFFASCAPHTVVSFEITTTFANKDARGRNDYLGAGEKSLPSVYYLFFLLDTCAAVA  181

Query  197  WLFHFMRGQGKR-IFRIHHLVTGIILLKLLTLLFEAIEFHYKKTTGHPGGWVIAYYIFSG  255
            W F   R  G+R + +IH L+  ++  K L++L +A  +H  + TG   GW  AYY+F+
```

```
Sbjct  182  WAFILGRANGQRGVRKIHWLMLALVCFKTLSVLAQAGRYHITRLTGSSRGWTAAYYVFTV  241

Query  256  LKGTMMFVVIALIGTGWAFIKPFLGEKDKNIFLVVIPLQILANIAIIVLEETAP------  309
             +  +MF VIAL+G GW+F+KPFL +++KN+ + VIPLQ+LANIA +V+ E  P
Sbjct  242  CRSMLMFSVIALVGMGWSFLKPFLHQREKNLLMAVIPLQVLANIAAVVIGEEGPADQGWF  301

Query  310  ---EVLRLVDIICCGAILVPIIWSIKHLRDAAAIDGKAKRNMEKLKLFREFYLLVVTYIY  366
                +  ++DI+CC A+LVPI+WSIKHLRD +    K  RN+EKL LFR FY++ V YIY
Sbjct  302  AWQNMFIVIDIMCCCAVLVPIVWSIKHLRDTSDSSEKKARNLEKLLLFRHFYVMTVAYIY  361

Query  367  FTRIIVFLLDATLHYQYVWLGEFFTELATLIFWGLTGYKFRPVADNPY--LKLDDEEEDA  424
            FTRIIV+LL +T+ Y+  W   FF ELATL ++  TGY FRP  +N Y  LK D++E+
Sbjct  362  FTRIIVYLLKSTVEYELRWTAAFFNELATLAYYVATGYMFRPEEENMYFSLKHDEDEDGV  421

Query  425  EREEAQR  431
            E  +  R
Sbjct  422  EMSDRAR  428
```

>uncharacterized protein HaLaN_07828 [Haematococcus lacustris]
Sequence ID: GFH12190.1 Length: 360
Range 1: 65 to 357

Score:292 bits(747), Expect:1e-91,
Method:Compositional matrix adjust.,
Identities:147/294(50%), Positives:198/294(67%), Gaps:14/294(4%)

```
Query  141  GFYTVVFLNCQPGTYVSFDLTLTNYN----PGPNYLSAGLTALPTLYAMLFVVWTVILGV  196
            G + + F NC+  T VSFD  +  YN       NY+S G T +  +Y ++F ++T I G
Sbjct  65   GLFYLYFANCERSTPVSFDSLIEMYNVDERGNKNYMSVGETEMEVVYWVMFGLFTAIGGA  124

Query  197  WLFHFMRGQGKRIFRIHHLVTGIILLKLLTLLFEAIEFHYKKTTGHPGGWVIAYYIFSGL  256
            W++ F+    +   RIH+L+  + L K LT++ +A  FHY + TGH  GW IAYYIF+
Sbjct  125  WIW-FVWTNRQHSHRIHYLMGLLCLFKALTVMSQAGMFHYVERTGHADGWNIAYYIFTFC  183

Query  257  KGTMMFVVIALIGTGWAFIKPFLGEKDKNIFLVVIPLQILANIAIIVLEETAP-------  309
            +G + F V+ LIGTGW+++KPFLG+K+K I +VVIPLQ+ ANIAII+ EE +P
Sbjct  184  RGVLFFTVVVLIGTGWSYMKPFLGDKEKRILMVVIPLQVFANIAIIITEEESPAIKDWFT  243

Query  310  --EVLRLVDIICCGAILVPIIWSIKHLRDAAAIDGKAKRNMEKLKLFREFYLLVVTYIYF  367
              +V  LVDIICC AIL PI+WSIKHLR+A+  DGKA RN+EKL LFR+FY++VV YIYF
Sbjct  244  WRDVFHLVDIICCCAILFPIVWSIKHLREASQTDGKAARNLEKLTLFRQFYVMVVVYIYF  303

Query  368  TRIIVFLLDATLHYQYVWLGEFFTELATLIFWGLTGYKFRPVADNPYLKLDDEE  421
            TRI+V+LL  T+ Y Y W+ E   +LATL F+  T  KFRP  +NPYLKL+  E
Sbjct  304  TRIVVYLLMRTMQYDYAWVSEAADQLATLAFYVWTAVKFRPQPNNPYLKLETAE  357
```

>protein GPR107 [Ricinus communis]
Sequence ID: XP_025013062.1 Length: 432
>lung seven transmembrane receptor, putative [Ricinus communis]
Sequence ID: EEF43452.1 Length: 432

Range 1: 25 to 414

Score:293 bits(751), Expect:3e-91,
Method:Compositional matrix adjust.,
Identities:162/409(40%), Positives:239/409(58%), Gaps:43/409(10%)

```
Query  35   IKNDRRLAFRIETFGFFTGGVMEMAIENFKVVDDKGSLWDDLS-AGF----------IIK  83
            I+ND R     + FGF   G +E+ + N  + +    L  DLS  GF          +I+
Sbjct  25   IRNDDRPIIPFDEFGFTHKGQLELNVSNIHLSNPNPDL--DLSRVGFFLCTRESWLHVIQ  82

Query  84   HIETDSGSFIEETDASKCVSLLDEPERGDTIAVKVTKPSKENEKQSKVITIDKTRPEGFY  143
             +E    +   ++D  K V   ++ ++G T    VT  +  ++                Y
Sbjct  83   QLEDGEIACALQSDLIKQVYTFNKLQKGQTFFSVVTSENDADQ----------------Y  126

Query  144  TVVFLNCQPGTYVSFDLTLTNYNPGPN----YLSAGLTALPTLYAMLFVVWTVILGVWLF  199
            T+VF NC     VS D+  T YN   N    YLSAG T LP +Y +L +V+ V+ G+W+
Sbjct  127  TLVFANCLNSVKVSMDVKSTMYNIERNGNRDYLSAGKTILPRVYFLLSIVYFVMAGLWI-  185

Query  200  HFMRGQGKRIFRIHHLVTGIILLKLLTLLFEAIEFHYKKTTGHPGGWVIAYYIFSGLKGT  259
            H +  +   +FRIH  +  ++++K L LLFEA +  Y K TG   GW + +YIFS  KG
Sbjct  186  HVLYKKRLTVFRIHFFMLAVVIMKALNLLFEAEDKSYIKRTGSAHGWDVLFYIFSFFKGI  245

Query  260  MMFVVIALIGTGWAFIKPFLGEKDKNIFLVVIPLQILANIAIIVLEETAP---------E  310
             +F +I LIGTGW+F+KP+L +K+K + ++VIPLQ++ANIA +V++ET P         +
Sbjct  246  ALFTLIVLIGTGWSFLKPYLQDKEKKVLMIVIPLQVIANIAQVVIDETGPYGQDWITWKQ  305

Query  311  VLRLVDIICCGAILVPIIWSIKHLRDAAAIDGKAKRNMEKLKLFREFYLLVVTYIYFTRI  370
            V  LVD++CC A+L PI+WSIK+LR+AA  DGKA  N+ KL LFR++Y++V+ YIYFTR+
Sbjct  306  VFLLVDVVCCCAVLFPIVWSIKNLREAARTDGKAAVNLMKLTLFRQYYIVVICYIYFTRV  365

Query  371  IVFLLDATLHYQYVWLGEFFTELATLIFWGLTGYKFRPVADNPYLKLDD  419
            +V+ L+    Y+Y+W      ELATL F+  TGYKF+P A NPY  +DD
Sbjct  366  VVYALETITSYRYLWTSVVAGELATLAFYAFTGYKFKPEAHNPYFVIDD  414
```

>hypothetical protein AXG93_1838s1160 [Marchantia polymorpha subsp. ruderalis]
Sequence ID: OAE31806.1 Length: 565
Range 1: 155 to 544

Score:296 bits(759), Expect:7e-91,
Method:Compositional matrix adjust.,
Identities:164/398(41%), Positives:227/398(57%), Gaps:23/398(5%)

```
Query  35   IKNDRRLAFRIETFGFFTGGVMEMAIENFKVVDDKGSLWDDLSAGFIIKHIETDSGSFIE  94
            I++D R     E FGF   G + + + N +V  D  S  D    GF +   E      +E
Sbjct  155  IRDDNRQMIMFEKFGFDRNGKINITVSNVQVTMDDASEADYDMLGFFLTTEEDLVAVLVE  214

Query  95   ETDASKCVSLLDEPERGDTIAVKVTKPSKENEKQSKVITIDKTRPEGF-YTVVFLNCQPG  153
              +      +L   +      +K  K        ++  T   + P+   YT+ F NC
Sbjct  215  TEEQMSQTCVLKNSQIKTLFTLKEVK-------LTETFTRSLSPPDANEYTLFFANCLRR  267

Query  154  TYVSFDLTLTNYN-----PGPNYLSAGLTALPTLYAMLFVVWTVILGVWLFHFMRGQGKR  208
            + VS D+    YN       P+YL AG T LP L+   FV++  +LGVW++  ++ +
```

```
Sbjct  268   SQVSMDVRTAMYNLEGKSNTPDYLPAGQTQLPKLFFSFFVIYVFLLGVWIYTCVKHRDT-  326

Query  209   IFRIHHLVTGIILLKLLTLLFEAIEFHYKKTTGHPGGWVIAYYIFSGLKGTMMFVVIALI  268
             + RIH L+  ++ LK L L+ EA E  Y K TG   GW IA+YIF  L+G M+F VI LI
Sbjct  327   VHRIHILMGVLVFLKALNLVAEATEKSYIKKTGLAHGWDIAFYIFGFLRGVMLFTVIVLI  386

Query  269   GTGWAFIKPFLGEKDKNIFLVVIPLQILANIAIIVLEETAP---------EVLRLVDIIC  319
             GTGW+F+KP+L EK+K + +VVIPLQ+ AN+A IV++ET P         +V  L+DIIC
Sbjct  387   GTGWSFLKPYLQEKEKKVLMVVIPLQVFANVASIVIDETGPSTKDWFTWKQVFLLLDIIC  446

Query  320   CGAILVPIIWSIKHLRDAAAIDGKAKRNMEKLKLFREFYLLVVTYIYFTRIIVFLLDATL  379
             C A+L PI+WSIKHLR+A+  DGKA RN+ KL LFR +Y++VV+YIYFTRI+VF +
Sbjct  447   CCAVLFPIVWSIKHLREASHTDGKAARNVVKLTLFRHYYMVVVSYIYFTRIVVFAVTTVT  506

Query  380   HYQYVWLGEFFTELATLIFWGLTGYKFRPVADNPYLKL  417
              Y Y W  +   ELA+L F+  TGYKFRPV  NPY  L
Sbjct  507   AYHYRWTSDLAEELASLAFYLFTGYKFRPVVHNPYFVL  544
```

>protein GPR107-like [Physcomitrium patens]
Sequence ID: XP_024386861.1 Length: 429
>hypothetical protein PHYPA_013976 [Physcomitrium patens]
Sequence ID: PNR46856.1 Length: 429
Range 1: 17 to 409

Score:291 bits(746), Expect:1e-90,
Method:Compositional matrix adjust.,
Identities:172/408(42%), Positives:230/408(56%), Gaps:27/408(6%)

```
Query  22    LAPTANGLIHKLSIKNDRRLAFRIETFGFFTGGVMEMAIENFKVVDDKGSLWDDLSAGFI  81
             L P  +G I  LSIKND R     ETFGF  GG + + + N KV D    L      GF
Sbjct  17    LMPLVSGEIKDLSIKNDARPIIPFETFGFSVGGEVSIEVTNVKVKDPAADLQ---RMGFF  73

Query  82    IKHIETDSGSFIEETDASKCVSLLDEPERGDTIAVKVTKPSKENEKQSKVITIDKTRPEG  141
             +  ++ D    I + +      LLD           +  P K N          +
Sbjct  74    LSTMD-DLVPVISQLEGLGKNCLLDSNLMHTLFT--MADPVKANFTH-------QVMQSN  123

Query  142   FYTVVFLNCQPGTYVSFDLTLTNYNPGPNY---LSAGLTALPTLYAMLFVVWTVILGVWL  198
              YT+ F NC   T VS ++  + YN    Y   L  G T LP +Y + F+    +LG W+
Sbjct  124   EYTLFFENCVK-TEVSMEVKTSMYNVENGYRDFLPVGQTVLPRMYFLFFLANVALLGTWI  182

Query  199   FHFMRGQGKRIFRIHHLVTGIILLKLLTLLFEAIEFHYKKTTGHPGGWVIAYYIFSGLKG  258
             +     Q + + RIH L+  +++LK L ++ EA +  Y K TG P GW +AYYIF+ L+G
Sbjct  183   YA-CWAQKESVHRIHFLMGLLVVLKALNMICEAEDKQYVKRTGTPHGWDVAYYIFNFLRG  241

Query  259   TMMFVVIALIGTGWAFIKPFLGEKDKNIFLVVIPLQILANIAIIVLEETAP---------  309
              ++F VI LIGTGW+F+KPFL +K+K + +VVIPLQ+ AN A IV +E+ P
Sbjct  242   LLLFSVIVLIGTGWSFLKPFLQDKEKKVLMVVIPLQVFANTAYIVYDESGPSTKDFFTWK  301

Query  310   EVLRLVDIICCGAILVPIIWSIKHLRDAAAIDGKAKRNMEKLKLFREFYLLVVTYIYFTR  369
             +V  L+DIICC A+L PI+WSIKHLR+AA  DGKA RN+ KL LFR+FY++VV+YIYFTR
Sbjct  302   QVFLLLDIICCCAVLFPIVWSIKHLREAARTDGKAARNLAKLTLFRQFYIVVVSYIYFTR  361
```

```
Query  370  IIVFLLDATLHYQYVWLGEFFTELATLIFWGLTGYKFRPVADNPYLKL  417
            I+VF L     Y+Y W  +F  E A L F+  TGYKFRPV  NPY  L
Sbjct  362  IVVFALFTVTVYRYQWTSKFAEEAANLAFYIYTGYKFRPVVHNPYFVL  409
```

>protein GPR107 isoform X2 [Cimex lectularius]
Sequence ID: XP_014261704.1 Length: 457
>protein GPR107 isoform X2 [Cimex lectularius]
Sequence ID: XP_024084687.1 Length: 457
Range 1: 7 to 415

Score:291 bits(746), Expect:3e-90,
Method:Compositional matrix adjust.,
Identities:167/423(39%), Positives:248/423(58%), Gaps:40/423(9%)

```
Query  30   IHKLSIKNDRRLAFRIETFGFFT---------GGVMEMAIENFKVVDDKGS---LWDDLS  77
            IH++   N+ R   R+ +  F               E+ +  FK      +   L D LS
Sbjct  7    IHQVHCLNEPRNPVRVPSVLFIMDRKNNVLRINSTYELKLHIFKDNSQLAAFLRLRDSLS  66

Query  78   AGFIIKHIETDSGSFIEETDASKCVSLLDEPERGDTIAVKVTKPSKENEKQSKVITIDKT  137
               +     T +G+F + + A          E   TI +   +P   N   S VI + +
Sbjct  67   DDVLFSTRGTVNGNFPKRSIA----------EEEQTIPLIQPQPGYYN--TSFVIYVSRE  114

Query  138  RPEGFYTVVFLNCQ---PGTY-VSFDLTLTNYNPGPNYLSAGLTALPTLYAMLFVVWTVI  193
            + EG Y + F NC+   P    V F++ +   N   NYLSAG   LP LY+M+ +++ +
Sbjct  115  KDEGLYNLYFFNCRNYRPNPIPVEFEIDIVEKN-KENYLSAGEMPLPALYSMMSLLFFLS  173

Query  194  LGVWLFHFMRGQGKRIFRIHHLVTGIILLKLLTLLFEAIEFHYKKTTG-HPGGWVIAYYI  252
               W+F  ++ +   +F+IH+L+  ++ LK L+LLF AI +H+ +  G H   W I YYI
Sbjct  174  GCFWVFILVKSK-HTVFKIHYLMATLVYLKSLSLLFHAINYHFIEKKGEHVAAWAILYYI  232

Query  253  FSGLKGTMMFVVIALIGTGWAFIKPFLGEKDKNIFLVVIPLQILANIAIIVLEETAP---  309
               LKG+++F+ I LIGTGW FIK  L E+DK IF++VIPLQ+LAN+  I++EE+
Sbjct  233  THLLKGSVLFITIVLIGTGWTFIKHLLFERDKKIFMIVIPLQVLANVVKIIIEESEEGDV  292

Query  310  ------EVLRLVDIICCGAILVPIIWSIKHLRDAAAIDGKAKRNMEKLKLFREFYLLVVT  363
                  +V  LVD++CCGAILVP++WSI++L++AA IDGKA  N+ KLKLFR FY+LVV
Sbjct  293  EHRTWLDVFILVDLLCCGAILVPVVWSIRYLQEAACIDGKAAINLRKLKLFRHFYILVVC  352

Query  364  YIYFTRIIVFLLDATLHYQYVWLGEFFTELATLIFWGLTGYKFRPVADNPYLKLDDEEED  423
            YIY TRI+V+LL+ T+ +QY WL E F E+ T +F+ LTGYKFRP + NPY ++++E++D
Sbjct  353  YIYVTRIVVYLLEMTVSFQYEWLDEMFKEMGTYVFFVLTGYKFRPASANPYFQVNNEDDD  412

Query  424  AER  426
             E
Sbjct  413  TEE  415
```

>PREDICTED: protein GPR107 isoform X2 [Galeopterus variegatus]
Sequence ID: XP_008577102.1 Length: 490
Range 1: 148 to 458

Score:292 bits(748), Expect:3e-90,
Method:Compositional matrix adjust.,
Identities:150/313(48%), Positives:205/313(65%), Gaps:19/313(6%)

```
Query  133  TIDKTRPEGFYTVVFLNC-----QPGTYVSF--DLTLTNYNPGPNYLSAGLTALPTLYAM  185
             I     EG Y++ F  C     +PG   SF  D+ +T  NP  +YLSAG   LP LY
Sbjct  148  NISTDDQEGLYSLYFHKCLRKELRPGDKYSFSLDIDITEKNPD-SYLSAGEIPLPKLYIS  206

Query  186  LFVVWTVILGVWLFHFMRGQGKRIFRIHHLVTGIILLKLLTLLFEAIEFHYKKTTGHP-G  244
            +   + V   +W+ H +R +   +F+IH L+  +   K L+L+F AI++HY  + G P
Sbjct  207  MAFFFFVSGTIWI-HILRKRRNDVFKIHWLMAALPFTKSLSLVFHAIDYHYISSQGFPIE  265

Query  245  GWVIAYYIFSGLKGTMMFVVIALIGTGWAFIKPFLGEKDKNIFLVVIPLQILANIAIIVL  304
            GW + YYI   LKG ++F+ IALIGTGWAFIK  L +KDK IF++VIPLQ+LAN+A I++
Sbjct  266  GWAVVYYITHLLKGALLFITIALIGTGWAFIKHILSDKDKRIFMIVIPLQVLANVAYIII  325

Query  305  EETA---------PEVLRLVDIICCGAILVPIIWSIKHLRDAAAIDGKAKRNMEKLKLFR  355
            E T           + L LVD++CCGAIL P++WSI+HL++A+A DGKA  N+ KLKLFR
Sbjct  326  ESTEEGTTEYGLWKDSLFLVDLLCCGAILFPVVWSIRHLQEASATDGKAAINLAKLKLFR  385

Query  356  EFYLLVVTYIYFTRIIVFLLDATLHYQYVWLGEFFTELATLIFWGLTGYKFRPVADNPYL  415
             +Y+L+V YIYFTRII FLL   + +Q+ WL +   E+ATL+F+ LTGYKFRP +DNPYL
Sbjct  386  HYYVLIVCYIYFTRIIAFLLKLAVPFQWKWLYQLLNEMATLVFFVLTGYKFRPASDNPYL  445

Query  416  KLDDEEEDAEREE  428
            +L  EE+D E E
Sbjct  446  QLSQEEDDLEMES  458
```

>protein GPR107 isoform X1 [Rhincodon typus]
Sequence ID: XP_020386814.1 Length: 556
Range 1: 127 to 524

Score:294 bits(753), Expect:4e-90,
Method:Compositional matrix adjust.,
Identities:173/408(42%), Positives:247/408(60%), Gaps:41/408(10%)

```
Query  52   TGGVMEMAIENFKV-----VDDKGSLWDDLS------AGFIIKHIETDSGSFIEETDASK  100
            T G  +  IE+F+      V DK S  D  S      AG + K  ET   S  E+   +K
Sbjct  127  TSGNNQPTIEDFRTAASQPVKDKSSKTDVKSQSESKKAGELQKKNETTKQS--EQVQMAK  184

Query  101  CVSLLDEPERGDTIAVKVTKPSKENEKQSKVI--TIDKTRPEGFYTVVFLNC------QP  152
              S +    + +T+ ++     K+N   S     TI+    +G Y++ F NC       P
Sbjct  185  PKSGISAERKDETLPLQ-----KKNLSYSFTFSFTINSEEEQGLYSLYFHNCYTDEKPSP  239

Query  153  GTY-VSFDLTLTNYNPGPNYLSAGLTALPTLY-AMLFVVWTVILGVWLFHFMRGQGKRIF  210
             T+    D+T+   NPG +YLSAG   LP LY  M F  +  + GV+  + +R +   +F
Sbjct  240  QTFQFDMDITIVEQNPG-SYLSAGEIPLPKLYICMAFFFF--LAGVFWVYILRKRSSDVF  296

Query  211  RIHHLVTGIILLKLLTLLFEAIEFHYKKTTGHP-GGWVIAYYIFSGLKGTMMFVVIALIG  269
            +IH L+  +   K L+L+F AI++H+  T G+P  GW + YYI   LKG ++F+ I+LIG
Sbjct  297  KIHWLMGALAFTKSLSLVFHAIDYHFISTQGYPIEGWAVVYYITHLLKGALLFITISLIG  356
```

```
Query  270  TGWAFIKPFLGEKDKNIFLVVIPLQILANIAIIVLEETA---------PEVLRLVDIICC  320
            TGWAF+K  L +KDK IF++VIPLQ+LAN+A I++E T           E+L LVD++CC
Sbjct  357  TGWAFVKHILSDKDKKIFMIVIPLQVLANVAYIIIESTEEGTTEYGLWKEILFLVDLLCC  416

Query  321  GAILVPIIWSIKHLRDAAAIDGKAKRNMEKLKLFREFYLLVVTYIYFTRIIVFLLDATLH  380
            GAIL P++WSI+HL++A+A DGKA  N+ KLKLFR +Y+++V YIYFTRII  L+   +
Sbjct  417  GAILFPVVWSIRHLQEASATDGKAAVNLAKLKLFRHYYVMIVCYIYFTRIIAILIKIIVP  476

Query  381  YQYVWLGEFFTELATLIFWGLTGYKFRPVADNPYLKLDDEEEDAEREE  428
            +Q+ WL +F  ELATL+F+ LTGYKFRP +DNPYL+L   EED E +E
Sbjct  477  FQWKWLYQFLDELATLVFFCLTGYKFRPASDNPYLQLPQNEEDVEMDE  524
```

Range 2: 13 to 99

Score:43.1 bits(100), Expect:2.7,
Method:Compositional matrix adjust.,
Identities:30/89(34%), Positives:41/89(46%), Gaps:4/89(4%)

```
Query  15   IFCLVAFLAPTANGLIHKLSIKNDRRLAFRIETFGFFTGGVMEMAIENF--KVVDDKGSL  72
            +F L    L +  G IH+L +K+D R   R+ TFGFF  G M + + N   K VD K
Sbjct  13   MFLLWILLLTSVRGRIHRLILKDDVRQKIRLNTFGFFKNGNMTVKMTNLTSKGVDFKAV-  71

Query  73   WDDLSAGFIIKHIETDSGSFIEETDASKC  101
             D  + G  +   + D  S   E D   C
Sbjct  72   -DTSALGLSLDRTKNDGFSSYLEEDIDFC  99
```

>PREDICTED: protein GPR107 isoform X2 [Latimeria chalumnae]
Sequence ID: XP_014351767.1 Length: 513
Range 1: 172 to 481

Score:293 bits(749), Expect:5e-90,
Method:Compositional matrix adjust.,
Identities:147/312(47%), Positives:206/312(66%), Gaps:18/312(5%)

```
Query  133  TIDKTRPEGFYTVVFLNCQPGTYVS------FDLTLTNYNPGPNYLSAGLTALPTLYAML  186
             +      EG Y++ F NC       S        D+ +   NPG ++LSAG   LP LY  +
Sbjct  172  NVSDVAQEGLYSLYFHNCYSKEASSRQLRFDLDIKIEEKNPG-SFLSAGEIPLPKLYISM  230

Query  187  FVVWTVILGVWLFHFMRGQGKRIFRIHHLVTGIILLKLLTLLFEAIEFHYKKTTGHP-GG  245
               + +  GVW+ H +R +   +F+IH L+  +   K L+L+F AI+++Y  T G P  G
Sbjct  231  AFFFLLSGGVWV-HLLRKRRNDVFKIHWLMAALPFTKSLSLVFHAIDYYYISTQGFPIEG  289

Query  246  WVIAYYIFSGLKGTMMFVVIALIGTGWAFIKPFLGEKDKNIFLVVIPLQILANIAIIVLE  305
            W + YYI   LKG ++F+ IALIGTGWAF+K  L +KDK IF++VIPLQ+LAN+A I++E
Sbjct  290  WAVVYYITHLLKGALLFITIALIGTGWAFVKHILSDKDKKIFMIVIPLQVLANVAYIIIE  349

Query  306  ETA---------PEVLRLVDIICCGAILVPIIWSIKHLRDAAAIDGKAKRNMEKLKLFRE  356
             T           E+L LVD++CCGAIL P++WSI+HL++A+A DGKA  N+ KLKLFR
Sbjct  350  STEEGTTEYGLWKEILFLVDLLCCGAILFPVVWSIRHLQEASATDGKAAINLAKLKLFRH  409
```

```
Query  357  FYLLVVTYIYFTRIIVFLLDATLHYQYVWLGEFFTELATLIFWGLTGYKFRPVADNPYLK  416
            +Y+++V YIYFTRII  L+  T+ +Q+ WL +   ELATLIF+ LTGYKFRP +DNPYL+
Sbjct  410  YYVMIVCYIYFTRIIAILIKFTIPFQWKWLYQLLDELATLIFFVLTGYKFRPASDNPYLQ  469

Query  417  LDDEEEDAEREE  428
            L  +E+DAE +E
Sbjct  470  LPQDEDDAEMDE  481
```

>hypothetical protein NSK_002311 [Nannochloropsis salina CCMP1776]
Sequence ID: TFJ86657.1 Length: 445
Range 1: 1 to 391

Score:290 bits(742), Expect:7e-90,
Method:Compositional matrix adjust.,
Identities:167/395(42%), Positives:244/395(61%), Gaps:26/395(6%)

```
Query  56   MEMAIENFKVVDDKGSLWDDLSAGFIIKHIETDSGS------FIEETDASKCVSLLDEPE  109
            ME+   +F V   + +    + AGF++   E+++ +       +E   A +    LD
Sbjct  1    MELNFNHFVVKVPEKTKDATVRAGFLMHLTESETTARQDLEEALERLAADQEDCWLDHRG  60

Query  110  RGDTIAVKVTKPSKENEKQSKVITIDKTRPEGFYTVVFLNCQPG-TYVSFDLTLTNYNPG  168
              D + + ++ P    EK  KV        EG Y+++F+ CQP  + VSF +    YNPG
Sbjct  61   PNDEV-IDLSDPESWAEK--KVQHTVAPGEEGLYSLIFVRCQPAASAVSFKIHAKFYNPG  117

Query  169  PNYLSAGLTALPTLYAMLFVVWTVILGVWLFHFMRGQGKRIFRIHHLVTGIILLKLLTLL  228
            PNYLSAG   LPTLY + F+ + V +  W+  F R + + + RIH+++  +++ K L+LL
Sbjct  118  PNYLSAGEAPLPTLYFVFFLCYLVAMAAWVAIFRR-RKEHVHRIHNMMLALLVFKTLSLL  176

Query  229  FEAIEFH-YKKT--TGHPGGWVIAYYIFSGLKGTMMFVVIALIGTGWAFIKPFLGEKDKN  285
            FEA+ F  Y+K   TGH  GW + YY+F+ +KG M FVVI LIGTGW+ +KP+L +++K
Sbjct  177  FEAVRFQAYQKHGLTGHADGWAVVYYVFAFVKGIMRFVVILLIGTGWSLLKPYLSDREKK  236

Query  286  IFLVVIPLQILANIAIIVLEE---TAP---------EVLRLVDIICCGAILVPIIWSIKH  333
            + LVV+ LQ++ N A++VL+E   T+P         ++  L+DI+CC AIL PI+WSI+H
Sbjct  237  VVLVVLALQVIDNTAMVVLDELSQTSPGSASWLTWRDIFHLIDIVCCCAILFPIVWSIRH  296

Query  334  LRDAAAIDGKAKRNMEKLKLFREFYLLVVTYIYFTRIIVFLLDATLHYQYVWLGEFFTEL  393
            LR AAA DGK +  + KL LFR+FYL+VV YIYFTRI+VFL+ ATL Y  +WL  FF E
Sbjct  297  LRQAAAADGKMEHILRKLTLFRQFYLMVVAYIYFTRIVVFLVKATLPYDLLWLQAFFDEG  356

Query  394  ATLIFWGLTGYKFRPVADNPYLKLDDEEEDAEREE  428
            AT +F+ +TG+KF P   NPYL +D +E DA   E
Sbjct  357  ATFLFYTVTGWKFCPADANPYLAVDTDERDAAELE  391
```

>protein GPR107 [Echinops telfairi]
Sequence ID: XP_012862854.1 Length: 333
Range 1: 22 to 332

Score:285 bits(730), Expect:2e-89,

Method:Compositional matrix adjust.,
Identities:147/313(47%), Positives:205/313(65%), Gaps:19/313(6%)

```
Query  133  TIDKTRPEGFYTVVFLNC-----QPGTYVSF--DLTLTNYNPGPNYLSAGLTALPTLYAM  185
             I     EG Y++ F  C     +    +SF  D+ +T  NP  +YLSAG   LP LY
Sbjct  22   NISTDDQEGLYSLYFHKCLGNEVRSSDKISFSLDIEITEKNPD-SYLSAGEIPLPKLYIS  80

Query  186  LFVVWTVILGVWLFHFMRGQGKRIFRIHHLVTGIILLKLLTLLFEAIEFHYKKTTGHP-G  244
            + + + +   +W+ H +R +   +F+IH L+  +   K L+L+F AI++HY  + G P
Sbjct  81   MALFFFLSGTIWI-HILRKRRNDVFKIHWLMAALPFTKSLSLVFHAIDYHYISSQGFPIE  139

Query  245  GWVIAYYIFSGLKGTMMFVVIALIGTGWAFIKPFLGEKDKNIFLVVIPLQILANIAIIVL  304
            GW + YYI   LKG ++F+ IALIGTGWAFIK  L +KDK IF++VIPLQ+LAN+A I++
Sbjct  140  GWAVVYYITHLLKGALLFITIALIGTGWAFIKHILSDKDKKIFMIVIPLQVLANVAYIII  199

Query  305  EETA---------PEVLRLVDIICCGAILVPIIWSIKHLRDAAAIDGKAKRNMEKLKLFR  355
            E T           + L LVD++CCGAIL P++WSI+HL++A+A DGKA  N+ KLKLFR
Sbjct  200  ESTEEGTTEYSLWKDSLFLVDLLCCGAILFPVVWSIRHLQEASATDGKAAINLAKLKLFR  259

Query  356  EFYLLVVTYIYFTRIIVFLLDATLHYQYVWLGEFFTELATLIFWGLTGYKFRPVADNPYL  415
             +Y+L+V YIYFTRII FLL   + +Q+ WL +   E+ATL+F+ LTGYKFRP +DNPYL
Sbjct  260  HYYVLIVCYIYFTRIIAFLLKLAVPFQWKWLYQLLDEMATLVFFVLTGYKFRPASDNPYL  319

Query  416  KLDDEEEDAEREE  428
            +L  EE+D E E
Sbjct  320  QLSQEEDDLEMES  332
```

>Transmembrane receptor, eukaryota [Nannochloropsis gaditana]
Sequence ID: EWM24894.1 Length: 512
Range 1: 67 to 458

Score:291 bits(745), Expect:2e-89,
Method:Compositional matrix adjust.,
Identities:166/396(42%), Positives:244/396(61%), Gaps:26/396(6%)

```
Query  55   VMEMAIENFKVVDDKGSLWDDLSAGFIIKHIETDSGS------FIIEETDASKCVSLLDEP  108
            VME+   +F V   + +    + AGF++   E+++ +       +E   A +    LD
Sbjct  67   VMELNFNHFVVKVPEKTKDATVRAGFLMHLTESETTARQDLEEALERLAADQEDCWLDHR  126

Query  109  ERGDTIAVKVTKPSKENEKQSKVITIDKTRPEGFYTVVFLNCQPG-TYVSFDLTLTNYNP  167
               D + + ++ P    EKQ +         EG Y+++F+ CQP  + VSF +    YNP
Sbjct  127  GPNDEV-IDLSDPESWAEKQVQHTVAPGE--EGLYSLIFVRCQPAPSAVSFKIHAKFYNP  183

Query  168  GPNYLSAGLTALPTLYAMLFVVWTVILGVWLFHFMRGQGKRIFRIHHLVTGIILLKLLTL  227
            GPNYLSAG   LPTLY + F+ + V +  W+    R + + + RIH+++  +++ K L+L
Sbjct  184  GPNYLSAGEAPLPTLYFVFFLCYLVAMAAWV-AVCRRRKEHVHRIHNMMLALLVFKTLSL  242

Query  228  LFEAIEFH-YKKT--TGHPGGWVIAYYIFSGLKGTMMFVVIALIGTGWAFIKPFLGEKDK  284
            LFEA+ F  Y+K   TGH  GW + YY+F+ +KG M FVVI LIGTGW+ +KP+L +++K
Sbjct  243  LFEAVRFQAYQKHGLTGHADGWAVVYYVFAFVKGIMRFVVILLIGTGWSLLKPYLSDREK  302

Query  285  NIFLVVIPLQILANIAIIVLEE---TAP---------EVLRLVDIICCGAILVPIIWSIK  332
```

```
                + LVV+ LQ++ N A++VL+E   T+P         ++  L+DI+CC AIL PI+WSI+
Sbjct  303  KVVLVVLALQVIDNTAMVVLDELSQTSPGSASWLTWRDIFHLIDIVCCCAILFPIVWSIR  362

Query  333  HLRDAAAIDGKAKRNMEKLKLFREFYLLVVTYIYFTRIIVFLLDATLHYQYVWLGEFFTE  392
            HLR AAA DGK +  + KL LFR+FYL+VV YIYFTRI+VFL+ ATL Y  +WL  FF E
Sbjct  363  HLRQAAAADGKMEHILRKLTLFRQFYLMVVAYIYFTRIVVFLVKATLPYDLLWLQAFFDE  422

Query  393  LATLIFWGLTGYKFRPVADNPYLKLDDEEEDAEREE  428
             AT +F+ +TG+KF P   NPYL +D +E DA   E
Sbjct  423  GATFLFYTVTGWKFCPADANPYLAVDTDERDAAELE  458
```

>protein GPR107 isoform X1 [Amblyraja radiata]
Sequence ID: XP_032904977.1 Length: 554
Range 1: 214 to 523

Score:292 bits(748), Expect:2e-89,
Method:Compositional matrix adjust.,
Identities:148/312(47%), Positives:208/312(66%), Gaps:18/312(5%)

```
Query  133  TIDKTRPEGFYTVVFLNCQPGTYV------SFDLTLTNYNPGPNYLSAGLTALPTLYAML  186
            TI+    +G Y++ F NC     +        D+T+  YNPG +YLSAG   LP LY  +
Sbjct  214  TINTEEEQGLYSLYFHNCYSDKAILQSFKFDMDITIVEYNPG-SYLSAGEIPLPKLYISM  272

Query  187  FVVWTVILGVWLFHFMRGQGKRIFRIHHLVTGIILLKLLTLLFEAIEFHYKKTTGHP-GG  245
               + V   VW++  +R +   +F+IH L+  +   K L+L+F AI++H+  T G+P  G
Sbjct  273  AFFFFVAGVVWVY-ILRKRSNDVFKIHWLMGALAFTKSLSLVFHAIDYHFISTQGYPIEG  331

Query  246  WVIAYYIFSGLKGTMMFVVIALIGTGWAFIKPFLGEKDKNIFLVVIPLQILANIAIIVLE  305
            W + YYI   LKG ++F+ IALIGTGWAF+K  L +KDK IF++VIPLQ+LAN+A I++E
Sbjct  332  WAVVYYITHLLKGALLFITIALIGTGWAFVKHILSDKDKKIFMIVIPLQVLANVAYIIIE  391

Query  306  ETA---------PEVLRLVDIICCGAILVPIIWSIKHLRDAAAIDGKAKRNMEKLKLFRE  356
             T           E+L LVD++CCGAIL P++WSI+HL++A+A DGKA  N+ KLKLFR
Sbjct  392  STEEGTTEYGLWKEILFLVDLLCCGAILFPVVWSIRHLQEASATDGKAAINLAKLKLFRH  451

Query  357  FYLLVVTYIYFTRIIVFLLDATLHYQYVWLGEFFTELATLIFWGLTGYKFRPVADNPYLK  416
            +Y+++V YIYFTRII  L+   + +Q+ WL +F  ELATL+F+ LTGYKFRP +DNPYL+
Sbjct  452  YYVMIVCYIYFTRIIAILIKIIVPFQWKWLYQFLDELATLVFFCLTGYKFRPASDNPYLQ  511

Query  417  LDDEEEDAEREE  428
            L   EED E +E
Sbjct  512  LSQNEEDVEMDE  523
```

>protein GPR108-like [Oncorhynchus mykiss]
Sequence ID: XP_021412141.1 Length: 338
Range 1: 8 to 314

Score:285 bits(729), Expect:2e-89,
Method:Compositional matrix adjust.,
Identities:147/309(48%), Positives:207/309(66%), Gaps:17/309(5%)

```
Query  140   EGFYTVVFLNCQ---PGTYVSFDLTL--TNYNPGPNYLSAGLTALPTLYAMLFVVWTVIL  194
             EG Y + F  C+   PG    + L +  T  NPG  YLSA    LP LY +   V+ +I
Sbjct  8     EGLYNLDFHYCENILPGNNRPYTLNVEVTEKNPG-GYLSAAEIPLPRLYIFMAGVFFIIA  66

Query  195   GVWLFHFMRGQGKRIFRIHHLVTGIILLKLLTLLFEAIEFHYKKTTGHP-GGWVIAYYIF  253
              VW++  ++ +   +F+IH L+  +   K ++L+F +I +H+  T GHP  GW + YYI
Sbjct  67    MVWVYTLLKHR-YSVFKIHWLMAALAFTKSISLVFHSINYHFINTEGHPIEGWAVMYYIT  125

Query  254   SGLKGTMMFVVIALIGTGWAFIKPFLGEKDKNIFLVVIPLQILANIAIIVLEETA-----  308
               LKG ++F+ +ALIGTGWAF+K  L +K+K IF++VIPLQ+LAN+A I++E T
Sbjct  126   HLLKGALLFITLALIGTGWAFVKYILSDKEKKIFMIVIPLQVLANVAYIIIESTEEGSSE  185

Query  309   ----PEVLRLVDIICCGAILVPIIWSIKHLRDAAAIDGKAKRNMEKLKLFREFYLLVVTY  364
                  EVL LVD+ICCGAIL P+IWSI+HL++A++ DGKA  N+EKLKLFR +Y+++V Y
Sbjct  186   YALWKEVLFLVDLICCGAILFPVIWSIRHLQEASSTDGKAAMNLEKLKLFRHYYVMIVCY  245

Query  365   IYFTRIIVFLLDATLHYQYVWLGEFFTELATLIFWGLTGYKFRPVADNPYLKLDDEEEDA  424
             IYFTRII  LL  T+ +Q+ W  EF  E++TLIF+ LTGYKFRP ++NPYL+L  +EED
Sbjct  246   IYFTRIIAILLKVTVPFQWQWCYEFLVEVSTLIFFVLTGYKFRPASNNPYLQLPLDEEDV  305

Query  425   EREEAQRQS  433
             E +E   +S
Sbjct  306   EMDEVVTES  314
```

>hypothetical protein CBR_g49976 [Chara braunii]
Sequence ID: GBG65182.1 Length: 373
Range 1: 67 to 367

Score:286 bits(732), Expect:3e-89,
Method:Compositional matrix adjust.,
Identities:148/302(49%), Positives:201/302(66%), Gaps:14/302(4%)

```
Query  143   YTVVFLNC-QPGTYVSFDLTLTNYNPG---PNYLSAGLTALPTLYAMLFVVWTVILGVWL  198
             YT+VF+NC  P   VS ++ +  +N     P++L AG T LP LY   F ++ +    W+
Sbjct  67    YTLVFVNCFSPRQAVSMNVRIELFNKEGTVPDFLPAGKTQLPKLYFSFFALFVLAGVAWV  126

Query  199   FHFMRGQGKRIFRIHHLVTGIILLKLLTLLFEAIEFHYKKTTGHPGGWVIAYYIFSGLKG  258
             +   R + +   R+H L+  ++ LK +TLL  A E+   K TG P GW +AYYIFS L+G
Sbjct  127   YVCYRHK-ETTHRVHILMGVLVSLKAITLLALAAEYTLVKQTGTPHGWNVAYYIFSFLRG  185

Query  259   TMMFVVIALIGTGWAFIKPFLGEKDKNIFLVVIPLQILANIAIIVLEETAP---------  309
              M+F VI LIGTGW+F+KPFL ++DK + L+VIPLQ+ ANIA  ++EE  P
Sbjct  186   VMLFTVILLIGTGWSFLKPFLQDRDKKVILIVIPLQVFANIAATIVEEYTPAAKGWFTWR  245

Query  310   EVLRLVDIICCGAILVPIIWSIKHLRDAAAIDGKAKRNMEKLKLFREFYLLVVTYIYFTR  369
             ++  L+DIICC  IL+PI+WSIKHLR+A+  DGKA RNM KL LFR+FY++VV+YIYFTR
Sbjct  246   DIFHLIDIICCCTILLPIVWSIKHLREASRTDGKAARNMIKLSLFRQFYVMVVSYIYFTR  305

Query  370   IIVFLLDATLHYQYVWLGEFFTELATLIFWGLTGYKFRPVADNPYLKLDDEEEDAEREEA  429
             I+V+LL  T+ Y Y W+ +     ATLIF+ LTGY FRPV  NPY  LDDEEE+A  + A
Sbjct  306   IVVYLLRNTVSYHYTWVSDLADNAATLIFYILTGYIFRPVERNPYFVLDDEEEEAAVQMA  365
```

```
Query  430  QR  431
             +
Sbjct  366  LK  367
```

>protein GPR107 isoform X3 [Balaenoptera acutorostrata scammoni]
Sequence ID: XP_028017752.1 Length: 396
Range 1: 15 to 332

Score:287 bits(734), Expect:3e-89,
Method:Compositional matrix adjust.,
Identities:149/321(46%), Positives:206/321(64%), Gaps:21/321(6%)

```
Query  128  QSKVITIDKTRPEGFYTVVFLNCQPGTYV--------SFDLTLTNYNPGPNYLSAGLTAL  179
            Q     I     EG Y++ F  C PG+ +        S D+ +T  NP  +YLSAG   L
Sbjct  15   QKFFFNISTEDQEGLYSLYFHKC-PGSEIRSNDKFSFSLDIDITEKNPD-SYLSAGEIPL  72

Query  180  PTLYAMLFVVWTVILGVWLFHFMRGQGKRIFRIHHLVTGIILLKLLTLLFEAIEFHYKKT  239
            P LY  +   + +   +W+ H +R +   +F+IH L+  +   K L+L+F AI++HY  +
Sbjct  73   PKLYISMAFFFFLSGTIWI-HILRKRRNDVFKIHWLMAALPFTKSLSLVFHAIDYHYISS  131

Query  240  TGHP-GGWVIAYYIFSGLKGTMMFVVIALIGTGWAFIKPFLGEKDKNIFLVVIPLQILAN  298
             G P  GW + YYI   LKG ++F+ IALIGTGWAFIK  L +KDK IF++VIPLQ+LAN
Sbjct  132  QGFPIEGWAVVYYITHLLKGALLFITIALIGTGWAFIKHILSDKDKKIFMIVIPLQVLAN  191

Query  299  IAIIVLEETA---------PEVLRLVDIICCGAILVPIIWSIKHLRDAAAIDGKAKRNME  349
            +A I++E T           + L LVD++CCGAIL P++WSI+HL++A+A DGKA  N+
Sbjct  192  VAYIIIESTEEGTTEYGLWKDSLFLVDLLCCGAILFPVVWSIRHLQEASATDGKAAINLA  251

Query  350  KLKLFREFYLLVVTYIYFTRIIVFLLDATLHYQYVWLGEFFTELATLIFWGLTGYKFRPV  409
            KLKLFR +Y+L+V YIYFTRII FLL   + +Q+ WL +   E+ATL+F+ LTGYKFRP
Sbjct  252  KLKLFRHYYVLIVCYIYFTRIIAFLLKLAVPFQWKWLYQLLDEMATLVFFVLTGYKFRPA  311

Query  410  ADNPYLKLDDEEEDAEREEAQ  430
            +DNPYL+L  EE+D E E
Sbjct  312  SDNPYLQLSQEEDDLEMESVS  332
```

>predicted protein [Micromonas pusilla CCMP1545]
Sequence ID: XP_003056720.1 Length: 451
>predicted protein [Micromonas pusilla CCMP1545]
Sequence ID: EEH58365.1 Length: 451
Range 1: 19 to 442

Score:288 bits(738), Expect:4e-89,
Method:Compositional matrix adjust.,
Identities:171/428(40%), Positives:248/428(57%), Gaps:25/428(5%)

```
Query  22   LAPTANGLIHKLSIKNDRRLAFRIE-TFGFFTGGVMEMAIEN-FKVVDDKGS-LWDDLSA  78
            +A   + LI    ++ D R    I   FGF   G +++ I+N   +   +G+   D
Sbjct  19   IATPTHALITNEGVEKDGRPKIPIALDFGFDPQGGIDIKIKNPVHIYTTEGAEPVDRTQM  78
```

```
Query  79   GFIIKHIETDSGSFIEETDASKCVSLLDEPERGDTIAVKVTKPSKENEKQSKVI--TIDK  136
            GF+I   + D+     E + +K   +LD         ++        E Q  V   TI K
Sbjct  79   GFLISFAKADAEL---EDELAKGTCILDSQYVKKLFTMQDIADQTTEEGQDFVYSNTIAK  135

Query  137  TRPEGFYTVVFLNCQPGTYVSFDLTLTNYNP----GPNYLSAGLTALPTLYAMLFVVWTV  192
                G Y++ F+NC   + VSFD+T+  YN       +YLSAG   LPTLY + F  +
Sbjct  136  DMA-GQYSLYFVNCVEHSAVSFDITVELYNHMEGGKKDYLSAGEKPLPTLYFLCFCAFVA  194

Query  193  ILGVWLF--HFMRG-QGKRIFRIHHLVTGIILLKLLTLLFEAIEFHYKKTTGHPGGWVIA  249
            +   W +  H  R  Q   + +IH L+  ++++K LT+L +A+ FH ++  G   GW IA
Sbjct  195  MGCAWAWNLHTSRNTQPGSVQKIHLLMLLLVMMKALTVLCQALMFHSRRVKGDADGWNIA  254

Query  250  YYIFSGLKGTMMFVVIALIGTGWAFIKPFLGEKDKNIFLVVIPLQILANIAIIVLEETA-  308
            YY F+ ++G M+F V+ LIGTGW+F+KPFL E++KN+ ++VIP+Q+ ANIA IVL++
Sbjct  255  YYFFTSVRGLMLFTVVVLIGTGWSFLKPFLNEREKNVLMIVIPMQVFANIATIVLDDNGV  314

Query  309  --------PEVLRLVDIICCGAILVPIIWSIKHLRDAAAIDGKAKRNMEKLKLFREFYLL  360
                     ++  L+DI CC AIL PI+WSIKHLR+AA  DGK  RNM KL LFR+FY++
Sbjct  315  ALAGWLEWRDLFHLLDIACCCAILFPIVWSIKHLREAALTDGKKVRNMNKLILFRQFYVM  374

Query  361  VVTYIYFTRIIVFLLDATLHYQYVWLGEFFTELATLIFWGLTGYKFRPVADNPYLKLDDE  420
            VV YIYFTRIIV+LL  T+ Y   WL +  +E AT+ F+ +TGY FRP+ DNPYL L+++
Sbjct  375  VVAYIYFTRIIVYLLKQTMAYNMTWLADLASEAATMCFYCVTGYMFRPIPDNPYLHLNED  434

Query  421  EEDAEREE  428
            E D E  E
Sbjct  435  ELDEESIE  442
```

>PREDICTED: protein GPR107 isoform X1 [Latimeria chalumnae]
Sequence ID: XP_006008808.1 Length: 565
Range 1: 224 to 533

Score:291 bits(746), Expect:5e-89,
Method:Compositional matrix adjust.,
Identities:147/312(47%), Positives:206/312(66%), Gaps:18/312(5%)

```
Query  133  TIDKTRPEGFYTVVFLNCQPGTYVS------FDLTLTNYNPGPNYLSAGLTALPTLYAML  186
             +     EG Y++ F NC      S       D+ +   NPG ++LSAG   LP LY  +
Sbjct  224  NVSDVAQEGLYSLYFHNCYSKEASSRQLRFDLDIKIEEKNPG-SFLSAGEIPLPKLYISM  282

Query  187  FVVWTVILGVWLFHFMRGQGKRIFRIHHLVTGIILLKLLTLLFEAIEFHYKKTTGHP-GG  245
               + +  GVW+ H +R +   +F+IH L+  +   K L+L+F AI+++Y  T G P  G
Sbjct  283  AFFFLLSGGVWV-HLLRKRRNDVFKIHWLMAALPFTKSLSLVFHAIDYYYISTQGFPIEG  341

Query  246  WVIAYYIFSGLKGTMMFVVIALIGTGWAFIKPFLGEKDKNIFLVVIPLQILANIAIIVLE  305
            W + YYI   LKG ++F+ IALIGTGWAF+K  L +KDK IF++VIPLQ+LAN+A I++E
Sbjct  342  WAVVYYITHLLKGALLFITIALIGTGWAFVKHILSDKDKKIFMIVIPLQVLANVAYIIIE  401

Query  306  ETA---------PEVLRLVDIICCGAILVPIIWSIKHLRDAAAIDGKAKRNMEKLKLFRE  356
             T           E+L LVD++CCGAIL P++WSI+HL++A+A DGKA  N+ KLKLFR
Sbjct  402  STEEGTTEYGLWKEILFLVDLLCCGAILFPVVWSIRHLQEASATDGKAAINLAKLKLFRH  461
```

```
Query  357  FYLLVVTYIYFTRIIVFLLDATLHYQYVWLGEFFTELATLIFWGLTGYKFRPVADNPYLK  416
            +Y+++V YIYFTRII  L+  T+ +Q+ WL +   ELATLIF+ LTGYKFRP +DNPYL+
Sbjct  462  YYVMIVCYIYFTRIIAILIKFTIPFQWKWLYQLLDELATLIFFVLTGYKFRPASDNPYLQ  521

Query  417  LDDEEEDAEREE  428
            L  +E+DAE +E
Sbjct  522  LPQDEDDAEMDE  533
```

>protein GPR107 isoform X4 [Pongo abelii]
Sequence ID: XP_009243232.1 Length: 332
Range 1: 22 to 331

Score:284 bits(727), Expect:5e-89,
Method:Compositional matrix adjust.,
Identities:149/312(48%), Positives:202/312(64%), Gaps:18/312(5%)

```
Query  133  TIDKTRPEGFYTVVFLNCQ----PGTYVSF--DLTLTNYNPGPNYLSAGLTALPTLYAML  186
             I     EG Y++ F  C     P    SF  D+ +T  NP  +YLSAG   LP LY  +
Sbjct  22   NISTDDQEGLYSLYFHKCLGKELPSDKFSFSLDIEITEKNPD-SYLSAGEIPLPKLYISM  80

Query  187  FVVWTVILGVWLFHFMRGQGKRIFRIHHLVTGIILLKLLTLLFEAIEFHYKKTTGHP-GG  245
               + +   +W+ H +R +   +F+IH L+  +   K L+L+F AI++HY  + G P  G
Sbjct  81   AFFFFLSGTIWI-HILRKRRNDVFKIHWLMAALPFTKSLSLVFHAIDYHYISSQGFPIEG  139

Query  246  WVIAYYIFSGLKGTMMFVVIALIGTGWAFIKPFLGEKDKNIFLVVIPLQILANIAIIVLE  305
            W + YYI   LKG ++F+ IALIGTGWAFIK  L +KDK IF++VIPLQ+LAN+A I++E
Sbjct  140  WAVVYYITHLLKGALLFITIALIGTGWAFIKHILSDKDKKIFMIVIPLQVLANVAYIIIE  199

Query  306  ETA---------PEVLRLVDIICCGAILVPIIWSIKHLRDAAAIDGKAKRNMEKLKLFRE  356
             T           + L LVD++CCGAIL P++WSI+HL++A+A DGKA  N+ KLKLFR
Sbjct  200  STEEGTTEYGLWKDSLFLVDLLCCGAILFPVVWSIRHLQEASATDGKAAINLAKLKLFRH  259

Query  357  FYLLVVTYIYFTRIIVFLLDATLHYQYVWLGEFFTELATLIFWGLTGYKFRPVADNPYLK  416
            +Y+L+V YIYFTRII FLL   + +Q+ WL +   E ATL+F+ LTGYKFRP +DNPYL+
Sbjct  260  YYVLIVCYIYFTRIIAFLLKLAVPFQWKWLYQLLDETATLVFFVLTGYKFRPASDNPYLQ  319

Query  417  LDDEEEDAEREE  428
            L  EEED E E
Sbjct  320  LSQEEEDLEMES  331
```

>protein GPR107 isoform X2 [Nomascus leucogenys]
Sequence ID: XP_030673890.1 Length: 363
Range 1: 22 to 336

Score:285 bits(728), Expect:8e-89,
Method:Compositional matrix adjust.,
Identities:150/317(47%), Positives:203/317(64%), Gaps:18/317(5%)

```
Query  133  TIDKTRPEGFYTVVFLNCQ----PGTYVSF--DLTLTNYNPGPNYLSAGLTALPTLYAML  186
```

```
            I     EG Y++ F  C     P    SF  D+ +T  NP  +YLSAG   LP LY  +
Sbjct  22   NISTDDQEGLYSLYFHKCLGKELPSDKFSFSLDIEITEKNPD-SYLSAGEIPLPKLYISM  80

Query  187  FVVWTVILGVWLFHFMRGQGKRIFRIHHLVTGIILLKLLTLLFEAIEFHYKKTTGHP-GG  245
               + +   +W+ H +R +   +F+IH L+  +   K L+L+F AI++HY  + G P  G
Sbjct  81   AFFFFLSGTIWI-HILRKRRNDVFKIHWLMAALPFTKSLSLVFHAIDYHYISSQGFPIEG  139

Query  246  WVIAYYIFSGLKGTMMFVVIALIGTGWAFIKPFLGEKDKNIFLVVIPLQILANIAIIVLE  305
            W + YYI   LKG ++F+ IALIGTGWAFIK  L +KDK IF++VIPLQ+LAN+A I++E
Sbjct  140  WAVVYYITHLLKGALLFITIALIGTGWAFIKHILSDKDKKIFMIVIPLQVLANVAYIIIE  199

Query  306  ETA---------PEVLRLVDIICCGAILVPIIWSIKHLRDAAAIDGKAKRNMEKLKLFRE  356
             T           + L LVD++CCGAIL P++WSI+HL++A+A DGKA  N+ KLKLFR
Sbjct  200  STEEGTTEYGLWKDSLFLVDLLCCGAILFPVVWSIRHLQEASATDGKAAINLAKLKLFRH  259

Query  357  FYLLVVTYIYFTRIIVFLLDATLHYQYVWLGEFFTELATLIFWGLTGYKFRPVADNPYLK  416
            +Y+L+V YIYFTRII FLL   + +Q+ WL +   E ATL+F+ LTGYKFRP +DNPYL+
Sbjct  260  YYVLIVCYIYFTRIIAFLLKLAVPFQWKWLYQLLDETATLVFFVLTGYKFRPASDNPYLQ  319

Query  417  LDDEEEDAEREEAQRQS  433
            L  EEED E E     S
Sbjct  320  LSQEEEDLEMESVVTTS  336
```

>protein GPR107-like [Durio zibethinus]
Sequence ID: XP_022735937.1 Length: 436
Range 1: 6 to 418

Score:287 bits(734), Expect:9e-89,
Method:Compositional matrix adjust.,
Identities:166/429(39%), Positives:246/429(57%), Gaps:37/429(8%)

```
Query  12   CFLIFCLVAFLAPTANGLIHKLSIKNDRRLAFRIETFGFFTGGVMEMAIENFKVVDDKGS  71
            CF +F L++         I    I++D R     + FGF   G +E+ +    + + K
Sbjct  6    CFFVFLLISSFVSLGFAEIRFTEIRSDDRSIIPFDEFGFTHTGRLELNVSQIALSNPKSD  65

Query  72   LWDDLSAGFIIKHIETDSGSFIEETDASKCVSLLDE-PERGDTIAVK------VTKPSKE  124
            L  DL          T  G F+   D+  C+ +L +  +R  T A+K      V+
Sbjct  66   L--DL----------TKVGFFLCTRDS--CMHVLQQLQDRETTCALKSDVVKLVSSFKSL  111

Query  125  NEKQSKVITIDKTRPEGFYTVVFLNCQPGTYVSFDLTLTNYN-PGPN----YLSAGLTAL  179
            N K S  I  ++   +  YT+VF NC     VS ++    YN  G N    YLSAG T L
Sbjct  112  NGKSSLNIVYEENDADQ-YTLVFANCLSQVKVSMNVRSAMYNLDGKNNRQDYLSAGKTIL  170

Query  180  PTLYAMLFVVWTVILGVWLFHFMRGQGKRIFRIHHLVTGIILLKLLTLLFEAIEFHYKKT  239
            P +Y +L +V+  + G+W++   + +   +FRIH  +  +++LK   L+ EA +  Y K
Sbjct  171  PRVYFLLSLVYFTLAGIWIYVLYKKR-LTVFRIHFFMLAVVILKAFNLVCEAEDKSYIKR  229

Query  240  TGHPGGWVIAYYIFSGLKGTMMFVVIALIGTGWAFIKPFLGEKDKNIFLVVIPLQILANI  299
            TG   GW + +YIFS LKG M+F +I LIGTGW+F+KP+L +K+K + ++VIPLQ +ANI
Sbjct  230  TGSAHGWDVLFYIFSFLKGIMLFTLIVLIGTGWSFLKPYLQDKEKKVLMIVIPLQFVANI  289

Query  300  AIIVLEETAP---------EVLRLVDIICCGAILVPIIWSIKHLRDAAAIDGKAKRNMEK  350
```

```
            A +V++ET P         +V  LVD++CC A+L PI+WSIK+LR+AA  DGKA  N+ K
Sbjct  290  AQVVIDETGPFGQDWITWRQVFLLVDVVCCCAVLFPIVWSIKNLREAARTDGKAAVNLMK  349

Query  351  LKLFREFYLLVVTYIYFTRIIVFLLDATLHYQYVWLGEFFTELATLIFWGLTGYKFRPVA  410
            L LFR++Y++V+ YIYFTR++V+ L+    Y+Y+W      ELATL F+  TGYKF+P A
Sbjct  350  LTLFRQYYIVVICYIYFTRVVVYALETITSYKYLWTSVVAGELATLAFYVFTGYKFKPEA  409

Query  411  DNPYLKLDD  419
             NPY  +DD
Sbjct  410  HNPYFVIDD  418
```

>PREDICTED: protein GPR107 isoform X1 [Saimiri boliviensis boliviensis]
Sequence ID: XP_010349023.1 Length: 332
Range 1: 22 to 331

Score:283 bits(725), Expect:9e-89,
Method:Compositional matrix adjust.,
Identities:148/312(47%), Positives:202/312(64%), Gaps:18/312(5%)

```
Query  133  TIDKTRPEGFYTVVFLNCQ----PGTYVSF--DLTLTNYNPGPNYLSAGLTALPTLYAML  186
             I     EG Y++ F  C     P    SF  D+ +T  NP  +YLSAG   LP LY  +
Sbjct  22   NISTDDQEGLYSLYFHKCLRNELPSDKFSFSLDIDITEKNPD-SYLSAGEIPLPKLYISM  80

Query  187  FVVWTVILGVWLFHFMRGQGKRIFRIHHLVTGIILLKLLTLLFEAIEFHYKKTTGHP-GG  245
               + +   +W+ H +R +   +F+IH L+  +   K L+L+F AI++HY  + G P  G
Sbjct  81   AFFFFLSGTIWI-HILRKRRNDVFKIHWLMAALPFTKSLSLVFHAIDYHYISSQGFPIEG  139

Query  246  WVIAYYIFSGLKGTMMFVVIALIGTGWAFIKPFLGEKDKNIFLVVIPLQILANIAIIVLE  305
            W + YYI   LKG ++F+ IALIGTGWAFIK  L +KDK IF++VIPLQ+LAN+A I++E
Sbjct  140  WAVVYYITHLLKGALLFITIALIGTGWAFIKHILSDKDKKIFMIVIPLQVLANVAYIIIE  199

Query  306  ETA---------PEVLRLVDIICCGAILVPIIWSIKHLRDAAAIDGKAKRNMEKLKLFRE  356
             T            + L LVD++CCGAIL P++WSI+HL++A+A DGKA  N+ KLKLFR
Sbjct  200  STEEGTTEYGLWKDSLFLVDLLCCGAILFPVVWSIRHLQEASATDGKAAINLAKLKLFRH  259

Query  357  FYLLVVTYIYFTRIIVFLLDATLHYQYVWLGEFFTELATLIFWGLTGYKFRPVADNPYLK  416
            +Y+L+V YIYFTRII FLL   + +Q+ WL +   E ATL+F+ LTGYKFRP +DNPYL+
Sbjct  260  YYVLIVCYIYFTRIIAFLLKLAVPFQWKWLYQLLDETATLVFFVLTGYKFRPASDNPYLQ  319

Query  417  LDDEEEDAEREE  428
            L  EE+D E E
Sbjct  320  LSQEEDDLEMES  331
```

>unknown [Picea sitchensis]
Sequence ID: ABR17223.1 Length: 445
Range 1: 20 to 444

Score:287 bits(735), Expect:1e-88,
Method:Compositional matrix adjust.,
Identities:174/440(40%), Positives:247/440(56%), Gaps:36/440(8%)

```
Query  18   LVAFLAPTANGLIHKLSIKNDRRLAFRIETFGFFTGGVMEMAIENFKVVDDKGSLWDDLS  77
            L+ FL P  +  I    I++D R     + FGF   G +E+ ++       KG   + +
Sbjct  20   LMGFL-PFCSAEIRHSEIRSDDRSIIPFDEFGFTHEGRLEIYVKEASYKHLKGETINPVH  78

Query  78   AGFIIKHIETDSGSFIEETDASKCVSLLDEPERGDTIAVKVTKPSKE---NEKQSKVITI  134
             GF           F+   DA   V  L + E G+   V  +K        +    +I+
Sbjct  79   MGF-----------FLSTRDAWAHV--LQDLEHGEIHCVLESKLIVHLFTFKDLENLISY  125

Query  135  DKTRPE---GFYTVVFLNCQPGTYVSFDLTLTNYN------PGPNYLSAGLTALPTLYAM  185
            +KT  +     YT+VF NC P   VS D+    YN         +YLSAG T LP LY
Sbjct  126  NKTFKDFEANQYTLVFANCIPDVVVSMDVKTVIYNLEGAGDGTKDYLSAGETLLPKLYFS  185

Query  186  LFVVWTVILGVWLFHFMRGQGKRIFRIHHLVTGIILLKLLTLLFEAIEFHYKKTTGHPGG  245
            L +++ V+ GVW+F  +R +    +RIH  +  +I LK L LL EA +  + K TG   G
Sbjct  186  LSLIYVVLAGVWIFVVVRNR-LTAYRIHLFMAVLICLKALNLLCEAEDKSFIKRTGTAHG  244

Query  246  WVIAYYIFSGLKGTMMFVVIALIGTGWAFIKPFLGEKDKNIFLVVIPLQILANIAIIVLE  305
            W + +YIFS LKG M+F +I LIGTGW+F+KP+L  K+K + +VVIPLQ+LANIA +V++
Sbjct  245  WDVLFYIFSFLKGIMLFTLIVLIGTGWSFLKPYLQGKEKKVLIVVIPLQVLANIATVVID  304

Query  306  ETAP---------EVLRLVDIICCGAILVPIIWSIKHLRDAAAIDGKAKRNMEKLKLFRE  356
            ET P         ++  LVD+ICC A+L PI+WSIK+LR AA  DGKA  N+ KL LFR+
Sbjct  305  ETGPYAKDWLAWKQMFLLVDVICCCAVLFPIVWSIKNLRQAAHTDGKAAVNLMKLTLFRQ  364

Query  357  FYLLVVTYIYFTRIIVFLLDATLHYQYVWLGEFFTELATLIFWGLTGYKFRPVADNPYLK  416
            +Y++VV YIYFTR++V+ L     Y+Y W      ELATL F+  TGY+FRP+  NPY
Sbjct  365  YYIVVVCYIYFTRVVVYALITITAYKYAWTSVMAGELATLAFYVFTGYRFRPIGHNPYFV  424

Query  417  LDDEEEDAEREEAQRQSRTE  436
            +DD+EE+A  +  + +   E
Sbjct  425  IDDDEEEAASQSLKLEDDFE  444
```

>protein GPR108-like isoform X1 [Etheostoma cragini]
Sequence ID: XP_034715705.1 Length: 549
Range 1: 219 to 525

Score:290 bits(743), Expect:1e-88,
Method:Compositional matrix adjust.,
Identities:147/309(48%), Positives:207/309(66%), Gaps:17/309(5%)

```
Query  140  EGFYTVVFLNCQ---PGTYV--SFDLTLTNYNPGPNYLSAGLTALPTLYAMLFVVWTVIL  194
            EG YT+ F  C+   PG  +  SF + +T  NPG  YLSA    L  LY  +  V+ +
Sbjct  219  EGLYTLKFYYCRNRIPGNKLPYSFSVEVTEENPG-GYLSAAEIPLSRLYICMAGVFFIAA  277

Query  195  GVWLFHFMRGQGKRIFRIHHLVTGIILLKLLTLLFEAIEFHYKKTTGHP-GGWVIAYYIF  253
             VW++  M+ +   +F+IH L+  +   K  +L+F +I +H+  T GHP  GW + YYI
Sbjct  278  MVWVYTLMKHR-YSVFKIHWLMAALAFTKATSLVFHSINYHFINTKGHPIEGWAVMYYIT  336

Query  254  SGLKGTMMFVVIALIGTGWAFIKPFLGEKDKNIFLVVIPLQILANIAIIVLEETA-----  308
              LKG ++F+ +ALIGTGWAF+K  L +K+K IF++VIPLQ+LAN+A I++E T
Sbjct  337  HLLKGALLFITLALIGTGWAFVKYILSDKEKKIFMIVIPLQVLANVAFIIIESTEEGSSE  396
```

```
Query  309  ----PEVLRLVDIICCGAILVPIIWSIKHLRDAAAIDGKAKRNMEKLKLFREFYLLVVTY  364
                 E+L LVD+ICCGAIL P++WSI+HL++A++ DGKA  N+EKLKLFR +Y+++V Y
Sbjct  397  YYLWKEILFLVDLICCGAILFPVVWSIRHLQEASSTDGKAAMNLEKLKLFRHYYVMIVCY  456

Query  365  IYFTRIIVFLLDATLHYQYVWLGEFFTELATLIFWGLTGYKFRPVADNPYLKLDDEEEDA  424
            IYFTRII  LL  T+ +Q+ W  EF  E++TLIF+ LTGYKFRP +DNPYL+L  +EED
Sbjct  457  IYFTRIIAILLKVTMPFQWQWCYEFLVEVSTLIFFVLTGYKFRPASDNPYLQLPLDEEDV  516

Query  425  EREEAQRQS  433
            E +E   +S
Sbjct  517  EMDEVMTES  525
```

>hypothetical protein GPECTOR_8g123 [Gonium pectorale]
Sequence ID: KXZ52730.1 Length: 425
Range 1: 22 to 424

Score:286 bits(733), Expect:1e-88,
Method:Compositional matrix adjust.,
Identities:170/417(41%), Positives:243/417(58%), Gaps:40/417(9%)

```
Query  31   HKLSIKNDRRLAFRIETFGFFTGGVMEMAIENFKVV-----DDKGSLWDDLSAGFIIKHI  85
            H +  K+DR L    + FGF  GG +++ I +  +      D++ S W++   GF +  +
Sbjct  22   HSIVDKDDRPLIPLTDAFGFAEGGKLDITIRDIGLYRLHGSDEEVSHWENF--GFFLSPV  79

Query  86   ETDSGSFIEETDASKCVSLLDEPERGDTIAVKVTKPSKENEKQSKVITIDK--------T  137
            E D     +  D+SKC+  L++     T          ++    KVIT D+
Sbjct  80   EADVALEQDLADSSKCI--LNDVNNLFTF---------KDSAVQKVITEDQDAFTFHFIV  128

Query  138  RPEGFYTVVFLNCQPGTYVSFDLTLTNYNPGP----NYLSAGLTALPTLYAMLFVVWTVI  193
               G + + F NC+P T VSFD  +  YN       +Y+S G T+L  +Y  +F ++TV
Sbjct  129  ENGGLFYLYFANCEPDTPVSFDSRIEMYNLDKYGRKDYMSVGDTSLDAVYWTMFALFTVC  188

Query  194  LGVWLFHFMRGQGKRIFRIHHLVTGIILLKLLTLLFEAIEFHYKKTTGHPGGWVIAYYIF  253
               W     R + +   +IH+L+  +   K LTLL +A+  +Y + TG   GW IAYY+F
Sbjct  189  TAAWALWMYRNK-QHSHKIHYLMFALGFFKALTLLSQALMVYYIERTGSADGWNIAYYVF  247

Query  254  SGLKGTMMFVVIALIGTGWAFIKPFLGEKDKNIFLVVIPLQILANIAIIVLEETAP----  309
            + L+G + F V+ LIGTGW+++KPFLGEK+  I ++V+PLQ+ ANIAII+ EE +P
Sbjct  248  TFLRGILFFTVVVLIGTGWSYMKPFLGEKEARIIMIVVPLQVFANIAIIITEEESPSVKD  307

Query  310  -----EVLRLVDIICCGAILVPIIWSIKHLRDAAAIDGKAKRNMEKLKLFREFYLLVVTY  364
                 +V  LVDIICC AIL PI+WSIKHLR+A+  DGKA RN+EKL LFR+FY++VV Y
Sbjct  308  WFTWRDVFHLVDIICCCAILFPIVWSIKHLREASQTDGKAARNLEKLTLFRQFYVMVVVY  367

Query  365  IYFTRIIVFLLDATLHYQYVWLGEFFTELATLIFWGLTGYKFRPVADNPYLKLDDEE  421
            IY TRI+V+LL +T+ YQY W+     EL TL F+  T  KFRP   NPYLKL + E
Sbjct  368  IYVTRIVVYLLKSTMQYQYSWVAAAVEELVTLAFYVWTATKFRPTDQNPYLKLQEIE  424
```

>protein GPR108-like [Oncorhynchus mykiss]

Sequence ID: XP_021412142.1 Length: 310
Range 1: 8 to 309

Score:282 bits(722), Expect:1e-88,
Method:Compositional matrix adjust.,
Identities:146/304(48%), Positives:205/304(67%), Gaps:17/304(5%)

```
Query  140  EGFYTVVFLNCQ---PGTYVSFDLTL--TNYNPGPNYLSAGLTALPTLYAMLFVVWTVIL  194
            EG Y + F  C+   PG    + L +  T  NPG  YLSA    LP LY  +  V+ +I
Sbjct  8    EGLYNLDFHYCENILPGNNRPYTLNVEVTEKNPG-GYLSAAEIPLPRLYIFMAGVFFIIA  66

Query  195  GVWLFHFMRGQGKRIFRIHHLVTGIILLKLLTLLFEAIEFHYKKTTGHP-GGWVIAYYIF  253
             VW++  ++ +   +F+IH L+  +   K ++L+F +I +H+  T GHP  GW + YYI
Sbjct  67   MVWVYTLLKHR-YSVFKIHWLMAALAFTKSISLVFHSINYHFINTEGHPIEGWAVMYYIT  125

Query  254  SGLKGTMMFVVIALIGTGWAFIKPFLGEKDKNIFLVVIPLQILANIAIIVLEETA-----  308
              LKG ++F+ +ALIGTGWAF+K  L +K+K IF++VIPLQ+LAN+A I++E T
Sbjct  126  HLLKGALLFITLALIGTGWAFVKYILSDKEKKIFMIVIPLQVLANVAYIIIESTEEGSSE  185

Query  309  ----PEVLRLVDIICCGAILVPIIWSIKHLRDAAAIDGKAKRNMEKLKLFREFYLLVVTY  364
                 EVL LVD+ICCGAIL P+IWSI+HL++A++ DGKA  N+EKLKLFR +Y+++V Y
Sbjct  186  YALWKEVLFLVDLICCGAILFPVIWSIRHLQEASSTDGKAAMNLEKLKLFRHYYVMIVCY  245

Query  365  IYFTRIIVFLLDATLHYQYVWLGEFFTELATLIFWGLTGYKFRPVADNPYLKLDDEEEDA  424
            IYFTRII  LL  T+ +Q+ W  EF  E++TLIF+ LTGYKFRP ++NPYL+L  +EED
Sbjct  246  IYFTRIIAILLKVTVPFQWQWCYEFLVEVSTLIFFVLTGYKFRPASNNPYLQLPLDEEDV  305

Query  425  EREE  428
            E +E
Sbjct  306  EMDE  309
```

>protein GPR107 isoform 4 [Homo sapiens]
Sequence ID: NP_001274275.1 Length: 332
>protein GPR107 isoform X3 [Gorilla gorilla gorilla]
Sequence ID: XP_018889017.1 Length: 332 >G protein-coupled receptor 107, isoform CRA_e [Homo sapiens]
Sequence ID: EAW87925.1 Length: 332
Range 1: 22 to 331

Score:283 bits(724), Expect:1e-88,
Method:Compositional matrix adjust.,
Identities:148/312(47%), Positives:202/312(64%), Gaps:18/312(5%)

```
Query  133  TIDKTRPEGFYTVVFLNCQ----PGTYVSF--DLTLTNYNPGPNYLSAGLTALPTLYAML  186
             I      EG Y++ F  C      P    +F  D+ +T  NP  +YLSAG   LP LY  +
Sbjct  22   NISTDDQEGLYSLYFHKCLGKELPSDKFTFSLDIEITEKNPD-SYLSAGEIPLPKLYISM  80

Query  187  FVVWTVILGVWLFHFMRGQGKRIFRIHHLVTGIILLKLLTLLFEAIEFHYKKTTGHP-GG  245
               + +   +W+ H +R +   +F+IH L+  +   K L+L+F AI++HY  + G P  G
Sbjct  81   AFFFFLSGTIWI-HILRKRRNDVFKIHWLMAALPFTKSLSLVFHAIDYHYISSQGFPIEG  139

Query  246  WVIAYYIFSGLKGTMMFVVIALIGTGWAFIKPFLGEKDKNIFLVVIPLQILANIAIIVLE  305
```

```
            W + YYI   LKG ++F+ IALIGTGWAFIK  L +KDK IF++VIPLQ+LAN+A I++E
Sbjct  140  WAVVYYITHLLKGALLFITIALIGTGWAFIKHILSDKDKKIFMIVIPLQVLANVAYIIIE  199

Query  306  ETA---------PEVLRLVDIICCGAILVPIIWSIKHLRDAAAIDGKAKRNMEKLKLFRE  356
             T           + L LVD++CCGAIL P++WSI+HL++A+A DGKA  N+ KLKLFR 
Sbjct  200  STEEGTTEYGLWKDSLFLVDLLCCGAILFPVVWSIRHLQEASATDGKAAINLAKLKLFRH  259

Query  357  FYLLVVTYIYFTRIIVFLLDATLHYQYVWLGEFFTELATLIFWGLTGYKFRPVADNPYLK  416
            +Y+L+V YIYFTRII FLL   + +Q+ WL +   E ATL+F+ LTGYKFRP +DNPYL+
Sbjct  260  YYVLIVCYIYFTRIIAFLLKLAVPFQWKWLYQLLDETATLVFFVLTGYKFRPASDNPYLQ  319

Query  417  LDDEEEDAEREE  428
            L  EEED E E 
Sbjct  320  LSQEEEDLEMES  331
```

>G protein-coupled receptor 107 [Molossus molossus]
Sequence ID: KAF6433526.1 Length: 362
Range 1: 15 to 335

Score:284 bits(726), Expect:1e-88,
Method:Compositional matrix adjust.,
Identities:148/324(46%), Positives:205/324(63%), Gaps:21/324(6%)

```
Query  128  QSKVITIDKTRPEGFYTVVFLNCQPGTYV--------SFDLTLTNYNPGPNYLSAGLTAL  179
            Q     I     EG Y++ F  C PG  +        S D+ +T  NP  +YLSAG   L
Sbjct  15   QKFFFNISTNDQEGLYSLYFHKC-PGPKIQSNDKFLFSLDIEITEKNPD-SYLSAGEIPL  72

Query  180  PTLYAMLFVVWTVILGVWLFHFMRGQGKRIFRIHHLVTGIILLKLLTLLFEAIEFHYKKT  239
            P LY  +   + +   +W+ H +R +   +F+IH L+  +   K L+L+F AI++HY  +
Sbjct  73   PKLYIFMAFFFFLSGTIWI-HILRKRRNDVFKIHWLMAALPFTKSLSLVFHAIDYHYISS  131

Query  240  TGHP-GGWVIAYYIFSGLKGTMMFVVIALIGTGWAFIKPFLGEKDKNIFLVVIPLQILAN  298
             G P  GW + YYI   LKG ++F+ IALIGTGWAFIK  L +KDK IF++VIPLQ+LAN
Sbjct  132  QGFPIEGWAVVYYITHLLKGALLFITIALIGTGWAFIKHILSDKDKKIFMIVIPLQVLAN  191

Query  299  IAIIVLEETA---------PEVLRLVDIICCGAILVPIIWSIKHLRDAAAIDGKAKRNME  349
            +A I++E T           + L LVD++CCGAIL P++WSI+HL++A+A DGKA  N+
Sbjct  192  VAYIIIESTEEGTTEYALWKDSLFLVDLLCCGAILFPVVWSIRHLQEASATDGKAAINLA  251

Query  350  KLKLFREFYLLVVTYIYFTRIIVFLLDATLHYQYVWLGEFFTELATLIFWGLTGYKFRPV  409
            KLKLFR +Y+L+V YIYFTRII F L   + +Q+ WL +   E+ATL+F+ LTGYKFRP 
Sbjct  252  KLKLFRHYYVLIVCYIYFTRIIAFFLKFAVPFQWKWLYQLLDEMATLVFFVLTGYKFRPA  311

Query  410  ADNPYLKLDDEEEDAEREEAQRQS  433
            +DNPYL+L  E++D E E     S
Sbjct  312  SDNPYLQLSQEDDDLEMESVVTTS  335
```

>protein GPR107 isoform X3 [Rhinopithecus roxellana]
Sequence ID: XP_010359244.1 Length: 332
>PREDICTED: protein GPR107 isoform X3 [Colobus angolensis palliatus]

Sequence ID: XP_011786597.1 Length: 332 >protein GPR107 isoform X3 [Trachypithecus francoisi]
Sequence ID: XP_033049780.1 Length: 332
Range 1: 22 to 331

Score:283 bits(724), Expect:1e-88,
Method:Compositional matrix adjust.,
Identities:148/312(47%), Positives:202/312(64%), Gaps:18/312(5%)

```
Query  133  TIDKTRPEGFYTVVFLNCQ----PGTYVSF--DLTLTNYNPGPNYLSAGLTALPTLYAML  186
             I     EG Y++ F  C     P    SF  D+ +T  NP  +YLSAG   LP LY  +
Sbjct  22   NISTDDQEGLYSLYFHKCLGKELPSDKFSFSLDIEITEKNPD-SYLSAGEIPLPKLYISM  80

Query  187  FVVWTVILGVWLFHFMRGQGKRIFRIHHLVTGIILLKLLTLLFEAIEFHYKKTTGHP-GG  245
               + +   +W+ H +R +   +F+IH L+  +   K L+L+F AI++HY  + G P  G
Sbjct  81   AFFFFLSGTIWI-HILRKRRNDVFKIHWLMAALPFTKSLSLVFHAIDYHYISSQGFPIEG  139

Query  246  WVIAYYIFSGLKGTMMFVVIALIGTGWAFIKPFLGEKDKNIFLVVIPLQILANIAIIVLE  305
            W + YYI   LKG ++F+ IALIGTGWAFIK  L +KDK IF++VIPLQ+LAN+A I++E
Sbjct  140  WAVVYYITHLLKGALLFITIALIGTGWAFIKHILSDKDKKIFMIVIPLQVLANVAYIIIE  199

Query  306  ETA---------PEVLRLVDIICCGAILVPIIWSIKHLRDAAAIDGKAKRNMEKLKLFRE  356
             T           + L LVD++CCGAIL P++WSI+HL++A+A DGKA  N+ KLKLFR
Sbjct  200  STEEGTTEYGLWKDSLFLVDLLCCGAILFPVVWSIRHLQEASATDGKAAINLAKLKLFRH  259

Query  357  FYLLVVTYIYFTRIIVFLLDATLHYQYVWLGEFFTELATLIFWGLTGYKFRPVADNPYLK  416
            +Y+L+V YIYFTRII FLL   + +Q+ WL +   E ATL+F+ LTGYKFRP +DNPYL+
Sbjct  260  YYVLIVCYIYFTRIIAFLLKLAVPFQWKWLYQLLDETATLVFFVLTGYKFRPASDNPYLQ  319

Query  417  LDDEEEDAEREE  428
            L  EE+D E E
Sbjct  320  LSQEEDDLEMES  331
```

>protein GPR107 isoform X2 [Pteropus vampyrus]
Sequence ID: XP_011362432.1 Length: 333
Range 1: 22 to 332

Score:283 bits(724), Expect:2e-88,
Method:Compositional matrix adjust.,
Identities:146/313(47%), Positives:201/313(64%), Gaps:19/313(6%)

```
Query  133  TIDKTRPEGFYTVVFLNC-------QPGTYVSFDLTLTNYNPGPNYLSAGLTALPTLYAM  185
             I     EG Y++ F  C             S D+ +T  NP  +YLSAG   LP LY
Sbjct  22   NISTDDQEGLYSLYFHKCLGNKMWSNDKFSFSLDIEITEKNPD-SYLSAGEIPLPKLYIS  80

Query  186  LFVVWTVILGVWLFHFMRGQGKRIFRIHHLVTGIILLKLLTLLFEAIEFHYKKTTGHP-G  244
            +   + +   VW+ H +R +   +F+IH L+  +   K L+L+F AI++HY  + G P
Sbjct  81   MAFFFFLSGTVWI-HILRKRRNEVFKIHWLMAALPFTKSLSLVFHAIDYHYISSQGFPIE  139

Query  245  GWVIAYYIFSGLKGTMMFVVIALIGTGWAFIKPFLGEKDKNIFLVVIPLQILANIAIIVL  304
            GW + YYI   LKG ++F+ IALIGTGWAFIK  L +KDK IF++VIPLQ+LAN+A I++
Sbjct  140  GWAVVYYITHLLKGALLFITIALIGTGWAFIKHILSDKDKKIFMIVIPLQVLANVAYIII  199
```

```
Query  305  EETA---------PEVLRLVDIICCGAILVPIIWSIKHLRDAAAIDGKAKRNMEKLKLFR  355
            E T           + L L+D++CCGAIL P++WSI+HL++A+A DGKA  N+ KLKLFR
Sbjct  200  ESTEEGTTEYSLWKDSLFLIDLLCCGAILFPVVWSIRHLQEASATDGKAAINLAKLKLFR  259

Query  356  EFYLLVVTYIYFTRIIVFLLDATLHYQYVWLGEFFTELATLIFWGLTGYKFRPVADNPYL  415
             +Y+L+V YIYFTRII FLL   + +Q+ WL +   E+ATL+F+ LTGYKFRP +DNPYL
Sbjct  260  HYYVLIVCYIYFTRIIAFLLKFAVPFQWKWLYQLLDEMATLVFFVLTGYKFRPASDNPYL  319

Query  416  KLDDEEEDAEREE  428
            +L  EE+D E E
Sbjct  320  QLSQEEDDLEMES  332
```

>protein GPR107 isoform X2 [Rhincodon typus]
Sequence ID: XP_020386815.1 Length: 555
Range 1: 127 to 523

Score:290 bits(742), Expect:2e-88,
Method:Compositional matrix adjust.,
Identities:173/408(42%), Positives:247/408(60%), Gaps:42/408(10%)

```
Query  52   TGGVMEMAIENFKV-----VDDKGSLWDDLS------AGFIIKHIETDSGSFIEETDASK  100
            T G  +  IE+F+      V DK S  D  S      AG + K  ET   S  E+   +K
Sbjct  127  TSGNNQPTIEDFRTAASQPVKDKSSKTDVKSQSESKKAGELQKKNETTKQS--EQVQMAK  184

Query  101  CVSLLDEPERGDTIAVKVTKPSKENEKQSKVI--TIDKTRPEGFYTVVFLNC------QP  152
              S +    + +T+ ++     K+N   S     TI+    +G Y++ F NC       P
Sbjct  185  PKSGISAERKDETLPLQ-----KKNLSYSFTFSFTINSEEEQGLYSLYFHNCYTDEKPSP  239

Query  153  GTY-VSFDLTLTNYNPGPNYLSAGLTALPTLY-AMLFVVWTVILGVWLFHFMRGQGKRIF  210
             T+    D+T+   NPG +YLSAG   LP LY  M F  +  + GV+  + +R +   +F
Sbjct  240  QTFQFDMDITIVEQNPG-SYLSAGEIPLPKLYICMAFFFF--LAGVFWVYILRKRSD-VF  295

Query  211  RIHHLVTGIILLKLLTLLFEAIEFHYKKTTGHP-GGWVIAYYIFSGLKGTMMFVVIALIG  269
            +IH L+  +   K L+L+F AI++H+  T G+P  GW + YYI   LKG ++F+ I+LIG
Sbjct  296  KIHWLMGALAFTKSLSLVFHAIDYHFISTQGYPIEGWAVVYYITHLLKGALLFITISLIG  355

Query  270  TGWAFIKPFLGEKDKNIFLVVIPLQILANIAIIVLEETA---------PEVLRLVDIICC  320
            TGWAF+K  L +KDK IF++VIPLQ+LAN+A I++E T           E+L LVD++CC
Sbjct  356  TGWAFVKHILSDKDKKIFMIVIPLQVLANVAYIIIESTEEGTTEYGLWKEILFLVDLLCC  415

Query  321  GAILVPIIWSIKHLRDAAAIDGKAKRNMEKLKLFREFYLLVVTYIYFTRIIVFLLDATLH  380
            GAIL P++WSI+HL++A+A DGKA  N+ KLKLFR +Y+++V YIYFTRII  L+   +
Sbjct  416  GAILFPVVWSIRHLQEASATDGKAAVNLAKLKLFRHYYVMIVCYIYFTRIIAILIKIIVP  475

Query  381  YQYVWLGEFFTELATLIFWGLTGYKFRPVADNPYLKLDDEEEDAEREE  428
            +Q+ WL +F  ELATL+F+ LTGYKFRP +DNPYL+L   EED E +E
Sbjct  476  FQWKWLYQFLDELATLVFFCLTGYKFRPASDNPYLQLPQNEEDVEMDE  523
```

```
Range 2: 13 to 99

Score:43.1 bits(100), Expect:2.6,
Method:Compositional matrix adjust.,
Identities:30/89(34%), Positives:41/89(46%), Gaps:4/89(4%)

Query  15   IFCLVAFLAPTANGLIHKLSIKNDRRLAFRIETFGFFTGGVMEMAIENF--KVVDDKGSL  72
            +F L   L  +  G IH+L +K+D R   R+ TFGFF  G M + + N   K VD K
Sbjct  13   MFLLWILLLTSVRGRIHRLILKDDVRQKIRLNTFGFFKNGNMTVKMTNLTSKGVDFKAV-  71

Query  73   WDDLSAGFIIKHIETDSGSFIEETDASKC  101
             D  + G  +   + D  S   E D   C
Sbjct  72   -DTSALGLSLDRTKNDGFSSYLEEDIDFC  99



>protein CANDIDATE G-PROTEIN COUPLED RECEPTOR 7 [Cucumis sativus]
Sequence ID: XP_011650293.1 Length: 439
>hypothetical protein Csa_011022 [Cucumis sativus]
Sequence ID: KAE8650004.1 Length: 439
Range 1: 11 to 421

Score:286 bits(732), Expect:2e-88,
Method:Compositional matrix adjust.,
Identities:166/431(39%), Positives:244/431(56%), Gaps:46/431(10%)

Query  15   IFCLVAFLAPTAN-GLIHKLSIKNDRRLAFRIETFGFFTGGVMEMAIENFKVVDDKGSLW  73
            +F L+ FL P ++   IH   I+ND R     + FGF  GG +E+ + +  + D    L
Sbjct  11   MFILLIFLLPISSFAEIHFTEIRNDNRPIIPFDVFGFSHGGRLELNVSHLTLSDSNPDL-  69

Query  74   DDLS-AGF----------IIKHIETDSGSFIEETDASKCVSLLDEPERGDTIAVKVTKPS  122
             DLS  GF          +I+ +E    S   ++D  K V   D  ++ D   V  ++
Sbjct  70   -DLSKVGFFLCTRESWLHVIQQLEEGDISCALQSDLVKPVYTFDSLKKQDRFGVLYSETD  128

Query  123  KENEKQSKVITIDKTRPEGFYTVVFLNCQPGTYVSFDLTLTNYN-PGPN----YLSAGLT  177
             +                  YT+VF NC     VS D+    YN  G N    YLSAG T
Sbjct  129  ADQ-----------------YTLVFANCLQQVKVSMDVQSAMYNLEGKNARRDYLSAGKT  171

Query  178  ALPTLYAMLFVVWTVILGVWLFHFMRGQGKRIFRIHHLVTGIILLKLLTLLFEAIEFHYK  237
             LP +Y +  +++  +  +W+ H +  +   ++ IH  +  +++LK L LL EA +  Y
Sbjct  172  ILPRIYFVFSLIYFSLAVIWI-HVLYKKRLTVYGIHFFMLAVVILKALNLLCEAEDKSYI  230

Query  238  KTTGHPGGWVIAYYIFSGLKGTMMFVVIALIGTGWAFIKPFLGEKDKNIFLVVIPLQILA  297
            K TG   GW + +YIFS LKG  +F +I LIGTGW+F+KP+L +K+K + ++VIPLQ++A
Sbjct  231  KRTGSAHGWDVLFYIFSFLKGITLFTLIVLIGTGWSFLKPYLQDKEKKVLMIVIPLQVVA  290

Query  298  NIAIIVLEETAP---------EVLRLVDIICCGAILVPIIWSIKHLRDAAAIDGKAKRNM  348
            NIA +V++ET P         +V  LVD+ICC A+L PI+WSIK+LR+AA  DGKA  N+
Sbjct  291  NIAQVVIDETGPFEQEWVTWKQVFLLVDVICCCAVLFPIVWSIKNLREAARTDGKAAVNL  350

Query  349  EKLKLFREFYLLVVTYIYFTRIIVFLLDATLHYQYVWLGEFFTELATLIFWGLTGYKFRP  408
             KL LFR++Y++V+ YIYFTR++V+ L+    Y+Y+W      ELATL F+  TGYKF+P
Sbjct  351  MKLTLFRQYYIVVICYIYFTRVVVYALETITSYRYLWTSVMAGELATLAFYVFTGYKFKP  410
```

```
Query  409  VADNPYLKLDD  419
             A NPY  +DD
Sbjct  411  EAHNPYFVVDD  421
```

>protein GPR107 isoform X2 [Papio anubis]
Sequence ID: XP_021782822.2 Length: 363
>protein GPR107 isoform X2 [Papio anubis]
Sequence ID: XP_031510497.1 Length: 363
Range 1: 22 to 336

Score:283 bits(725), Expect:3e-88,
Method:Compositional matrix adjust.,
Identities:149/317(47%), Positives:203/317(64%), Gaps:18/317(5%)

```
Query  133  TIDKTRPEGFYTVVFLNCQ----PGTYVSF--DLTLTNYNPGPNYLSAGLTALPTLYAML  186
             I     EG Y++ F  C      P    SF  D+ +T  NP  +YLSAG   LP LY  +
Sbjct  22   NISTDDQEGLYSLYFHKCLGKELPSDKFSFSLDIEITEKNPD-SYLSAGEIPLPKLYISM  80

Query  187  FVVWTVILGVWLFHFMRGQGKRIFRIHHLVTGIILLKLLTLLFEAIEFHYKKTTGHP-GG  245
               + +   +W+ H +R +   +F+IH L+  +   K L+L+F AI++HY  + G P  G
Sbjct  81   AFFFFLSGTIWI-HILRKRRNDVFKIHWLMAALPFTKSLSLVFHAIDYHYISSQGFPIEG  139

Query  246  WVIAYYIFSGLKGTMMFVVIALIGTGWAFIKPFLGEKDKNIFLVVIPLQILANIAIIVLE  305
            W + YYI   LKG ++F+ IALIGTGWAFIK  L +KDK IF++VIPLQ+LAN+A I++E
Sbjct  140  WAVVYYITHLLKGALLFITIALIGTGWAFIKHILSDKDKKIFMIVIPLQVLANVAYIIIE  199

Query  306  ETA---------PEVLRLVDIICCGAILVPIIWSIKHLRDAAAIDGKAKRNMEKLKLFRE  356
             T           + L LVD++CCGAIL P++WSI+HL++A+A DGKA  N+ KLKLFR
Sbjct  200  STEEGTTEYGLWKDSLFLVDLLCCGAILFPVVWSIRHLQEASATDGKAAINLAKLKLFRH  259

Query  357  FYLLVVTYIYFTRIIVFLLDATLHYQYVWLGEFFTELATLIFWGLTGYKFRPVADNPYLK  416
            +Y+L+V YIYFTRII FLL   + +Q+ WL +   E ATL+F+ LTGYKFRP +DNPYL+
Sbjct  260  YYVLIVCYIYFTRIIAFLLKLAVPFQWKWLYQLLDETATLVFFVLTGYKFRPASDNPYLQ  319

Query  417  LDDEEEDAEREEAQRQS  433
            L  EE+D E E     S
Sbjct  320  LSQEEDDLEMESVVTTS  336
```

>G protein-coupled receptor 107 [Molossus molossus]
Sequence ID: KAF6433525.1 Length: 412
Range 1: 68 to 385

Score:285 bits(729), Expect:3e-88,
Method:Compositional matrix adjust.,
Identities:147/321(46%), Positives:206/321(64%), Gaps:21/321(6%)

```
Query  131  VITIDKTRPEGFYTVVFLNCQPGTYV--------SFDLTLTNYNPGPNYLSAGLTALPTL  182
            V+ +     EG Y++ F  C PG  +        S D+ +T  NP  +YLSAG   LP L
Sbjct  68   VVNVSTNDQEGLYSLYFHKC-PGPKIQSNDKFLFSLDIEITEKNPD-SYLSAGEIPLPKL  125
```

```
Query  183  YAMLFVVWTVILGVWLFHFMRGQGKRIFRIHHLVTGIILLKLLTLLFEAIEFHYKKTTGH  242
            Y  +   + +   +W+ H +R +   +F+IH L+  +   K L+L+F AI++HY  + G
Sbjct  126  YIFMAFFFFLSGTIWI-HILRKRRNDVFKIHWLMAALPFTKSLSLVFHAIDYHYISSQGF  184

Query  243  P-GGWVIAYYIFSGLKGTMMFVVIALIGTGWAFIKPFLGEKDKNIFLVVIPLQILANIAI  301
            P  GW + YYI   LKG ++F+ IALIGTGWAFIK  L +KDK IF++VIPLQ+LAN+A
Sbjct  185  PIEGWAVVYYITHLLKGALLFITIALIGTGWAFIKHILSDKDKKIFMIVIPLQVLANVAY  244

Query  302  IVLEETA---------PEVLRLVDIICCGAILVPIIWSIKHLRDAAAIDGKAKRNMEKLK  352
            I++E T           + L LVD++CCGAIL P++WSI+HL++A+A DGKA  N+ KLK
Sbjct  245  IIIESTEEGTTEYALWKDSLFLVDLLCCGAILFPVVWSIRHLQEASATDGKAAINLAKLK  304

Query  353  LFREFYLLVVTYIYFTRIIVFLLDATLHYQYVWLGEFFTELATLIFWGLTGYKFRPVADN  412
            LFR +Y+L+V YIYFTRII F L   + +Q+ WL +   E+ATL+F+ LTGYKFRP +DN
Sbjct  305  LFRHYYVLIVCYIYFTRIIAFFLKFAVPFQWKWLYQLLDEMATLVFFVLTGYKFRPASDN  364

Query  413  PYLKLDDEEEDAEREEAQRQS  433
            PYL+L  E++D E E     S
Sbjct  365  PYLQLSQEDDDLEMESVVTTS  385
```

>hypothetical protein [Chiloscyllium punctatum]
Sequence ID: GCC30454.1 Length: 560
Range 1: 217 to 528

Score:289 bits(740), Expect:3e-88,
Method:Compositional matrix adjust.,
Identities:148/315(47%), Positives:212/315(67%), Gaps:21/315(6%)

```
Query  132  ITIDKTRPEGFYTVVFLNC------QPGTY-VSFDLTLTNYNPGPNYLSAGLTALPTLY-  183
            +TI+    +G Y++ F NC       P ++    D+T+   NPG +YLSAG   LP LY
Sbjct  217  LTINSEEEQGLYSLYFHNCYTDEKPSPQSFQFDMDITIVEQNPG-SYLSAGEIPLPKLYI  275

Query  184  AMLFVVWTVILGVWLFHFMRGQGKRIFRIHHLVTGIILLKLLTLLFEAIEFHYKKTTGHP  243
            +M F  +  + GV+  + +R +   +F+IH L+  +   K L+L+F AI++H+  T G+P
Sbjct  276  SMAFFFF--LAGVFWVYILRKRSSDVFKIHWLMGALAFTKSLSLVFHAIDYHFISTQGYP  333

Query  244  -GGWVIAYYIFSGLKGTMMFVVIALIGTGWAFIKPFLGEKDKNIFLVVIPLQILANIAII  302
              GW + YYI   LKG ++F+ I+LIGTGWAF+K  L +KDK IF++VIPLQ+LAN+A I
Sbjct  334  IEGWAVVYYITHLLKGALLFITISLIGTGWAFVKHILSDKDKKIFMIVIPLQVLANVAYI  393

Query  303  VLEETA---------PEVLRLVDIICCGAILVPIIWSIKHLRDAAAIDGKAKRNMEKLKL  353
            ++E T           E+L LVD++CCGAIL P++WSI+HL++A+A DGKA  N+ KLKL
Sbjct  394  IIESTEEGTTEYGLWKEILFLVDLLCCGAILFPVVWSIRHLQEASATDGKAAVNLAKLKL  453

Query  354  FREFYLLVVTYIYFTRIIVFLLDATLHYQYVWLGEFFTELATLIFWGLTGYKFRPVADNP  413
            FR +Y+++V YIYFTRII  L+   + +Q+ WL +F  ELATL+F+ LTGYKFRP +DNP
Sbjct  454  FRHYYVMIVCYIYFTRIIAILIKIIVPFQWKWLYQFLDELATLVFFCLTGYKFRPASDNP  513

Query  414  YLKLDDEEEDAEREE  428
            YL+L   EED E +E
Sbjct  514  YLQLPQNEEDVEMDE  528
```

```
Range 2: 19 to 104

Score:42.7 bits(99), Expect:3.7,
Method:Compositional matrix adjust.,
Identities:30/88(34%), Positives:40/88(45%), Gaps:4/88(4%)

Query  16   FCLVAFLAPTANGLIHKLSIKNDRRLAFRIETFGFFTGGVM--EMAIENFKVVDDKGSLW  73
            F +  FL+ +  G IH+L +K+D R    + TFGFF  G M  EM     K VD K
Sbjct  19   FLICMFLSASVQGRIHRLILKDDIRQKIHLNTFGFFKNGNMTVEMTSLTSKEVDFKTV--  76

Query  74   DDLSAGFIIKHIETDSGSFIEETDASKC  101
            D  + G  +   + D  S   E D   C
Sbjct  77   DTSALGLSLDKTKNDGFSSYLEEDIDFC  104



>PREDICTED: protein GPR107 [Daucus carota subsp. sativus]
Sequence ID: XP_017248308.1 Length: 438
>hypothetical protein DCAR_014560 [Daucus carota subsp. sativus]
Sequence ID: KZM98078.1 Length: 438
Range 1: 10 to 420

Score:285 bits(730), Expect:4e-88,
Method:Compositional matrix adjust.,
Identities:164/422(39%), Positives:238/422(56%), Gaps:27/422(6%)

Query  14   LIFCLVAFLAPTANGLIHKLSIKNDRRLAFRIETFGFFTGGVMEMAIENFKVVDDKGSLW  73
            LI   +A   P     I    I++D RL    + FGF   G +++ I N K+ + +
Sbjct  10   LIIFFIASSIPLTLAEIRVTQIRSDNRLTIPFDEFGFTHSGRLDLNISNIKLSNPRVPKT  69

Query  74   DDLSAGFIIKHIETDSGSFIEETDASKCVSLLDEPERGDTIAVKVTKP--SKENEKQSKV  131
            +    GF +    T   S++   D       L E E   ++   + KP    +N    +V
Sbjct  70   ELSQVGFFL----TTRESWLHVIDQ------LIEREIPCSLNSNLIKPVFKFDNLDGDRV  119

Query  132  ITIDKTRPEGFYTVVFLNCQPGTYVSFDLTLTNYNPGPN-----YLSAGLTALPTLYAML  186
                +      YT+VF NC P   VS ++    YN  P      YLSAG T+LP++Y  +
Sbjct  120  DVSFEANDANQYTLVFANCVPNLKVSMNVKSAMYNVEPRTGQRVYLSAGKTSLPSIYFVF  179

Query  187  FVVWTVILGVWLFHFMRGQGKRIFRIHHLVTGIILLKLLTLLFEAIEFHYKKTTGHPGGW  246
            F+++  + G W+F  +  +   +FRIH  +  +++LK L LL E  +  Y K TG   GW
Sbjct  180  FLMYISLAGFWIFT-IYSKILSVFRIHFFMLAVVILKALNLLCETEDKSYIKRTGTAHGW  238

Query  247  VIAYYIFSGLKGTMMFVVIALIGTGWAFIKPFLGEKDKNIFLVVIPLQILANIAIIVLEE  306
             + +YIFS LKG  +F +I LIGTGW+FIKP+L  K+K + ++VIPLQ++AN+A IV++E
Sbjct  239  DVLFYIFSFLKGITLFTLIVLIGTGWSFIKPYLQGKEKKVLMIVIPLQVIANVAQIVIDE  298

Query  307  TAP---------EVLRLVDIICCGAILVPIIWSIKHLRDAAAIDGKAKRNMEKLKLFREF  357
            T P         +V  LVDI+CC A+L PI+WSIK+LR+AA  DGKA  N+ KL LFR++
Sbjct  299  TGPFGEDSDMWKKVFLLVDIVCCCAVLFPIVWSIKNLREAARTDGKAAVNLMKLTLFRQY  358

Query  358  YLLVVTYIYFTRIIVFLLDATLHYQYVWLGEFFTELATLIFWGLTGYKFRPVADNPYLKL  417
```

```
            Y++V+ YIYFTR++V+ L+    Y+Y W      ELATL F+  TGY FRP A NPY  +
Sbjct  359  YVVVICYIYFTRVVVYALETITSYRYEWTSVVAAELATLAFYIFTGYNFRPKAHNPYFAI  418

Query  418  DD  419
            DD
Sbjct  419  DD  420
```

>protein GPR107-like [Cynara cardunculus var. scolymus]
Sequence ID: XP_024985659.1 Length: 438
>Transmembrane receptor, eukaryota [Cynara cardunculus var. scolymus]
Sequence ID: KVH90584.1 Length: 438
Range 1: 1 to 437

Score:285 bits(729), Expect:5e-88,
Method:Compositional matrix adjust.,
Identities:174/449(39%), Positives:250/449(55%), Gaps:26/449(5%)

```
Query  2    MRDRRQTLMGCFLIFCLVAFLAPTANGLIHKLSIKNDRRLAFRIETFGFFTGGVMEMAIE  61
            MR   + L    L F L  F AP+  G I  L I++D R     E FGF   G + +AI
Sbjct  1    MRILGKILPAVLLFFFL--FTAPSM-GEIKSLKIRSDNRPMILFEKFGFTHTGFVSIAIS  57

Query  62   NFKVVDDKGSLWDDLSAGFIIKHIETDSGSFIE-ETDASKCVSLLDEPERGDTIAVKVTK  120
            +  V        D    GF +   E+     +E + + + CV  +D          +
Sbjct  58   SVSVTSTLSQP-DPSRLGFFLLSEESQIQVLLELQQNPNFCV--VDSKFIARLFTFRDLS  114

Query  121  PSKENEKQSKVITIDKTRPEGFYTVVFLNCQPGTYVSFDLTLTNYN----PGPNYLSAGL  176
            P  ++       T   T P   Y++ F NC P + V+ D+    YN       +YLSAGL
Sbjct  115  PPPQSSFNR---TYPVTHPNE-YSLFFANCNPQSLVTMDVHTELYNTDDGSTKDYLSAGL  170

Query  177  TALPTLYAMLFVVWTVILGVWLFHFMRGQGKRIFRIHHLVTGIILLKLLTLLFEAIEFHY  236
            T LP+LY +  +++   LG W+    + Q + + RIH L+ G++++K L L+  A + HY
Sbjct  171  TQLPSLYFIFSLIYLSFLGFWISICFKNQ-RSVHRIHLLMGGLLVMKALNLICAAEDKHY  229

Query  237  KKTTGHPGGWVIAYYIFSGLKGTMMFVVIALIGTGWAFIKPFLGEKDKNIFLVVIPLQIL  296
             K TG P GW + +YIF  ++  ++F VI LIGTGW+F+KPFL EK+K + ++VIPLQ+L
Sbjct  230  VKVTGTPHGWDVLFYIFQFIRVVLLFTVIVLIGTGWSFLKPFLQEKEKKVLMIVIPLQVL  289

Query  297  ANIAIIVLEETAP---------EVLRLVDIICCGAILVPIIWSIKHLRDAAAIDGKAKRN  347
            AN+A IV+ ET P         +V  LVDIICC AI+ PI+WSI+ LR+ +  DGKA RN
Sbjct  290  ANVASIVIGETGPFIRDWVTWNQVFLLVDIICCCAIIFPIVWSIRSLRETSKTDGKAARN  349

Query  348  MEKLKLFREFYLLVVTYIYFTRIIVFLLDATLHYQYVWLGEFFTELATLIFWGLTGYKFR  407
            + KL LFR+FY++V+ Y+YFTRI+VF L     Y+Y W+     E+A+L+F+ +  Y FR
Sbjct  350  LAKLTLFRQFYIVVIGYLYFTRIVVFALKTIAAYKYQWVANAAEEIASLVFYMVMYYMFR  409

Query  408  PVADNPYLKLDDEEEDAEREEAQRQSRTE  436
            PV  N Y  LDDE+E+A  E A R    E
Sbjct  410  PVEKNEYFVLDDEDEEAA-EMALRDEEFE  437
```

>PREDICTED: protein GPR107 [Charadrius vociferus]

Sequence ID: XP_009886300.1 Length: 334
Range 1: 22 to 333

Score:281 bits(719), Expect:9e-88,
Method:Compositional matrix adjust.,
Identities:144/314(46%), Positives:202/314(64%), Gaps:19/314(6%)

```
Query  133  TIDKTRPEGFYTVVFLNC-------QPGTYVSFDLTLTNYNPGPNYLSAGLTALPTLYAM  185
             I     EG Y++ F  C             S D+ +T  NP  +YLSAG   LP LY
Sbjct  22   NISSDNQEGLYSLYFHKCFGNGGPTNDQQLFSLDIEITEKNP-ESYLSAGEIPLPKLYIS  80

Query  186  LFVVWTVILGVWLFHFMRGQGKRIFRIHHLVTGIILLKLLTLLFEAIEFHYKKTTGHP-G  244
            + + + +   VW+ H +R +   +F+IH L+  +   K L+L+F AI++HY  + G P
Sbjct  81   MAIFFFLSGTVWI-HILRKRRNDVFKIHWLMAALPFTKSLSLVFHAIDYHYISSQGFPIE  139

Query  245  GWVIAYYIFSGLKGTMMFVVIALIGTGWAFIKPFLGEKDKNIFLVVIPLQILANIAIIVL  304
            GW + YYI   LKG ++F+ IALIGTGWAFIK  L +KDK IF++VIPLQ+LAN+A I++
Sbjct  140  GWAVVYYITHLLKGALLFITIALIGTGWAFIKHILSDKDKKIFMIVIPLQVLANVAYIII  199

Query  305  EETA---------PEVLRLVDIICCGAILVPIIWSIKHLRDAAAIDGKAKRNMEKLKLFR  355
            E T           E+L LVD++CCGAIL P++WSI+HL++A+A DGKA  N+ KLKLFR
Sbjct  200  ESTEEGTTEYGLWKEILFLVDLLCCGAILFPVVWSIRHLQEASATDGKAAINLAKLKLFR  259

Query  356  EFYLLVVTYIYFTRIIVFLLDATLHYQYVWLGEFFTELATLIFWGLTGYKFRPVADNPYL  415
             +Y+++V YIYFTRII  L+   + +Q+ WL +   E+ATL+F+ LTGYKFRP +DNPYL
Sbjct  260  HYYVMIVCYIYFTRIIAILIKIAVPFQWKWLYQLLDEMATLVFFVLTGYKFRPASDNPYL  319

Query  416  KLDDEEEDAEREEA  429
            +L  ++ED    EA
Sbjct  320  QLSQDDEDDLEMEA  333
```

>hypothetical protein [Gossypium davidsonii]
Sequence ID: MBA0621736.1 Length: 436
>hypothetical protein [Gossypium klotzschianum]
Sequence ID: MBA0657215.1 Length: 436
Range 1: 7 to 435

Score:283 bits(724), Expect:3e-87,
Method:Compositional matrix adjust.,
Identities:163/442(37%), Positives:251/442(56%), Gaps:31/442(7%)

```
Query  13   FLIFCLVAFLAPTANGLIHKLSIKNDRRLAFRIETFGFFTGGVMEMAIENFKVVDDKGSL  72
            F++   ++FL    +  I    I++D R     + FGF   G +E+ +   ++ D   +L
Sbjct  7    FVLLLFMSFLVSFGSAEIRFTEIRSDGRPIIPFDKFGFTHTGRLELNVSQVELSDSNRNL  66

Query  73   WDDLSAGFIIKHIETDSGSFIEETDA-SKCVSLLDEPERG---DTIAVKVTKPSKENEKQ  128
              DLS            G F+   DA  + +  L++ E     D+  +K+    K    +
Sbjct  67   --DLS----------KVGFFLCTLDAWMRVLQQLEDEEVACVLDSDLIKLVSNFKSLNGK  114

Query  129  SKVITIDKTRPEGFYTVVFLNCQPGTYVSFDLTLTNYN-----PGPNYLSAGLTALPTLY  183
            S    + + +    YT+VF NC     VS  +    YN        +YLSAG T LP +Y
Sbjct  115  SSFNFLYEEKDADQYTLVFANCLNQVKVSMKVRSAMYNLDGKKNRRDYLSAGETVLPRVY  174
```

```
Query  184  AMLFVVWTVILGVWLFHFMRGQGKRIFRIHHLVTGIILLKLLTLLFEAIEFHYKKTTGHP  243
             +L +V+  + GVW++   + +   +F IH  +  +++LK   L+ EA +  Y K TG
Sbjct  175  FLLSLVYFTLAGVWIYVLYKKR-LTVFGIHFFMLAVVILKAFNLVCEAEDKSYIKRTGSA  233

Query  244  GGWVIAYYIFSGLKGTMMFVVIALIGTGWAFIKPFLGEKDKNIFLVVIPLQILANIAIIV  303
             GW + +YIFS LKG  +F +I LIGTGW+F+KP+L +K+K + ++VIPLQ++ANIA +V
Sbjct  234  HGWDVLFYIFSFLKGITLFTLIVLIGTGWSFLKPYLQDKEKKVLMIVIPLQVVANIAQVV  293

Query  304  LEETAP---------EVLRLVDIICCGAILVPIIWSIKHLRDAAAIDGKAKRNMEKLKLF  354
            ++ET P         +V  LVD+ICC A+L PI+WSIK+LR+AA  DGKA  N+ KL LF
Sbjct  294  IDETGPFSQDWNTWKQVFLLVDVICCCAVLFPIVWSIKNLREAAKTDGKAAVNLMKLTLF  353

Query  355  REFYLLVVTYIYFTRIIVFLLDATLHYQYVWLGEFFTELATLIFWGLTGYKFRPVADNPY  414
            R++Y++V+ YIYFTR++V+ L+    Y+Y+W      ELATL F+  TG+KF+P A NPY
Sbjct  354  RQYYIVVICYIYFTRVVVYALETITSYKYLWTSVVAGELATLAFYVFTGFKFKPEAHNPY  413

Query  415  LKLDDEEEDAEREEAQRQSRTE  436
              +DDEEE+A   + + +   E
Sbjct  414  FVIDDEEEEAAAGQLKLEDEFE  435
```

>hypothetical protein [Gossypium raimondii]
Sequence ID: MBA0588979.1 Length: 436
Range 1: 31 to 435

Score:283 bits(724), Expect:3e-87,
Method:Compositional matrix adjust.,
Identities:159/416(38%), Positives:240/416(57%), Gaps:25/416(6%)

```
Query  35   IKNDRRLAFRIETFGFFTGGVMEMAIENFKVVDDKGSLWDDLSAGFIIKHIETDSGSFIE  94
            I++D R     + FGF   G +E+   N   +D  G   D    GF +   +T      +
Sbjct  31   IRSDVRPIIPFDEFGFTHNGRLEL---NLSQIDLSGKNLDLNKIGFFLCTRDTWFHVLEQ  87

Query  95   ETDASKCVSLLDEPERGDTIAVKVTKPSKENEKQSKVITIDKTRPEGFYTVVFLNCQPGT  154
              D     +L       D+  VKV    +  + ++ V  +        YT++F NC
Sbjct  88   LNDHHVTCAL-------DSDLVKVVFRFESLKGKTSVNAVFPVNNADQYTLLFANCLTQV  140

Query  155  YVSFDLTLTNYN-----PGPNYLSAGLTALPTLYAMLFVVWTVILGVWLFHFMRGQGKRI  209
             VS  +    YN        +YLSAG T LP +Y +L +V+  + G+W++ F+  +   +
Sbjct  141  KVSMTVRSAMYNLEGKQNSRDYLSAGKTILPRVYFLLSLVYFSLAGIWVY-FLYKKRLTV  199

Query  210  FRIHHLVTGIILLKLLTLLFEAIEFHYKKTTGHPGGWVIAYYIFSGLKGTMMFVVIALIG  269
            FRIH  +  +I+LK   L+FEA +  Y K TG   GW + +YIFS LKG M+F +I LIG
Sbjct  200  FRIHFFMLAVIVLKAFNLVFEAEDKSYIKRTGSAHGWDVLFYIFSFLKGIMLFTLIVLIG  259

Query  270  TGWAFIKPFLGEKDKNIFLVVIPLQILANIAIIVLEETAP---------EVLRLVDIICC  320
            TGW+F+KP+L +K+K + ++VIPLQ++ANIA +V++E +P         ++  LVD+ICC
Sbjct  260  TGWSFLKPYLQDKEKKVLMIVIPLQVVANIAQVVIDEASPFGQDRVTWKQLFLLVDVICC  319

Query  321  GAILVPIIWSIKHLRDAAAIDGKAKRNMEKLKLFREFYLLVVTYIYFTRIIVFLLDATLH  380
             A+L PI+WSIK+LR+AA  DGKA  N+ KL LFR++Y++V+ YIYFTR++V+ L+
Sbjct  320  CAVLFPIVWSIKNLREAARTDGKAAVNLMKLTLFRQYYVVVICYIYFTRVVVYALETITS  379
```

```
Query  381  YQYVWLGEFFTELATLIFWGLTGYKFRPVADNPYLKLDDEEEDAEREEAQRQSRTE  436
            Y+Y+W      ELATL F+  TGYKF+P A NPY  +D E+E+A  E+ + +   E
Sbjct  380  YKYLWTAVLAGELATLAFYVFTGYKFKPEAHNPYFAIDGEDEEAAAEQLKLEDEFE  435
```

>Lung seven transmembrane receptor-like [Parasponia andersonii]
Sequence ID: PON73519.1 Length: 441
Range 1: 13 to 423

Score:283 bits(724), Expect:4e-87,
Method:Compositional matrix adjust.,
Identities:160/419(38%), Positives:239/419(57%), Gaps:23/419(5%)

```
Query  16   FCLVAFLAPTANGLIHKLSIKNDRRLAFRIETFGFFTGGVMEMAIENFKVVDDKGSLWDD  75
            F L+    P A   I    I+ND R    ++ FGF   G +E+++    +         +
Sbjct  13   FFLLLLTIPMAFSEIRFSEIRNDLRPIIPLDEFGFTHKGRLELSVSQISIFAQNSD--PN  70

Query  76   LS-AGFIIKHIETDSGSFIEETDASKCVSLLDEPERGDTIAVKVTKPSKENEKQSKVITI  134
            LS  GF +   E     F +  D       +D   + D +   +T  S +     K+ ++
Sbjct  71   LSNVGFFLCTRELWIHVFQQLEDGE-----IDCALKSDLVKKVLTFASLKTLTGGKIDSV  125

Query  135  DKTRPEGFYTVVFLNCQPGTYVSFDLTLTNYNPG-----PNYLSAGLTALPTLYAMLFVV  189
                    YT++F NC  GT VS D+    YN        +YLSAG T LP +Y +  +
Sbjct  126  YNETDADQYTLLFANCNSGTKVSMDVRSVMYNLDGKGNVRDYLSAGKTVLPRVYFLFSLS  185

Query  190  WTVILGVWLFHFMRGQGKRIFRIHHLVTGIILLKLLTLLFEAIEFHYKKTTGHPGGWVIA  249
            + V+ G+W++   + +   +FRIH  +  +++LK L LL EA +  Y K TG   GW +
Sbjct  186  YFVLAGLWVYVLYKKR-LTVFRIHFFMLAVVILKALNLLCEAEDKSYIKRTGSAHGWDVL  244

Query  250  YYIFSGLKGTMMFVVIALIGTGWAFIKPFLGEKDKNIFLVVIPLQILANIAIIVLEETAP  309
            +YIFS LKG  +F +I LIGTGW+F+KP+L +K+K + ++VIPLQ++ANIA +V++ET P
Sbjct  245  FYIFSFLKGITLFTLIVLIGTGWSFLKPYLQDKEKKVLMIVIPLQVIANIAQVVIDETGP  304

Query  310  ---------EVLRLVDIICCGAILVPIIWSIKHLRDAAAIDGKAKRNMEKLKLFREFYLL  360
                     +V  LVD++CC A+L PI+WSIK+LR+AA  DGKA  N+ KL LFR++Y++
Sbjct  305  FGQDWFTWRQVFLLVDVVCCCAVLFPIVWSIKNLREAAKTDGKAAVNLMKLTLFRQYYIV  364

Query  361  VVTYIYFTRIIVFLLDATLHYQYVWLGEFFTELATLIFWGLTGYKFRPVADNPYLKLDD  419
            V+ YIYFTR++ F L+    Y+Y+W     +ELATL F+  TGYKF+P A NPY  +DD
Sbjct  365  VICYIYFTRVVFFALETITSYRYLWTSVVASELATLAFYVFTGYKFKPEAHNPYFVIDD  423
```

>hypothetical protein B456_007G019200 [Gossypium raimondii]
Sequence ID: KJB39560.1 Length: 433
Range 1: 28 to 432

Score:283 bits(723), Expect:4e-87,
Method:Compositional matrix adjust.,
Identities:159/416(38%), Positives:240/416(57%), Gaps:25/416(6%)

```
Query  35   IKNDRRLAFRIETFGFFTGGVMEMAIENFKVVDDKGSLWDDLSAGFIIKHIETDSGSFIE  94
```

```
            I++D R     + FGF   G +E+   N   +D  G   D    GF +   +T      +
Sbjct  28   IRSDVRPIIPFDEFGFTHNGRLEL---NLSQIDLSGKNLDLNKIGFFLCTRDTWFHVLEQ  84

Query  95   ETDASKCVSLLDEPERGDTIAVKVTKPSKENEKQSKVITIDKTRPEGFYTVVFLNCQPGT  154
              D     +L       D+  VKV    +  + ++ V  +        YT++F NC
Sbjct  85   LNDHHVTCAL-------DSDLVKVVFRFESLKGKTSVNAVFPVNNADQYTLLFANCLTQV  137

Query  155  YVSFDLTLTNYN-----PGPNYLSAGLTALPTLYAMLFVVWTVILGVWLFHFMRGQGKRI  209
             VS  +    YN        +YLSAG T LP +Y +L +V+  + G+W++ F+  +   +
Sbjct  138  KVSMTVRSAMYNLEGKQNSRDYLSAGKTILPRVYFLLSLVYFSLAGIWVY-FLYKKRLTV  196

Query  210  FRIHHLVTGIILLKLLTLLFEAIEFHYKKTTGHPGGWVIAYYIFSGLKGTMMFVVIALIG  269
            FRIH  +  +I+LK   L+FEA +  Y K TG   GW + +YIFS LKG M+F +I LIG
Sbjct  197  FRIHFFMLAVIVLKAFNLVFEAEDKSYIKRTGSAHGWDVLFYIFSFLKGIMLFTLIVLIG  256

Query  270  TGWAFIKPFLGEKDKNIFLVVIPLQILANIAIIVLEETAP---------EVLRLVDIICC  320
            TGW+F+KP+L +K+K + ++VIPLQ++ANIA +V++E +P         ++  LVD+ICC
Sbjct  257  TGWSFLKPYLQDKEKKVLMIVIPLQVVANIAQVVIDEASPFGQDRVTWKQLFLLVDVICC  316

Query  321  GAILVPIIWSIKHLRDAAAIDGKAKRNMEKLKLFREFYLLVVTYIYFTRIIVFLLDATLH  380
             A+L PI+WSIK+LR+AA  DGKA  N+ KL LFR++Y++V+ YIYFTR++V+ L+
Sbjct  317  CAVLFPIVWSIKNLREAARTDGKAAVNLMKLTLFRQYYVVVICYIYFTRVVVYALETITS  376

Query  381  YQYVWLGEFFTELATLIFWGLTGYKFRPVADNPYLKLDDEEEDAEREEAQRQSRTE  436
            Y+Y+W      ELATL F+  TGYKF+P A NPY  +D E+E+A  E+ + +   E
Sbjct  377  YKYLWTAVLAGELATLAFYVFTGYKFKPEAHNPYFAIDGEDEEAAAEQLKLEDEFE  432
```

>PREDICTED: protein GPR107 [Beta vulgaris subsp. vulgaris]
Sequence ID: XP_010684174.1 Length: 440
>hypothetical protein BVRB_7g162510 [Beta vulgaris subsp. vulgaris]
Sequence ID: KMT06178.1 Length: 440
Range 1: 26 to 439

Score:283 bits(723), Expect:6e-87,
Method:Compositional matrix adjust.,
Identities:170/423(40%), Positives:235/423(55%), Gaps:25/423(5%)

```
Query  30   IHKLSIKNDRRLAFRIETFGF-FTGGVMEMAIENFKVVDDKGSLWDDLSAGFIIKHIETD  88
            I  L I +D R     E FGF  TG V               S  D    GF +   E+
Sbjct  26   IKTLKITSDSRPMILFEKFGFTHTGHVSVAVSSVSVTSTSNLSPPDPSRLGFFLLSEESL  85

Query  89   SGSFIE-ETDASKCVSLLDEPERGDTIAVK-VTKPSKENEKQSKVITIDKTRPEGFYTVV  146
                +E + + S CV  LD          + ++ P   +  QS  ITI        Y++
Sbjct  86   LQVLLEIQQNPSFCV--LDSQYTTHLFTFRDLSPPPVSSFNQSYPITIPNE-----YSLF  138

Query  147  FLNCQPGTYVSFDLTLTNYNPGPN----YLSAGLTALPTLYAMLFVVWTVILGVWLFHFM  202
            F NC P + VS  +T   YN   N    YL AG T LPTLY    V++ V L  W+   +
Sbjct  139  FANCNPQSSVSMTVTTLMYNLEFNGVKDYLPAGQTQLPTLYFFFSVLYCVFLAYWIAACV  198

Query  203  RGQGKRIFRIHHLVTGIILLKLLTLLFEAIEFHYKKTTGHPGGWVIAYYIFSGLKGTMMF  262
            R + K + RIH L+ G++++K + L+  A + HY K TG P GW + +Y+F  L+  ++F
Sbjct  199  RNK-KSVHRIHLLMVGLLVIKAVNLICAAEDKHYVKVTGSPHGWDVLFYLFQFLRVVLLF  257
```

```
Query  263  VVIALIGTGWAFIKPFLGEKDKNIFLVVIPLQILANIAIIVLEETAP---------EVLR  313
             VI L+GTGW+F+KPFL EK+K + ++VIPLQ+LANIA IV+ ET P         +V
Sbjct  258  TVIVLVGTGWSFLKPFLQEKEKKVLMIVIPLQVLANIASIVIGETGPFIKDWVTWNQVFL  317

Query  314  LVDIICCGAILVPIIWSIKHLRDAAAIDGKAKRNMEKLKLFREFYLLVVTYIYFTRIIVF  373
            LVDIICC AI+ PI+WSI+ LR+ +  DGKA RN+ KL LFR+FY++V+ Y+YFTRI+VF
Sbjct  318  LVDIICCCAIIFPIVWSIRSLRETSKTDGKAARNLAKLTLFRQFYIVVIGYLYFTRIVVF  377

Query  374  LLDATLHYQYVWLGEFFTELATLIFWGLTGYKFRPVADNPYLKLDDEEEDAEREEAQRQS  433
             L     Y+Y W+     E+A+L+F+ +  Y FRPV  N Y  LDD+EE+A  E A R
Sbjct  378  ALKTISAYKYQWVSNAAEEIASLVFYAIMFYMFRPVERNEYFALDDDEEEAA-EMALRDE  436

Query  434  RTE  436
              E
Sbjct  437  EFE  439
```

```
>hypothetical protein F3Y22_tig00113156pilonHSYRG00094 [Hibiscus syriacus]
Sequence ID: KAE8662714.1 Length: 437
Range 1: 9 to 419

Score:282 bits(722), Expect:6e-87,
Method:Compositional matrix adjust.,
Identities:157/420(37%), Positives:241/420(57%), Gaps:23/420(5%)

Query  14   LIFCLVAFLAPTANGLIHKLSIKNDRRLAFRIETFGFFTGGVMEMAIENFKVVDDKGSLW  73
            L+F L++      +  I    I++D      ++ FGF   G +E+ +    +  +K
Sbjct  9    LLFLLISLFVSHVSSEIRITEIRSDAHPIIPLDQFGFTHTGRLELNVSRIDL-SNKNPDI  67

Query  74   DDLSAGFIIKHIETDSGSFIEETDASKCVSLLDEPERGDTIAVKVTKPSKENEKQSKVIT  133
            D    GF +  ++       +  D+    +L  +      +  +V+       K S  ++
Sbjct  68   DLAKVGFFLSTLDAWMHVLQQLADSEVTCALESD------LVKRVSDFKAAKGKSSFDVS  121

Query  134  IDKTRPEGFYTVVFLNCQPGTYVSFDLTLTNYN-----PGPNYLSAGLTALPTLYAMLFV  188
            +  +  E  YT+VF NC     VS  +    YN        +YLSAG T LP +Y +L +
Sbjct  122  VPVSDAEQ-YTLVFANCLSDAKVSMTVRSAMYNLEGRDGHRDYLSAGKTILPRVYFLLSL  180

Query  189  VWTVILGVWLFHFMRGQGKRIFRIHHLVTGIILLKLLTLLFEAIEFHYKKTTGHPGGWVI  248
            V+  + G+W++   + +   +FRIH  +  +++LK   L+FEA +  Y K TG   GW +
Sbjct  181  VYFSLAGIWIYVLYKKR-LTVFRIHFFMLAVVVLKAFNLIFEAEDKSYIKRTGSAHGWDV  239

Query  249  AYYIFSGLKGTMMFVVIALIGTGWAFIKPFLGEKDKNIFLVVIPLQILANIAIIVLEETA  308
             +YIFS LKG M+F +I LIGTGW+F+KPFL +K+K + ++VIPLQ++ANIA +V++ET+
Sbjct  240  LFYIFSFLKGIMLFTLIVLIGTGWSFVKPFLQDKEKKVLMIVIPLQVVANIAQVVVDETS  299

Query  309  P---------EVLRLVDIICCGAILVPIIWSIKHLRDAAAIDGKAKRNMEKLKLFREFYL  359
            P         +VL LVD+ICC A+L PI+WSIK+LR+AA  DGKA  N+ KL LFR++Y+
Sbjct  300  PFGENRVTWKQVLLLVDVICCCAVLFPIVWSIKNLREAARTDGKAAVNLMKLTLFRQYYV  359

Query  360  LVVTYIYFTRIIVFLLDATLHYQYVWLGEFFTELATLIFWGLTGYKFRPVADNPYLKLDD  419
            +V+ YIYFTR++V+ L+    Y+Y W      ELATL F+  TGYKF+P A NPY  +DD
Sbjct  360  VVICYIYFTRVVVYALETITSYKYFWTSVLAGELATLAFYVFTGYKFKPEAHNPYFAIDD  419
```

```
>Lung_7-TM_R domain-containing protein [Cephalotus follicularis]
Sequence ID: GAV88895.1 Length: 437
Range 1: 17 to 419

Score:282 bits(722), Expect:6e-87,
Method:Compositional matrix adjust.,
Identities:162/418(39%), Positives:239/418(57%), Gaps:37/418(8%)

Query  24   PTANGLIHKLSIKNDRRLAFRIETFGFFTGGVMEMAIENFKVVDDKGSLWDDLSAGFIIK  83
            P +   I    I++D R     + FGF   G +E+ +    + D       DLS
Sbjct  17   PISYSEIRFSEIRSDDRQIIPFDEFGFTHTGRLELNVSKISLSDPNPDF--DLS------  68

Query  84   HIETDSGSFIEETDA-SKCVSLLDEPERGDTIAVKVTKPS------KENEKQS-KVITID  135
                  G F+   DA  + +  L++ E    +  K+ KP       K N+K S KV+ I+
Sbjct  69   ----KLGFFLCTRDAWMQVLQQLEDREVTCALQSKLVKPVFTFNNLKSNQKDSFKVLYIE  124

Query  136  KTRPEGFYTVVFLNCQPGTYVSFDLTLTNYNPG-----PNYLSAGLTALPTLYAMLFVVW  190
                +  YT+VF NC     VS  +    YN        +YLSAG T LP +Y + F+ +
Sbjct  125  TEADQ--YTLVFANCLSQMKVSMSVKSAMYNLDGKSGRRDYLSAGKTILPRVYFLFFLAY  182

Query  191  TVILGVWLFHFMRGQGKRIFRIHHLVTGIILLKLLTLLFEAIEFHYKKTTGHPGGWVIAY  250
              I  +W+ H +  +   ++RIH  +  ++++K L L+ E+ +  Y K TG   GW + +
Sbjct  183  FAIAAIWI-HVLYKKRLTVYRIHFFMLAVVVMKALNLICESEDKSYIKRTGSAHGWDVLF  241

Query  251  YIFSGLKGTMMFVVIALIGTGWAFIKPFLGEKDKNIFLVVIPLQILANIAIIVLEETAP-  309
            YIFS LKG M+F +I LIGTGW+F+KP+L +K+K + ++VIPLQI+ANIA +V++ET P
Sbjct  242  YIFSFLKGIMLFTLIVLIGTGWSFLKPYLQDKEKKVLMIVIPLQIVANIAQVVIDETGPY  301

Query  310  --------EVLRLVDIICCGAILVPIIWSIKHLRDAAAIDGKAKRNMEKLKLFREFYLLV  361
                    +V  LVD++CC A+L PI+WSIK+LR+AA  DGKA  N+ KL LFR++Y++V
Sbjct  302  GQDLITWKQVFLLVDVVCCCAVLFPIVWSIKNLREAARTDGKAAVNLMKLTLFRQYYIVV  361

Query  362  VTYIYFTRIIVFLLDATLHYQYVWLGEFFTELATLIFWGLTGYKFRPVADNPYLKLDD  419
            + YIYFTR++V+ L+    Y+Y+W      ELATL F+  TGYKFRP A NPY  +DD
Sbjct  362  ICYIYFTRVVVYALETITSYKYLWTSLVAGELATLAFYVFTGYKFRPEAHNPYFVIDD  419
```

```
>protein GPR107 [Carex littledalei]
Sequence ID: KAF3332350.1 Length: 448
Range 1: 18 to 431

Score:283 bits(723), Expect:7e-87,
Method:Compositional matrix adjust.,
Identities:160/417(38%), Positives:239/417(57%), Gaps:23/417(5%)

Query  24   PTANGLIHKLSIKNDRRLAFRIETFGFFTGGVMEMAIENFKVVDDKGSLWDDLSA-GFII  82
            P A   I  + +K+D R     + FGF   GV+E+ + +  ++D    L  DLS  GF +
Sbjct  18   PIALAEIRTMQVKSDARSIIPFDEFGFTHTGVLELNVSSISLLDPSPDL--DLSQLGFFL  75

Query  83   KHIETDSGSFIEETDASKCVSLLDEPERGDTIAVKVTKPSKE----NEKQSKVITIDKTR  138
```

```
                ++       +  D     +L  +  +      K+  P+      N + S      K
Sbjct  76   STLDAWVHVLRQLQDLDITCALQSDLVKLVYTFDKLHPPTNPSGVVNARSSSFSAAYKVS  135

Query  139  PEGFYTVVFLNCQPGTYVSFDLTLTNYNPGPN------YLSAGLTALPTLYAMLFVVWTV  192
              G YT+VF NC PG  VS  +    YN  P       YLSAG   LP +Y +LF+V+
Sbjct  136  DPGQYTLVFANCLPGLKVSMSVNSAMYNLDPRSPDKRIYLSAGSAVLPFIYFVLFLVYVG  195

Query  193  ILGVWLFHFMRGQGKRIFRIHHLVTGIILLKLLTLLFEAIEFHYKKTTGHPGGWVIAYYI  252
            +  +WL   +R +   ++RIH+ +  +++LK L L+ EA +    + TG   GW + +YI
Sbjct  196  LGALWLAILLRKRTA-VYRIHYFMLAVVMLKALNLVAEAEDKSCIERTGTAHGWDVLFYI  254

Query  253  FSGLKGTMMFVVIALIGTGWAFIKPFLGEKDKNIFLVVIPLQILANIAIIVLEETAP---  309
            FS LKG  +F +I LIGTGW+F+KPFL +++K + +VVIPLQI+ANIA +V++E+ P
Sbjct  255  FSFLKGISLFTLIVLIGTGWSFLKPFLQDREKKVLMVVIPLQIVANIAQVVIDESGPFAS  314

Query  310  ------EVLRLVDIICCGAILVPIIWSIKHLRDAAAIDGKAKRNMEKLKLFREFYLLVVT  363
                  +V  LVD+ICC A+L PI+WSIK+LR+AA  DGKA  N+ KL LFR++Y++V+
Sbjct  315  DWLTWKQVFLLVDVICCCAVLFPIVWSIKNLREAARSDGKAAVNLMKLTLFRQYYVVVIC  374

Query  364  YIYFTRIIVFLLDATLHYQYVWLGEFFTELATLIFWGLTGYKFRPVADNPYLKLDDE  420
            YIYFTR++V+ L     Y+Y+W      ELATL F+  TGY+FRP + NPY  ++D+
Sbjct  375  YIYFTRVVVYALVTITSYRYLWTSVVAGELATLGFYVFTGYRFRPESHNPYFVVNDD  431
```

>PREDICTED: protein GPR107 [Cucumis melo]
Sequence ID: XP_008448554.1 Length: 439
>seven transmembrane receptor [Cucumis melo subsp. melo]
Sequence ID: ADN33938.1 Length: 439 >protein GPR107 [Cucumis melo var. makuwa]
Sequence ID: KAA0052964.1 Length: 439 >protein GPR107 [Cucumis melo var. makuwa]
Sequence ID: TYK11420.1 Length: 439
Range 1: 11 to 421

Score:282 bits(721), Expect:8e-87,
Method:Compositional matrix adjust.,
Identities:166/431(39%), Positives:243/431(56%), Gaps:46/431(10%)

```
Query  15   IFCLVAFLAPTAN-GLIHKLSIKNDRRLAFRIETFGFFTGGVMEMAIENFKVVDDKGSLW  73
            +F L+ FL P ++   I    I+ND R     + FGF  GG +E+ + +  + D    L
Sbjct  11   MFILIIFLLPFSSFAEIRFTEIRNDNRPIIPFDVFGFSHGGRLELNVTHLTLSDSNPDL-  69

Query  74   DDLS-AGF----------IIKHIETDSGSFIEETDASKCVSLLDEPERGDTIAVKVTKPS  122
             DLS  GF          +I+ +E    S   ++D  K V   D  ++ D   V  ++
Sbjct  70   -DLSKVGFFLCTRESWLHVIQQLEEGDISCALQSDLVKPVYTFDSLKKQDRFGVLYSETD  128

Query  123  KENEKQSKVITIDKTRPEGFYTVVFLNCQPGTYVSFDLTLTNYN-PGPN----YLSAGLT  177
             +                  YT+VF NC     VS D+    YN  G N    YLSAG T
Sbjct  129  ADQ-----------------YTLVFANCLQQFKVSMDVQSAMYNLEGKNARRDYLSAGKT  171

Query  178  ALPTLYAMLFVVWTVILGVWLFHFMRGQGKRIFRIHHLVTGIILLKLLTLLFEAIEFHYK  237
             LP +Y +  +++  +  VW+ H +  +   ++ IH  +  +++LK L LL EA +  Y
Sbjct  172  ILPRIYFVFSLIYFSLAVVWI-HVLYKKRLTVYGIHFFMLAVVILKALNLLCEAEDKSYI  230

Query  238  KTTGHPGGWVIAYYIFSGLKGTMMFVVIALIGTGWAFIKPFLGEKDKNIFLVVIPLQILA  297
```

```
            K TG   GW + +YIFS LKG  +F +I LIGTGW+F+KP+L +K+K + ++VIPLQ++A
Sbjct  231  KRTGSAHGWDVLFYIFSFLKGITLFTLIVLIGTGWSFLKPYLQDKEKKVLMIVIPLQVVA  290

Query  298  NIAIIVLEETAP---------EVLRLVDIICCGAILVPIIWSIKHLRDAAAIDGKAKRNM  348
            NIA +V++ET P         +V  LVD+ICC A+L PI+WSIK+LR+AA  DGKA  N+
Sbjct  291  NIAQVVIDETGPFEQEWVTWKQVFLLVDVICCCAVLFPIVWSIKNLREAARTDGKAAVNL  350

Query  349  EKLKLFREFYLLVVTYIYFTRIIVFLLDATLHYQYVWLGEFFTELATLIFWGLTGYKFRP  408
             KL LFR++Y++V+ YIYFTR++V+ L+    Y+Y+W      ELATL F+  TGYKF+P
Sbjct  351  MKLTLFRQYYIVVICYIYFTRVVVYALETITSYRYLWTSVVAGELATLAFYVFTGYKFKP  410

Query  409  VADNPYLKLDD  419
             A NPY  +DD
Sbjct  411  EAHNPYFVVDD  421
```

>protein GPR107-like [Physcomitrium patens]
Sequence ID: XP_024394221.1 Length: 432
>hypothetical protein PHYPA_018463 [Physcomitrium patens]
Sequence ID: PNR41060.1 Length: 432
Range 1: 7 to 412

Score:281 bits(720), Expect:1e-86,
Method:Compositional matrix adjust.,
Identities:169/421(40%), Positives:240/421(57%), Gaps:27/421(6%)

```
Query  9    LMGCFLIFCLVAFLAPTANGLIHKLSIKNDRRLAFRIETFGFFTGGVMEMAIENFKVVDD  68
            L+   ++  L   L P A   I  LSIKND R     ETFGF  GG + + +     V +
Sbjct  7    LLRALVVMVLAVALFPVAFAEIKHLSIKNDARPIIPFETFGFSIGGEVSVIVSE-VEVKE  65

Query  69   KGSLWDDLSAGFIIKHIETDSGSFIEETDASKCVSLLDEPERGDTIAVKVTKPSKENEKQ  128
             G+  D    GF +  ++ D    I + +A     LLD          +       E +K
Sbjct  66   PGA--DRSRMGFFLSTMD-DLVPVIAQLEAQGKNCLLDSASMHTLFTLA------ETDKF  116

Query  129  SKVITIDKTRPEGFYTVVFLNCQPGTYVSFDLTLTNYNPGP---NYLSAGLTALPTLYAM  185
            +    + ++      Y++ F NC   T VS ++  + YN      ++L  G T LP +Y +
Sbjct  117  NYTHQVFQSNE---YSLFFENCVK-TEVSMEVVTSMYNVENGFRDFLPVGQTVLPRMYFL  172

Query  186  LFVVWTVILGVWLFHFMRGQGKRIFRIHHLVTGIILLKLLTLLFEAIEFHYKKTTGHPGG  245
             F++   +LG+W++     Q + + RIH L+  +++LK L ++ EA +  Y K TG P G
Sbjct  173  FFLINVALLGIWIY-VCWAQKESVHRIHLLMGLLVVLKALNMICEAEDKQYVKRTGEPHG  231

Query  246  WVIAYYIFSGLKGTMMFVVIALIGTGWAFIKPFLGEKDKNIFLVVIPLQILANIAIIVLE  305
            W +AYYIF+ L+G ++F VI LIGTGW+F+KPFL +K+K + +V+IPLQ+ AN A IV +
Sbjct  232  WDVAYYIFNFLRGLLLFSVIVLIGTGWSFLKPFLQDKEKKVLMVIIPLQVFANTAYIVYD  291

Query  306  ETAP---------EVLRLVDIICCGAILVPIIWSIKHLRDAAAIDGKAKRNMEKLKLFRE  356
            E  P         +V  L+DIICC A+L PI+WSIKHLR+AA  DGKA RN+ KL LFR+
Sbjct  292  EAGPSTKDFFTWKQVFLLLDIICCCAVLFPIVWSIKHLREAARTDGKAARNLAKLTLFRQ  351

Query  357  FYLLVVTYIYFTRIIVFLLDATLHYQYVWLGEFFTELATLIFWGLTGYKFRPVADNPYLK  416
            FY++VV+YIYFTRI+VF L     Y+Y W  +F  E A L F+  TGYKFRPV  NPY
Sbjct  352  FYIVVVSYIYFTRIVVFALFTVTVYRYRWTSKFAEEAANLAFYIYTGYKFRPVVHNPYFV  411
```

```
Query  417  L  417
            L
Sbjct  412  L  412
```

>hypothetical protein F3Y22_tig00110785pilonHSYRG00198 [Hibiscus syriacus]
Sequence ID: KAE8694305.1 Length: 433
Range 1: 1 to 415

Score:281 bits(719), Expect:2e-86,
Method:Compositional matrix adjust.,
Identities:159/424(38%), Positives:241/424(56%), Gaps:23/424(5%)

```
Query  10   MGCFLIFCLVAFLAPTANGLIHKLSIKNDRRLAFRIETFGFFTGGVMEMAIENFKVVDDK  69
            M     F L++      +  I    I++D      ++ FGF   G +E+ +    + +
Sbjct  1    MSLRFAFLLISIFVSYVSSEIRLAEIRSDANPIIPLDQFGFTHTGRLELHVSRIDLSNKN  60

Query  70   GSLWDDLSAGFIIKHIETDSGSFIEETDASKCVSLLDEPERGDTIAVKVTKPSKENEKQS  129
              L D    GF +  +  D+   + +  A+  V+   E +      VK+    K  + +S
Sbjct  61   PDL-DLAKVGFFLSTL--DAWMHVLQKLANGEVTCALESD-----LVKLVSDFKAAKGKS  112

Query  130  KVITIDKTRPEGFYTVVFLNCQPGTYVSFDLTLTNYN-----PGPNYLSAGLTALPTLYA  184
             +  +         YT+VF NC     VS ++    YN        +YLSAG T LP +Y
Sbjct  113  SLDVVVPVSNAEQYTLVFANCLSDVKVSMNVRSAMYNLEGRDGHRDYLSAGKTILPRVYF  172

Query  185  MLFVVWTVILGVWLFHFMRGQGKRIFRIHHLVTGIILLKLLTLLFEAIEFHYKKTTGHPG  244
            +L +V+  + G+W++   + Q   +FRIH  +  +++LK   L+FEA +  Y K TG
Sbjct  173  LLSLVYFSLAGLWIYVLYKKQ-LTVFRIHFFMLAVVVLKAFNLIFEAEDKSYIKRTGSAH  231

Query  245  GWVIAYYIFSGLKGTMMFVVIALIGTGWAFIKPFLGEKDKNIFLVVIPLQILANIAIIVL  304
            GW + +YIFS LKG M+F +I LIGTGW+F+KPFL +K+K + ++VIPLQ++ANIA +V+
Sbjct  232  GWDVLFYIFSFLKGIMLFTLIVLIGTGWSFLKPFLQDKEKKVLMIVIPLQVVANIAQVVV  291

Query  305  EETAP---------EVLRLVDIICCGAILVPIIWSIKHLRDAAAIDGKAKRNMEKLKLFR  355
            +ET+P          +V  LVD+ICC A+L PI+WSIK+LR+AA  DGKA  N+ KL LFR
Sbjct  292  DETSPFGEDRVTWKQVFLLVDVICCCAVLFPIVWSIKNLREAARTDGKAAVNLMKLTLFR  351

Query  356  EFYLLVVTYIYFTRIIVFLLDATLHYQYVWLGEFFTELATLIFWGLTGYKFRPVADNPYL  415
            ++Y++V+ YIYFTR+V+ L+    Y+Y W      ELATL F+  TGYKF+P A NPY
Sbjct  352  QYYIVVICYIYFTRVVVYALETITSYKYFWTSVLAGELATLAFYVFTGYKFKPEAHNPYF  411

Query  416  KLDD  419
             +DD
Sbjct  412  AIDD  415
```

>hypothetical protein C1H46_025649 [Malus baccata]
Sequence ID: TQD88760.1 Length: 439
Range 1: 29 to 421

Score:281 bits(719), Expect:2e-86,

Method:Compositional matrix adjust.,
Identities:159/404(39%), Positives:231/404(57%), Gaps:29/404(7%)

```
Query  34   SIKNDRRLAFRIETFGFFTGGVMEMAIENFKVVDDKGSLWDDLSAGFIIKHIETDSGSFI  93
            SI++DR      + FGF   G +E+ +    +VD      D    GF +        S+I
Sbjct  29   SIRSDRSQIIPFDEFGFTHKGRLELNVSKIALVDLPSGDPDLSKVGFFL----CTRDSWI  84

Query  94   EETDASKCVSLLDEPERGDTIAVKVTKPS---KENEKQSKVITIDKTRPEGFYTVVFLNC  150
                    +  L+E E G  +   + KP    K      +  T+        YT++F NC
Sbjct  85   H------VLQQLEEGEIGCALDSTLVKPVYTFKSLNGVDQFSTVFSETDADQYTLLFANC  138

Query  151  QPGTYVSFDLTLTNYN------PGPNYLSAGLTALPTLYAMLFVVWTVILGVWLFHFMRG  204
                 VS D+    YN         +YLSAG T LP +Y +L +V+  + G+W+F   +
Sbjct  139  LQALKVSMDVKSAAYNLEGKTNDRRDYLSAGKTILPRVYFLLSLVYASLAGLWVFVLYKK  198

Query  205  QGKRIFRIHHLVTGIILLKLLTLLFEAIEFHYKKTTGHPGGWVIAYYIFSGLKGTMMFVV  264
            +   +FRIH  +  +++LK   LL EA +  Y K TG   GW + +YIFS LKG  +F +
Sbjct  199  R-LTVFRIHFFMLAVVILKAFNLLCEAEDKSYIKRTGSAHGWDVLFYIFSFLKGVTLFTL  257

Query  265  IALIGTGWAFIKPFLGEKDKNIFLVVIPLQILANIAIIVLEETAP---------EVLRLV  315
            I LIGTGW+F+KPFL +K+K + ++VIPLQ++ANIA +V++ET+P         +V  LV
Sbjct  258  IVLIGTGWSFLKPFLQDKEKKVLMIVIPLQVIANIAQVVIDETSPFGQDWVTWKQVFLLV  317

Query  316  DIICCGAILVPIIWSIKHLRDAAAIDGKAKRNMEKLKLFREFYLLVVTYIYFTRIIVFLL  375
            D+ICC A+L PI+WSIK+LR+AA  DGKA  N+ KL LFR++Y++V+ YIYFTR++V+ L
Sbjct  318  DVICCCAVLFPIVWSIKNLREAARTDGKAAVNLMKLTLFRQYYIVVICYIYFTRVVVYAL  377

Query  376  DATLHYQYVWLGEFFTELATLIFWGLTGYKFRPVADNPYLKLDD  419
            +    Y+Y+W      ELATL F+  TGYKF+P A NPY  +DD
Sbjct  378  ETITSYRYLWTSVVAAELATLAFYVFTGYKFKPEAHNPYFVIDD  421
```

>hypothetical protein E2562_016196, partial [Oryza meyeriana var. granulata]
Sequence ID: KAF0902345.1 Length: 480
Range 1: 25 to 432

Score:282 bits(721), Expect:3e-86,
Method:Compositional matrix adjust.,
Identities:158/409(39%), Positives:232/409(56%), Gaps:20/409(4%)

```
Query  30   IHKLSIKNDRRLAFRIETFGFFTGGVMEMAIENFKVVDDKGSLWDDLSAGFIIKHIETDS  89
            I +  I++D R    ++ FGF   GV+E+ +          S  D    GF +  ++
Sbjct  25   IRETVIRSDPRSIIPLDEFGFSHSGVLELNVSGIAFDPPASSELDLSQLGFFLSTLDAWV  84

Query  90   GSFIEETDASKCVSLLDEPERGDTIAVKVTKPSK----ENEKQSKVITIDKTRPEGFYTV  145
                +  D     +L  +  +      ++  PS     E  + S   T       G YT+
Sbjct  85   HVLRQLQDLDVTCALQADLVKLAYSFDRLRPPSNPAGVEVARSSSFYTAFPVSEPGQYTL  144

Query  146  VFLNC-QPGTYVSFDLTLTNYNPGP-----NYLSAGLTALPTLYAMLFVVWTVILGVWLF  199
            VF NC   G  VS D+    YN  P     +YLSAG TALP+++    VV+  +   W+
Sbjct  145  VFANCLGGGLKVSMDVRSAMYNVDPPTGERSYLSAGATALPSIFGFFGVVYAALAAGWVA  204

Query  200  HFMRGQGKRIFRIHHLVTGIILLKLLTLLFEAIEFHYKKTTGHPGGWVIAYYIFSGLKGT  259
```

```
              +R +   +FRIH+ +  +++LK + LL EA +  Y + TG   GW + +YIFS LKG
Sbjct  205  ILLRKRAA-VFRIHYFMLAVLVLKAVNLLAEAEDKSYIERTGTAHGWDVLFYIFSFLKGI  263

Query  260  MMFVVIALIGTGWAFIKPFLGEKDKNIFLVVIPLQILANIAIIVLEETAP---------E  310
             +F +I LIGTGW+F+KP+L +++K + +VVIPLQ++ANIA +V++E+ P         +
Sbjct  264  SLFTLIVLIGTGWSFLKPYLADREKKVLMVVIPLQVVANIAQVVIDESGPYARDWVTWKQ  323

Query  311  VLRLVDIICCGAILVPIIWSIKHLRDAAAIDGKAKRNMEKLKLFREFYLLVVTYIYFTRI  370
            VL LVD+ICC A+L PI+WSIK+LR+AA  DGKA  N+ KL LFR++Y++V+ YIYFTR+
Sbjct  324  VLLLVDVICCCAVLFPIVWSIKNLREAARSDGKAAVNLMKLTLFRQYYVVVICYIYFTRV  383

Query  371  IVFLLDATLHYQYVWLGEFFTELATLIFWGLTGYKFRPVADNPYLKLDD  419
            +V+ L     YQY W  +   ELATL F+  TGYKFRP   NPY  +DD
Sbjct  384  VVYALMTITSYQYQWTSDVAKELATLAFYVFTGYKFRPEVHNPYFAIDD  432
```

>beta-carotene isomerase D27 [Hibiscus syriacus]
Sequence ID: KAE8727010.1 Length: 437
Range 1: 8 to 419

Score:280 bits(717), Expect:3e-86,
Method:Compositional matrix adjust.,
Identities:154/421(37%), Positives:237/421(56%), Gaps:23/421(5%)

```
Query  13   FLIFCLVAFLAPTANGLIHKLSIKNDRRLAFRIETFGFFTGGVMEMAIENFKVVDDKGSL  72
            FL+F L++      +  I     ++D R    ++ FGF   G +E+ +    + +    L
Sbjct  8    FLLFMLISLFVSFGSAEIRFTEFRSDGRPIIPLDQFGFTHTGRLELNLSQIGLSNKNPDL  67

Query  73   WDDLSAGFIIKHIETDSGSFIEETDASKCVSLLDEPERGDTIAVKVTKPSKENEKQSKVI  132
             D    GF +  ++       +  D     +L       ++  VK+    +  +  S+
Sbjct  68   -DLAKVGFFLSTLDAWMHVLQQLADGEVTCAL-------ESDLVKLVSDFRAAKGNSRFD  119

Query  133  TIDKTRPEGFYTVVFLNCQPGTYVSFDLTLTNYN-----PGPNYLSAGLTALPTLYAMLF  187
             +        YT+VF NC     VS  +    YN        +YLSAG T LP +Y +L
Sbjct  120  VVFPVNNADQYTLVFANCLSDVKVSMTVRSAMYNLEGKQNSRDYLSAGRTVLPRVYFLLS  179

Query  188  VVWTVILGVWLFHFMRGQGKRIFRIHHLVTGIILLKLLTLLFEAIEFHYKKTTGHPGGWV  247
            +++  + G+W++   + +   +F IH  +  +++LK   L+FEA +  Y K TG   GW
Sbjct  180  LIYFSLAGIWIYVLYKKR-LTVFTIHFFMLAVVVLKAFNLIFEAEDKSYIKRTGSAHGWD  238

Query  248  IAYYIFSGLKGTMMFVVIALIGTGWAFIKPFLGEKDKNIFLVVIPLQILANIAIIVLEET  307
            + +YIFS LKG M+F +I LIGTGW+F+KPFL +K+K + ++VIPLQ++ANIA +V++ET
Sbjct  239  VLFYIFSFLKGIMLFTLIVLIGTGWSFLKPFLQDKEKKVLMIVIPLQVVANIAQVVIDET  298

Query  308  AP---------EVLRLVDIICCGAILVPIIWSIKHLRDAAAIDGKAKRNMEKLKLFREFY  358
            +P         +V  LVD+ICC A+L PI+WSIK+LR+AA  DGKA  N+ KL LFR++Y
Sbjct  299  SPFGQDRVTWKQVFLLVDVICCCAVLFPIVWSIKNLREAARTDGKAAVNLMKLTLFRQYY  358

Query  359  LLVVTYIYFTRIIVFLLDATLHYQYVWLGEFFTELATLIFWGLTGYKFRPVADNPYLKLD  418
            ++V+ YIYFTR++V+ L+    Y+Y W      ELATL F+  TGYKF+P A NPY  +D
Sbjct  359  IVVICYIYFTRVVVYALETITSYKYFWTSVLAGELATLAFYVFTGYKFKPEAHNPYFAID  418

Query  419  D  419
```

```
            D
Sbjct  419  D  419
```

>G protein-coupled seven transmembrane receptor [Chlamydomonas reinhardtii]
Sequence ID: XP_001696905.1 Length: 422
Range 1: 23 to 421

Score:280 bits(716), Expect:3e-86,
Method:Compositional matrix adjust.,
Identities:167/409(41%), Positives:240/409(58%), Gaps:27/409(6%)

```
Query  31   HKLSIKNDRRLAFRIETFGFFTGGVMEMAIENFKVVDDKGSLWDDLSA----GFIIKHIE  86
            H +  K+DR L    E FGF  GG +++ I +  +    GS  +D+S     GF +  +E
Sbjct  23   HSVVDKDDRPLIPLTEAFGFAEGGKLDITIRDIGLYRLHGSQ-EDVSNWENFGFFLSPVE  81

Query  87   TDSGSFIEETDASKCVSLLDEPERGDTIAVKVTKPSKENEKQSKVITIDKTRPEGFYTVV  146
             D+    + TD+SKC+         D   +   K S     +S        +  G + +
Sbjct  82   ADAALEQDLTDSSKCI-------LNDVNNLFTFKDSSVQNLESFTFHF-VVQNGGLFYLY  133

Query  147  FLNCQPGTYVSFDLTLTNYNPG----PNYLSAGLTALPTLYAMLFVVWTVILGVWLFHFM  202
            F NC+  T +SF   +  YN       +YLS G T+L  +Y  +F ++T+    W+ +
Sbjct  134  FANCEVDTPISFSSVIEMYNVDTYGRKDYLSVGDTSLDAVYWAMFALFTLCTVAWVVYCY  193

Query  203  RGQGKRIFRIHHLVTGIILLKLLTLLFEAIEFHYKKTTGHPGGWVIAYYIFSGLKGTMMF  262
            R +   + +IH+L+ G+ + K LTLL +A+   Y + TG   GW IAYY+F+  +G + F
Sbjct  194  RNKS-LVHKIHYLMFGLGVFKALTLLSQALMVFYIERTGSADGWNIAYYVFTFFRGVLFF  252

Query  263  VVIALIGTGWAFIKPFLGEKDKNIFLVVIPLQILANIAIIVLEETAP---------EVLR  313
             VI LIGTGW+++KPFLGEK+  I ++VIPLQ+ ANIAI++ +E +P         +V
Sbjct  253  TVIVLIGTGWSYMKPFLGEKEARIIMIVIPLQVFANIAIVITDEESPSVKDWFTWRDVFH  312

Query  314  LVDIICCGAILVPIIWSIKHLRDAAAIDGKAKRNMEKLKLFREFYLLVVTYIYFTRIIVF  373
            LVDIICC AIL PI+WSIKHLR+A+  DGKA RN+EKL LFR+FY++VV YIY TRI+V+
Sbjct  313  LVDIICCCAILFPIVWSIKHLREASQTDGKAARNLEKLTLFRQFYVMVVVYIYVTRIVVY  372

Query  374  LLDATLHYQYVWLGEFFTELATLIFWGLTGYKFRPVADNPYLKLDDEEE  422
            LL +T+ Y+Y W+     EL TL F+  T  KFRP  +NPYLKL  E E
Sbjct  373  LLRSTMQYEYSWVAAMVEELVTLAFYVWTAVKFRPNNENPYLKLQQEIE  421
```

>unnamed protein product [Digitaria exilis]
Sequence ID: CAB3469365.1 Length: 459
Range 1: 9 to 441

Score:281 bits(719), Expect:3e-86,
Method:Compositional matrix adjust.,
Identities:159/435(37%), Positives:241/435(55%), Gaps:22/435(5%)

```
Query  5    RRQTLMGCFLIFCLVAFLAPTANGLIHKLSIKNDRRLAFRIETFGFFTGGVMEMAIENFK  64
            R +  +   ++F   A LAP     I +  I+ D R    ++ FGF   GV+E+ +
Sbjct  9    RPRAALVLLILFLAAASLAPPTAAEIRETLIRADPRSIIPLDEFGFSHSGVLELNVSGIA  68
```

```
Query  65    VVDDKGSLWDDLSA-GFIIKHIETDSGSFIEETDASKCVSLLDEPERGDTIAVKVTKPSK  123
               D   S   DLS  GF +  ++       +  D     +L  +  +      ++  P+
Sbjct  69    F-DPPASAELDLSQLGFFLSTLDAWVHVLRQLQDLDVTCALQSDLVKLAFTFDRLRPPAN  127

Query  124   ----ENEKQSKVITIDKTRPEGFYTVVFLNC-QPGTYVSFDLTLTNYNPGP-----NYLS  173
                 E  + S   T       G YT+VF NC   G  V  D+    YN   P      YLS
Sbjct  128   PAGVEVARSSSFSTAFSVSDPGQYTLVFANCLGGGLKVDMDVRSAMYNVDPVTRERQYLS  187

Query  174   AGLTALPTLYAMLFVVWTVILGVWLFHFMRGQGKRIFRIHHLVTGIILLKLLTLLFEAIE  233
             AG ++LPT Y +  + +  +   W+   +R +   +FRIH+ +  +++LK L LL EA +
Sbjct  188   AGASSLPTFYFLFCLAYAGLAAAWVAILLRKRAA-VFRIHYFMLAVLVLKALNLLAEAED  246

Query  234   FHYKKTTGHPGGWVIAYYIFSGLKGTMMFVVIALIGTGWAFIKPFLGEKDKNIFLVVIPL  293
               Y + TG   GW + +YIFS LKG  +F +I LIGTGW+F+KP+L +++K + +VVIPL
Sbjct  247   KSYIERTGTAHGWDVLFYIFSFLKGISLFTLIVLIGTGWSFLKPYLADREKKVLMVVIPL  306

Query  294   QILANIAIIVLEETAP---------EVLRLVDIICCGAILVPIIWSIKHLRDAAAIDGKA  344
             Q++ANIA +V++E+ P         ++  LVD++CC A+L PI+WSIK+LR+AA  DGKA
Sbjct  307   QVVANIAQVVIDESGPYARDWVTWKQIFLLVDVVCCCAVLFPIVWSIKNLREAARSDGKA  366

Query  345   KRNMEKLKLFREFYLLVVTYIYFTRIIVFLLDATLHYQYVWLGEFFTELATLIFWGLTGY  404
               N+ KL LFR++Y++V+ YIYFTR++V+ L     Y+Y+W     +ELATL F+  TGY
Sbjct  367   AVNLMKLTLFRQYYVVVICYIYFTRVVVYALQTITSYRYLWTSVVASELATLAFYVFTGY  426

Query  405   KFRPVADNPYLKLDD  419
             +FRP   NPY  +DD
Sbjct  427   RFRPEVHNPYFAIDD  441
```

>protein GPR107 [Momordica charantia]
Sequence ID: XP_022145456.1 Length: 440
Range 1: 33 to 422

Score:280 bits(717), Expect:3e-86,
Method:Compositional matrix adjust.,
Identities:159/410(39%), Positives:234/410(57%), Gaps:45/410(10%)

```
Query  35    IKNDRRLAFRIETFGFFTGGVMEMAIENFKVVDDKGSLWDDLS-AGF----------IIK  83
             ++ND R     + FGF  GG +E+ + +  + D    L  DLS AGF          +I+
Sbjct  33    VRNDNRQIIPFDVFGFSHGGRLELNVSHISLSDSNPDL--DLSKAGFFLCTRDSWLHVIQ  90

Query  84    HIETDSGSFIEETDASKCVSLLDEPERGDTIAVKVTKPSKENEKQSKVITIDKTRPEGFY  143
              +E    S   ++D  K V   ++    D   V  ++   +                  Y
Sbjct  91    QLEEGEISCALQSDLVKAVYTFNKLAGRDRFDVLYSETDADQ-----------------Y  133

Query  144   TVVFLNCQPGTYVSFDLTLTNYNPGP-----NYLSAGLTALPTLYAMLFVVWTVILGVWL  198
             T+VF NC     VS D+    YN  P     +YLSAG T LP +Y +L +++  +  VW+
Sbjct  134   TLVFANCLLQLQVSMDVRSAMYNLEPKTGRRDYLSAGKTVLPRIYFLLSLIYFTLAVVWI  193

Query  199   FHFMRGQGKRIFRIHHLVTGIILLKLLTLLFEAIEFHYKKTTGHPGGWVIAYYIFSGLKG  258
             +   + +   ++ IH  +  +++LK L LL EA +  Y K TG   GW + +YIFS LKG
Sbjct  194   YVLYKKR-LTVYGIHFFMLAVVVLKALNLLCEAEDKSYIKRTGSAHGWDVLFYIFSFLKG  252
```

```
Query  259  TMMFVVIALIGTGWAFIKPFLGEKDKNIFLVVIPLQILANIAIIVLEETAP---------  309
              +F +I LIGTGW+F+KP+L +K+K + +VVIPLQ++ANIA +V++ET P
Sbjct  253  ITLFTLIVLIGTGWSFLKPYLQDKEKKVLMVVIPLQVVANIAQVVIDETGPFEQDWVTWK  312

Query  310  EVLRLVDIICCGAILVPIIWSIKHLRDAAAIDGKAKRNMEKLKLFREFYLLVVTYIYFTR  369
            +V  LVD+ICC A+L PI+WSIK+LR+AA  DGKA  N+ KL LFR++Y++V+ YIYFTR
Sbjct  313  QVFLLVDVICCCAVLFPIVWSIKNLREAARTDGKAAVNLMKLTLFRQYYIVVICYIYFTR  372

Query  370  IIVFLLDATLHYQYVWLGEFFTELATLIFWGLTGYKFRPVADNPYLKLDD  419
            ++V+ L+    Y+Y+W       ELATL F+  TGYKF+P A NPY  +DD
Sbjct  373  VVVYALETITSYRYLWTSVVAGELATLAFYAFTGYKFKPEAHNPYFVVDD  422
```

>hypothetical protein SO13M23_000006 [Saccharum officinarum]
Sequence ID: AWA44644.1 Length: 462
Range 1: 20 to 444

Score:281 bits(718), Expect:4e-86,
Method:Compositional matrix adjust.,
Identities:161/427(38%), Positives:242/427(56%), Gaps:23/427(5%)

```
Query  14   LIFCLVAFLA-PTANGLIHKLSIKNDRRLAFRIETFGFFTGGVMEMAIENFKVVDDKGSL  72
            L+F +VA ++ P A   I + +I+ D R    ++ FGF   GV+E+ +      D + S
Sbjct  20   LLFLVVAMVSVPPAAAEIRETAIRADPRSIIPLDEFGFSHSGVLELNVSGIAF-DPQASA  78

Query  73   WDDLSA-GFIIKHIETDSGSFIEETDASKCVSLLDEPERGDTIAVKVTKPSK----ENEK  127
              DLS  GF +  ++        +  D     +L  E  +      ++  PS      E  +
Sbjct  79   ELDLSQLGFFLSTLDAWVHVLRQLQDLDVTCALQSELVKLAFSFDRLRPPSNPAGVEVAR  138

Query  128  QSKVITIDKTRPEGFYTVVFLNC-QPGTYVSFDLTLTNYNPGP-----NYLSAGLTALPT  181
             S   T  +    G YT+VF NC   G  V  D+    YN  P      YLSAG +ALP+
Sbjct  139  SSSFSTAFRVSEPGQYTLVFANCLGGGLKVDMDVRSAMYNVDPATGERQYLSAGASALPS  198

Query  182  LYAMLFVVWTVILGVWLFHFMRGQGKRIFRIHHLVTGIILLKLLTLLFEAIEFHYKKTTG  241
             Y +  + +  +   W+   +R +   +FRIH+ +  +++LK L LL EA +    + TG
Sbjct  199  FYFLFCLAYAGLAAAWVAILLRKRAA-VFRIHYFMLAVLVLKALNLLAEAEDKSCIERTG  257

Query  242  HPGGWVIAYYIFSGLKGTMMFVVIALIGTGWAFIKPFLGEKDKNIFLVVIPLQILANIAI  301
               GW + +YIFS LKG  +F +I LIGTGW+F+KP+L +++K + +VVIPLQ++ANIA
Sbjct  258  TAHGWDVLFYIFSFLKGISLFTLIVLIGTGWSFLKPYLADREKKVLMVVIPLQVVANIAQ  317

Query  302  IVLEETAP---------EVLRLVDIICCGAILVPIIWSIKHLRDAAAIDGKAKRNMEKLK  352
            +V++E+ P         ++  LVD++CC A+L PI+WSIK+LR+AA  DGKA  N+ KL
Sbjct  318  VVIDESGPYARDWVTWKQIFLLVDVVCCCAVLFPIVWSIKNLREAARSDGKAAVNLMKLT  377

Query  353  LFREFYLLVVTYIYFTRIIVFLLDATLHYQYVWLGEFFTELATLIFWGLTGYKFRPVADN  412
            LFR++Y++V+ YIYFTR++V+ L     Y+Y+W       ELATL F+  TGYKFRP   N
Sbjct  378  LFRQYYVVVICYIYFTRVVVYALQTITSYRYLWTSVVAGELATLAFYVFTGYKFRPEVHN  437

Query  413  PYLKLDD  419
            PY  +DD
Sbjct  438  PYFAIDD  444
```

>unnamed protein product [Digitaria exilis]
Sequence ID: CAB3466863.1 Length: 459
Range 1: 9 to 441

Score:281 bits(718), Expect:5e-86,
Method:Compositional matrix adjust.,
Identities:160/435(37%), Positives:241/435(55%), Gaps:22/435(5%)

```
Query  5    RRQTLMGCFLIFCLVAFLAPTANGLIHKLSIKNDRRLAFRIETFGFFTGGVMEMAIENFK  64
            R    +   ++F   A LAP A   I +  I++D R    ++ FGF   GV+E+ +
Sbjct  9    RPHAALVLLILFLAGASLAPPAAAEIRETLIRSDPRSIIPLDEFGFSHSGVLELNVSGIA  68

Query  65   VVDDKGSLWDDLSA-GFIIKHIETDSGSFIEETDASKCVSLLDEPERGDTIAVKVTKPSK  123
              D   S   DLS  GF +  ++       +  D     +L  +  +      ++  P+
Sbjct  69   F-DPPASAELDLSQLGFFLSTLDAWVHVLRQLQDLDVTCALQSDLVKLAFTFDRLRPPAN  127

Query  124  ----ENEKQSKVITIDKTRPEGFYTVVFLNC-QPGTYVSFDLTLTNYNPGP-----NYLS  173
                E  + S   T       G YT+VF NC   G  V  D+    YN  P      YLS
Sbjct  128  PAGVEVARSSSFSTAFSVSDPGQYTLVFANCLGGGLKVDMDVRSAMYNVDPVTRERQYLS  187

Query  174  AGLTALPTLYAMLFVVWTVILGVWLFHFMRGQGKRIFRIHHLVTGIILLKLLTLLFEAIE  233
            AG ++LPT Y +  + +  +   W+   +R +   +FRIH+ +  +++LK L LL EA +
Sbjct  188  AGASSLPTFYFLFCLAYAGLAAAWVAILLRKRAA-VFRIHYFMLAVLVLKALNLLAEAED  246

Query  234  FHYKKTTGHPGGWVIAYYIFSGLKGTMMFVVIALIGTGWAFIKPFLGEKDKNIFLVVIPL  293
              Y + TG   GW + +YIFS LKG  +F +I LIGTGW+F+KP+L +++K + +VVIPL
Sbjct  247  KSYIERTGTAHGWDVLFYIFSFLKGISLFTLIVLIGTGWSFLKPYLADREKKVLMVVIPL  306

Query  294  QILANIAIIVLEETAP---------EVLRLVDIICCGAILVPIIWSIKHLRDAAAIDGKA  344
            Q++ANIA +V++E+ P         ++  LVD++CC A+L PI+WSIK+LR+AA  DGKA
Sbjct  307  QVVANIAQVVIDESGPYARDWVAWKQIFLLVDVVCCCAVLFPIVWSIKNLREAARSDGKA  366

Query  345  KRNMEKLKLFREFYLLVVTYIYFTRIIVFLLDATLHYQYVWLGEFFTELATLIFWGLTGY  404
              N+ KL LFR++Y++V+ YIYFTR++V+ L     Y+Y+W      ELATL F+  TGY
Sbjct  367  AVNLMKLTLFRQYYVVVICYIYFTRVVVYALQTITSYRYLWTSVVAGELATLAFYVFTGY  426

Query  405  KFRPVADNPYLKLDD  419
            +FRP   NPY  +DD
Sbjct  427  RFRPEVHNPYFAIDD  441
```

>protein GPR107-like [Malus domestica]
Sequence ID: XP_008356560.2 Length: 439
Range 1: 29 to 421

Score:280 bits(715), Expect:7e-86,
Method:Compositional matrix adjust.,
Identities:158/404(39%), Positives:231/404(57%), Gaps:29/404(7%)

```
Query  34   SIKNDRRLAFRIETFGFFTGGVMEMAIENFKVVDDKGSLWDDLSAGFIIKHIETDSGSFI  93
```

```
            SI++DRR     + FGF   G +E+ +    + D      D    GF +   ++    F
Sbjct  29   SIRSDRRQIIPFDEFGFTHKGRLELNVSKIALADLPRGDPDLSKVGFFLCTRDSWIHVFQ  88

Query  94   EETDASKCVSLLDEPERGDTIAVKVTKPS---KENEKQSKVITIDKTRPEGFYTVVFLNC  150
            +          L+E E G  +   + KP    K      +  T+        YT++F NC
Sbjct  89   Q----------LEEGEIGCALDSTLVKPVYTFKSLNGVDQFSTVFSETDADQYTLLFANC  138

Query  151  QPGTYVSFDLTLTNYN------PGPNYLSAGLTALPTLYAMLFVVWTVILGVWLFHFMRG  204
                 VS D+    YN         +YLSAG T LP +Y +L +V+  + G+W+F   +
Sbjct  139  LQPLKVSMDVKSAAYNLEGKTNDRRDYLSAGKTILPRVYFLLSLVYASLAGLWVFVLYKK  198

Query  205  QGKRIFRIHHLVTGIILLKLLTLLFEAIEFHYKKTTGHPGGWVIAYYIFSGLKGTMMFVV  264
            +   +FRIH  +  +++LK   LL EA +  Y K TG   GW + +YIFS LKG  +F +
Sbjct  199  R-LTVFRIHFFMLAVVILKAFNLLCEAEDKSYIKRTGSAHGWDVLFYIFSFLKGVTLFTL  257

Query  265  IALIGTGWAFIKPFLGEKDKNIFLVVIPLQILANIAIIVLEETAP---------EVLRLV  315
            I LIGTGW+F+KPFL +K+K + ++VIPLQ++ANIA +V++ET+P         +V  LV
Sbjct  258  IVLIGTGWSFLKPFLQDKEKKVLMIVIPLQVIANIAQVVIDETSPFGQDWVTWKQVFLLV  317

Query  316  DIICCGAILVPIIWSIKHLRDAAAIDGKAKRNMEKLKLFREFYLLVVTYIYFTRIIVFLL  375
            D+ICC A+L PI+WSIK+LR+AA  DGKA  N+ KL LFR++Y++V+ YIYFTR++V+ L
Sbjct  318  DVICCCAVLFPIVWSIKNLREAARTDGKAAVNLMKLTLFRQYYIVVICYIYFTRVVVYAL  377

Query  376  DATLHYQYVWLGEFFTELATLIFWGLTGYKFRPVADNPYLKLDD  419
            +    Y+Y+W      ELATL F+  TGYKF+P A NPY  +DD
Sbjct  378  ETITSYRYLWTSVVAAELATLAFYVFTGYKFKPEAHNPYFVIDD  421
```

>protein GPR107-like [Cucurbita pepo subsp. pepo]
Sequence ID: XP_023551801.1 Length: 435
Range 1: 7 to 417

Score:279 bits(714), Expect:9e-86,
Method:Compositional matrix adjust.,
Identities:164/431(38%), Positives:241/431(55%), Gaps:46/431(10%)

```
Query  15   IFCLVAFLAPTAN-GLIHKLSIKNDRRLAFRIETFGFFTGGVMEMAIENFKVVDDKGSLW  73
            IF L+ FL P ++   I    I+ND R     + FGF  GG +E+ + +  + D    L
Sbjct  7    IFLLLVFLLPISSFAEIRFTDIRNDNRPIIPFDVFGFSHGGRLELNVSHVSLSDTNPDL-  65

Query  74   DDLS-AGF----------IIKHIETDSGSFIEETDASKCVSLLDEPERGDTIAVKVTKPS  122
             DLS AGF          +I+ +E    S   ++D  K V   +  +  D   V  ++
Sbjct  66   -DLSKAGFFLCTRESWLHVIQQLEEAEISCALQSDLVKPVYTFNSLKGQDRFDVLYSQSD  124

Query  123  KENEKQSKVITIDKTRPEGFYTVVFLNCQPGTYVSFDLTLTNYN-----PGPNYLSAGLT  177
             +                  YT+VF NC     VS D+    YN        +YLSAG T
Sbjct  125  ADQ-----------------YTLVFANCLQQLKVSMDVRSAMYNLEGKSGRRDYLSAGKT  167

Query  178  ALPTLYAMLFVVWTVILGVWLFHFMRGQGKRIFRIHHLVTGIILLKLLTLLFEAIEFHYK  237
             LP +Y +  +++ ++  VW+ H +  +   ++ IH  +  +++LK L L+ EA +  Y
Sbjct  168  ILPRIYFIFSMIYFLLAIVWI-HVLYKKRLTVYGIHFFMLAVVILKALNLICEAEDKSYI  226

Query  238  KTTGHPGGWVIAYYIFSGLKGTMMFVVIALIGTGWAFIKPFLGEKDKNIFLVVIPLQILA  297
```

```
            K TG   GW I +YIFS LKG  +F +I LIGTGW+F+KP+L +K+K + ++VIPLQ++A
Sbjct  227  KRTGSAHGWDILFYIFSFLKGITLFTLIVLIGTGWSFLKPYLQDKEKKVLMIVIPLQVVA  286

Query  298  NIAIIVLEETAP---------EVLRLVDIICCGAILVPIIWSIKHLRDAAAIDGKAKRNM  348
            NIA +V +ET P         +V  LVD+ICC A+L PI+WSIK+LR+AA  DGKA  N+
Sbjct  287  NIAQVVTDETGPFEQEWVTWKQVFLLVDVICCCAVLFPIVWSIKNLREAARTDGKAAVNL  346

Query  349  EKLKLFREFYLLVVTYIYFTRIIVFLLDATLHYQYVWLGEFFTELATLIFWGLTGYKFRP  408
             KL LFR++Y++V+ YIYFTR++V+ L+    Y+Y+W      ELAT  F+  TGYKF+P
Sbjct  347  MKLTLFRQYYIVVICYIYFTRVVVYALETITSYRYLWTSVVAGELATFAFYAFTGYKFKP  406

Query  409  VADNPYLKLDD  419
             A NPY  +DD
Sbjct  407  EAHNPYFVVDD  417
```

>hypothetical protein GOBAR_AA25214 [Gossypium barbadense]
Sequence ID: PPR95457.1 Length: 437
>hypothetical protein ES288_A11G019500v1 [Gossypium darwinii]
Sequence ID: TYG92308.1 Length: 437
Range 1: 31 to 419

Score:279 bits(714), Expect:1e-85,
Method:Compositional matrix adjust.,
Identities:155/398(39%), Positives:233/398(58%), Gaps:22/398(5%)

```
Query  35   IKNDRRLAFRIETFGFFTGGVMEMAIENFKVVDDKGSLWDDLSAGFIIKHIETDSGSFIE  94
            I++D R     + FGF   G +E+ +    + D   +L D    GF +   +T      +
Sbjct  31   IRSDVRPIIPFDEFGFTHNGRLELNLSQIDLSDKNKNL-DLNKIGFFLCTRDTWFHVLEQ  89

Query  95   ETDASKCVSLLDEPERGDTIAVKVTKPSKENEKQSKVITIDKTRPEGFYTVVFLNCQPGT  154
              D     +L       D+  VKV    K  + ++ +  +        YT++F NC
Sbjct  90   LNDHHVTCAL-------DSELVKVVFRFKSLDGKTSINPVFPVNNADQYTLLFANCLTQV  142

Query  155  YVSFDLTLTNYN----PGPNYLSAGLTALPTLYAMLFVVWTVILGVWLFHFMRGQGKRIF  210
             VS  +    YN       +YLSAG T LP +Y +L +V+  + G+W++ F+  +   +F
Sbjct  143  KVSMTVRSAMYNIEGKQSRDYLSAGKTILPRVYFLLSLVYFSLAGIWVY-FLYKKRLTVF  201

Query  211  RIHHLVTGIILLKLLTLLFEAIEFHYKKTTGHPGGWVIAYYIFSGLKGTMMFVVIALIGT  270
            RIH  +  +I+LK   L+FEA +  Y K TG   GW + +YIFS LKG M+F +I LIGT
Sbjct  202  RIHFFMLAVIVLKAFNLVFEAEDKSYIKRTGSAHGWDVLFYIFSFLKGIMLFTLIVLIGT  261

Query  271  GWAFIKPFLGEKDKNIFLVVIPLQILANIAIIVLEETAP---------EVLRLVDIICCG  321
            GW+F+KP+L +K+K + ++VIPLQ++ANIA +V++E +P         ++  LVD+ICC
Sbjct  262  GWSFLKPYLQDKEKKVLMIVIPLQVVANIAQVVIDEASPFGQDRATWKQLFLLVDVICCC  321

Query  322  AILVPIIWSIKHLRDAAAIDGKAKRNMEKLKLFREFYLLVVTYIYFTRIIVFLLDATLHY  381
            A+L PI+WSIK+LR+AA  DGKA  N+ KL LFR++Y++V+ YIYFTR++V+ L+    Y
Sbjct  322  AVLFPIVWSIKNLREAARTDGKAAVNLMKLTLFRQYYVVVICYIYFTRVVVYALETITSY  381

Query  382  QYVWLGEFFTELATLIFWGLTGYKFRPVADNPYLKLDD  419
            +Y+W   F  ELATL F+  TGYKF+P A NPY  +DD
Sbjct  382  KYLWTAVFAGELATLAFYVFTGYKFKPEAHNPYFAIDD  419
```

```
>hypothetical protein ES319_A11G019500v1 [Gossypium barbadense]
Sequence ID: KAB2055199.1 Length: 434
Range 1: 28 to 416

Score:279 bits(713), Expect:1e-85,
Method:Compositional matrix adjust.,
Identities:155/398(39%), Positives:233/398(58%), Gaps:22/398(5%)

Query  35   IKNDRRLAFRIETFGFFTGGVMEMAIENFKVVDDKGSLWDDLSAGFIIKHIETDSGSFIE  94
            I++D R     + FGF   G +E+ +    + D   +L D    GF +   +T      +
Sbjct  28   IRSDVRPIIPFDEFGFTHNGRLELNLSQIDLSDKNKNL-DLNKIGFFLCTRDTWFHVLEQ  86

Query  95   ETDASKCVSLLDEPERGDTIAVKVTKPSKENEKQSKVITIDKTRPEGFYTVVFLNCQPGT  154
              D     +L       D+  VKV    K  + ++ +  +        YT++F NC
Sbjct  87   LNDHHVTCAL-------DSELVKVVFRFKSLDGKTSINPVFPVNNADQYTLLFANCLTQV  139

Query  155  YVSFDLTLTNYN----PGPNYLSAGLTALPTLYAMLFVVWTVILGVWLFHFMRGQGKRIF  210
             VS  +    YN       +YLSAG T LP +Y +L +V+  + G+W++ F+  +   +F
Sbjct  140  KVSMTVRSAMYNIEGKQSRDYLSAGKTILPRVYFLLSLVYFSLAGIWVY-FLYKKRLTVF  198

Query  211  RIHHLVTGIILLKLLTLLFEAIEFHYKKTTGHPGGWVIAYYIFSGLKGTMMFVVIALIGT  270
            RIH  +  +I+LK   L+FEA +  Y K TG   GW + +YIFS LKG M+F +I LIGT
Sbjct  199  RIHFFMLAVIVLKAFNLVFEAEDKSYIKRTGSAHGWDVLFYIFSFLKGIMLFTLIVLIGT  258

Query  271  GWAFIKPFLGEKDKNIFLVVIPLQILANIAIIVLEETAP---------EVLRLVDIICCG  321
            GW+F+KP+L +K+K + ++VIPLQ++ANIA +V++E +P         ++  LVD+ICC
Sbjct  259  GWSFLKPYLQDKEKKVLMIVIPLQVVANIAQVVIDEASPFGQDRATWKQLFLLVDVICCC  318

Query  322  AILVPIIWSIKHLRDAAAIDGKAKRNMEKLKLFREFYLLVVTYIYFTRIIVFLLDATLHY  381
            A+L PI+WSIK+LR+AA  DGKA  N+ KL LFR++Y++V+ YIYFTR++V+ L+    Y
Sbjct  319  AVLFPIVWSIKNLREAARTDGKAAVNLMKLTLFRQYYVVVICYIYFTRVVVYALETITSY  378

Query  382  QYVWLGEFFTELATLIFWGLTGYKFRPVADNPYLKLDD  419
            +Y+W   F  ELATL F+  TGYKF+P A NPY  +DD
Sbjct  379  KYLWTAVFAGELATLAFYVFTGYKFKPEAHNPYFAIDD  416
```

```
>protein GPR108 precursor [Zea mays]
Sequence ID: ACG34585.1 Length: 462
>Protein GPR107 [Zea mays]
Sequence ID: PWZ18845.1 Length: 462
Range 1: 22 to 444

Score:280 bits(715), Expect:1e-85,
Method:Compositional matrix adjust.,
Identities:160/427(37%), Positives:240/427(56%), Gaps:24/427(5%)

Query  13   FLIFCLVAFLAPTANGLIHKLSIKNDRRLAFRIETFGFFTGGVMEMAIENFKVVDDKGSL  72
            FL+  +V  L P A   I + +I+ D R    ++ FGF   GV+E+ +      D + S
Sbjct  22   FLVAAMV--LVPPAAAEIRETAIRADPRSIIPLDEFGFSHSGVLELNVSGIAF-DPQASA  78
```

```
Query  73   WDDLSA-GFIIKHIETDSGSFIEETDASKCVSLLDEPERGDTIAVKVTKPSK----ENEK  127
              DLS  GF +  ++       +  D     +L  E  +      ++  PS     E  +
Sbjct  79   ELDLSQLGFFLSTLDAWVHVLRQLQDLDVTCALQSELVKLAFSFDRLRPPSNPAGVEVAR  138

Query  128  QSKVITIDKTRPEGFYTVVFLNCQPGTY-VSFDLTLTNYNPGP-----NYLSAGLTALPT  181
             S   T  +    G YT+VF NC  G   V D+     YN  P      YLSAG +ALP+
Sbjct  139  SSSFSTAFRVSEPGQYTLVFANCLGGGLKVDMDVRSAMYNVDPATGERQYLSAGASALPS  198

Query  182  LYAMLFVVWTVILGVWLFHFMRGQGKRIFRIHHLVTGIILLKLLTLLFEAIEFHYKKTTG  241
             Y +  + +  +   W+   +R +   +FRIH+ +  +++LK L LL EA +    + TG
Sbjct  199  FYFLFCLAYAGLAAAWVAILLRKRAA-VFRIHYFMLAVLVLKALNLLAEAEDKSCIERTG  257

Query  242  HPGGWVIAYYIFSGLKGTMMFVVIALIGTGWAFIKPFLGEKDKNIFLVVIPLQILANIAI  301
               GW + +YIFS LKG  +F +I LIGTGW+F+KP+L +++K + +VVIPLQ++ANIA
Sbjct  258  TAHGWDVLFYIFSFLKGISLFTLIVLIGTGWSFLKPYLADREKKVLMVVIPLQVVANIAQ  317

Query  302  IVLEETAP---------EVLRLVDIICCGAILVPIIWSIKHLRDAAAIDGKAKRNMEKLK  352
            +V++E+ P         ++  LVD++CC A+L PI+WSIK+LR+AA +DGKA  N+ KL
Sbjct  318  VVIDESGPYARDWVTWKQIFLLVDVVCCCAVLFPIVWSIKNLREAARLDGKAAVNLMKLT  377

Query  353  LFREFYLLVVTYIYFTRIIVFLLDATLHYQYVWLGEFFTELATLIFWGLTGYKFRPVADN  412
            LFR++Y++V+ YIYFTR++V+ L     Y+Y+W      EL TL F+  TGYKFRP   N
Sbjct  378  LFRQYYVVVICYIYFTRVVVYALQTITSYRYLWTSVVAGELVTLAFYVFTGYKFRPEVHN  437

Query  413  PYLKLDD  419
            PY  +DD
Sbjct  438  PYFAIDD  444
```

>protein GPR107-like [Cucurbita moschata]
Sequence ID: XP_022923496.1 Length: 435
Range 1: 7 to 417

Score:279 bits(713), Expect:1e-85,
Method:Compositional matrix adjust.,
Identities:164/426(38%), Positives:243/426(57%), Gaps:36/426(8%)

```
Query  15   IFCLVAFLAPTAN-GLIHKLSIKNDRRLAFRIETFGFFTGGVMEMAIENFKVVDDKGSLW  73
            IF L+ FL P ++   I    I+ND R     + FGF  GG +E+ + +  + D    L
Sbjct  7    IFLLLVFLLPISSFAEIRFTDIRNDNRPIIPFDVFGFSHGGRLELNVSHVSLSDTNPDL-  65

Query  74   DDLS-AGFIIKHIETDSGSFIEETDASKCVSLLDEPERGDTIAVKVTKP-----SKENEK  127
             DLS AGF +   E+              +  L+E E    +   + KP     S + +
Sbjct  66   -DLSKAGFFLCTRES----------WLHVIQQLEEAEISCALQSDLVKPVYTFNSLKGQD  114

Query  128  QSKVITIDKTRPEGFYTVVFLNCQPGTYVSFDLTLTNYN-----PGPNYLSAGLTALPTL  182
            +  V+  +    +  YT+VF NC     VS D+    YN        +YLSAG T LP +
Sbjct  115  RFDVLYSESDADQ--YTLVFANCLQQLKVSMDVRSAMYNLEGKSGRRDYLSAGKTILPRI  172

Query  183  YAMLFVVWTVILGVWLFHFMRGQGKRIFRIHHLVTGIILLKLLTLLFEAIEFHYKKTTGH  242
            Y +  +++ ++  VW+ H +  +   ++ IH  +  +++LK L L+ EA +  Y K TG
Sbjct  173  YFIFSMIYFLLAIVWI-HVLYKKRLTVYGIHFFMLAVVILKALNLICEAEDKSYIKRTGS  231
```

```
Query  243   PGGWVIAYYIFSGLKGTMMFVVIALIGTGWAFIKPFLGEKDKNIFLVVIPLQILANIAII  302
               GW I +YIFS LKG  +F +I LIGTGW+F+KP+L +K+K + ++VIPLQ++ANIA +
Sbjct  232   AHGWDILFYIFSFLKGITLFTLIVLIGTGWSFLKPYLQDKEKKVLMIVIPLQVVANIAQV  291

Query  303   VLEETAP---------EVLRLVDIICCGAILVPIIWSIKHLRDAAAIDGKAKRNMEKLKL  353
             V +ET P         +V  LVD+ICC A+L PI+WSIK+LR+AA  DGKA  N+ KL L
Sbjct  292   VTDETGPFEQEWVTWKQVFLLVDVICCCAVLFPIVWSIKNLREAARTDGKAAVNLMKLTL  351

Query  354   FREFYLLVVTYIYFTRIIVFLLDATLHYQYVWLGEFFTELATLIFWGLTGYKFRPVADNP  413
             FR++Y++V+ YIYFTR++V+ L+    Y+Y+W      ELAT  F+  TGYKF+P A NP
Sbjct  352   FRQYYIVVICYIYFTRVVVYALETITSYRYLWTSVVAGELATFAFYAFTGYKFKPEAHNP  411

Query  414   YLKLDD  419
             Y  +DD
Sbjct  412   YFVVDD  417
```

>hypothetical protein [Colocasia esculenta]
Sequence ID: MQL86691.1 Length: 591
Range 1: 173 to 590

Score:283 bits(725), Expect:2e-85,
Method:Compositional matrix adjust.,
Identities:163/427(38%), Positives:237/427(55%), Gaps:48/427(11%)

```
Query  35    IKNDRRLAFRIETFGFFTGGVMEMAIENFKVVDDKGSLWDDLSA-GF----------IIK  83
             I+ D R     + FGF   GV+E+ +      +    L  DLS  GF          +++
Sbjct  173   IRKDARSIIPFDEFGFTHAGVLELNVSKIAFSNPSPDL--DLSQLGFFLSTRDAWIHVVQ  230

Query  84    HIETDSGSFIEETDASKCVSLLDEPERGDTIAVKVTKPSKENEKQSKVITIDKTRPEGFY  143
              I+    +    ++  K V  LD   R  +    +  P+      S  + + +  P+ F
Sbjct  231   QIQDTEITCALHSNLIKTVFTLDHLPRLPSTGAAIPPPA-----DSFGVVVPEGEPDQF-  284

Query  144   TVVFLNCQPGTYVSFDLTLTNYNPGP-----NYLSAGLTALPTLYAMLFVVWTVILGVWL  198
             T+VF NC P   VS D+    YN  P     +YLSAG T LP ++ + F+V+  +  VW+
Sbjct  285   TLVFANCLPQVQVSMDVLSVMYNKDPRTGHRDYLSAGATVLPRIFFLFFLVYAALACVWV  344

Query  199   FHFMRGQGKRIFRIHHLVTGIILLKLLTLLFEAIEFHYKKTTGHPGGWVIAYYIFSGLKG  258
             +   R +    FRIH+ +  +++LK   LL EA +  Y K TG   GW + +YIFS LKG
Sbjct  345   YVLYRKRST-AFRIHYFMLAVVVLKAFNLLCEAEDKSYIKRTGSAHGWDVLFYIFSFLKG  403

Query  259   TMMFVVIALIGTGWAFIKPFLGEKDKNIFLVVIPLQILANIAIIVLEETAP---------  309
               +F +I LIGTGW+F+KP+L +K+K + ++VIPLQ++ANIA +V++E+ P
Sbjct  404   ITLFTLIVLIGTGWSFLKPYLQDKEKRVLMIVIPLQVVANIAQVVIDESGPYARDWITWK  463

Query  310   EVLRLVDIICCGAILVPIIWSIKHLRDAAAIDGKAKRNMEKLKLFREFYLLVVTYIYFTR  369
             +V  LVD+ICC A+L PI+WSIK+LR+AA  DGKA  N+ KL LFR++Y++V+ YIYFTR
Sbjct  464   QVFLLVDVICCCAVLFPIVWSIKNLREAARTDGKAAVNLMKLTLFRQYYIVVICYIYFTR  523

Query  370   IIVFLLDATLHYQYVWLGEFFTELATLIFWGLTGYKFRPVADNPY--------------L  415
             ++V+       Y+Y+W      ELATL F+  TGYKFRP   NPY              L
Sbjct  524   VVVYAFATITSYRYLWTSVVAGELATLAFYVFTGYKFRPEVHNPYFVIDDDEEEAAAEAL  583
```

```
Query  416  KLDDEEE  422
            KLDDE E
Sbjct  584  KLDDEFE  590
```

>protein GPR107-like [Cucurbita maxima]
Sequence ID: XP_022965377.1 Length: 434
Range 1: 6 to 416

Score:278 bits(712), Expect:2e-85,
Method:Compositional matrix adjust.,
Identities:164/426(38%), Positives:243/426(57%), Gaps:36/426(8%)

```
Query  15   IFCLVAFLAPTAN-GLIHKLSIKNDRRLAFRIETFGFFTGGVMEMAIENFKVVDDKGSLW  73
            IF L+ FL P ++   I    I+ND R      + FGF  GG +E+ + +  + D    L
Sbjct  6    IFLLLVFLLPISSFAEIRFTDIRNDNRPIIPFDVFGFSHGGRLELNVSHVSLSDTNPDL-  64

Query  74   DDLS-AGFIIKHIETDSGSFIEETDASKCVSLLDEPERGDTIAVKVTKP-----SKENEK  127
             DLS AGF +   E+                +  L+E E    +   + KP     S + +
Sbjct  65   -DLSKAGFFLCTRES----------WLHVIQQLEEAEISCALQSDLVKPVYTFNSLKGQD  113

Query  128  QSKVITIDKTRPEGFYTVVFLNCQPGTYVSFDLTLTNYN-----PGPNYLSAGLTALPTL  182
            +  V+  +    +  YT+VF NC     VS D+    YN        +YLSAG T LP +
Sbjct  114  RFDVLYSESDADQ--YTLVFANCLQQLKVSMDVRSAMYNLEGKSGRRDYLSAGKTILPRI  171

Query  183  YAMLFVVWTVILGVWLFHFMRGQGKRIFRIHHLVTGIILLKLLTLLFEAIEFHYKKTTGH  242
            Y +  +++ ++  VW+ H +  +   ++ IH  +  +++LK L L+ EA +  Y K TG
Sbjct  172  YFIFSMIYFLLAIVWI-HVLYKKRLTVYGIHFFMLAVVILKALNLICEAEDKSYIKRTGS  230

Query  243  PGGWVIAYYIFSGLKGTMMFVVIALIGTGWAFIKPFLGEKDKNIFLVVIPLQILANIAII  302
              GW I +YIFS LKG  +F +I LIGTGW+F+KP+L +K+K + ++VIPLQ++ANIA +
Sbjct  231  AHGWDILFYIFSFLKGITLFTLIVLIGTGWSFLKPYLQDKEKKVLMIVIPLQVVANIAQV  290

Query  303  VLEETAP---------EVLRLVDIICCGAILVPIIWSIKHLRDAAAIDGKAKRNMEKLKL  353
            V +ET P          +V  LVD+ICC A+L PI+WSIK+LR+AA  DGKA  N+ KL L
Sbjct  291  VTDETGPFEQEWVTWKQVFLLVDVICCCAVLFPIVWSIKNLREAARTDGKAAVNLMKLTL  350

Query  354  FREFYLLVVTYIYFTRIIVFLLDATLHYQYVWLGEFFTELATLIFWGLTGYKFRPVADNP  413
            FR++Y++V+ YIYFTR++V+ L+    Y+Y+W      ELAT  F+  TGYKF+P A NP
Sbjct  351  FRQYYIVVICYIYFTRVVVYALETITSYRYLWTSVVAGELATFAFYAFTGYKFKPEAHNP  410

Query  414  YLKLDD  419
            Y  +DD
Sbjct  411  YFVVDD  416
```

>protein GPR107 [Populus trichocarpa]
Sequence ID: XP_024453523.1 Length: 436
>hypothetical protein POPTR_003G147700 [Populus trichocarpa]
Sequence ID: PNT45656.1 Length: 436 >hypothetical protein POPTR_003G147700 [Populus trichocarpa]

```
Sequence ID: PNT45657.1 Length: 436
Range 1: 27 to 418

Score:278 bits(712), Expect:2e-85,
Method:Compositional matrix adjust.,
Identities:156/407(38%), Positives:240/407(58%), Gaps:36/407(8%)

Query  34   SIKNDRRLAFRIETFGFFTGGVMEMAIENFKVVDDKGSLWDDLSAGFIIKHIETDSGSFI  93
             I++D R      + FGF   G +E+ + N ++ +    L D    GF +     DS   +
Sbjct  27   DIRSDDRQIIPFDEFGFTHFGRLELNVTNIRLSNPNPDL-DRSKIGFFL--CTRDSWLHV  83

Query  94   EETDASKCVSLLDEPERGDTIAVKVTKP-------SKENEKQSKVITIDKTRPEGFYTVV  146
                    ++ L++ E    +   + KP         K+++  SK++T +       YT+V
Sbjct  84   --------INQLEDGEIACALQSDLIKPVFTFNDLKKDHDSLSKIVTQNDADQ---YTLV  132

Query  147  FLNCQPGTYVSFDLTLTNYNPGP-----NYLSAGLTALPTLYAMLFVVWTVILGVWLFHF  201
            F NC     VS D+    YN        +YLSAG T LP +Y +L +++  ++GVW++
Sbjct  133  FANCLTSLKVSMDVKSVMYNLDRGGKVRDYLSAGKTILPRVYYLLSLIYFGLVGVWIYVL  192

Query  202  MRGQGKRIFRIHHLVTGIILLKLLTLLFEAIEFHYKKTTGHPGGWVIAYYIFSGLKGTMM  261
             R +   ++RIH  +  +++LK + LL EA +  Y K TG+  GW + +YIFS LKG  +
Sbjct  193  YRKR-LTVYRIHFFMLAVVILKTVNLLCEAEDKSYIKRTGYAHGWDVLFYIFSFLKGITL  251

Query  262  FVVIALIGTGWAFIKPFLGEKDKNIFLVVIPLQILANIAIIVLEETAP---------EVL  312
            F +I LIGTGW+F+KP+L +K+K + ++VIPLQ++ANIA +V++ET P         +V
Sbjct  252  FTLIVLIGTGWSFLKPYLQDKEKKVLMIVIPLQVVANIAQVVIDETGPYGQDWITWKQVF  311

Query  313  RLVDIICCGAILVPIIWSIKHLRDAAAIDGKAKRNMEKLKLFREFYLLVVTYIYFTRIIV  372
             LVD++CC A+L PI+WSIK+LR+AA  DGKA  N+ KL LFR++Y++V+ YIYFTR++V
Sbjct  312  LLVDVVCCCAVLFPIVWSIKNLREAARTDGKAAVNLMKLTLFRQYYIVVICYIYFTRVVV  371

Query  373  FLLDATLHYQYVWLGEFFTELATLIFWGLTGYKFRPVADNPYLKLDD  419
            + L+    Y+Y+W      ELATL F+  TGYKF+P A NPY  +DD
Sbjct  372  YALETITSYKYLWTSVVAGELATLAFYVFTGYKFKPEAHNPYFVVDD  418



>Lung seven transmembrane receptor family protein [Tripterygium wilfordii]
Sequence ID: KAF5734950.1 Length: 438
Range 1: 13 to 420

Score:278 bits(711), Expect:2e-85,
Method:Compositional matrix adjust.,
Identities:162/422(38%), Positives:239/422(56%), Gaps:28/422(6%)

Query  12   CFLIFCLVAFLAPTANGLIHKLSIKNDRRLAFRIETFGFFTGGVMEMAIENFKVVDDKGS  71
            C LI C V+         I    I++D R     + FGF   G +E+ +    + +
Sbjct  13   CVLILCSVSI------AEIRSTEIRSDERPIIPFDEFGFTHTGRIELNVSQISLSNPNPD  66

Query  72   LWDDLSAGFIIKHIETDSGSFIEETDASKCVSLLDEPERGDTIAVKVTKPSKENEKQSKV  131
            L D    GF +   ++      +  D     +L     + D I    T  S + ++ +
Sbjct  67   L-DRSKLGFFLCTRDSWLHVLQQLEDGEIACAL-----QSDLIKSVYTFNSLKKDQVTFT  120

Query  132  ITIDKTRPEGFYTVVFLNCQPGTYVSFDLTLTNYN-PGPN----YLSAGLTALPTLYAML  186
```

```
                 T  + F T+VF NC     VS D+    YN  G N    YLSAG T LP +Y +L
Sbjct  121  TVYSMTDADQF-TLVFANCLTELKVSMDVKSAMYNLEGKNGRRDYLSAGKTVLPRVYFLL  179

Query  187  FVVWTVILGVWLFHFMRGQGKRIFRIHHLVTGIILLKLLTLLFEAIEFHYKKTTGHPGGW  246
             +V+  + G+W++   + +   +FRIH  +  ++LLK L L+ EA +  Y K TG   GW
Sbjct  180  SLVYCGLAGLWIYVLYKKR-LTVFRIHFFMLAVVLLKALNLICEAEDKSYIKRTGSAHGW  238

Query  247  VIAYYIFSGLKGTMMFVVIALIGTGWAFIKPFLGEKDKNIFLVVIPLQILANIAIIVLEE  306
             + +YIFS LKGT +F +I LIGTGW+F+KP+L +K+K + ++VIPLQ++ANIA +V++E
Sbjct  239  DVLFYIFSFLKGTTLFTLIVLIGTGWSFLKPYLQDKEKKVLMIVIPLQVVANIAQVVIDE  298

Query  307  TAP---------EVLRLVDIICCGAILVPIIWSIKHLRDAAAIDGKAKRNMEKLKLFREF  357
            T P         +V  LVD++CC A+L PI+WSIK+LR+AA  DGKA  N+ KL LFR++
Sbjct  299  TGPFGQDWITWKQVFLLVDVVCCCAVLFPIVWSIKNLREAARTDGKAAVNLMKLTLFRQY  358

Query  358  YLLVVTYIYFTRIIVFLLDATLHYQYVWLGEFFTELATLIFWGLTGYKFRPVADNPYLKL  417
            Y++V+ YIYFTR++V+ L+    Y+Y+W      ELATL F+  TGYKF+P A NPY  +
Sbjct  359  YIVVICYIYFTRVVVYALETITSYKYLWTSVVAGELATLAFYVFTGYKFKPEAHNPYFVI  418

Query  418  DD  419
            DD
Sbjct  419  DD  420


>PREDICTED: protein GPR107 [Populus euphratica]
Sequence ID: XP_011022403.1 Length: 436
Range 1: 27 to 418

Score:278 bits(711), Expect:3e-85,
Method:Compositional matrix adjust.,
Identities:155/410(38%), Positives:239/410(58%), Gaps:42/410(10%)

Query  34   SIKNDRRLAFRIETFGFFTGGVMEMAIENFKVVDDKGSLWDDLSAGF----------IIK  83
             I++D R     + FGF   G +E+ + N ++ +    L D    GF          +I
Sbjct  27   DIRSDDRQIIPFDEFGFTHFGRLELNVTNIRLSNPNPDL-DRSKIGFFLCTRDSWLHVIN  85

Query  84   HIETDSGSFIEETDASKCVSLLDEPERGDTIAVKVTKPSKENEKQSKVITIDKTRPEGFY  143
             +E    +   ++D  K V   ++ ++G             ++  SK++T +       Y
Sbjct  86   QLEDGEITCALQSDLIKPVFTFNDLKKG-------------HDNHSKIVTQNDADQ---Y  129

Query  144  TVVFLNCQPGTYVSFDLTLTNYNPGP-----NYLSAGLTALPTLYAMLFVVWTVILGVWL  198
            T+VF NC     VS D+    YN        +YLSAG T LP +Y +L +++  ++GVW+
Sbjct  130  TLVFANCLTSLKVSMDVKSVMYNLDRGGKVRDYLSAGKTILPRVYYLLSLIYFGLVGVWI  189

Query  199  FHFMRGQGKRIFRIHHLVTGIILLKLLTLLFEAIEFHYKKTTGHPGGWVIAYYIFSGLKG  258
            +   R +   ++RIH  +  +++LK + LL EA +  Y K TG+  GW + +YIFS LKG
Sbjct  190  YVLYRKR-LTVYRIHFFMLAVVILKTVNLLCEAEDKSYIKRTGYAHGWDVLFYIFSFLKG  248

Query  259  TMMFVVIALIGTGWAFIKPFLGEKDKNIFLVVIPLQILANIAIIVLEETAP---------  309
              +F +I LIGTGW+F+KP+L +K+K + ++VIPLQ++ANIA +V++ET P
Sbjct  249  ITLFTLIVLIGTGWSFLKPYLQDKEKKVLMIVIPLQVVANIAQVVIDETGPYGQDWITWK  308

Query  310  EVLRLVDIICCGAILVPIIWSIKHLRDAAAIDGKAKRNMEKLKLFREFYLLVVTYIYFTR  369
```

```
            +V  LVD++CC A+L PI+WSIK+LR+AA  DGKA  N+ KL LFR++Y++V+ YIYFTR
Sbjct  309  QVFLLVDVVCCCAVLFPIVWSIKNLREAARTDGKAAVNLMKLTLFRQYYIVVICYIYFTR  368

Query  370  IIVFLLDATLHYQYVWLGEFFTELATLIFWGLTGYKFRPVADNPYLKLDD  419
            ++V+ L+    Y+Y+W      ELATL F+  TGYKF+P A NPY  +DD
Sbjct  369  VVVYALETITSYKYLWTSVVAGELATLAFYVFTGYKFKPEAHNPYFVVDD  418
```

```
>hypothetical protein EJD97_007360 [Solanum chilense]
Sequence ID: TMW81933.1 Length: 440
Range 1: 20 to 422

Score:278 bits(711), Expect:3e-85,
Method:Compositional matrix adjust.,
Identities:155/422(37%), Positives:237/422(56%), Gaps:46/422(10%)

Query  25   TANGLIHKLSIKNDRRLAFRIETFGFFTGGVMEMAIENFKVVDDKGSLWDDLSAGFIIKH  84
            TA   I  + I++D R     + FG+   G + + + +    + K      LS
Sbjct  20   TAIAEIRSIQIRSDSRSTIPFDEFGYTHFGRLNLTVTDISFSNPKSGPEPVLS-------  72

Query  85   IETDSGSFIEETDASKCVSLLDEPERGDTIAVKVTKPSKENEKQSKVITIDKTRPEGF--  142
               + G F+   +A + V  L++ + GD       + +  ++   +V T D+ +P
Sbjct  73   ---ELGFFLVTREAWQHV--LEQLQDGDI------RCTLHSDLVKRVFTFDQLQPSARQF  121

Query  143  -----------YTVVFLNCQPGTYVSFDLTLTNYNPGP-----NYLSAGLTALPTLYAML  186
                       +T+VF NC P   VS ++    YN  P     ++LSAG TALP +Y +
Sbjct  122  TTSFSVSDANQFTLVFANCMPNLVVSMNVHSVMYNFNPKSGQLDFLSAGKTALPAIYYLF  181

Query  187  FVVWTVILGVWLFHFMRGQGKRIFRIHHLVTGIILLKLLTLLFEAIEFHYKKTTGHPGGW  246
            FVV+ ++   W+F   + +   ++ IH  +  +++LK L LL EA +  Y K TG   GW
Sbjct  182  FVVYVLMGAFWVFALYKKR-LSVYGIHFFMLAVVILKALNLLCEAEDKSYIKKTGTAHGW  240

Query  247  VIAYYIFSGLKGTMMFVVIALIGTGWAFIKPFLGEKDKNIFLVVIPLQILANIAIIVLEE  306
             + +YIFS LKG  +F +I LIGTGW+F+KP+L +K+K + ++VIPLQ++AN+A +V++E
Sbjct  241  DVLFYIFSFLKGITLFTLIVLIGTGWSFLKPYLQDKEKKVLMIVIPLQVVANLAQVVIDE  300

Query  307  TAP---------EVLRLVDIICCGAILVPIIWSIKHLRDAAIDGKAKRNMEKLKLFREF  357
            T P         +V  LVDI+CC A+L PI+WSIK+LR+AA  DGKA  N+ KL LFR++
Sbjct  301  TGPFGENSYTWKQVFLLVDIVCCCAVLFPIVWSIKNLREAAKTDGKAAVNLMKLTLFRQY  360

Query  358  YLLVVTYIYFTRIIVFLLDATLHYQYVWLGEFFTELATLIFWGLTGYKFRPVADNPYLKL  417
            Y++V+ YIYFTR++V+ L+    Y+Y W      E ATL F+  TGY FRP A NPY  +
Sbjct  361  YVIVICYIYFTRVVVYALETITSYRYQWTSVMAAEAATLAFYAFTGYNFRPKAHNPYFAI  420

Query  418  DD  419
            DD
Sbjct  421  DD  422
```

```
>Transmembrane receptor, eukaryota [Corchorus olitorius]
Sequence ID: OMP02814.1 Length: 435
Range 1: 6 to 417
```

Score:278 bits(711), Expect:3e-85,
Method:Compositional matrix adjust.,
Identities:159/422(38%), Positives:237/422(56%), Gaps:24/422(5%)

```
Query  12   CFLIFCLVAFLAPTANGLIHKLSIKNDRRLAFRIETFGFFTGGVMEMAIENFKVVDDKGS  71
            C L   L++  A      I    I++D R     + FGF   G +E+ +   K+ D
Sbjct  6    CLLFCLLISLFASFGFAEIRYTEIRSDNRPIIPFDEFGFTHTGRLELNVSGIKLSDSNPD  65

Query  72   LWDDLSAGFIIKHIETDSGSFIEETDASKCVSLLDEPERGDTIAVKVTKPSKENEKQSKV  131
            L D    GF +   +       E  DA    +L       ++  VKV    K  +  S
Sbjct  66   L-DYSKIGFFLCTRDAWLHVLQELEDAEVTCAL-------NSRLVKVVSDFKSMKGDSFN  117

Query  132  ITIDKTRPEGFYTVVFLNCQPGTYVSFDLTLTNYNPG-----PNYLSAGLTALPTLYAML  186
               ++   +  YT+VF NC     V+ ++    YN        +YLSAG T LP +Y  L
Sbjct  118  ALYEEKDADQ-YTLVFANCLSQIKVTMNVRSAMYNLDGKKNVRDYLSAGKTILPRVYFFL  176

Query  187  FVVWTVILGVWLFHFMRGQGKRIFRIHHLVTGIILLKLLTLLFEAIEFHYKKTTGHPGGW  246
             +V+  + G+W++   + +   +FRIH  +  +++LK   L+ EA +  Y K TG   GW
Sbjct  177  SLVYVTLAGIWIYVLYKKR-LTVFRIHFFMLAVVILKAFNLICEAEDKSYIKRTGSAHGW  235

Query  247  VIAYYIFSGLKGTMMFVVIALIGTGWAFIKPFLGEKDKNIFLVVIPLQILANIAIIVLEE  306
             + +YIFS LKG M+F +I LIGTGW+F+KP+L +K+K + ++VIPLQ++ANIA +V++E
Sbjct  236  DVLFYIFSFLKGIMLFTLIVLIGTGWSFLKPYLQDKEKKVLMIVIPLQVVANIAQVVIDE  295

Query  307  TAP---------EVLRLVDIICCGAILVPIIWSIKHLRDAAAIDGKAKRNMEKLKLFREF  357
            T P         +V  LVD++CC A+L PI+WSIK+LR+AA  DGKA  N+ KL LFR++
Sbjct  296  TGPFGQDWITWKQVFLLVDVVCCCAVLFPIVWSIKNLREAARTDGKAAVNLMKLTLFRQY  355

Query  358  YLLVVTYIYFTRIIVFLLDATLHYQYVWLGEFFTELATLIFWGLTGYKFRPVADNPYLKL  417
            Y++V+ YIYFTR++V+ L+    Y+Y+W      ELATL F+  TGYKF+P A NPY  +
Sbjct  356  YIVVICYIYFTRVVVYALETITSYKYLWTSVVAGELATLAFYVFTGYKFKPEAHNPYFVI  415

Query  418  DD  419
            DD
Sbjct  416  DD  417
```

>hypothetical protein ES319_A02G106800v1 [Gossypium barbadense]
Sequence ID: KAB2093650.1 Length: 437
>hypothetical protein GOBAR_AA20287 [Gossypium barbadense]
Sequence ID: PPS00373.1 Length: 437 >hypothetical protein ES288_A02G117200v1
[Gossypium darwinii]
Sequence ID: TYH28093.1 Length: 437
Range 1: 6 to 419

Score:278 bits(710), Expect:4e-85,
Method:Compositional matrix adjust.,
Identities:158/422(37%), Positives:238/422(56%), Gaps:22/422(5%)

```
Query  12   CFLIFCLVAFLAPTANGLIHKLSIKNDRRLAFRIETFGFFTGGVMEMAIENFKVVDDKGS  71
            C L F LV+         I    I++D R     + FGF   G +E+ + N  + +
Sbjct  6    CCLSFLLVSLFVSFCYAEIRLTEIRHDDRPIIPFDEFGFTHTGRLELNVSNIALSNQNPD  65
```

```
Query  72   LWDDLSAGFIIKHIETDSGSFIEETDASKCVSLLDEPERGDTIAVKVTKPSKENEKQSKV  131
            L D    GF +   +      ++  D       ++ P   D + + V +  +   K +
Sbjct  66   L-DMSKVGFFLCTRDAWMQVLLQIEDG-----YINCPLNSDFVKL-VFEFKQLKGKSNSF  118

Query  132  ITIDKTRPEGFYTVVFLNCQPGTYVSFDLTLTNYN-----PGPNYLSAGLTALPTLYAML  186
             T+ +      YT+VF NC     VS D+    YN        +YLSAG T LP +Y +
Sbjct  119  NTVYQENTADQYTLVFANCLSHVKVSMDVRSAMYNLDGKEKHRDYLSAGKTILPRVYFIF  178

Query  187  FVVWTVILGVWLFHFMRGQGKRIFRIHHLVTGIILLKLLTLLFEAIEFHYKKTTGHPGGW  246
             +V+  + G+W++  +  +   +FRIH  +  +I+LK + LL EA +  Y K TG   GW
Sbjct  179  ALVYFTLAGIWIY-VLYLKRLAVFRIHLFMLAVIILKAVNLLCEAEDKSYIKRTGSAHGW  237

Query  247  VIAYYIFSGLKGTMMFVVIALIGTGWAFIKPFLGEKDKNIFLVVIPLQILANIAIIVLEE  306
             + +YIFS LKG M+F +I LIGTGW+F+KP+L +K+KN+ ++VIPLQ++AN+A IV++E
Sbjct  238  DVLFYIFSFLKGVMLFTLIVLIGTGWSFLKPYLQDKEKNVLMIVIPLQVVANVAQIVIDE  297

Query  307  TAP---------EVLRLVDIICCGAILVPIIWSIKHLRDAAAIDGKAKRNMEKLKLFREF  357
            T P         +V  LVD++CC A+L PI+WSIK+LR+AA  DGKA  N+ KL LFR++
Sbjct  298  TGPFGHDWMVWKQVFLLVDVVCCCAVLFPIVWSIKNLREAAKTDGKAAVNLMKLTLFRQY  357

Query  358  YLLVVTYIYFTRIIVFLLDATLHYQYVWLGEFFTELATLIFWGLTGYKFRPVADNPYLKL  417
            Y++V+ YIYFTR++V+ L     Y+Y+W      E+ATL F+  TGYKF+P   NPY  +
Sbjct  358  YIVVICYIYFTRVVVYALVTITSYKYLWTSMVAGEVATLAFYVFTGYKFKPEPHNPYFVI  417

Query  418  DD  419
            DD
Sbjct  418  DD  419
```

>PREDICTED: protein GPR107-like [Gossypium hirsutum]
Sequence ID: XP_016741571.1 Length: 437
>PREDICTED: protein GPR107-like [Gossypium arboreum]
Sequence ID: XP_017629713.1 Length: 437 >hypothetical protein ES332_A02G119900v1 [Gossypium tomentosum]
Sequence ID: TYI39812.1 Length: 437 >hypothetical protein E1A91_A02G110200v1 [Gossypium mustelinum]
Sequence ID: TYJ46290.1 Length: 437
Range 1: 6 to 419

Score:278 bits(710), Expect:5e-85,
Method:Compositional matrix adjust.,
Identities:158/422(37%), Positives:238/422(56%), Gaps:22/422(5%)

```
Query  12   CFLIFCLVAFLAPTANGLIHKLSIKNDRRLAFRIETFGFFTGGVMEMAIENFKVVDDKGS  71
            C L F LV+         I    I++D R     + FGF   G +E+ + N  + +
Sbjct  6    CCLSFLLVSLFVSFCYAEIRLTEIRHDDRPIIPFDEFGFTHTGRLELNVSNIALSNQNPD  65

Query  72   LWDDLSAGFIIKHIETDSGSFIEETDASKCVSLLDEPERGDTIAVKVTKPSKENEKQSKV  131
            L D    GF +   +      ++  D       ++ P   D + + V +  +   K +
Sbjct  66   L-DMSKVGFFLCTRDAWMQVLLQIEDG-----YINCPLNSDFVKL-VFEFKQLKGKSNSF  118

Query  132  ITIDKTRPEGFYTVVFLNCQPGTYVSFDLTLTNYN-----PGPNYLSAGLTALPTLYAML  186
```

```
              T+ +      YT+VF NC      VS D+     YN        +YLSAG T LP  +Y +
Sbjct  119   NTVYQENTADQYTLVFANCLSHVKVSMDVRSAMYNLDGKEKHRDYLSAGKTILPRVYFIF  178

Query  187   FVVWTVILGVWLFHFMRGQGKRIFRIHHLVTGIILLKLLTLLFEAIEFHYKKTTGHPGGW  246
              +V+  + G+W++  +  +   +FRIH  +  +I+LK + LL EA +  Y K TG   GW
Sbjct  179   ALVYFTLAGIWIY-VLYLKRLTVFRIHLFMLAVIILKAVNLLCEAEDKSYIKRTGSAHGW  237

Query  247   VIAYYIFSGLKGTMMFVVIALIGTGWAFIKPFLGEKDKNIFLVVIPLQILANIAIIVLEE  306
              + +YIFS LKG M+F +I LIGTGW+F+KP+L +K+KN+ ++VIPLQ++AN+A IV++E
Sbjct  238   DVLFYIFSFLKGVMLFTLIVLIGTGWSFLKPYLQDKEKNVLMIVIPLQVVANVAQIVIDE  297

Query  307   TAP---------EVLRLVDIICCGAILVPIIWSIKHLRDAAAIDGKAKRNMEKLKLFREF  357
             T P         +V  LVD++CC A+L PI+WSIK+LR+AA  DGKA  N+ KL LFR++
Sbjct  298   TGPFGHDWMVWKQVFLLVDVVCCCAVLFPIVWSIKNLREAAKTDGKAAVNLMKLTLFRQY  357

Query  358   YLLVVTYIYFTRIIVFLLDATLHYQYVWLGEFFTELATLIFWGLTGYKFRPVADNPYLKL  417
             Y++V+ YIYFTR++V+ L     Y+Y+W      E+ATL F+  TGYKF+P   NPY  +
Sbjct  358   YIVVICYIYFTRVVVYALVTITSYKYLWTSMVAGEVATLAFYVFTGYKFKPEPHNPYFVI  417

Query  418   DD  419
             DD
Sbjct  418   DD  419
```

>hypothetical protein EE612_031732, partial [Oryza sativa]
Sequence ID: KAB8101064.1 Length: 446
Range 1: 21 to 428

Score:278 bits(710), Expect:5e-85,
Method:Compositional matrix adjust.,
Identities:157/409(38%), Positives:230/409(56%), Gaps:20/409(4%)

```
Query  30    IHKLSIKNDRRLAFRIETFGFFTGGVMEMAIENFKVVDDKGSLWDDLSAGFIIKHIETDS  89
             I +  I++D R    ++ FGF   GV+E+ +          S  D    GF +  ++
Sbjct  21    IRETVIRSDPRSIIPLDEFGFSHSGVLELNVSGIAFDPPASSELDLSQLGFFLSTLDAWV  80

Query  90    GSFIEETDASKCVSLLDEPERGDTIAVKVTKPSK----ENEKQSKVITIDKTRPEGFYTV  145
                 +  D     +L  +  +      ++  PS     E  + S   T       G YT+
Sbjct  81    HVLRQLQDLDVTCALQADLVKLAYSFDRLRPPSNPAGVEVARSSSFSTAFPVSEPGQYTL  140

Query  146   VFLNC-QPGTYVSFDLTLTNYNPGP-----NYLSAGLTALPTLYAMLFVVWTVILGVWLF  199
             VF NC   G  VS D+    YN  P     +YLSAG TALPT++    V +  +   W+
Sbjct  141   VFANCLGGGLKVSMDVRSAMYNVDPPTGERSYLSAGATALPTIFGFFGVAYAALAAGWIA  200

Query  200   HFMRGQGKRIFRIHHLVTGIILLKLLTLLFEAIEFHYKKTTGHPGGWVIAYYIFSGLKGT  259
               +R +   +FRIH+ +  +++LK + LL EA +  Y + TG   GW + +YIFS LKG
Sbjct  201   ILLRKRAA-VFRIHYFMLAVLVLKAVNLLAEAEDKSYIERTGTAHGWDVLFYIFSFLKGI  259

Query  260   MMFVVIALIGTGWAFIKPFLGEKDKNIFLVVIPLQILANIAIIVLEETAP---------E  310
              +F +I LIGTGW+F+KP+L +++K + +VVIPLQ++ANIA +V++E+ P         +
Sbjct  260   SLFTLIVLIGTGWSFLKPYLADREKKVLMVVIPLQVVANIAQVVIDESGPYARAWVTWKQ  319

Query  311   VLRLVDIICCGAILVPIIWSIKHLRDAAAIDGKAKRNMEKLKLFREFYLLVVTYIYFTRI  370
```

```
            VL LVD+ICC A+L PI+WSIK+LR+AA  DGKA  N+ KL LFR++Y++V+ YIYFTR+
Sbjct  320  VLLLVDVICCCAVLFPIVWSIKNLREAARSDGKAAVNLMKLTLFRQYYVVVICYIYFTRV  379

Query  371  IVFLLDATLHYQYVWLGEFFTELATLIFWGLTGYKFRPVADNPYLKLDD  419
            +V+ L     Y+Y W      ELATL F+  TGYKFRP   NPY  +DD
Sbjct  380  VVYALMTITSYRYQWTSYVAKELATLAFYVFTGYKFRPEVHNPYFAIDD  428
```

>hypothetical protein EE612_031732 [Oryza sativa]
Sequence ID: KAB8101063.1 Length: 453
Range 1: 28 to 435

Score:278 bits(711), Expect:5e-85,
Method:Compositional matrix adjust.,
Identities:157/409(38%), Positives:230/409(56%), Gaps:20/409(4%)

```
Query  30   IHKLSIKNDRRLAFRIETFGFFTGGVMEMAIENFKVVDDKGSLWDDLSAGFIIKHIETDS  89
            I +  I++D R    ++ FGF   GV+E+ +          S  D    GF +  ++
Sbjct  28   IRETVIRSDPRSIIPLDEFGFSHSGVLELNVSGIAFDPPASSELDLSQLGFFLSTLDAWV  87

Query  90   GSFIEETDASKCVSLLDEPERGDTIAVKVTKPSK----ENEKQSKVITIDKTRPEGFYTV  145
                +  D     +L  +  +      ++  PS     E  + S   T       G YT+
Sbjct  88   HVLRQLQDLDVTCALQADLVKLAYSFDRLRPPSNPAGVEVARSSSFSTAFPVSEPGQYTL  147

Query  146  VFLNC-QPGTYVSFDLTLTNYNPGP-----NYLSAGLTALPTLYAMLFVVWTVILGVWLF  199
            VF NC   G  VS D+    YN  P     +YLSAG TALPT++    V +  +   W+
Sbjct  148  VFANCLGGGLKVSMDVRSAMYNVDPPTGERSYLSAGATALPTIFGFFGVAYAALAAGWIA  207

Query  200  HFMRGQGKRIFRIHHLVTGIILLKLLTLLFEAIEFHYKKTTGHPGGWVIAYYIFSGLKGT  259
              +R +   +FRIH+ +  +++LK + LL EA +  Y + TG   GW + +YIFS LKG
Sbjct  208  ILLRKRAA-VFRIHYFMLAVLVLKAVNLLAEAEDKSYIERTGTAHGWDVLFYIFSFLKGI  266

Query  260  MMFVVIALIGTGWAFIKPFLGEKDKNIFLVVIPLQILANIAIIVLEETAP---------E  310
             +F +I LIGTGW+F+KP+L +++K + +VVIPLQ++ANIA +V++E+ P         +
Sbjct  267  SLFTLIVLIGTGWSFLKPYLADREKKVLMVVIPLQVVANIAQVVIDESGPYARAWVTWKQ  326

Query  311  VLRLVDIICCGAILVPIIWSIKHLRDAAAIDGKAKRNMEKLKLFREFYLLVVTYIYFTRI  370
            VL LVD+ICC A+L PI+WSIK+LR+AA  DGKA  N+ KL LFR++Y++V+ YIYFTR+
Sbjct  327  VLLLVDVICCCAVLFPIVWSIKNLREAARSDGKAAVNLMKLTLFRQYYVVVICYIYFTRV  386

Query  371  IVFLLDATLHYQYVWLGEFFTELATLIFWGLTGYKFRPVADNPYLKLDD  419
            +V+ L     Y+Y W      ELATL F+  TGYKFRP   NPY  +DD
Sbjct  387  VVYALMTITSYRYQWTSYVAKELATLAFYVFTGYKFRPEVHNPYFAIDD  435
```

>protein GPR107 [Oryza sativa Japonica Group]
Sequence ID: XP_015643947.1 Length: 454
>hypothetical protein OsI_21496 [Oryza sativa Indica Group]
Sequence ID: EAY99527.1 Length: 454 >unknown [Oryza sativa Japonica Group]
Sequence ID: AAO33145.1 Length: 454 >hypothetical protein OsJ_20008 [Oryza sativa Japonica Group]
Sequence ID: EAZ35718.1 Length: 454 >hypothetical protein DAI22_06g021100 [Oryza

sativa Japonica Group]
Sequence ID: KAF2925012.1 Length: 454 >putative lung seven transmembrane receptor 1 [Oryza sativa Japonica Group]
Sequence ID: BAA84809.1 Length: 454
Range 1: 29 to 436

Score:278 bits(711), Expect:5e-85,
Method:Compositional matrix adjust.,
Identities:157/409(38%), Positives:230/409(56%), Gaps:20/409(4%)

```
Query  30   IHKLSIKNDRRLAFRIETFGFFTGGVMEMAIENFKVVDDKGSLWDDLSAGFIIKHIETDS  89
            I +  I++D R    ++ FGF   GV+E+ +          S  D    GF +  ++
Sbjct  29   IRETVIRSDPRSIIPLDEFGFSHSGVLELNVSGIAFDPPASSELDLSQLGFFLSTLDAWV  88

Query  90   GSFIEETDASKCVSLLDEPERGDTIAVKVTKPSK----ENEKQSKVITIDKTRPEGFYTV  145
                +  D     +L  +  +      ++  PS     E  + S   T       G YT+
Sbjct  89   HVLRQLQDLDVTCALQADLVKLAYSFDRLRPPSNPAGVEVARSSSFSTAFPVSEPGQYTL  148

Query  146  VFLNC-QPGTYVSFDLTLTNYNPGP-----NYLSAGLTALPTLYAMLFVVWTVILGVWLF  199
            VF NC   G  VS D+    YN  P     +YLSAG TALPT++    V +  +   W+
Sbjct  149  VFANCLGGGLKVSMDVRSAMYNVDPPTGERSYLSAGATALPTIFGFFGVAYAALAAGWIA  208

Query  200  HFMRGQGKRIFRIHHLVTGIILLKLLTLLFEAIEFHYKKTTGHPGGWVIAYYIFSGLKGT  259
              +R +   +FRIH+ +  +++LK + LL EA +  Y + TG   GW + +YIFS LKG
Sbjct  209  ILLRKRAA-VFRIHYFMLAVLVLKAVNLLAEAEDKSYIERTGTAHGWDVLFYIFSFLKGI  267

Query  260  MMFVVIALIGTGWAFIKPFLGEKDKNIFLVVIPLQILANIAIIVLEETAP---------E  310
             +F +I LIGTGW+F+KP+L +++K + +VVIPLQ++ANIA +V++E+ P         +
Sbjct  268  SLFTLIVLIGTGWSFLKPYLADREKKVLMVVIPLQVVANIAQVVIDESGPYARAWVTWKQ  327

Query  311  VLRLVDIICCGAILVPIIWSIKHLRDAAAIDGKAKRNMEKLKLFREFYLLVVTYIYFTRI  370
            VL LVD+ICC A+L PI+WSIK+LR+AA  DGKA  N+ KL LFR++Y++V+ YIYFTR+
Sbjct  328  VLLLVDVICCCAVLFPIVWSIKNLREAARSDGKAAVNLMKLTLFRQYYVVVICYIYFTRV  387

Query  371  IVFLLDATLHYQYVWLGEFFTELATLIFWGLTGYKFRPVADNPYLKLDD  419
            +V+ L     Y+Y W      ELATL F+  TGYKFRP   NPY  +DD
Sbjct  388  VVYALMTITSYRYQWTSYVAKELATLAFYVFTGYKFRPEVHNPYFAIDD  436
```

>hypothetical protein [Gossypium harknessii]
Sequence ID: MBA0819879.1 Length: 437
Range 1: 6 to 419

Score:277 bits(709), Expect:6e-85,
Method:Compositional matrix adjust.,
Identities:160/423(38%), Positives:239/423(56%), Gaps:24/423(5%)

```
Query  12   CFLIFCLVAFLAPTANGLIHKLSIKNDRRLAFRIETFGFFTGGVMEMAIENFKVVDDKGS  71
            CF  F LV+         I    I++D R     + FGF   G +E+ + N  + +
Sbjct  6    CFFSFLLVSLFVSFCYAEIRLTEIRHDDRPIIPFDEFGFTHTGRLELNVSNIALSNQNPD  65

Query  72   LWDDLSAGFIIKHIETDSGSFIEETDASKCVSLLDEPERGDTIAVKVTKPSKENEKQSKV  131
            L D    GF +   +      ++  D     +L       ++  VK+    K+ + +S
```

```
Sbjct  66   L-DMSKVGFFLCTRDAWMQVLLQIEDGYINCAL-------NSDFVKLVFEFKQLKGKSNS  117

Query  132  I-TIDKTRPEGFYTVVFLNCQPGTYVSFDLTLTNYN-----PGPNYLSAGLTALPTLYAM  185
              T+ +      YT+VF NC     VS D+    YN        +YLSAG T LP +Y +
Sbjct  118  FNTVYQENTADQYTLVFANCLSHVKVSMDVRSAMYNLDGKEKHRDYLSAGKTILPRVYFI  177

Query  186  LFVVWTVILGVWLFHFMRGQGKRIFRIHHLVTGIILLKLLTLLFEAIEFHYKKTTGHPGG  245
              +V+  + G+W++  +  +   +FRIH  +  +I+LK + LL EA +  Y K TG   G
Sbjct  178  FALVYFTLAGIWIY-VLYLKRLTVFRIHLFMLAVIILKAVNLLCEAEDKSYIKRTGSAHG  236

Query  246  WVIAYYIFSGLKGTMMFVVIALIGTGWAFIKPFLGEKDKNIFLVVIPLQILANIAIIVLE  305
            W + +YIFS LKG M+F +I LIGTGW+F+KP+L +K+KN+ +VVIPLQ++AN+A IV++
Sbjct  237  WDVLFYIFSFLKGVMLFTLIVLIGTGWSFLKPYLQDKEKNVLMVVIPLQVVANVAQIVID  296

Query  306  ETAP---------EVLRLVDIICCGAILVPIIWSIKHLRDAAAIDGKAKRNMEKLKLFRE  356
            ET P         +V  LVD++CC A+L PI+WSIK+LR+AA  DGKA  N+ KL LFR+
Sbjct  297  ETGPFGHDWMVWKQVFLLVDVVCCCAVLFPIVWSIKNLREAAKTDGKAAVNLMKLTLFRQ  356

Query  357  FYLLVVTYIYFTRIIVFLLDATLHYQYVWLGEFFTELATLIFWGLTGYKFRPVADNPYLK  416
            +Y++V+ YIYFTR++V+ L     Y+Y+W      E+ATL F+  TGYKF+P   NPY
Sbjct  357  YYIVVICYIYFTRVVVYALVTITSYKYLWTSVVAGEVATLAFYVFTGYKFKPEPHNPYFV  416

Query  417  LDD  419
            +DD
Sbjct  417  IDD  419
```

>Lung seven transmembrane receptor-like [Trema orientale]
Sequence ID: PON38231.1 Length: 441
Range 1: 21 to 423

Score:277 bits(709), Expect:7e-85,
Method:Compositional matrix adjust.,
Identities:157/411(38%), Positives:235/411(57%), Gaps:23/411(5%)

```
Query  24   PTANGLIHKLSIKNDRRLAFRIETFGFFTGGVMEMAIENFKVVDDKGSLWDDLS-AGFII  82
            P A   I    I+ND R    ++ FGF   G +E+++    +         +LS  GF +
Sbjct  21   PMAFAEIRFSEIRNDIRPIIPLDEFGFTHKGRLELSVSQISIFAQNSD--PNLSNVGFFL  78

Query  83   KHIETDSGSFIEETDASKCVSLLDEPERGDTIAVKVTKPSKENEKQSKVITIDKTRPEGF  142
               E     F +       V  +D   + D +   +T  S +      + ++
Sbjct  79   CTRELWIHVFQQLE-----VGEVDCALKSDLVKKVLTFASLKTLTGGNIDSVYNETDADQ  133

Query  143  YTVVFLNCQPGTYVSFDLTLTNYNPG-----PNYLSAGLTALPTLYAMLFVVWTVILGVW  197
            YT++F NC  GT VS D+    YN        +YLSAG T LP +Y +  + + V+ G+W
Sbjct  134  YTLLFANCNSGTKVSMDVRSAMYNLDGRGNVRDYLSAGKTVLPRVYFLFSLSYFVLAGLW  193

Query  198  LFHFMRGQGKRIFRIHHLVTGIILLKLLTLLFEAIEFHYKKTTGHPGGWVIAYYIFSGLK  257
            ++   + +   +FRIH  +  +++LK L LL EA +  Y K TG   GW + +YIFS LK
Sbjct  194  VYVLYKKR-LTVFRIHFFMLAVVILKALNLLCEAEDKSYIKRTGSAHGWDVLFYIFSFLK  252

Query  258  GTMMFVVIALIGTGWAFIKPFLGEKDKNIFLVVIPLQILANIAIIVLEETAP--------  309
            G  +F +I LIGTGW+F+KP+L +K+K + ++VIPLQ++ANIA +V++ET P
```

```
Sbjct  253  GITLFTLIVLIGTGWSFLKPYLQDKEKKVLMIVIPLQVIANIAQVVIDETGPFGQDWFTW  312

Query  310  -EVLRLVDIICCGAILVPIIWSIKHLRDAAAIDGKAKRNMEKLKLFREFYLLVVTYIYFT  368
             +V  LVD++CC A+L PI+WSIK+LR+AA  DGKA  N+ KL LFR++Y++V+ YIYFT
Sbjct  313  RQVFLLVDVVCCCAVLFPIVWSIKNLREAAKTDGKAAVNLMKLTLFRQYYIVVICYIYFT  372

Query  369  RIIVFLLDATLHYQYVWLGEFFTELATLIFWGLTGYKFRPVADNPYLKLDD  419
            R++ F L+    Y+Y+W     +ELATL F+  TGYKF+P A NPY  +DD
Sbjct  373  RVVFFALETITSYRYLWTSVVASELATLAFYVFTGYKFKPEAHNPYFVIDD  423
```

>protein GPR107-like [Lactuca sativa]
Sequence ID: XP_023754948.1 Length: 441
>hypothetical protein LSAT_8X4561 [Lactuca sativa]
Sequence ID: PLY92155.1 Length: 441
Range 1: 33 to 440

Score:277 bits(709), Expect:7e-85,
Method:Compositional matrix adjust.,
Identities:159/418(38%), Positives:236/418(56%), Gaps:26/418(6%)

```
Query  35   IKNDRRLAFRIETFGFFTGGVMEMAIENFKVVDDKGSLWDDLSA-GFIIKHIETDSG-SF  92
            I++D      I+ FGF   G + + +        +     DLS  GF +     DS
Sbjct  33   IRSDSNFNIPIDEFGFTHFGRLHLNLSQISFNPPE----PDLSRIGFFL--CTRDSWIHV  86

Query  93   IEETDASKCVSLLDEPERGDTIAVKVTKPSKENEKQSKVITIDKTRPEGFYTVVFLNCQP  152
            +E+  + K    LD P            KP+        V  +  T     +T+VF NC
Sbjct  87   VEQLQSEKIRCPLDSPAVKPVFTFNHLKPTTTTPSYDAVFNVTDTNQ---FTLVFANCAG  143

Query  153  GTYVSFDLTLTNYNPGP-----NYLSAGLTALPTLYAMLFVVWTVILGVWLFHFMRGQGK  207
            G  VS ++    YN  P     +YLSAG + LP++Y   F+++  +  +W++  +  +
Sbjct  144  GIKVSVNVLSVMYNLNPQNNRRDYLSAGKSNLPSIYFSFFLIYASLSCIWIYT-LHHRKS  202

Query  208  RIFRIHHLVTGIILLKLLTLLFEAIEFHYKKTTGHPGGWVIAYYIFSGLKGTMMFVVIAL  267
              +RIH+ +  ++LLK L LL E  +  Y K +G P GW + +YIFS LKG  +F +I L
Sbjct  203  SAYRIHYFMLAVLLLKALNLLCETEDKSYIKRSGTPHGWDVLFYIFSFLKGITLFTLIVL  262

Query  268  IGTGWAFIKPFLGEKDKNIFLVVIPLQILANIAIIVLEETAP---------EVLRLVDII  318
            IGTGW+F+KP+L +K+KN+ ++VIPLQ+++N+A +V++ET P         ++  LVDII
Sbjct  263  IGTGWSFLKPYLQDKEKNVLMIVIPLQVVSNLAQVVIDETGPFGQDLSTWRQIFLLVDII  322

Query  319  CCGAILVPIIWSIKHLRDAAAIDGKAKRNMEKLKLFREFYLLVVTYIYFTRIIVFLLDAT  378
            CC A+L PIIWSIK+LR+AA  DGKA  N+ KL LFR++Y++V+ YIYFTR+ V+ L+
Sbjct  323  CCCAVLFPIIWSIKNLREAAMTDGKAAVNLMKLTLFRQYYVVVICYIYFTRVAVYFLEMI  382

Query  379  LHYQYVWLGEFFTELATLIFWGLTGYKFRPVADNPYLKLDDEEEDAEREEAQRQSRTE  436
              Y+Y W   F  ELATL F+  TGY FRP   NPY  +DDEEE+A  +  + +   E
Sbjct  383  TSYRYAWTSVFLAELATLAFYVFTGYNFRPKTHNPYFAIDDEEEEAAVQALKLEDEFE  440
```

>hypothetical protein E1A91_A11G019800v1 [Gossypium mustelinum]
Sequence ID: TYJ07649.1 Length: 437

Range 1: 31 to 419

Score:277 bits(708), Expect:8e-85,
Method:Compositional matrix adjust.,
Identities:154/398(39%), Positives:232/398(58%), Gaps:22/398(5%)

```
Query  35   IKNDRRLAFRIETFGFFTGGVMEMAIENFKVVDDKGSLWDDLSAGFIIKHIETDSGSFIE  94
            I++D R     + FGF   G +E+ +    + D   +L D    GF +   +T      +
Sbjct  31   IRSDVRPIIPFDEFGFTHNGRLELNLSQIDLSDKNKNL-DLNKIGFFLCTRDTWFHVLEQ  89

Query  95   ETDASKCVSLLDEPERGDTIAVKVTKPSKENEKQSKVITIDKTRPEGFYTVVFLNCQPGT  154
              D     +L       D+  VKV    K  + ++ +  +        YT++F NC
Sbjct  90   LNDHHVTCAL-------DSELVKVVFRFKSLDGKTSINPVFPVNNADQYTLLFANCLTQV  142

Query  155  YVSFDLTLTNYN----PGPNYLSAGLTALPTLYAMLFVVWTVILGVWLFHFMRGQGKRIF  210
             VS  +    YN       +YLSAG T LP +Y +L +V+  + G+W++ F+  +   +F
Sbjct  143  KVSMTVRSAMYNIEGKQSRDYLSAGKTILPRVYFLLSLVYFSLAGIWVY-FLYKKRLTVF  201

Query  211  RIHHLVTGIILLKLLTLLFEAIEFHYKKTTGHPGGWVIAYYIFSGLKGTMMFVVIALIGT  270
            RIH  +  +I+LK   L+FEA +  Y K TG   GW + +YIFS LKG M+F +I LIGT
Sbjct  202  RIHFFMLAVIVLKAFNLVFEAEDKSYIKRTGSAHGWDVLFYIFSFLKGIMLFTLIVLIGT  261

Query  271  GWAFIKPFLGEKDKNIFLVVIPLQILANIAIIVLEETAP---------EVLRLVDIICCG  321
            GW+F+KP+L +K+K + ++VIPLQ++ANIA +V++E +P         ++  LVD+ICC
Sbjct  262  GWSFLKPYLQDKEKKVLMIVIPLQVVANIAQVVIDEASPFGQDRATWKQLFLLVDVICCC  321

Query  322  AILVPIIWSIKHLRDAAAIDGKAKRNMEKLKLFREFYLLVVTYIYFTRIIVFLLDATLHY  381
            A+L PI+WSIK+LR+AA  DGKA  N+ KL LFR++Y++V+ YIYFTR++V+ L+    Y
Sbjct  322  AVLFPIVWSIKNLREAARTDGKAAVNLMKLTLFRQYYVVVICYIYFTRVVVYALETITSY  381

Query  382  QYVWLGEFFTELATLIFWGLTGYKFRPVADNPYLKLDD  419
            +Y+W      ELATL F+  TGYKF+P A NPY  +DD
Sbjct  382  KYLWTAVLAGELATLAFYVFTGYKFKPEAHNPYFAIDD  419
```

>protein GPR107 [Carica papaya]
Sequence ID: XP_021905116.1 Length: 443
Range 1: 3 to 425

Score:277 bits(708), Expect:9e-85,
Method:Compositional matrix adjust.,
Identities:163/436(37%), Positives:245/436(56%), Gaps:36/436(8%)

```
Query  7    QTLMG---CFLIFCLV-AFLAPTANGLIHKLSIKNDRRLAFRIETFGFFTGGVMEMAIEN  62
            Q+ MG   CFL   L+ AF +  A+  I    I++D R     + FGF   G +E+ +
Sbjct  3    QSAMGFSHCFLSIVLIFAFTSFLASAEIRSSEIRSDDRPIIPFDEFGFNHAGRLELNVSR  62

Query  63   FKVVDDKGSLWDDLSAGFIIKHIETDSGSFIEETDA-SKCVSLLDEPERGDTIAVKVTKP  121
              + +    L  DLS            G F+   DA    +  L+E E    +   + KP
Sbjct  63   ISLSNSNPDL--DLS----------KVGFFLCTRDAWVHVIQQLEEGEVSCALQSDLVKP  110

Query  122  ----SKENEKQSKVITIDKTRPEGFYTVVFLNCQPGTYVSFDLTLTNYN-PGPN----YL  172
                +K    +    T+        YT+VF NC     VS ++    YN  G N    YL
```

```
Sbjct  111  VFTFNKLKGDKKSFETVYSENDADQYTLVFANCLNQVKVSMNVKSAMYNLEGKNRSRDYL  170

Query  173  SAGLTALPTLYAMLFVVWTVILGVWLFHFMRGQGKRIFRIHHLVTGIILLKLLTLLFEAI  232
            SAG T LP +Y +L +++  + G+W+   +  +   ++RIH  +  +++LK L LL EA 
Sbjct  171  SAGKTILPKVYFLLSLIYFCLAGLWI-SVLYKKRLTVYRIHFFMLAVVILKALNLLCEAE  229

Query  233  EFHYKKTTGHPGGWVIAYYIFSGLKGTMMFVVIALIGTGWAFIKPFLGEKDKNIFLVVIP  292
            +  Y K TG   GW + +Y+F+ LKG  +F +I LIGTGW+F+KP+L +K+K + ++VIP
Sbjct  230  DKSYIKRTGSAHGWDVLFYMFNFLKGITLFTLIVLIGTGWSFLKPYLQDKEKKVLMIVIP  289

Query  293  LQILANIAIIVLEETAP---------EVLRLVDIICCGAILVPIIWSIKHLRDAAAIDGK  343
            LQ++ANIA +V++E  P         ++  LVD++CC A+L PI+WSIK+LR+AA  DGK
Sbjct  290  LQVVANIAQVVIDENGPYGHDWITWKQIFLLVDVVCCCAVLFPIVWSIKNLREAARTDGK  349

Query  344  AKRNMEKLKLFREFYLLVVTYIYFTRIIVFLLDATLHYQYVWLGEFFTELATLIFWGLTG  403
            A  N+ KL LFR++Y++V+ YIYFTR++V+ L+    Y+Y+W      ELATL F+  TG
Sbjct  350  AAVNLMKLTLFRQYYIVVICYIYFTRVVVYALETITSYKYLWTSVVAGELATLAFYVFTG  409

Query  404  YKFRPVADNPYLKLDD  419
            YKF+P A NPY  +DD
Sbjct  410  YKFKPEAHNPYFVVDD  425
```
