## Supplemental Data 1 for "Overexpression of a G-protein coupled receptor-like gene affects encystment of *Acanthamoeba castellanii*"

RID: M0G0P173016
Job Title:Protein Sequence
Program: BLASTP
Query: None ID: lcl|Query_63602(amino acid) Length: 456
Database: nr All non-redundant GenBank CDS translations+PDB+SwissProt+PIR+PRF excluding environmental samples from WGS projects

Sequences producing significant alignments:

| Description | Max Score | Total Score | Query cover | E Value | Per. Ident | Accession |
|---|---|---|---|---|---|---|
| protein GPR107 isoform X1 [Homo sapiens] | 297 | 297 | 87% | 2e-94 | 37.17 | XP_016870442.1 |
| protein GPR107 isoform 3 [Homo sapiens] | 296 | 296 | 87% | 6e-94 | 37.17 | NP_066011.2 |
| unnamed protein product [Homo sapiens] | 295 | 295 | 87% | 3e-93 | 36.96 | BAF85781.1 |
| hypothetical protein [Homo sapiens] | 291 | 291 | 86% | 4e-92 | 36.93 | CAI46205.1 |
| protein GPR107 isoform 4 [Homo sapiens] | 283 | 283 | 64% | 2e-91 | 47.44 | NP_001274275.1 |
| protein GPR107 isoform 2 [Homo sapiens] | 285 | 285 | 87% | 4e-89 | 35.77 | NP_001130030.1 |
| KIAA1624 protein [Homo sapiens] | 274 | 274 | 87% | 8e-85 | 33.83 | BAB13450.1 |
| protein GPR107 isoform 1 precursor [Homo sapiens] | 274 | 274 | 87% | 1e-84 | 33.83 | NP_001130029.1 |
| protein GPR108 isoform X4 [Homo sapiens] | 250 | 250 | 72% | 4e-78 | 38.89 | XP_024307386.1 |
| protein GPR108 isoform 1 precursor [Homo sapiens] | 249 | 249 | 67% | 1e-75 | 41.30 | NP_001073921.1 |
| G protein-coupled receptor 108 [Homo sapiens] | 249 | 249 | 67% | 1e-75 | 41.30 | AAI46910.1 |
| hypothetical protein, similar to (AAF46469.1) CG12121 predicte... | 248 | 282 | 75% | 1e-75 | 41.30 | CAB96950.1 |
| unnamed protein product [Homo sapiens] | 247 | 281 | 75% | 6e-75 | 40.99 | BAF84657.1 |
| hCG1811164, isoform CRA_a [Homo sapiens] | 247 | 247 | 65% | 1e-74 | 41.99 | EAW69066.1 |
| protein GPR108 isoform 2 [Homo sapiens] | 233 | 233 | 59% | 2e-72 | 42.96 | NP_064556.1 |
| G protein-coupled receptor 108 [Homo sapiens] | 232 | 232 | 58% | 4e-72 | 42.86 | AAI50658.1 |
| protein GPR108 isoform X2 [Homo sapiens] | 221 | 221 | 67% | 2e-65 | 38.20 | XP_016882502.1 |
| G protein-coupled receptor 107, isoform CRA_a [Homo sapiens] | 185 | 185 | 68% | 2e-52 | 31.99 | EAW87920.1 |
| hCG1811164, isoform CRA_d [Homo sapiens] | 177 | 177 | 59% | 3e-51 | 37.72 | EAW69069.1 |
| protein GPR108 isoform X3 [Homo sapiens] | 178 | 178 | 50% | 3e-49 | 39.92 | XP_016882503.1 |
| protein GPR108 isoform X1 [Homo sapiens] | 177 | 177 | 50% | 7e-49 | 39.92 | XP_016882501.1 |

```
protein GPR108 isoform X6 [Homo sapiens]                            161    161
42%   5e-45 41.95  XP_016882504.1
GPR107 protein [Homo sapiens]                                       137    137
58%   2e-35 30.06  AAI43656.1
GPR108 protein [Homo sapiens]                                       75.1   75.1
17%   1e-15 45.00  AAH07862.1
unnamed protein product [Homo sapiens]                              47.4   47.4
40%   1e-04 22.52  BAB15408.1
```

Alignments:
>protein GPR107 isoform X1 [Homo sapiens]
Sequence ID: XP_016870442.1 Length: 521
Range 1: 39 to 520

Score:297 bits(760), Expect:2e-94,
Method:Compositional matrix adjust.,
Identities:181/487(37%), Positives:256/487(52%), Gaps:91/487(18%)

```
Query  28   GLIHKLSIKNDRRLAFRIETFGFFTGGVMEMAIENFKVVDDKGSLWDDLSAGFIIKHIET  87
            G +H L++K+D R    + TFGFF  G M + + +  + + +     D++ GF +   +
Sbjct  39   GRVHHLALKDDVRHKVHLNTFGFFKDGYMVVNVSSLSLNEPEDK---DVTIGFSLDRTKN  95

Query  88   DSGSFIEETDASKC--------VSLL----------------------------DEPERG  111
            D  S   + D + C        V+LL                            DE   G
Sbjct  96   DGFSSYLDEDVNYCILKKQSVSVTLLILDISRSEVRVKSPPEAGTQLPKIIFSRDEKVLG  155

Query  112  DTIAVKVTKPSKENEKQ-------SKVITIDKTR--------------------------  138
             +    V   S  N+ Q       SK  T+D
Sbjct  156  QSQEPNVNPASAGNQTQKTQDGGKSKRSTVDSKAMGEKSFSVHNNGGAVSFQFFNISTD  215

Query  139  -PEGFYTVVFLNCQ----PGTYVSF--DLTLTNYNPGPNYLSAGLTALPTLYAMLFVVWT  191
              EG Y++ F  C     P   +F  D+ +T  NP  +YLSAG   LP LY  +   +
Sbjct  216  DQEGLYSLYFHKCLGKELPSDKFTFSLDIEITEKNPD-SYLSAGEIPLPKLYISMAFFFF  274

Query  192  VILGVWLFHFMRGQGKRIFRIHHLVTGIILLKLLTLLFEAIEFHYKKTTGHP-GGWVIAY  250
            +   +W+ H +R +   +F+IH L+  +   K L+L+F AI++HY  + G P  GW + Y
Sbjct  275  LSGTIWI-HILRKRRNDVFKIHWLMAALPFTKSLSLVFHAIDYHYISSQGFPIEGWAVVY  333

Query  251  YIFSGLKGTMMFVVIALIGTGWAFIKPFLGEKDKNIFLVVIPLQILANIAIIVLEETA--  308
            YI   LKG ++F+ IALIGTGWAFIK  L +KDK IF++VIPLQ+LAN+A I++E T
Sbjct  334  YITHLLKGALLFITIALIGTGWAFIKHILSDKDKKIFMIVIPLQVLANVAYIIIESTEEG  393

Query  309  -------PEVLRLVDIICCGAILVPIIWSIKHLRDAAAIDGKAKRNMEKLKLFREFYLLV  361
                    + L LVD++CCGAIL P++WSI+HL++A+A DGKA  N+ KLKLFR +Y+L+
Sbjct  394  TTEYGLWKDSLFLVDLLCCGAILFPVVWSIRHLQEASATDGKAAINLAKLKLFRHYYVLI  453

Query  362  VTYIYFTRIIVFLLDATLHYQYVWLGEFFTELATLIFWGLTGYKFRPVADNPYLKLDDEE  421
            V YIYFTRII FLL   + +Q+ WL +   E ATL+F+ LTGYKFRP +DNPYL+L  EE
Sbjct  454  VCYIYFTRIIAFLLKLAVPFQWKWLYQLLDETATLVFFVLTGYKFRPASDNPYLQLSQEE  513

Query  422  EDAEREE  428
            ED E E
Sbjct  514  EDLEMES  520
```

>protein GPR107 isoform 3 [Homo sapiens]
Sequence ID: NP_066011.2 Length: 552
>G protein-coupled receptor 107 [Homo sapiens]
Sequence ID: AAI10519.1 Length: 552 >lung seven transmembrane receptor 1 [Homo sapiens]
Sequence ID: AAK57695.1 Length: 552 >G protein-coupled receptor 107, isoform CRA_d [Homo sapiens]
Sequence ID: EAW87924.1 Length: 552
Range 1: 39 to 520

Score:296 bits(759), Expect:6e-94,
Method:Compositional matrix adjust.,
Identities:181/487(37%), Positives:256/487(52%), Gaps:91/487(18%)

```
Query  28   GLIHKLSIKNDRRLAFRIETFGFFTGGVMEMAIENFKVVDDKGSLWDDLSAGFIIKHIET  87
            G +H L++K+D R    + TFGFF  G M + + +  + + +     D++ GF +   +
Sbjct  39   GRVHHLALKDDVRHKVHLNTFGFFKDGYMVVNVSSLSLNEPEDK---DVTIGFSLDRTKN  95

Query  88   DSGSFIEETDASKC--------VSLL----------------------------DEPERG  111
            D  S   + D + C        V+LL                            DE   G
Sbjct  96   DGFSSYLDEDVNYCILKKQSVSVTLLILDISRSEVRVKSPPEAGTQLPKIIFSRDEKVLG  155

Query  112  DTIAVKVTKPSKENEKQ-------SKVITIDKTR-------------------------  138
             +    V   S  N+ Q       SK  T+D
Sbjct  156  QSQEPNVNPASAGNQTQKTQDGGKSKRSTVDSKAMGEKSFSVHNNGGAVSFQFFFNISTD  215

Query  139  -PEGFYTVVFLNCQ----PGTYVSF--DLTLTNYNPGPNYLSAGLTALPTLYAMLFVVWT  191
              EG Y++ F  C     P    +F  D+ +T  NP  +YLSAG   LP LY  +   +
Sbjct  216  DQEGLYSLYFHKCLGKELPSDKFTFSLDIEITEKNPD-SYLSAGEIPLPKLYISMAFFFF  274

Query  192  VILGVWLFHFMRGQGKRIFRIHHLVTGIILLKLLTLLFEAIEFHYKKTTGHP-GGWVIAY  250
            +   +W+ H +R +   +F+IH L+  +   K L+L+F AI++HY  + G P  GW + Y
Sbjct  275  LSGTIWI-HILRKRRNDVFKIHWLMAALPFTKSLSLVFHAIDYHYISSQGFPIEGWAVVY  333

Query  251  YIFSGLKGTMMFVVIALIGTGWAFIKPFLGEKDKNIFLVVIPLQILANIAIIVLEETA--  308
            YI   LKG ++F+ IALIGTGWAFIK  L +KDK IF++VIPLQ+LAN+A I++E T
Sbjct  334  YITHLLKGALLFITIALIGTGWAFIKHILSDKDKKIFMIVIPLQVLANVAYIIIESTEEG  393

Query  309  -------PEVLRLVDIICCGAILVPIIWSIKHLRDAAAIDGKAKRNMEKLKLFREFYLLV  361
                    + L LVD++CCGAIL P++WSI+HL++A+A DGKA  N+ KLKLFR +Y+L+
Sbjct  394  TTEYGLWKDSLFLVDLLCCGAILFPVVWSIRHLQEASATDGKAAINLAKLKLFRHYYVLI  453

Query  362  VTYIYFTRIIVFLLDATLHYQYVWLGEFFTELATLIFWGLTGYKFRPVADNPYLKLDDEE  421
            V YIYFTRII FLL   + +Q+ WL +   E ATL+F+ LTGYKFRP +DNPYL+L  EE
Sbjct  454  VCYIYFTRIIAFLLKLAVPFQWKWLYQLLDETATLVFFVLTGYKFRPASDNPYLQLSQEE  513

Query  422  EDAEREE  428
            ED E E
Sbjct  514  EDLEMES  520
```

```
>unnamed protein product [Homo sapiens]
Sequence ID: BAF85781.1 Length: 552
Range 1: 39 to 520

Score:295 bits(754), Expect:3e-93,
Method:Compositional matrix adjust.,
Identities:180/487(37%), Positives:255/487(52%), Gaps:91/487(18%)

Query  28   GLIHKLSIKNDRRLAFRIETFGFFTGGVMEMAIENFKVVDDKGSLWDDLSAGFIIKHIET  87
            G +H L++K+D R    + TFGFF  G M + + +  + + +     D++ GF +   +
Sbjct  39   GRVHHLALKDDVRHKVHLNTFGFFKDGYMVVNVSSLSLNEPEDK---DVTIGFSLDRTKN  95

Query  88   DSGSFIEETDASKC--------VSLL----------------------------DEPERG  111
            D  S   + D + C        V+LL                            DE   G
Sbjct  96   DGFSSYLDEDVNYCILKKQSVSVTLLILDISRSEVRVKSPPEAGSQLPKIIFSRDEKVLG  155

Query  112  DTIAVKVTKPSKENEKQ-------SKVITIDKTR-------------------------  138
             +    V   S  N+ Q       SK  T+D
Sbjct  156  QSQEPNVNPASAGNQTQKTQDGGKSKRSTVDSKAMGEKSFSVHNNGGAVSFQFFFNISTD  215

Query  139  -PEGFYTVVFLNCQ----PGTYVSF--DLTLTNYNPGPNYLSAGLTALPTLYAMLFVVWT  191
              EG Y++ F  C     P    +F  D+ +T  NP  +YLSAG   LP LY  +   +
Sbjct  216  DQEGLYSLYFHKCLGKELPSDKFTFSLDIEITEKNPD-SYLSAGEIPLPKLYISMAFFFF  274

Query  192  VILGVWLFHFMRGQGKRIFRIHHLVTGIILLKLLTLLFEAIEFHYKKTTGHP-GGWVIAY  250
            +   +W+ H +R +   +F+IH L+  +   K L+L+F AI++HY  + G P  GW + Y
Sbjct  275  LSGTIWI-HILRKRRNDVFKIHWLMAALPFTKSLSLVFHAIDYHYISSQGFPIEGWAVVY  333

Query  251  YIFSGLKGTMMFVVIALIGTGWAFIKPFLGEKDKNIFLVVIPLQILANIAIIVLEETA--  308
            YI   LKG ++F+ IALIGTGWAFIK  L +KDK IF++VIPLQ+LAN+A I++E T
Sbjct  334  YITHLLKGALLFITIALIGTGWAFIKHILSDKDKKIFMIVIPLQVLANVAYIIIESTEEG  393

Query  309  -------PEVLRLVDIICCGAILVPIIWSIKHLRDAAAIDGKAKRNMEKLKLFREFYLLV  361
                    + L LVD++CCGAIL P++WSI+HL++A+A DGKA  N+ KLKLFR +Y+L+
Sbjct  394  TTEYGLWKDSLFLVDLLCCGAILFPVVWSIRHLQEASATDGKAAINLAKLKLFRHYYVLI  453

Query  362  VTYIYFTRIIVFLLDATLHYQYVWLGEFFTELATLIFWGLTGYKFRPVADNPYLKLDDEE  421
            V YIYF RII FLL   + +Q+ WL +   E ATL+F+ LTGYKFRP +DNPYL+L  EE
Sbjct  454  VCYIYFARIIAFLLKLAVPFQWKWLYQLLDETATLVFFVLTGYKFRPASDNPYLQLSQEE  513

Query  422  EDAEREE  428
            ED E E
Sbjct  514  EDLEMES  520



>hypothetical protein [Homo sapiens]
Sequence ID: CAI46205.1 Length: 509
Range 1: 1 to 477

Score:291 bits(744), Expect:4e-92,
Method:Compositional matrix adjust.,
Identities:178/482(37%), Positives:253/482(52%), Gaps:91/482(18%)
```

```
Query  33   LSIKNDRRLAFRIETFGFFTGGVMEMAIENFKVVDDKGSLWDDLSAGFIIKHIETDSGSF  92
            +++K+D R    + TFGFF  G M + + +  + + +     D++ GF +   + D  S
Sbjct  1    MALKDDVRHKVHLNTFGFFKDGYMVVNVSSLSLNEPEDK---DVTIGFSLDRTKNDGFSS  57

Query  93   IEETDASKC--------VSLL---------------------------DEPERGDTIAV  116
              + D + C        V+LL                            DE   G +
Sbjct  58   YLDEDVNYCILKKQSVSVTLLILDISRSEVRVKSPPEAGTQLPKIIFSRDEKVLGQSQEP  117

Query  117  KVTKPSKENEKQ-------SKVITIDKTR---------------------------PEGF  142
             V   S  N+ Q       SK  T+D                               EG
Sbjct  118  NVNPASAGNQTQKTQDGGKSKRSTVDSKAMGEKSFSVHNNGGAVSFQFFFNISTDDQEGL  177

Query  143  YTVVFLNCQ----PGTYVSF--DLTLTNYNPGPNYLSAGLTALPTLYAMLFVVWTVILGV  196
            Y++ F  C     P    +F  D+ +T  NP  +YLSAG   LP LY  +   + +   +
Sbjct  178  YSLYFHKCLGKELPSDKFTFSLDIEITEKNPD-SYLSAGEIPLPKLYISMAFFFFLSGTI  236

Query  197  WLFHFMRGQGKRIFRIHHLVTGIILLKLLTLLFEAIEFHYKKTTGHP-GGWVIAYYIFSG  255
            W+ H +R +   +F+IH L+  +   K L+L+F AI++HY  + G P  GW + YYI
Sbjct  237  WI-HILRKRRNDVFKIHWLMAALPFTKSLSLVFHAIDYHYISSQGFPIEGWAVVYYITHL  295

Query  256  LKGTMMFVVIALIGTGWAFIKPFLGEKDKNIFLVVIPLQILANIAIIVLEETA-------  308
            LKG ++F+ IALIGTGWAFIK  L +KDK IF++VIPLQ+LAN+A I++E T
Sbjct  296  LKGALLFITIALIGTGWAFIKHILSDKDKKIFMIVIPLQVLANVAYIIIESTEEGTTEYG  355

Query  309  --PEVLRLVDIICCGAILVPIIWSIKHLRDAAAIDGKAKRNMEKLKLFREFYLLVVTYIY  366
               + L LVD++CCGAIL P++WSI+HL++A+A DGKA  N+ KLKLFR +Y+L+V YIY
Sbjct  356  LWKDSLFLVDLLCCGAILFPVVWSIRHLQEASATDGKAAINLAKLKLFRHYYVLIVCYIY  415

Query  367  FTRIIVFLLDATLHYQYVWLGEFFTELATLIFWGLTGYKFRPVADNPYLKLDDEEEDAER  426
            FTRII FLL   + +Q+ WL +   E ATL+F+ LTGYKFRP +DNPYL+L  EEED E
Sbjct  416  FTRIIAFLLKLAVPFQWKWLYQLLDETATLVFFVLTGYKFRPASDNPYLQLSQEEEDLEM  475

Query  427  EE  428
            E
Sbjct  476  ES  477
```

>protein GPR107 isoform 4 [Homo sapiens]
Sequence ID: NP_001274275.1 Length: 332
>G protein-coupled receptor 107, isoform CRA_e [Homo sapiens]
Sequence ID: EAW87925.1 Length: 332
Range 1: 22 to 331

Score:283 bits(724), Expect:2e-91,
Method:Compositional matrix adjust.,
Identities:148/312(47%), Positives:202/312(64%), Gaps:18/312(5%)

```
Query  133  TIDKTRPEGFYTVVFLNCQ----PGTYVSF--DLTLTNYNPGPNYLSAGLTALPTLYAML  186
             I      EG Y++ F  C     P    +F  D+ +T  NP  +YLSAG   LP LY  +
Sbjct  22   NISTDDQEGLYSLYFHKCLGKELPSDKFTFSLDIEITEKNPD-SYLSAGEIPLPKLYISM  80

Query  187  FVVWTVILGVWLFHFMRGQGKRIFRIHHLVTGIILLKLLTLLFEAIEFHYKKTTGHP-GG  245
               + +   +W+ H +R +   +F+IH L+  +   K L+L+F AI++HY  + G P  G
```

```
Sbjct  81   AFFFFLSGTIWI-HILRKRRNDVFKIHWLMAALPFTKSLSLVFHAIDYHYISSQGFPIEG  139

Query  246  WVIAYYIFSGLKGTMMFVVIALIGTGWAFIKPFLGEKDKNIFLVVIPLQILANIAIIVLE  305
            W + YYI   LKG ++F+ IALIGTGWAFIK  L +KDK IF++VIPLQ+LAN+A I++E
Sbjct  140  WAVVYYITHLLKGALLFITIALIGTGWAFIKHILSDKDKKIFMIVIPLQVLANVAYIIIE  199

Query  306  ETA---------PEVLRLVDIICCGAILVPIIWSIKHLRDAAAIDGKAKRNMEKLKLFRE  356
             T           + L LVD++CCGAIL P++WSI+HL++A+A DGKA  N+ KLKLFR
Sbjct  200  STEEGTTEYGLWKDSLFLVDLLCCGAILFPVVWSIRHLQEASATDGKAAINLAKLKLFRH  259

Query  357  FYLLVVTYIYFTRIIVFLLDATLHYQYVWLGEFFTELATLIFWGLTGYKFRPVADNPYLK  416
            +Y+L+V YIYFTRII FLL   + +Q+ WL +   E ATL+F+ LTGYKFRP +DNPYL+
Sbjct  260  YYVLIVCYIYFTRIIAFLLKLAVPFQWKWLYQLLDETATLVFFVLTGYKFRPASDNPYLQ  319

Query  417  LDDEEEDAEREE  428
            L  EEED E E
Sbjct  320  LSQEEEDLEMES  331
```

>protein GPR107 isoform 2 [Homo sapiens]
Sequence ID: NP_001130030.1 Length: 571
>G protein-coupled receptor 107, isoform CRA_c [Homo sapiens]
Sequence ID: EAW87922.1 Length: 571 >G protein-coupled receptor 107, isoform CRA_c [Homo sapiens]
Sequence ID: EAW87923.1 Length: 571
Range 1: 39 to 539

Score:285 bits(728), Expect:4e-89,
Method:Compositional matrix adjust.,
Identities:181/506(36%), Positives:256/506(50%), Gaps:110/506(21%)

```
Query  28   GLIHKLSIKNDRRLAFRIETFGFFTGGVMEMAIENFKVVDDKGSLWDDLSAGFIIKHIET  87
            G +H L++K+D R    + TFGFF  G M + + +  + + +     D++ GF +   +
Sbjct  39   GRVHHLALKDDVRHKVHLNTFGFFKDGYMVVNVSSLSLNEPEDK---DVTIGFSLDRTKN  95

Query  88   DSGSFIEETDASKC--------VSLL----------------------------DEPERG  111
            D  S   + D + C        V+LL                            DE   G
Sbjct  96   DGFSSYLDEDVNYCILKKQSVSVTLLILDISRSEVRVKSPPEAGTQLPKIIFSRDEKVLG  155

Query  112  DTIAVKVTKPSKENEKQ-------SKVITIDKTR--------------------------  138
             +    V   S  N+ Q       SK  T+D
Sbjct  156  QSQEPNVNPASAGNQTQKTQDGGKSKRSTVDSKAMGEKSFSVHNNGGAVSFQFFNISTD  215

Query  139  -PEGFYTVVFLNCQ----PGTYVSF--DLTLTNYNPGPNYLSAGLTALPTLYAMLFVVWT  191
              EG Y++ F  C     P    +F  D+ +T  NP  +YLSAG   LP LY  +   +
Sbjct  216  DQEGLYSLYFHKCLGKELPSDKFTFSLDIEITEKNPD-SYLSAGEIPLPKLYISMAFFFF  274

Query  192  VILGVWLFHFMRGQGKRIFRIHHLVTGIILLKLLTLLFEAIEFHYKKTTGHP-GGWVIAY  250
            +   +W+ H +R +   +F+IH L+  +   K L+L+F AI++HY  + G P  GW + Y
Sbjct  275  LSGTIWI-HILRKRRNDVFKIHWLMAALPFTKSLSLVFHAIDYHYISSQGFPIEGWAVVY  333

Query  251  YIFSGLKGTMMFVVIALIGTGWAFIKPFLGEKDKNIFLVVIPLQILANIAIIVLEETA--  308
            YI   LKG ++F+ IALIGTGWAFIK  L +KDK IF++VIPLQ+LAN+A I++E T
```

```
Sbjct  334  YITHLLKGALLFITIALIGTGWAFIKHILSDKDKKIFMIVIPLQVLANVAYIIIESTEEG  393

Query  309  -------PEVLRLVDIICCGAILVPIIWSIKHLRDAAAIDGK------------------  343
                    + L LVD++CCGAIL P++WSI+HL++A+A DGK
Sbjct  394  TTEYGLWKDSLFLVDLLCCGAILFPVVWSIRHLQEASATDGKGDSMGPLQQRANLRAGSR  453

Query  344  -AKRNMEKLKLFREFYLLVVTYIYFTRIIVFLLDATLHYQYVWLGEFFTELATLIFWGLT  402
             A  N+ KLKLFR +Y+L+V YIYFTRII FLL   + +Q+ WL +   E ATL+F+ LT
Sbjct  454  IAAINLAKLKLFRHYYVLIVCYIYFTRIIAFLLKLAVPFQWKWLYQLLDETATLVFFVLT  513

Query  403  GYKFRPVADNPYLKLDDEEEDAEREE  428
            GYKFRP +DNPYL+L  EEED E E
Sbjct  514  GYKFRPASDNPYLQLSQEEEDLEMES  539
```

>KIAA1624 protein, partial [Homo sapiens]
Sequence ID: BAB13450.1 Length: 599
Range 1: 38 to 567

Score:274 bits(701), Expect:8e-85,
Method:Compositional matrix adjust.,
Identities:181/535(34%), Positives:256/535(47%), Gaps:139/535(25%)

```
Query  28   GLIHKLSIKNDRRLAFRIETFGFFTGGVMEMAIENFKVVDDKGSLWDDLSAGFIIKHIET  87
            G +H L++K+D R    + TFGFF  G M + + +  + + +     D++ GF +   +
Sbjct  38   GRVHHLALKDDVRHKVHLNTFGFFKDGYMVVNVSSLSLNEPEDK---DVTIGFSLDRTKN  94

Query  88   DSGSFIEETDASKC--------VSLL----------------------------DEPERG  111
            D  S   + D + C        V+LL                            DE   G
Sbjct  95   DGFSSYLDEDVNYCILKKQSVSVTLLILDISRSEVRVKSPPEAGTQLPKIIFSRDEKVLG  154

Query  112  DTIAVKVTKPSKENEKQ-------SKVITIDKTR--------------------------  138
             +    V   S  N+ Q       SK  T+D
Sbjct  155  QSQEPNVNPASAGNQTQKTQDGGKSKRSTVDSKAMGEKSFSVHNNGGAVSFQFFFNISTD  214

Query  139  -PEGFYTVVFLNCQ----PGTYVSF--DLTLTNYNPGPNYLSAGLTALPTLYAMLFVVWT  191
              EG Y++ F  C     P    +F  D+ +T  NP  +YLSAG   LP LY  +   +
Sbjct  215  DQEGLYSLYFHKCLGKELPSDKFTFSLDIEITEKNPD-SYLSAGEIPLPKLYISMAFFFF  273

Query  192  VILGVWLFHFMRGQGKRIFRIHHLVTGIILLKLLTLLFEAIEFHYKKTTGHP-GGWVIAY  250
            +   +W+ H +R +   +F+IH L+  +   K L+L+F AI++HY  + G P  GW + Y
Sbjct  274  LSGTIWI-HILRKRRNDVFKIHWLMAALPFTKSLSLVFHAIDYHYISSQGFPIEGWAVVY  332

Query  251  YIFSGLKGTMMFVVIALIGTGWAFIKPFLGEKDKNIFLVVIPLQILANIAIIVLEETA--  308
            YI   LKG ++F+ IALIGTGWAFIK  L +KDK IF++VIPLQ+LAN+A I++E T
Sbjct  333  YITHLLKGALLFITIALIGTGWAFIKHILSDKDKKIFMIVIPLQVLANVAYIIIESTEEG  392

Query  309  -------PEVLRLVDIICCGAILVPIIWSIKHLRDAAAIDGK------------------  343
                    + L LVD++CCGAIL P++WSI+HL++A+A DGK
Sbjct  393  TTEYGLWKDSLFLVDLLCCGAILFPVVWSIRHLQEASATDGKGDSMGPLQQRANLRAGSR  452

Query  344  ------------------------------AKRNMEKLKLFREFYLLVVTYIYFTRIIVF  373
                                          A  N+ KLKLFR +Y+L+V YIYFTRII F
```

```
Sbjct  453  IESRHFARADLELLASSCPPASVSQRAGITAAINLAKLKLFRHYYVLIVCYIYFTRIIAF  512

Query  374  LLDATLHYQYVWLGEFFTELATLIFWGLTGYKFRPVADNPYLKLDDEEEDAEREE  428
            LL   + +Q+ WL +   E ATL+F+ LTGYKFRP +DNPYL+L  EEED E E
Sbjct  513  LLKLAVPFQWKWLYQLLDETATLVFFVLTGYKFRPASDNPYLQLSQEEEDLEMES  567
```

>protein GPR107 isoform 1 precursor [Homo sapiens]
Sequence ID: NP_001130029.1 Length: 600
>RecName: Full=Protein GPR107; AltName: Full=Lung seven transmembrane receptor 1; Flags: Precursor [Homo sapiens]
Sequence ID: Q5VW38.1 Length: 600 >G protein-coupled receptor 107, isoform CRA_b [Homo sapiens]
Sequence ID: EAW87921.1 Length: 600
Range 1: 39 to 568

Score:274 bits(701), Expect:1e-84,
Method:Compositional matrix adjust.,
Identities:181/535(34%), Positives:256/535(47%), Gaps:139/535(25%)

```
Query  28   GLIHKLSIKNDRRLAFRIETFGFFTGGVMEMAIENFKVVDDKGSLWDDLSAGFIIKHIET  87
            G +H L++K+D R    + TFGFF  G M + + +  + + +     D++ GF +   +
Sbjct  39   GRVHHLALKDDVRHKVHLNTFGFFKDGYMVVNVSSLSLNEPEDK---DVTIGFSLDRTKN  95

Query  88   DSGSFIEETDASKC--------VSLL---------------------------DEPERG  111
            D  S   + D + C        V+LL                           DE   G
Sbjct  96   DGFSSYLDEDVNYCILKKQSVSVTLLILDISRSEVRVKSPPEAGTQLPKIIFSRDEKVLG  155

Query  112  DTIAVKVTKPSKENEKQ-------SKVITIDKTR-------------------------  138
             +    V   S  N+ Q       SK  T+D
Sbjct  156  QSQEPNVNPASAGNQTQKTQDGGKSKRSTVDSKAMGEKSFSVHNNGGAVSFQFFFNISTD  215

Query  139  -PEGFYTVVFLNCQ----PGTYVSF--DLTLTNYNPGPNYLSAGLTALPTLYAMLFVVWT  191
              EG Y++ F  C      P    +F  D+ +T  NP  +YLSAG   LP LY  +   +
Sbjct  216  DQEGLYSLYFHKCLGKELPSDKFTFSLDIEITEKNPD-SYLSAGEIPLPKLYISMAFFFF  274

Query  192  VILGVWLFHFMRGQGKRIFRIHHLVTGIILLKLLTLLFEAIEFHYKKTTGHP-GGWVIAY  250
            +   +W+ H +R +   +F+IH L+  +   K L+L+F AI++HY  + G P  GW + Y
Sbjct  275  LSGTIWI-HILRKRRNDVFKIHWLMAALPFTKSLSLVFHAIDYHYISSQGFPIEGWAVVY  333

Query  251  YIFSGLKGTMMFVVIALIGTGWAFIKPFLGEKDKNIFLVVIPLQILANIAIIVLEETA--  308
            YI   LKG ++F+ IALIGTGWAFIK  L +KDK IF++VIPLQ+LAN+A I++E T
Sbjct  334  YITHLLKGALLFITIALIGTGWAFIKHILSDKDKKIFMIVIPLQVLANVAYIIIESTEEG  393

Query  309  -------PEVLRLVDIICCGAILVPIIWSIKHLRDAAAIDGK------------------  343
                   + L LVD++CCGAIL P++WSI+HL++A+A DGK
Sbjct  394  TTEYGLWKDSLFLVDLLCCGAILFPVVWSIRHLQEASATDGKGDSMGPLQQRANLRAGSR  453

Query  344  ------------------------------AKRNMEKLKLFREFYLLVVTYIYFTRIIVF  373
                                          A  N+ KLKLFR +Y+L+V YIYFTRII F
Sbjct  454  IESHHFAQADLELLASSCPPASVSQRAGITAAINLAKLKLFRHYYVLIVCYIYFTRIIAF  513

Query  374  LLDATLHYQYVWLGEFFTELATLIFWGLTGYKFRPVADNPYLKLDDEEEDAEREE  428
```

```
            LL   + +Q+ WL +   E ATL+F+ LTGYKFRP +DNPYL+L  EEED E E
Sbjct  514  LLKLAVPFQWKWLYQLLDETATLVFFVLTGYKFRPASDNPYLQLSQEEEDLEMES  568
```

>protein GPR108 isoform X4 [Homo sapiens]
Sequence ID: XP_024307386.1 Length: 381
Range 1: 9 to 367

Score:250 bits(639), Expect:4e-78,
Method:Compositional matrix adjust.,
Identities:140/360(39%), Positives:207/360(57%), Gaps:29/360(8%)

```
Query  107  EPERGDTIAVKVTKPSKENEKQ-------------SKVITIDKTRPEGFYTVVFLNCQ--  151
            +PE  D  A+ +  PS +++               S  + I     EG Y++ F NC
Sbjct  9    DPEPADGGALSLQGPSGKDKDLVLGLSHLNNSYNFSFHVVIGSQAEEGQYSLNFHNCNNS  68

Query  152  -PGTYVSFDLTLTNYNPGPN-YLSAGLTALPTLYAMLFVVWTVILGVWLFHFMRGQGKRI  209
             PG    FD+T+      P+ +LSA    L  LY M+     +  G++    +      +
Sbjct  69   VPGKEHPFDITVMIREKNPDGFLSAAEMPLFKLY-MVMSACFLAAGIFWVSILCRNTYSV  127

Query  210  FRIHHLVTGIILLKLLTLLFEAIEFHYKKTTGHP-GGWVIAYYIFSGLKGTMMFVVIALI  268
            F+IH L+  +   K ++LLF +I +++  + GHP  G  + YYI   LKG ++F+ IALI
Sbjct  128  FKIHWLMAALAFTKSISLLFHSINYYFINSQGHPIEGLAVMYYIAHLLKGALLFITIALI  187

Query  269  GTGWAFIKPFLGEKDKNIFLVVIPLQILANIAIIVLEETA---------PEVLRLVDIIC  319
            G+GWAFIK  L +K+K +F +VIP+Q+LAN+A I++E             E+L LVD+IC
Sbjct  188  GSGWAFIKYVLSDKEKKVFGIVIPMQVLANVAYIIIESREEGASDYVLWKEILFLVDLIC  247

Query  320  CGAILVPIIWSIKHLRDAAAIDGKAKRNMEKLKLFREFYLLVVTYIYFTRIIVFLLDATL  379
            CGAIL P++WSI+HL+DA+  DGK   N+ KLKLFR +Y++V+ Y+YFTRII  LL   +
Sbjct  248  CGAILFPVVWSIRHLQDASGTDGKVAVNLAKLKLFRHYYVMVICYVYFTRIIAILLQVAV  307

Query  380  HYQYVWLGEFFTELATLIFWGLTGYKFRPVADNPYLKLDDE-EEDAEREEAQRQSRTEAG  438
             +Q+ WL +   E +TL F+ LTGYKF+P  +NPYL+L  E EED + E+    S    G
Sbjct  308  PFQWQWLYQLLVEGSTLAFFVLTGYKFQPTGNNPYLQLPQEDEEDVQMEQVMTDSGFREG  367
```

>protein GPR108 isoform 1 precursor [Homo sapiens]
Sequence ID: NP_001073921.1 Length: 543
>RecName: Full=Protein GPR108; AltName: Full=Lung seven transmembrane receptor 2; Flags: Precursor [Homo sapiens]
Sequence ID: Q9NPR9.3 Length: 543
Range 1: 209 to 529

Score:249 bits(635), Expect:1e-75,
Method:Compositional matrix adjust.,
Identities:133/322(41%), Positives:194/322(60%), Gaps:16/322(4%)

```
Query  132  ITIDKTRPEGFYTVVFLNCQ---PGTYVSFDLTLTNYNPGPN-YLSAGLTALPTLYAMLF  187
            + I     EG Y++ F NC    PG    FD+T+      P+ +LSA    L  LY M+
Sbjct  209  VVIGSQAEEGQYSLNFHNCNNSVPGKEHPFDITVMIREKNPDGFLSAAEMPLFKLY-MVM  267
```

```
Query  188  VVWTVILGVWLFHFMRGQGKRIFRIHHLVTGIILLKLLTLLFEAIEFHYKKTTGHP-GGW  246
                +  G++    +      +F+IH L+  +   K ++LLF +I +++  + GHP  G
Sbjct  268  SACFLAAGIFWVSILCRNTYSVFKIHWLMAALAFTKSISLLFHSINYYFINSQGHPIEGL  327

Query  247  VIAYYIFSGLKGTMMFVVIALIGTGWAFIKPFLGEKDKNIFLVVIPLQILANIAIIVLEE  306
             + YYI   LKG ++F+ IALIG+GWAFIK  L +K+K +F +VIP+Q+LAN+A I++E
Sbjct  328  AVMYYIAHLLKGALLFITIALIGSGWAFIKYVLSDKEKKVFGIVIPMQVLANVAYIIIES  387

Query  307  TA---------PEVLRLVDIICCGAILVPIIWSIKHLRDAAAIDGKAKRNMEKLKLFREF  357
                        E+L LVD+ICCGAIL P++WSI+HL+DA+  DGK   N+ KLKLFR +
Sbjct  388  REEGASDYVLWKEILFLVDLICCGAILFPVVWSIRHLQDASGTDGKVAVNLAKLKLFRHY  447

Query  358  YLLVVTYIYFTRIIVFLLDATLHYQYVWLGEFFTELATLIFWGLTGYKFRPVADNPYLKL  417
            Y++V+ Y+YFTRII  LL    + +Q+ WL +    E +TL F+ LTGYKF+P  +NPYL+L
Sbjct  448  YVMVICYVYFTRIIAILLQVAVPFQWQWLYQLLVEGSTLAFFVLTGYKFQPTGNNPYLQL  507

Query  418  DDE-EEDAEREEAQRQSRTEAG  438
              E EED + E+    S    G
Sbjct  508  PQEDEEDVQMEQVMTDSGFREG  529
```

>G protein-coupled receptor 108 [Homo sapiens]
Sequence ID: AAI46910.1 Length: 543
>hCG1811164, isoform CRA_b [Homo sapiens]
Sequence ID: EAW69067.1 Length: 543
Range 1: 209 to 529

Score:249 bits(635), Expect:1e-75,
Method:Compositional matrix adjust.,
Identities:133/322(41%), Positives:194/322(60%), Gaps:16/322(4%)

```
Query  132  ITIDKTRPEGFYTVVFLNCQ---PGTYVSFDLTLTNYNPGPN-YLSAGLTALPTLYAMLF  187
            + I     EG Y++ F NC    PG    FD+T+      P+ +LSA    L  LY M+
Sbjct  209  VVIGSQAEEGQYSLNFHNCNNSVPGKEHPFDITVMIREKNPDGFLSAAEMPLFKLY-MVM  267

Query  188  VVWTVILGVWLFHFMRGQGKRIFRIHHLVTGIILLKLLTLLFEAIEFHYKKTTGHP-GGW  246
                +  G++    +      +F+IH L+  +   K ++LLF +I +++  + GHP  G
Sbjct  268  SACFLAAGIFWVSILCRNTYSVFKIHWLMAALAFTKSISLLFHSINYYFINSQGHPIEGL  327

Query  247  VIAYYIFSGLKGTMMFVVIALIGTGWAFIKPFLGEKDKNIFLVVIPLQILANIAIIVLEE  306
             + YYI   LKG ++F+ IALIG+GWAFIK  L +K+K +F +VIP+Q+LAN+A I++E
Sbjct  328  AVMYYIAHLLKGALLFITIALIGSGWAFIKYVLSDKEKKVFGIVIPMQVLANVAYIIIES  387

Query  307  TA---------PEVLRLVDIICCGAILVPIIWSIKHLRDAAAIDGKAKRNMEKLKLFREF  357
                        E+L LVD+ICCGAIL P++WSI+HL+DA+  DGK   N+ KLKLFR +
Sbjct  388  REEGASDYVLWKEILFLVDLICCGAILFPVVWSIRHLQDASGTDGKVAVNLAKLKLFRHY  447

Query  358  YLLVVTYIYFTRIIVFLLDATLHYQYVWLGEFFTELATLIFWGLTGYKFRPVADNPYLKL  417
            Y++V+ Y+YFTRII  LL    + +Q+ WL +    E +TL F+ LTGYKF+P  +NPYL+L
Sbjct  448  YVMVICYVYFTRIIAILLQVAVPFQWQWLYQLLVEGSTLAFFVLTGYKFQPTGNNPYLQL  507

Query  418  DDE-EEDAEREEAQRQSRTEAG  438
              E EED + E+    S    G
```

```
Sbjct  508  PQEDEEDVQMEQVMTDSGFREG  529
```

>hypothetical protein, similar to (AAF46469.1) CG12121 predicted protein [Drosophila melanogaster], partial [Homo sapiens]
Sequence ID: CAB96950.1 Length: 536
Range 1: 202 to 522

Score:248 bits(634), Expect:1e-75,
Method:Compositional matrix adjust.,
Identities:133/322(41%), Positives:194/322(60%), Gaps:16/322(4%)

```
Query  132  ITIDKTRPEGFYTVVFLNCQ---PGTYVSFDLTLTNYNPGPN-YLSAGLTALPTLYAMLF  187
            + I     EG Y++ F NC    PG    FD+T+      P+ +LSA    L  LY M+
Sbjct  202  VVIGSQAEEGQYSLNFHNCNNSVPGKEHPFDITVMIREKNPDGFLSAAEMPLFKLY-MVM  260

Query  188  VVWTVILGVWLFHFMRGQGKRIFRIHHLVTGIILLKLLTLLFEAIEFHYKKTTGHP-GGW  246
                +  G++    +      +F+IH L+  +   K ++LLF +I +++  + GHP  G
Sbjct  261  SACFLAAGIFWVSILCRNTYSVFKIHWLMAALAFTKSISLLFHSINYYFINSQGHPIEGL  320

Query  247  VIAYYIFSGLKGTMMFVVIALIGTGWAFIKPFLGEKDKNIFLVVIPLQILANIAIIVLEE  306
             + YYI   LKG ++F+ IALIG+GWAFIK  L +K+K +F +VIP+Q+LAN+A I++E
Sbjct  321  AVMYYIAHLLKGALLFITIALIGSGWAFIKYVLSDKEKKVFGIVIPMQVLANVAYIIIES  380

Query  307  TA---------PEVLRLVDIICCGAILVPIIWSIKHLRDAAAIDGKAKRNMEKLKLFREF  357
                        E+L LVD+ICCGAIL P++WSI+HL+DA+  DGK   N+ KLKLFR +
Sbjct  381  REEGASDYVLWKEILFLVDLICCGAILFPVVWSIRHLQDASGTDGKVAVNLAKLKLFRHY  440

Query  358  YLLVVTYIYFTRIIVFLLDATLHYQYVWLGEFFTELATLIFWGLTGYKFRPVADNPYLKL  417
            Y++V+ Y+YFTRII  LL   + +Q+ WL +   E +TL F+ LTGYKF+P  +NPYL+L
Sbjct  441  YVMVICYVYFTRIIAILLQVAVPFQWQWLYQLLVEGSTLAFFVLTGYKFQPTGNNPYLQL  500

Query  418  DDE-EEDAEREEAQRQSRTEAG  438
              E EED + E+    S    G
Sbjct  501  PQEDEEDVQMEQVMTDSGFREG  522
```

Range 2: 23 to 58

Score:33.9 bits(76), Expect:3.0,
Method:Compositional matrix adjust.,
Identities:11/36(31%), Positives:24/36(66%), Gaps:0/36(0%)

```
Query  26  ANGLIHKLSIKNDRRLAFRIETFGFFTGGVMEMAIE  61
            +G IH+L++  ++R   ++ +FGF+T G +E+ +
Sbjct  23  CSGRIHRLALTGEKRADIQLNSFGFYTNGSLEVELS  58
```

>unnamed protein product [Homo sapiens]
Sequence ID: BAF84657.1 Length: 543

Range 1: 209 to 529

Score:247 bits(630), Expect:6e-75,
Method:Compositional matrix adjust.,
Identities:132/322(41%), Positives:193/322(59%), Gaps:16/322(4%)

```
Query  132  ITIDKTRPEGFYTVVFLNCQ---PGTYVSFDLTLTNYNPGPN-YLSAGLTALPTLYAMLF  187
            + I     EG Y++ F NC    PG    FD+T+      P+ +LSA    L  LY M+
Sbjct  209  VVIGSQAEEGQYSLNFHNCNNSVPGKEHPFDITVMIREKNPDGFLSAAEMPLFKLY-MVM  267

Query  188  VVWTVILGVWLFHFMRGQGKRIFRIHHLVTGIILLKLLTLLFEAIEFHYKKTTGHP-GGW  246
                +  G++    +      +F+IH L+  +   K ++LLF +I +++   + GHP  G
Sbjct  268  SACFLAAGIFWVSILCRNTYSVFKIHWLMAALAFTKSISLLFHSINYYFINSQGHPIEGL  327

Query  247  VIAYYIFSGLKGTMMFVVIALIGTGWAFIKPFLGEKDKNIFLVVIPLQILANIAIIVLEE  306
             + YYI   LKG ++F+ IALIG+GWAFIK  L +K+K +F +VIP+Q+LAN+  I++E
Sbjct  328  AVMYYIAHLLKGALLFITIALIGSGWAFIKYVLSDKEKKVFGIVIPMQVLANVTYIIIES  387

Query  307  TA---------PEVLRLVDIICCGAILVPIIWSIKHLRDAAAIDGKAKRNMEKLKLFREF  357
                        E+L LVD+ICCGAIL P++WSI+HL+DA+  DGK   N+ KLKLFR +
Sbjct  388  REEGASDYVLWKEILFLVDLICCGAILFPVVWSIRHLQDASGTDGKVAVNLAKLKLFRHY  447

Query  358  YLLVVTYIYFTRIIVFLLDATLHYQYVWLGEFFTELATLIFWGLTGYKFRPVADNPYLKL  417
            Y++V+ Y+YFTRII  LL   + +Q+ WL +   E +TL F+ LTGYKF+P  +NPYL+L
Sbjct  448  YVMVICYVYFTRIIAILLQVAVPFQWQWLYQLLVEGSTLAFFVLTGYKFQPTGNNPYLQL  507

Query  418  DDE-EEDAEREEAQRQSRTEAG  438
              E EED + E+    S    G
Sbjct  508  PQEDEEDVQMEQVMTDSGFREG  529
```

Range 2: 30 to 65

Score:33.9 bits(76), Expect:3.0,
Method:Compositional matrix adjust.,
Identities:11/36(31%), Positives:24/36(66%), Gaps:0/36(0%)

```
Query  26  ANGLIHKLSIKNDRRLAFRIETFGFFTGGVMEMAIE  61
            +G IH+L++  ++R   ++ +FGF+T G +E+ +
Sbjct  30  CSGRIHRLALTGEKRADIQLNSFGFYTNGSLEVELS  65
```

>hCG1811164, isoform CRA_a [Homo sapiens]
Sequence ID: EAW69066.1 Length: 575
Range 1: 209 to 519

Score:247 bits(630), Expect:1e-74,
Method:Compositional matrix adjust.,
Identities:131/312(42%), Positives:192/312(61%), Gaps:16/312(5%)

```
Query  132  ITIDKTRPEGFYTVVFLNCQ---PGTYVSFDLTLTNYNPGPN-YLSAGLTALPTLYAMLF  187
```

```
                + I     EG Y++ F NC    PG    FD+T+      P+ +LSA    L  LY M+
Sbjct  209  VVIGSQAEEGQYSLNFHNCNNSVPGKEHPFDITVMIREKNPDGFLSAAEMPLFKLY-MVM  267

Query  188  VVWTVILGVWLFHFMRGQGKRIFRIHHLVTGIILLKLLTLLFEAIEFHYKKTTGHP-GGW  246
                +  G++    +      +F+IH L+  +   K ++LLF +I +++  + GHP  G
Sbjct  268  SACFLAAGIFWVSILCRNTYSVFKIHWLMAALAFTKSISLLFHSINYYFINSQGHPIEGL  327

Query  247  VIAYYIFSGLKGTMMFVVIALIGTGWAFIKPFLGEKDKNIFLVVIPLQILANIAIIVLEE  306
              + YYI   LKG ++F+ IALIG+GWAFIK  L +K+K +F +VIP+Q+LAN+A I++E
Sbjct  328  AVMYYIAHLLKGALLFITIALIGSGWAFIKYVLSDKEKKVFGIVIPMQVLANVAYIIIES  387

Query  307  TA---------PEVLRLVDIICCGAILVPIIWSIKHLRDAAAIDGKAKRNMEKLKLFREF  357
                        E+L LVD+ICCGAIL P++WSI+HL+DA+  DGK   N+ KLKLFR +
Sbjct  388  REEGASDYVLWKEILFLVDLICCGAILFPVVWSIRHLQDASGTDGKVAVNLAKLKLFRHY  447

Query  358  YLLVVTYIYFTRIIVFLLDATLHYQYVWLGEFFTELATLIFWGLTGYKFRPVADNPYLKL  417
            Y++V+ Y+YFTRII  LL   + +Q+ WL +   E +TL F+ LTGYKF+P  +NPYL+L
Sbjct  448  YVMVICYVYFTRIIAILLQVAVPFQWQWLYQLLVEGSTLAFFVLTGYKFQPTGNNPYLQL  507

Query  418  DDE-EEDAEREE  428
              E EED + E+
Sbjct  508  PQEDEEDVQMEQ  519
```

>protein GPR108 isoform 2 [Homo sapiens]
Sequence ID: NP_064556.1 Length: 301
>protein GPR108 isoform X5 [Homo sapiens]
Sequence ID: XP_011526439.1 Length: 301 >hCG1811164, isoform CRA_c [Homo sapiens]
Sequence ID: EAW69068.1 Length: 301
Range 1: 6 to 287

Score:233 bits(594), Expect:2e-72,
Method:Compositional matrix adjust.,
Identities:122/284(43%), Positives:177/284(62%), Gaps:13/284(4%)

```
Query  166  NPGPNYLSAGLTALPTLYAMLFVVWTVILGVWLFHFMRGQGKRIFRIHHLVTGIILLKLL  225
            NP   +LSA    L  LY M+     +  G++    +      +F+IH L+  +   K +
Sbjct  6    NPD-GFLSAAEMPLFKLY-MVMSACFLAAGIFWVSILCRNTYSVFKIHWLMAALAFTKSI  63

Query  226  TLLFEAIEFHYKKTTGHP-GGWVIAYYIFSGLKGTMMFVVIALIGTGWAFIKPFLGEKDK  284
            +LLF +I +++  + GHP  G  + YYI   LKG ++F+ IALIG+GWAFIK  L +K+K
Sbjct  64   SLLFHSINYYFINSQGHPIEGLAVMYYIAHLLKGALLFITIALIGSGWAFIKYVLSDKEK  123

Query  285  NIFLVVIPLQILANIAIIVLEETA---------PEVLRLVDIICCGAILVPIIWSIKHLR  335
             +F +VIP+Q+LAN+A I++E            E+L LVD+ICCGAIL P++WSI+HL+
Sbjct  124  KVFGIVIPMQVLANVAYIIIESREEGASDYVLWKEILFLVDLICCGAILFPVVWSIRHLQ  183

Query  336  DAAAIDGKAKRNMEKLKLFREFYLLVVTYIYFTRIIVFLLDATLHYQYVWLGEFFTELAT  395
            DA+  DGK   N+ KLKLFR +Y++V+ Y+YFTRII  LL   + +Q+ WL +   E +T
Sbjct  184  DASGTDGKVAVNLAKLKLFRHYYVMVICYVYFTRIIAILLQVAVPFQWQWLYQLLVEGST  243

Query  396  LIFWGLTGYKFRPVADNPYLKLDDE-EEDAEREEAQRQSRTEAG  438
```

```
             L F+ LTGYKF+P  +NPYL+L  E EED + E+    S    G
Sbjct  244  LAFFVLTGYKFQPTGNNPYLQLPQEDEEDVQMEQVMTDSGFREG  287



>G protein-coupled receptor 108 [Homo sapiens]
Sequence ID: AAI50658.1 Length: 301
Range 1: 9 to 287

Score:232 bits(592), Expect:4e-72,
Method:Compositional matrix adjust.,
Identities:120/280(43%), Positives:175/280(62%), Gaps:12/280(4%)

Query  170  NYLSAGLTALPTLYAMLFVVWTVILGVWLFHFMRGQGKRIFRIHHLVTGIILLKLLTLLF  229
             +LSA    L  LY M+     +  G++    +      +F+IH L+  +   K ++LLF
Sbjct  9    GFLSAAEMPLFKLY-MVMSACFLAAGIFWVSILCRNTYSVFKIHWLMAALAFTKSISLLF  67

Query  230  EAIEFHYKKTTGHP-GGWVIAYYIFSGLKGTMMFVVIALIGTGWAFIKPFLGEKDKNIFL  288
             +I +++  + GHP  G  + YYI   LKG ++F+ IALIG+GWAFIK  L +K+K +F
Sbjct  68   HSINYYFINSQGHPIEGLAVMYYIAHLLKGALLFITIALIGSGWAFIKYVLSDKEKKVFG  127

Query  289  VVIPLQILANIAIIVLEETA---------PEVLRLVDIICCGAILVPIIWSIKHLRDAAA  339
            +VIP+Q+LAN+A I++E             E+L LVD+ICCGAIL P++WSI+HL+DA+
Sbjct  128  IVIPMQVLANVAYIIIESREEGASDYVLWKEILFLVDLICCGAILFPVVWSIRHLQDASG  187

Query  340  IDGKAKRNMEKLKLFREFYLLVVTYIYFTRIIVFLLDATLHYQYVWLGEFFTELATLIFW  399
             DGK   N+ KLKLFR +Y++V+ Y+YFTRII  LL   + +Q+ WL +   E +TL F+
Sbjct  188  TDGKVAVNLAKLKLFRHYYVMVICYVYFTRIIAILLQVAVPFQWQWLYQLLVEGSTLAFF  247

Query  400  GLTGYKFRPVADNPYLKLDDE-EEDAEREEAQRQSRTEAG  438
             LTGYKF+P  +NPYL+L  E EED + E+    S    G
Sbjct  248  VLTGYKFQPTGNNPYLQLPQEDEEDVQMEQVMTDSGFREG  287



>protein GPR108 isoform X2 [Homo sapiens]
Sequence ID: XP_016882502.1 Length: 524
Range 1: 209 to 510

Score:221 bits(564), Expect:2e-65,
Method:Compositional matrix adjust.,
Identities:123/322(38%), Positives:182/322(56%), Gaps:35/322(10%)

Query  132  ITIDKTRPEGFYTVVFLNCQ---PGTYVSFDLTLTNYNPGPN-YLSAGLTALPTLYAMLF  187
            + I     EG Y++ F NC    PG    FD+T+      P+ +LSA    L  LY M+
Sbjct  209  VVIGSQAEEGQYSLNFHNCNNSVPGKEHPFDITVMIREKNPDGFLSAAEMPLFKLY-MVM  267

Query  188  VVWTVILGVWLFHFMRGQGKRIFRIHHLVTGIILLKLLTLLFEAIEFHYKKTTGHP-GGW  246
                +  G++    +      +F+IH L+  +   K ++LLF +I +++  + GHP  G
Sbjct  268  SACFLAAGIFWVSILCRNTYSVFKIHWLMAALAFTKSISLLFHSINYYFINSQGHPIEGL  327

Query  247  VIAYYIFSGLKGTMMFVVIALIGTGWAFIKPFLGEKDKNIFLVVIPLQILANIAIIVLEE  306
             + YYI   LKG ++F+ IALIG+GWAFIK  L +K+K +F +VIP+Q+LAN+A I++E
Sbjct  328  AVMYYIAHLLKGALLFITIALIGSGWAFIKYVLSDKEKKVFGIVIPMQVLANVAYIIIES  387
```

```
Query  307  TA---------PEVLRLVDIICCGAILVPIIWSIKHLRDAAAIDGKAKRNMEKLKLFREF  357
                        E+L LVD+ICCGAIL P++WSI+HL+DA+  DGK   N+ KLKLFR +
Sbjct  388  REEGASDYVLWKEILFLVDLICCGAILFPVVWSIRHLQDASGTDGKVAVNLAKLKLFRHY  447

Query  358  YLLVVTYIYFTRIIVFLLDATLHYQYVWLGEFFTELATLIFWGLTGYKFRPVADNPYLKL  417
            Y++V+ Y+YFTRII  LL   + +Q+ WL                   ++P  +NPYL+L
Sbjct  448  YVMVICYVYFTRIIAILLQVAVPFQWQWL-------------------YQPTGNNPYLQL  488

Query  418  DDE-EEDAEREEAQRQSRTEAG  438
              E EED + E+    S    G
Sbjct  489  PQEDEEDVQMEQVMTDSGFREG  510
```

>G protein-coupled receptor 107, isoform CRA_a [Homo sapiens]
Sequence ID: EAW87920.1 Length: 440
Range 1: 39 to 430

Score:185 bits(469), Expect:2e-52,
Method:Compositional matrix adjust.,
Identities:127/397(32%), Positives:190/397(47%), Gaps:91/397(22%)

```
Query  28   GLIHKLSIKNDRRLAFRIETFGFFTGGVMEMAIENFKVVDDKGSLWDDLSAGFIIKHIET  87
            G +H L++K+D R    + TFGFF  G M + + +  + + +     D++ GF +   +
Sbjct  39   GRVHHLALKDDVRHKVHLNTFGFFKDGYMVVNVSSLSLNEPEDK---DVTIGFSLDRTKN  95

Query  88   DSGSFIEETDASKC--------VSLL----------------------------DEPERG  111
            D  S   + D + C        V+LL                            DE   G
Sbjct  96   DGFSSYLDEDVNYCILKKQSVSVTLLILDISRSEVRVKSPPEAGTQLPKIIFSRDEKVLG  155

Query  112  DTIAVKVTKPSKENEKQ-------SKVITIDKTR--------------------------  138
             +    V   S  N+ Q       SK  T+D
Sbjct  156  QSQEPNVNPASAGNQTQKTQDGGKSKRSTVDSKAMGEKSFSVHNNGGAVSFQFFFNISTD  215

Query  139  -PEGFYTVVFLNCQ----PGTYVSF--DLTLTNYNPGPNYLSAGLTALPTLYAMLFVVWT  191
              EG Y++ F  C     P    +F  D+ +T  NP  +YLSAG   LP LY  +   +
Sbjct  216  DQEGLYSLYFHKCLGKELPSDKFTFSLDIEITEKNPD-SYLSAGEIPLPKLYISMAFFFF  274

Query  192  VILGVWLFHFMRGQGKRIFRIHHLVTGIILLKLLTLLFEAIEFHYKKTTGHP-GGWVIAY  250
            +   +W+ H +R +   +F+IH L+  +   K L+L+F AI++HY  + G P  GW + Y
Sbjct  275  LSGTIWI-HILRKRRNDVFKIHWLMAALPFTKSLSLVFHAIDYHYISSQGFPIEGWAVVY  333

Query  251  YIFSGLKGTMMFVVIALIGTGWAFIKPFLGEKDKNIFLVVIPLQILANIAIIVLEETA--  308
            YI   LKG ++F+ IALIGTGWAFIK  L +KDK IF++VIPLQ+LAN+A I++E T
Sbjct  334  YITHLLKGALLFITIALIGTGWAFIKHILSDKDKKIFMIVIPLQVLANVAYIIIESTEEG  393

Query  309  -------PEVLRLVDIICCGAILVPIIWSIKHLRDAA  338
                    + L LVD++CCGAIL P++WSI+HL++A+
Sbjct  394  TTEYGLWKDSLFLVDLLCCGAILFPVVWSIRHLQEAS  430
```

>hCG1811164, isoform CRA_d [Homo sapiens]

```
Sequence ID: EAW69069.1 Length: 297
Range 1: 6 to 283

Score:177 bits(450), Expect:3e-51,
Method:Compositional matrix adjust.,
Identities:109/289(38%), Positives:160/289(55%), Gaps:27/289(9%)

Query  166  NPGPNYLSAGLTALPTLYAMLFVVWTVILGVWLFHFMRGQGKRIFRIHHLVTGIILLKLL  225
            NP   +LSA    L  LY M+     +  G++    +      +F+IH L+  +   K +
Sbjct  6    NPD-GFLSAAEMPLFKLY-MVMSACFLAAGIFWVSILCRNTYSVFKIHWLMAALAFTKSI  63

Query  226  TLLFEAIEFHYKKTTGHP-GGWVIAYYIFSGLKGTMMFVVIALIGTGWAFIKPFLGEKDK  284
            +LLF +I +++  + GHP  G  + YYI   LKG ++F+ IALIG+GWAFIK  L +K+K
Sbjct  64   SLLFHSINYYFINSQGHPIEGLAVMYYIAHLLKGALLFITIALIGSGWAFIKYVLSDKEK  123

Query  285  NIFLVVIPLQILANIAIIVLEETA---------PEVLRLVDIICCGAILVPIIWSIKHLR  335
             +F +VIP+Q+LAN+A I++E             E+L LVD+ICCGAIL P++
Sbjct  124  KVFGIVIPMQVLANVAYIIIESREEGASDYVLWKEILFLVDLICCGAILFPVVCG-SEPG  182

Query  336  DAAAIDG-----KAKRNMEKLKLFREFYLLVVTYIYFTRIIVFLLDATLHYQYVWLGEFF  390
             A A+        A R +  L+        V+ Y+YFTRII  LL   + +Q+ WL +
Sbjct  183  QAEAVPALLCHVLAPRLLAPLQ--------VICYVYFTRIIAILLQVAVPFQWQWLYQLL  234

Query  391  TELATLIFWGLTGYKFRPVADNPYLKLDDE-EEDAEREEAQRQSRTEAG  438
             E +TL F+ LTGYKF+P  +NPYL+L  E EED + E+    S    G
Sbjct  235  VEGSTLAFFVLTGYKFQPTGNNPYLQLPQEDEEDVQMEQVMTDSGFREG  283


>protein GPR108 isoform X3 [Homo sapiens]
Sequence ID: XP_016882503.1 Length: 511
Range 1: 209 to 450

Score:178 bits(451), Expect:3e-49,
Method:Compositional matrix adjust.,
Identities:97/243(40%), Positives:145/243(59%), Gaps:15/243(6%)

Query  132  ITIDKTRPEGFYTVVFLNCQ---PGTYVSFDLTLTNYNPGPN-YLSAGLTALPTLYAMLF  187
            + I     EG Y++ F NC    PG    FD+T+      P+ +LSA    L  LY M+
Sbjct  209  VVIGSQAEEGQYSLNFHNCNNSVPGKEHPFDITVMIREKNPDGFLSAAEMPLFKLY-MVM  267

Query  188  VVWTVILGVWLFHFMRGQGKRIFRIHHLVTGIILLKLLTLLFEAIEFHYKKTTGHP-GGW  246
                +  G++    +      +F+IH L+  +   K ++LLF +I +++  + GHP  G
Sbjct  268  SACFLAAGIFWVSILCRNTYSVFKIHWLMAALAFTKSISLLFHSINYYFINSQGHPIEGL  327

Query  247  VIAYYIFSGLKGTMMFVVIALIGTGWAFIKPFLGEKDKNIFLVVIPLQILANIAIIVLEE  306
              + YYI   LKG ++F+ IALIG+GWAFIK  L +K+K +F +VIP+Q+LAN+A I++E
Sbjct  328  AVMYYIAHLLKGALLFITIALIGSGWAFIKYVLSDKEKKVFGIVIPMQVLANVAYIIIES  387

Query  307  TA---------PEVLRLVDIICCGAILVPIIWSIKHLRDAAAIDGKAKRNMEKLKLFREF  357
                        E+L LVD+ICCGAIL P++WSI+HL+DA+  DGK   N+ KLKLFR +
Sbjct  388  REEGASDYVLWKEILFLVDLICCGAILFPVVWSIRHLQDASGTDGKVAVNLAKLKLFRHY  447

Query  358  YLL  360
```

```
            Y++
Sbjct  448  YVM  450
```

>protein GPR108 isoform X1 [Homo sapiens]
Sequence ID: XP_016882501.1 Length: 530
Range 1: 209 to 450

Score:177 bits(450), Expect:7e-49,
Method:Compositional matrix adjust.,
Identities:97/243(40%), Positives:145/243(59%), Gaps:15/243(6%)

```
Query  132  ITIDKTRPEGFYTVVFLNCQ---PGTYVSFDLTLTNYNPGPN-YLSAGLTALPTLYAMLF  187
            + I     EG Y++ F NC    PG    FD+T+      P+ +LSA    L  LY M+
Sbjct  209  VVIGSQAEEGQYSLNFHNCNNSVPGKEHPFDITVMIREKNPDGFLSAAEMPLFKLY-MVM  267

Query  188  VVWTVILGVWLFHFMRGQGKRIFRIHHLVTGIILLKLLTLLFEAIEFHYKKTTGHP-GGW  246
                +  G++    +      +F+IH L+  +   K ++LLF +I +++  + GHP  G
Sbjct  268  SACFLAAGIFWVSILCRNTYSVFKIHWLMAALAFTKSISLLFHSINYYFINSQGHPIEGL  327

Query  247  VIAYYIFSGLKGTMMFVVIALIGTGWAFIKPFLGEKDKNIFLVVIPLQILANIAIIVLEE  306
             + YYI   LKG ++F+ IALIG+GWAFIK  L +K+K +F +VIP+Q+LAN+A I++E
Sbjct  328  AVMYYIAHLLKGALLFITIALIGSGWAFIKYVLSDKEKKVFGIVIPMQVLANVAYIIIES  387

Query  307  TA---------PEVLRLVDIICCGAILVPIIWSIKHLRDAAAIDGKAKRNMEKLKLFREF  357
                        E+L LVD+ICCGAIL P++WSI+HL+DA+  DGK   N+ KLKLFR +
Sbjct  388  REEGASDYVLWKEILFLVDLICCGAILFPVVWSIRHLQDASGTDGKVAVNLAKLKLFRHY  447

Query  358  YLL  360
            Y++
Sbjct  448  YVM  450
```

>protein GPR108 isoform X6 [Homo sapiens]
Sequence ID: XP_016882504.1 Length: 288
Range 1: 6 to 208

Score:161 bits(407), Expect:5e-45,
Method:Compositional matrix adjust.,
Identities:86/205(42%), Positives:128/205(62%), Gaps:12/205(5%)

```
Query  166  NPGPNYLSAGLTALPTLYAMLFVVWTVILGVWLFHFMRGQGKRIFRIHHLVTGIILLKLL  225
            NP   +LSA    L  LY M+     +  G++    +      +F+IH L+  +   K +
Sbjct  6    NPD-GFLSAAEMPLFKLY-MVMSACFLAAGIFWVSILCRNTYSVFKIHWLMAALAFTKSI  63

Query  226  TLLFEAIEFHYKKTTGHP-GGWVIAYYIFSGLKGTMMFVVIALIGTGWAFIKPFLGEKDK  284
            +LLF +I +++  + GHP  G  + YYI   LKG ++F+ IALIG+GWAFIK  L +K+K
Sbjct  64   SLLFHSINYYFINSQGHPIEGLAVMYYIAHLLKGALLFITIALIGSGWAFIKYVLSDKEK  123

Query  285  NIFLVVIPLQILANIAIIVLEETA---------PEVLRLVDIICCGAILVPIIWSIKHLR  335
             +F +VIP+Q+LAN+A I++E             E+L LVD+ICCGAIL P++WSI+HL+
Sbjct  124  KVFGIVIPMQVLANVAYIIIESREEGASDYVLWKEILFLVDLICCGAILFPVVWSIRHLQ  183
```

```
Query  336  DAAAIDGKAKRNMEKLKLFREFYLL  360
            DA+  DGK   N+ KLKLFR +Y++
Sbjct  184  DASGTDGKVAVNLAKLKLFRHYYVM  208
```

>GPR107 protein, partial [Homo sapiens]
Sequence ID: AAI43656.1 Length: 379
Range 1: 39 to 379

Score:137 bits(346), Expect:2e-35,
Method:Compositional matrix adjust.,
Identities:104/346(30%), Positives:155/346(44%), Gaps:82/346(23%)

```
Query  28   GLIHKLSIKNDRRLAFRIETFGFFTGGVMEMAIENFKVVDDKGSLWDDLSAGFIIKHIET  87
            G +H L++K+D R    + TFGFF  G M + + +  + + +     D++ GF +   +
Sbjct  39   GRVHHLALKDDVRHKVHLNTFGFFKDGYMVVNVSSLSLNEPEDK---DVTIGFSLDRTKN  95

Query  88   DSGSFIEETDASKC--------VSLL----------------------------DEPERG  111
            D  S   + D + C        V+LL                            DE   G
Sbjct  96   DGFSSYLDEDVNYCILKKQSVSVTLLILDISRSEVRVKSPPEAGTQLPKIIFSRDEKVLG  155

Query  112  DTIAVKVTKPSKENEKQ-------SKVITIDKTR--------------------------  138
             +    V   S  N+ Q       SK  T+D
Sbjct  156  QSQEPNVNPASAGNQTQKTQDGGKSKRSTVDSKAMGEKSFSVHHNNGGAVSFQFFFNISTD  215

Query  139  -PEGFYTVVFLNCQ----PGTYVSF--DLTLTNYNPGPNYLSAGLTALPTLYAMLFVVWT  191
              EG Y++ F  C     P    +F  D+ +T  NP  +YLSAG   LP LY  +   +
Sbjct  216  DQEGLYSLYFHKCLGKELPSDKFTFSLDIEITEKNPD-SYLSAGEIPLPKLYISMAFFFF  274

Query  192  VILGVWLFHFMRGQGKRIFRIHHLVTGIILLKLLTLLFEAIEFHYKKTTGHP-GGWVIAY  250
            +   +W+ H +R +   +F+IH L+  +   K L+L+F AI++HY  + G P  GW + Y
Sbjct  275  LSGTIWI-HILRKRRNDVFKIHWLMAALPFTKSLSLVFHAIDYHYISSQGFPIEGWAVVY  333

Query  251  YIFSGLKGTMMFVVIALIGTGWAFIKPFLGEKDKNIFLVVIPLQIL  296
            YI   LKG ++F+ IALIGTGWAFIK  L +KDK IF++VIPLQ+L
Sbjct  334  YITHLLKGALLFITIALIGTGWAFIKHILSDKDKKIFMIVIPLQVL  379
```

>GPR108 protein [Homo sapiens]
Sequence ID: AAH07862.1 Length: 94
Range 1: 1 to 80

Score:75.1 bits(183), Expect:1e-15,
Method:Composition-based stats.,
Identities:36/80(45%), Positives:50/80(62%), Gaps:1/80(1%)

```
Query  360  LVVTYIYFTRIIVFLLDATLHYQYVWLGEFFTELATLIFWGLTGYKFRPVADNPYLKLDD  419
            +V+ Y+YFTRII  LL   + +Q+ WL +   E +TL F+ LTGYKF+P  +NPYL+L
Sbjct  1    MVICYVYFTRIIAILLQVAVPFQWQWLYQLLVEGSTLAFFVLTGYKFQPTGNNPYLQLPQ  60

Query  420  E-EEDAEREEAQRQSRTEAG  438
```

```
            E EED + E+    S    G
Sbjct  61   EDEEDVQMEQVMTDSGFREG  80
```

>unnamed protein product [Homo sapiens]
Sequence ID: BAB15408.1 Length: 300
Range 1: 39 to 295

Score:47.4 bits(111), Expect:1e-04,
Method:Compositional matrix adjust.,
Identities:59/262(23%), Positives:96/262(36%), Gaps:81/262(30%)

```
Query  28   GLIHKLSIKNDRRLAFRIETFGFFTGGVMEMAIENFKVVDDKGSLWDDLSAGFIIKHIET  87
            G +H L++K+D R    + TFGFF  G M + + +  + + +     D++ GF +   +
Sbjct  39   GRVHHLALKDDVRHKVHLNTFGFFKDGYMVVNVSSLSLNEPEDK---DVTIGFSLDRTKN  95

Query  88   DSGSFIEETDASKC--------VSLL----------------------------DEPERG  111
            D  S   + D + C        V+LL                            DE   G
Sbjct  96   DGFSSYLDEDVNYCILKKQSVSVTLLILDISRSEVRVKSPPEAGTQLPKIIFSRDEKVLG  155

Query  112  DTIAVKVTKPSKENEKQ-------SKVITIDKT---------------------------  137
             +    V   S  N+ Q       SK  T+D
Sbjct  156  QSQEPNVNPASAGNQTQKTQDGGKSKRSTVDSKAMGEKSFSVHNNGGAVSFQFFFNISTD  215

Query  138  RPEGFYTVVFLNCQ----PGTYVSF--DLTLTNYNPGPNYLSAGLTALPTLYAMLFVVWT  191
              EG Y++ F  C      P    +F  D+ +T  +P  +YLSAG   LP LY  +   +
Sbjct  216  DQEGLYSLYFHKCLGKELPSDKFTFSLDIEITEKDPD-SYLSAGEIPLPKLYISMAFFFF  274

Query  192  VILGVWLFHFMRGQGKRIFRIH  213
            +   +W+ H +R +   +F+IH
Sbjct  275  LSGTIWI-HILRKRRNDVFKIH  295
```
