## Supplemental Data 2 for "Overexpression of a G-protein coupled receptor-like gene affects encystment of *Acanthamoeba castellanii*"

RID: M0G1A0XE014
Job Title:Protein Sequence
Program: BLASTP
Query: None ID: lcl|Query_5642(amino acid) Length: 456
Database: nr All non-redundant GenBank CDS translations+PDB+SwissProt+PIR+PRF excluding environmental samples from WGS projects

Sequences producing significant alignments:

```
                                                                    Max    Total
Query   E   Per.
Description                                                         Score  Score
cover Value Ident  Accession
mKIAA1624 protein [Mus musculus]                                    305    305
90%   6e-98 36.47  BAC98217.1
unnamed protein product [Mus musculus]                              305    305
87%   2e-97 36.89  BAC38445.1
protein GPR107 precursor [Mus musculus]                             305    305
87%   2e-97 36.89  NP_848875.2
unnamed protein product [Mus musculus]                              305    305
87%   2e-97 36.89  BAC40414.1
unnamed protein product [Mus musculus]                              305    305
87%   2e-97 36.89  BAC28840.1
protein GPR108 isoform 3 [Mus musculus]                             250    250
67%   4e-78 42.24  NP_001295003.1
protein GPR108 isoform 2 precursor [Mus musculus]                   251    285
75%   2e-76 42.24  NP_001295001.1
G protein-coupled receptor 108, isoform CRA_c [Mus musculus]        250    284
75%   3e-76 42.24  EDL38250.1
protein GPR108 isoform 1 precursor [Mus musculus]                   250    284
75%   3e-76 42.24  NP_084360.2
G protein-coupled receptor 108, isoform CRA_e [Mus musculus]        251    285
80%   4e-76 42.24  EDL38252.1
lung seven transmembrane receptor 2 [Mus musculus]                  249    283
75%   1e-75 41.93  AAK57696.1
unnamed protein product [Mus musculus]                              248    282
75%   1e-75 41.93  BAC33036.1
unnamed protein product [Mus musculus]                              207    207
69%   5e-61 33.09  BAE25070.1
unnamed protein product [Mus musculus]                              77.8   77.8
17%   7e-17 47.50  BAB31509.2
G protein-coupled receptor 108, isoform CRA_b [Mus musculus]        66.6   66.6
11%   2e-12 51.85  EDL38249.1
```

Alignments:
>mKIAA1624 protein, partial [Mus musculus]
Sequence ID: BAC98217.1 Length: 530
Range 1: 1 to 498

Score:305 bits(782), Expect:6e-98,
Method:Compositional matrix adjust.,
Identities:182/499(36%), Positives:262/499(52%), Gaps:88/499(17%)

```
Query  17   CLVAFLAPTANGLIHKLSIKNDRRLAFRIETFGFFTGGVMEMAIENFKVVDDKGSLWDDL  76
             L+  L     G +H L++K+D R    + TFGFF  G M + + +  V + +G+   D
```

```
Sbjct  1    ALLELLVHPVLGRVHHLALKDDVRHKVHLNTFGFFKDGYMVVNVSSLSVNEPEGATDKDA  60

Query  77   SAGFIIKHIETDSGSFIEETDASKC---------------------VSLLDEPERG----  111
              GF +   + D  S   + D + C                     V +   PE G
Sbjct  61   EIGFSLDRTKNDGFSSYLDEDVNYCILKKKSMSSVTLVILDISGSIVKVRSPPEAGKQLP  120

Query  112  -------DTIAVKVTKPS-----KENEKQ--------------SKVIT------------  133
                   + I  +  +P+     K++E +              SK IT
Sbjct  121  EIVFSKDEKILSRSQEPAVSSNPKDSEARRTLDGFKAGRSTVDSKAITERSFSIHKNDGV  180

Query  134  --------IDKTRPEGFYTVVFLNC-----QPGTYVSFDLTLTNYNPGPN-YLSAGLTAL  179
                    I     EG Y++ F  C     +PG   SF L +      PN YLSAG   L
Sbjct  181  VSFQFFFNISTDDQEGLYSLYFHKCSGNNVKPGEQASFSLNIAITEKNPNSYLSAGEIPL  240

Query  180  PTLYAMLFVVWTVILGVWLFHFMRGQGKRIFRIHHLVTGIILLKLLTLLFEAIEFHYKKT  239
            P LY  + + + +   +W+ H +R +   +F+IH L+  +   K L+L+F AI++HY  +
Sbjct  241  PKLYVSMALFFFLSGTIWI-HILRKRRNDVFKIHWLMAALPFTKSLSLVFHAIDYHYISS  299

Query  240  TGHP-GGWVIAYYIFSGLKGTMMFVVIALIGTGWAFIKPFLGEKDKNIFLVVIPLQILAN  298
             G P  GW + YYI   LKG ++F+ IALIGTGWAFIK  L +KDK IF++VIPLQ+LAN
Sbjct  300  QGFPIEGWAVVYYITHLLKGALLFITIALIGTGWAFIKHILSDKDKKIFMIVIPLQVLAN  359

Query  299  IAIIVLEETA---------PEVLRLVDIICCGAILVPIIWSIKHLRDAAAIDGKAKRNME  349
            +A I++E T           + L LVD++CCGAIL P++WSI+HL++A+A DGKA  N+
Sbjct  360  VAYIIIESTEEGTTEYGLWKDSLFLVDLLCCGAILFPVVWSIRHLQEASATDGKAAINLA  419

Query  350  KLKLFREFYLLVVTYIYFTRIIVFLLDATLHYQYVWLGEFFTELATLIFWGLTGYKFRPV  409
            KL+LFR +Y+L+V YIYFTRII FLL   + +Q+ WL +   E ATL+F+ LTGYKFRP
Sbjct  420  KLRLFRHYYVLIVCYIYFTRIIAFLLKFAVPFQWKWLYQLLDETATLVFFVLTGYKFRPA  479

Query  410  ADNPYLKLDDEEEDAEREE  428
            +DNPYL+L  E++D E E
Sbjct  480  SDNPYLQLSQEDDDLEMES  498



>unnamed protein product [Mus musculus]
Sequence ID: BAC38445.1 Length: 551
Range 1: 33 to 519

Score:305 bits(781), Expect:2e-97,
Method:Compositional matrix adjust.,
Identities:180/488(37%), Positives:259/488(53%), Gaps:88/488(18%)

Query  28   GLIHKLSIKNDRRLAFRIETFGFFTGGVMEMAIENFKVVDDKGSLWDDLSAGFIIKHIET  87
            G +H L++K+D R    + TFGFF  G M + + +  V + +G+   D   GF +   +
Sbjct  33   GRVHHLALKDDVRHKVHLNTFGFFKDGYMVVNVSSLSVNEPEGATDKDAEIGFSLDRTKN  92

Query  88   DSGSFIEETDASKC---------------------VSLLDEPERG-----------DTIA  115
            D  S   + D + C                     V +   PE G           + I
Sbjct  93   DGFSSYLDEDVNYCILKKKSMSSVTLVILDISGSLVKVRSPPEAGKQLPEIVFSKDEKIL  152

Query  116  VKVTKPS-----KENEKQ--------------SKVIT--------------------IDK  136
             +  +P+     K++E +              SK IT                    I
```

```
Sbjct  153  SQSQEPAVSSNPKDSEARRTLDGFKAGRSTVDSKAITERSFSIHKNDGVVSFQFFFNIST  212

Query  137  TRPEGFYTVVFLNC-----QPGTYVSFDLTLTNYNPGPN-YLSAGLTALPTLYAMLFVVW  190
               EG Y++ F  C     +PG   SF L +      PN YLSAG   LP LY  + + +
Sbjct  213  DDQEGLYSLYFHKCSGNNVKPGEQASFSLNIAITEKNPNSYLSAGEIPLPKLYVSMALFF  272

Query  191  TVILGVWLFHFMRGQGKRIFRIHHLVTGIILLKLLTLLFEAIEFHYKKTTGHP-GGWVIA  249
             +   +W+ H +R +   +F+IH L+  +   K L+L+F AI++HY  + G P  GW +
Sbjct  273  FLSGTIWI-HILRKRRNDVFKIHWLMAALPFTKSLSLVFHAIDYHYISSQGFPIEGWAVV  331

Query  250  YYIFSGLKGTMMFVVIALIGTGWAFIKPFLGEKDKNIFLVVIPLQILANIAIIVLEETA-  308
            YYI   LKG ++F+ IALIGTGWAFIK  L +KDK IF++VIPLQ+LAN+A I++E T
Sbjct  332  YYITHLLKGALLFITIALIGTGWAFIKHILSDKDKKIFMIVIPLQVLANVAYIIIESTEE  391

Query  309  --------PEVLRLVDIICCGAILVPIIWSIKHLRDAAAIDGKAKRNMEKLKLFREFYLL  360
                     + L LVD++CCGAIL P++WSI+HL++A+A DGKA  N+ KL+LFR +Y+L
Sbjct  392  GTTEYGLWKDSLFLVDLLCCGAILFPVVWSIRHLQEASATDGKAAINLAKLRLFRHYYVL  451

Query  361  VVTYIYFTRIIVFLLDATLHYQYVWLGEFFTELATLIFWGLTGYKFRPVADNPYLKLDDE  420
            +V YIYFTRII FLL   + +Q+ WL +   E ATL+F+ LTGYKFRP +DNPYL+L  E
Sbjct  452  IVCYIYFTRIIAFLLKFAVPFQWKWLYQLLDETATLVFFVLTGYKFRPASDNPYLQLSQE  511

Query  421  EEDAEREE  428
            ++D E E
Sbjct  512  DDDLEMES  519
```

>protein GPR107 precursor [Mus musculus]
Sequence ID: NP_848875.2 Length: 551
>RecName: Full=Protein GPR107; Flags: Precursor [Mus musculus]
Sequence ID: Q8BUV8.2 Length: 551 >G protein-coupled receptor 107 [Mus musculus]
Sequence ID: AAH92231.1 Length: 551 >G protein-coupled receptor 107 [Mus musculus]
Sequence ID: AAI17901.1 Length: 551 >G protein-coupled receptor 107 [Mus musculus]
Sequence ID: AAI17902.1 Length: 551 >expressed sequence AI790205 [Mus musculus]
Sequence ID: EDL08503.1 Length: 551 >unnamed protein product [Mus musculus]
Sequence ID: BAC26961.1 Length: 551
Range 1: 33 to 519

Score:305 bits(781), Expect:2e-97,
Method:Compositional matrix adjust.,
Identities:180/488(37%), Positives:259/488(53%), Gaps:88/488(18%)

```
Query  28   GLIHKLSIKNDRRLAFRIETFGFFTGGVMEMAIENFKVVDDKGSLWDDLSAGFIIKHIET  87
            G +H L++K+D R    + TFGFF  G M + + +  V + +G+   D   GF +   +
Sbjct  33   GRVHHLALKDDVRHKVHLNTFGFFKDGYMVVNVSSLSVNEPEGATDKDAEIGFSLDRTKN  92

Query  88   DSGSFIEETDASKC---------------------VSLLDEPERG-----------DTIA  115
            D  S   + D + C                     V +   PE G           + I
Sbjct  93   DGFSSYLDEDVNYCILKKKSMSSVTLVILDISGSIVKVRSPPEAGKQLPEIVFSKDEKIL  152

Query  116  VKVTKPS-----KENEKQ--------------SKVIT--------------------IDK  136
```

```
                +  +P+     K++E +              SK IT                    I
Sbjct  153  SQSQEPAVSSNPKDSEARRTLDGFKAGRSTVDSKAITERSFSIHKNDGVVSFQFFFNIST  212

Query  137  TRPEGFYTVVFLNC-----QPGTYVSFDLTLTNYNPGPN-YLSAGLTALPTLYAMLFVVW  190
               EG Y++ F  C     +PG   SF L +      PN YLSAG   LP LY  + + +
Sbjct  213  DDQEGLYSLYFHKCSGNNVKPGEQASFSLNIAITEKNPNSYLSAGEIPLPKLYVSMALFF  272

Query  191  TVILGVWLFHFMRGQGKRIFRIHHLVTGIILLKLLTLLFEAIEFHYKKTTGHP-GGWVIA  249
             +   +W+ H +R +   +F+IH L+  +   K L+L+F AI++HY  + G P  GW +
Sbjct  273  FLSGTIWI-HILRKRRNDVFKIHWLMAALPFTKSLSLVFHAIDYHYISSQGFPIEGWAVV  331

Query  250  YYIFSGLKGTMMFVVIALIGTGWAFIKPFLGEKDKNIFLVVIPLQILANIAIIVLEETA-  308
            YYI   LKG ++F+ IALIGTGWAFIK  L +KDK IF++VIPLQ+LAN+A I++E T
Sbjct  332  YYITHLLKGALLFITIALIGTGWAFIKHILSDKDKKIFMIVIPLQVLANVAYIIIESTEE  391

Query  309  --------PEVLRLVDIICCGAILVPIIWSIKHLRDAAAIDGKAKRNMEKLKLFREFYLL  360
                     + L LVD++CCGAIL P++WSI+HL++A+A DGKA  N+ KL+LFR +Y+L
Sbjct  392  GTTEYGLWKDSLFLVDLLCCGAILFPVVWSIRHLQEASATDGKAAINLAKLRLFRHYYVL  451

Query  361  VVTYIYFTRIIVFLLDATLHYQYVWLGEFFTELATLIFWGLTGYKFRPVADNPYLKLDDE  420
            +V YIYFTRII FLL   + +Q+ WL +   E ATL+F+ LTGYKFRP +DNPYL+L  E
Sbjct  452  IVCYIYFTRIIAFLLKFAVPFQWKWLYQLLDETATLVFFVLTGYKFRPASDNPYLQLSQE  511

Query  421  EEDAEREE  428
            ++D E E
Sbjct  512  DDDLEMES  519
```

>unnamed protein product [Mus musculus]
Sequence ID: BAC40414.1 Length: 551
>unnamed protein product [Mus musculus]
Sequence ID: BAE32655.1 Length: 551
Range 1: 33 to 519

Score:305 bits(781), Expect:2e-97,
Method:Compositional matrix adjust.,
Identities:180/488(37%), Positives:259/488(53%), Gaps:88/488(18%)

```
Query  28   GLIHKLSIKNDRRLAFRIETFGFFTGGVMEMAIENFKVVDDKGSLWDDLSAGFIIKHIET  87
            G +H L++K+D R    + TFGFF  G M + + +  V + +G+   D   GF +   +
Sbjct  33   GRVHHLALKDDVRHKVHLNTFGFFKDGYMVVNVSSLSVNEPEGATDKDAEIGFSLDRTKN  92

Query  88   DSGSFIEETDASKC---------------------VSLLDEPERG-----------DTIA  115
            D  S   + D + C                     V +   PE G           + I
Sbjct  93   DGFSSYLDEDVNYCILKKKSMSSVTLVILDISGSIVKVRSPPEAGKQLPEIVFSKDEKIL  152

Query  116  VKVTKPS-----KENEKQ--------------SKVIT--------------------IDK  136
             +  +P+     K++E +              SK IT                    I
Sbjct  153  SRSQEPAVSSNPKDSEARRTLDGFKAGRSTVDSKAITERSFSIHKNDGVVSFQFFFNIST  212

Query  137  TRPEGFYTVVFLNC-----QPGTYVSFDLTLTNYNPGPN-YLSAGLTALPTLYAMLFVVW  190
               EG Y++ F  C     +PG   SF L +      PN YLSAG   LP LY  + + +
Sbjct  213  DDQEGLYSLYFHKCSGNNVKPGEQASFSLNIAITEKNPNSYLSAGEIPLPKLYVSMALFF  272
```

```
Query  191  TVILGVWLFHFMRGQGKRIFRIHHLVTGIILLKLLTLLFEAIEFHYKKTTGHP-GGWVIA  249
             +   +W+ H +R +   +F+IH L+  +   K L+L+F AI++HY  + G P  GW +
Sbjct  273  FLSGTIWI-HILRKRRNDVFKIHWLMAALPFTKSLSLVFHAIDYHYISSQGFPIEGWAVV  331

Query  250  YYIFSGLKGTMMFVVIALIGTGWAFIKPFLGEKDKNIFLVVIPLQILANIAIIVLEETA-  308
            YYI   LKG ++F+ IALIGTGWAFIK  L +KDK IF++VIPLQ+LAN+A I++E T
Sbjct  332  YYITHLLKGALLFITIALIGTGWAFIKHILSDKDKKIFMIVIPLQVLANVAYIIIESTEE  391

Query  309  --------PEVLRLVDIICCGAILVPIIWSIKHLRDAAAIDGKAKRNMEKLKLFREFYLL  360
                     + L LVD++CCGAIL P++WSI+HL++A+A DGKA  N+ KL+LFR +Y+L
Sbjct  392  GTTEYGLWKDSLFLVDLLCCGAILFPVVWSIRHLQEASATDGKAAINLAKLRLFRHYYVL  451

Query  361  VVTYIYFTRIIVFLLDATLHYQYVWLGEFFTELATLIFWGLTGYKFRPVADNPYLKLDDE  420
            +V YIYFTRII FLL   + +Q+ WL +   E ATL+F+ LTGYKFRP +DNPYL+L  E
Sbjct  452  IVCYIYFTRIIAFLLKFAVPFQWKWLYQLLDETATLVFFVLTGYKFRPASDNPYLQLSQE  511

Query  421  EEDAEREE  428
            ++D E E
Sbjct  512  DDDLEMES  519
```

>unnamed protein product [Mus musculus]
Sequence ID: BAC28840.1 Length: 551
Range 1: 33 to 519

Score:305 bits(780), Expect:2e-97,
Method:Compositional matrix adjust.,
Identities:180/488(37%), Positives:259/488(53%), Gaps:88/488(18%)

```
Query  28   GLIHKLSIKNDRRLAFRIETFGFFTGGVMEMAIENFKVVDDKGSLWDDLSAGFIIKHIET  87
            G +H L++K+D R    + TFGFF  G M + + +  V + +G+   D   GF +    +
Sbjct  33   GRVHHLALKDDVRHKVHLNTFGFFKDGYMVVNVSSLSVNEPEGATDKDAEIGFSLDRTKN  92

Query  88   DSGSFIEETDASKC---------------------VSLLDEPERG-----------DTIA  115
            D  S   + D + C                     V +   PE G           + I
Sbjct  93   DGFSSYLDEDVNYCILKKKSMSSVTLVILDISGSIVKVRSPPEAGKQLPEIVFSKDEKIL  152

Query  116  VKVTKPS-----KENEKQ--------------SKVIT--------------------IDK  136
             +  +P+     K++E +              SK IT                    I
Sbjct  153  SQSQEPAVSSNPKDSEARRTLDGFKAGRSTVDSKAITERSFSIHKNDGVVSFQFFFNIST  212

Query  137  TRPEGFYTVVFLNC-----QPGTYVSFDLTLTNYNPGPN-YLSAGLTALPTLYAMLFVVW  190
               EG Y++ F  C     +PG   SF L +      PN YLSAG   LP LY  + + +
Sbjct  213  DDQEGLYSLYFHKCSGNNVKPGEQASFSLNIAITEKNPNSYLSAGEIPLPKLYVSMALFF  272

Query  191  TVILGVWLFHFMRGQGKRIFRIHHLVTGIILLKLLTLLFEAIEFHYKKTTGHP-GGWVIA  249
             +   +W+ H +R +   +F+IH L+  +   K L+L+F AI++HY  + G P  GW +
Sbjct  273  FLSGTIWI-HILRKRRNDVFKIHWLMAALPFTKSLSLVFHAIDYHYISSQGFPIEGWAVV  331

Query  250  YYIFSGLKGTMMFVVIALIGTGWAFIKPFLGEKDKNIFLVVIPLQILANIAIIVLEETA-  308
            YYI   LKG ++F+ IALIGTGWAFIK  L +KDK IF++VIPLQ+LAN+A I++E T
Sbjct  332  YYITHLLKGALLFITIALIGTGWAFIKHILSDKDKKIFMIVIPLQVLANVAYIIIESTEE  391
```

```
Query  309  --------PEVLRLVDIICCGAILVPIIWSIKHLRDAAAIDGKAKRNMEKLKLFREFYLL  360
                     + L LVD++CCGAIL P++WSI+HL++A+A DGKA  N+ KL+LFR +Y+L
Sbjct  392  GTTEYGLWKDSLFLVDLLCCGAILFPVVWSIRHLQEASATDGKAAINLAKLRLFRHYYVL  451

Query  361  VVTYIYFTRIIVFLLDATLHYQYVWLGEFFTELATLIFWGLTGYKFRPVADNPYLKLDDE  420
            +V YIYFTRII FLL   + +Q+ WL +   E ATL+F+ LTGYKFRP +DNPYL+L  E
Sbjct  452  IVCYIYFTRIIAFLLKFAVPFQWKWLYQLLDETATLVFFVLTGYKFRPASDNPYLQLSQE  511

Query  421  EEDAEREE  428
            ++D E E
Sbjct  512  DDDLEMES  519
```

>protein GPR108 isoform 3 [Mus musculus]
Sequence ID: NP_001295003.1 Length: 394
>protein GPR108 isoform X1 [Mus musculus]
Sequence ID: XP_011245021.1 Length: 394 >protein GPR108 isoform X1 [Mus musculus]
Sequence ID: XP_017173215.1 Length: 394 >protein GPR108 isoform X1 [Mus musculus]
Sequence ID: XP_017173216.1 Length: 394
Range 1: 60 to 380

Score:250 bits(639), Expect:4e-78,
Method:Compositional matrix adjust.,
Identities:136/322(42%), Positives:192/322(59%), Gaps:16/322(4%)

```
Query  132  ITIDKTRPEGFYTVVFLNCQ---PGTYVSFDLTLTNYNPGPN-YLSAGLTALPTLYAMLF  187
            I I     EG Y++ F NC    PG    FDLT+      P  +LSA    L  LY ++
Sbjct  60   IVISSRAEEGQYSLNFHNCHNSIPGQEQPFDLTVMIREKNPEGFLSAAEIPLFKLYLIMS  119

Query  188  VVWTVILGVWLFHFMRGQGKRIFRIHHLVTGIILLKLLTLLFEAIEFHYKKTTGHP-GGW  246
              +      W+    +     +F+IH L+  +   K ++LLF +I +++  + GHP  G
Sbjct  120  ACFLAADIFWVSVLCKNT-YSVFKIHWLMAALAFTKSVSLLFHSINYYFINSQGHPIEGL  178

Query  247  VIAYYIFSGLKGTMMFVVIALIGTGWAFIKPFLGEKDKNIFLVVIPLQILANIAIIVLEE  306
             + +YI   LKG ++F+ IALIG+GWAF+K  L +K+K IF +VIPLQ+LAN+A IV+E
Sbjct  179  AVMHYITHLLKGALLFITIALIGSGWAFVKYMLSDKEKKIFGIVIPLQVLANVAYIVIES  238

Query  307  TA---------PEVLRLVDIICCGAILVPIIWSIKHLRDAAAIDGKAKRNMEKLKLFREF  357
                        E+L LVD+ICCGAIL P++WSI+HL+DA+  DGK   N+ +LKLFR +
Sbjct  239  REEGASDYGLWKEILFLVDLICCGAILFPVVWSIRHLQDASGTDGKVAVNLARLKLFRHY  298

Query  358  YLLVVTYIYFTRIIVFLLDATLHYQYVWLGEFFTELATLIFWGLTGYKFRPVADNPYLKL  417
            Y++V+ YIYFTRII  LL   + +Q+ WL +   E +TL F+ LTGYKF+P  DNPYL+L
Sbjct  299  YVMVICYIYFTRIIAILLQVAVPFQWQWLYQLLVESSTLAFFVLTGYKFQPAGDNPYLQL  358

Query  418  DDE-EEDAEREEAQRQSRTEAG  438
              E EED + E+    S    G
Sbjct  359  PQEDEEDVQMEQVMTDSGFREG  380
```

>protein GPR108 isoform 2 precursor [Mus musculus]
Sequence ID: NP_001295001.1 Length: 562
>G protein-coupled receptor 108, isoform CRA_d [Mus musculus]
Sequence ID: EDL38251.1 Length: 562
Range 1: 228 to 548

Score:251 bits(640), Expect:2e-76,
Method:Compositional matrix adjust.,
Identities:136/322(42%), Positives:192/322(59%), Gaps:16/322(4%)

```
Query  132  ITIDKTRPEGFYTVVFLNCQ---PGTYVSFDLTLTNYNPGP-NYLSAGLTALPTLYAMLF  187
            I I     EG Y++ F NC    PG    FDLT+      P  +LSA    L  LY ++
Sbjct  228  IVISSRAEEGQYSLNFHNCHNSIPGQEQPFDLTVMIREKNPEGFLSAAEIPLFKLYLIMS  287

Query  188  VVWTVILGVWLFHFMRGQGKRIFRIHHLVTGIILLKLLTLLFEAIEFHYKKTTGHP-GGW  246
              +      W+    +     +F+IH L+  +   K ++LLF +I +++  + GHP  G
Sbjct  288  ACFLAADIFWVSVLCKNT-YSVFKIHWLMAALAFTKSVSLLFHSINYYFINSQGHPIEGL  346

Query  247  VIAYYIFSGLKGTMMFVVIALIGTGWAFIKPFLGEKDKNIFLVVIPLQILANIAIIVLEE  306
             + +YI   LKG ++F+ IALIG+GWAF+K  L +K+K IF +VIPLQ+LAN+A IV+E
Sbjct  347  AVMHYITHLLKGALLFITIALIGSGWAFVKYMLSDKEKKIFGIVIPLQVLANVAYIVIES  406

Query  307  TA---------PEVLRLVDIICCGAILVPIIWSIKHLRDAAAIDGKAKRNMEKLKLFREF  357
                        E+L LVD+ICCGAIL P++WSI+HL+DA+  DGK   N+ +LKLFR +
Sbjct  407  REEGASDYGLWKEILFLVDLICCGAILFPVVWSIRHLQDASGTDGKVAVNLARLKLFRHY  466

Query  358  YLLVVTYIYFTRIIVFLLDATLHYQYVWLGEFFTELATLIFWGLTGYKFRPVADNPYLKL  417
            Y++V+ YIYFTRII  LL    + +Q+ WL +   E +TL F+ LTGYKF+P  DNPYL+L
Sbjct  467  YVMVICYIYFTRIIAILLQVAVPFQWQWLYQLLVESSTLAFFVLTGYKFQPAGDNPYLQL  526

Query  418  DDE-EEDAEREEAQRQSRTEAG  438
              E EED + E+    S    G
Sbjct  527  PQEDEEDVQMEQVMTDSGFREG  548
```

Range 2: 32 to 67

Score:33.9 bits(76), Expect:1.4,
Method:Compositional matrix adjust.,
Identities:11/36(31%), Positives:24/36(66%), Gaps:0/36(0%)

```
Query  26  ANGLIHKLSIKNDRRLAFRIETFGFFTGGVMEMAIE  61
            +G IH+L++  ++R   ++ +FGF+T G +E+ +
Sbjct  32  CSGRIHRLTLTGEKRADIQLNSFGFYTNGSLEVELS  67
```

>G protein-coupled receptor 108, isoform CRA_c, partial [Mus musculus]
Sequence ID: EDL38250.1 Length: 570
Range 1: 236 to 556

Score:250 bits(639), Expect:3e-76,
Method:Compositional matrix adjust.,
Identities:136/322(42%), Positives:192/322(59%), Gaps:16/322(4%)

```
Query  132  ITIDKTRPEGFYTVVFLNCQ---PGTYVSFDLTLTNYNPGP-NYLSAGLTALPTLYAMLF  187
            I I     EG Y++ F NC    PG    FDLT+      P  +LSA    L  LY ++
Sbjct  236  IVISSRAEEGQYSLNFHNCHNSIPGQEQPFDLTVMIREKNPEGFLSAAEIPLFKLYLIMS  295

Query  188  VVWTVILGVWLFHFMRGQGKRIFRIHHLVTGIILLKLLTLLFEAIEFHYKKTTGHP-GGW  246
              +      W+    +     +F+IH L+  +   K ++LLF +I +++  + GHP  G
Sbjct  296  ACFLAADIFWVSVLCKNT-YSVFKIHWLMAALAFTKSVSLLFHSINYYFINSQGHPIEGL  354

Query  247  VIAYYIFSGLKGTMMFVVIALIGTGWAFIKPFLGEKDKNIFLVVIPLQILANIAIIVLEE  306
             + +YI   LKG ++F+ IALIG+GWAF+K  L +K+K IF +VIPLQ+LAN+A IV+E
Sbjct  355  AVMHYITHLLKGALLFITIALIGSGWAFVKYMLSDKEKKIFGIVIPLQVLANVAYIVIES  414

Query  307  TA---------PEVLRLVDIICCGAILVPIIWSIKHLRDAAAIDGKAKRNMEKLKLFREF  357
                        E+L LVD+ICCGAIL P++WSI+HL+DA+  DGK   N+ +LKLFR +
Sbjct  415  REEGASDYGLWKEILFLVDLICCGAILFPVVWSIRHLQDASGTDGKVAVNLARLKLFRHY  474

Query  358  YLLVVTYIYFTRIIVFLLDATLHYQYVWLGEFFTELATLIFWGLTGYKFRPVADNPYLKL  417
            Y++V+ YIYFTRII  LL   + +Q+ WL +   E +TL F+ LTGYKF+P  DNPYL+L
Sbjct  475  YVMVICYIYFTRIIAILLQVAVPFQWQWLYQLLVESSTLAFFVLTGYKFQPAGDNPYLQL  534

Query  418  DDE-EEDAEREEAQRQSRTEAG  438
              E EED + E+    S    G
Sbjct  535  PQEDEEDVQMEQVMTDSGFREG  556
```

Range 2: 33 to 67

Score:33.9 bits(76), Expect:1.3,
Method:Compositional matrix adjust.,
Identities:11/35(31%), Positives:24/35(68%), Gaps:0/35(0%)

```
Query  26  ANGLIHKLSIKNDRRLAFRIETFGFFTGGVMEMAI  60
            +G IH+L++  ++R   ++ +FGF+T G +E+ +
Sbjct  33  CSGRIHRLTLTGEKRADIQLNSFGFYTNGSLEVEL  67
```

>protein GPR108 isoform 1 precursor [Mus musculus]
Sequence ID: NP_084360.2 Length: 569
>RecName: Full=Protein GPR108; AltName: Full=Lung seven transmembrane receptor 2; Flags: Precursor [Mus musculus]
Sequence ID: Q91WD0.1 Length: 569 >G protein-coupled receptor 108 [Mus musculus]
Sequence ID: AAH16104.1 Length: 569 >unnamed protein product [Mus musculus]
Sequence ID: BAE34981.1 Length: 569
Range 1: 235 to 555

Score:250 bits(639), Expect:3e-76,
Method:Compositional matrix adjust.,

Identities:136/322(42%), Positives:192/322(59%), Gaps:16/322(4%)

```
Query  132  ITIDKTRPEGFYTVVFLNCQ---PGTYVSFDLTLTNYNPGP-NYLSAGLTALPTLYAMLF  187
            I I     EG Y++ F NC    PG    FDLT+      P  +LSA    L  LY ++
Sbjct  235  IVISSRAEEGQYSLNFHNCHNSIPGQEQPFDLTVMIREKNPEGFLSAAEIPLFKLYLIMS  294

Query  188  VVWTVILGVWLFHFMRGQGKRIFRIHHLVTGIILLKLLTLLFEAIEFHYKKTTGHP-GGW  246
              +      W+    +     +F+IH L+  +   K ++LLF +I +++  + GHP  G
Sbjct  295  ACFLAADIFWVSVLCKNT-YSVFKIHWLMAALAFTKSVSLLFHSINYYFINSQGHPIEGL  353

Query  247  VIAYYIFSGLKGTMMFVVIALIGTGWAFIKPFLGEKDKNIFLVVIPLQILANIAIIVLEE  306
             + +YI   LKG ++F+ IALIG+GWAF+K  L +K+K IF +VIPLQ+LAN+A IV+E
Sbjct  354  AVMHYITHLLKGALLFITIALIGSGWAFVKYMLSDKEKKIFGIVIPLQVLANVAYIVIES  413

Query  307  TA---------PEVLRLVDIICCGAILVPIIWSIKHLRDAAAIDGKAKRNMEKLKLFREF  357
                        E+L LVD+ICCGAIL P++WSI+HL+DA+  DGK   N+ +LKLFR +
Sbjct  414  REEGASDYGLWKEILFLVDLICCGAILFPVVWSIRHLQDASGTDGKVAVNLARLKLFRHY  473

Query  358  YLLVVTYIYFTRIIVFLLDATLHYQYVWLGEFFTELATLIFWGLTGYKFRPVADNPYLKL  417
            Y++V+ YIYFTRII  LL   + +Q+ WL +   E +TL F+ LTGYKF+P  DNPYL+L
Sbjct  474  YVMVICYIYFTRIIAILLQVAVPFQWQWLYQLLVESSTLAFFVLTGYKFQPAGDNPYLQL  533

Query  418  DDE-EEDAEREEAQRQSRTEAG  438
              E EED + E+    S    G
Sbjct  534  PQEDEEDVQMEQVMTDSGFREG  555
```

Range 2: 32 to 67

Score:33.9 bits(76), Expect:1.3,
Method:Compositional matrix adjust.,
Identities:11/36(31%), Positives:24/36(66%), Gaps:0/36(0%)

```
Query  26  ANGLIHKLSIKNDRRLAFRIETFGFFTGGVMEMAIE  61
            +G IH+L++  ++R   ++ +FGF+T G +E+ +
Sbjct  32  CSGRIHRLTLTGEKRADIQLNSFGFYTNGSLEVELS  67
```

>G protein-coupled receptor 108, isoform CRA_e [Mus musculus]
Sequence ID: EDL38252.1 Length: 598
Range 1: 228 to 548

Score:251 bits(640), Expect:4e-76,
Method:Compositional matrix adjust.,
Identities:136/322(42%), Positives:192/322(59%), Gaps:16/322(4%)

```
Query  132  ITIDKTRPEGFYTVVFLNCQ---PGTYVSFDLTLTNYNPGP-NYLSAGLTALPTLYAMLF  187
            I I     EG Y++ F NC    PG    FDLT+      P  +LSA    L  LY ++
Sbjct  228  IVISSRAEEGQYSLNFHNCHNSIPGQEQPFDLTVMIREKNPEGFLSAAEIPLFKLYLIMS  287

Query  188  VVWTVILGVWLFHFMRGQGKRIFRIHHLVTGIILLKLLTLLFEAIEFHYKKTTGHP-GGW  246
```

```
              +      W+    +     +F+IH L+  +   K ++LLF +I +++  + GHP  G
Sbjct  288  ACFLAADIFWVSVLCKNT-YSVFKIHWLMAALAFTKSVSLLFHSINYYFINSQGHPIEGL  346

Query  247  VIAYYIFSGLKGTMMFVVIALIGTGWAFIKPFLGEKDKNIFLVVIPLQILANIAIIVLEE  306
             + +YI   LKG ++F+ IALIG+GWAF+K  L +K+K IF +VIPLQ+LAN+A IV+E
Sbjct  347  AVMHYITHLLKGALLFITIALIGSGWAFVKYMLSDKEKKIFGIVIPLQVLANVAYIVIES  406

Query  307  TA---------PEVLRLVDIICCGAILVPIIWSIKHLRDAAAIDGKAKRNMEKLKLFREF  357
                        E+L LVD+ICCGAIL P++WSI+HL+DA+  DGK   N+ +LKLFR +
Sbjct  407  REEGASDYGLWKEILFLVDLICCGAILFPVVWSIRHLQDASGTDGKVAVNLARLKLFRHY  466

Query  358  YLLVVTYIYFTRIIVFLLDATLHYQYVWLGEFFTELATLIFWGLTGYKFRPVADNPYLKL  417
            Y++V+ YIYFTRII  LL   + +Q+ WL +   E +TL F+ LTGYKF+P  DNPYL+L
Sbjct  467  YVMVICYIYFTRIIAILLQVAVPFQWQWLYQLLVESSTLAFFVLTGYKFQPAGDNPYLQL  526

Query  418  DDE-EEDAEREEAQRQSRTEAG  438
              E EED + E+    S    G
Sbjct  527  PQEDEEDVQMEQVMTDSGFREG  548
```

Range 2: 1 to 67

Score:34.3 bits(77), Expect:1.2,
Method:Compositional matrix adjust.,
Identities:17/67(25%), Positives:33/67(49%), Gaps:6/67(8%)

```
Query  1   MMRDRRQTLMGCFLIFCLVAFLA------PTANGLIHKLSIKNDRRLAFRIETFGFFTGG  54
           M    R+ L G     C   +L+         +G IH+L++  ++R   ++ +FGF+T G
Sbjct  1   MAVSERRGLSGESPTQCRWGYLSLLVLTLSGCSGRIHRLTLTGEKRADIQLNSFGFYTNG  60

Query  55  VMEMAIE  61
            +E+ +
Sbjct  61  SLEVELS  67
```

>lung seven transmembrane receptor 2 [Mus musculus]
Sequence ID: AAK57696.1 Length: 562
Range 1: 228 to 548

Score:249 bits(635), Expect:1e-75,
Method:Compositional matrix adjust.,
Identities:135/322(42%), Positives:191/322(59%), Gaps:16/322(4%)

```
Query  132  ITIDKTRPEGFYTVVFLNCQ---PGTYVSFDLTLTNYNPGP-NYLSAGLTALPTLYAMLF  187
            I I     EG Y++ F NC    PG    FDLT+      P  +LSA    L  LY ++
Sbjct  228  IVISSRAEEGQYSLNFHNCHNSIPGQEQPFDLTVMIREKNPEGFLSAAEIPLFKLYLIMS  287

Query  188  VVWTVILGVWLFHFMRGQGKRIFRIHHLVTGIILLKLLTLLFEAIEFHYKKTTGHP-GGW  246
              +      W+    +     +F+IH L+  +   K ++LLF +I +++  + GHP  G
Sbjct  288  ACFLAADIFWVSVLCKNT-YSVFKIHWLMAALAFTKSVSLLFHSINYYFINSQGHPIEGL  346
```

```
Query  247  VIAYYIFSGLKGTMMFVVIALIGTGWAFIKPFLGEKDKNIFLVVIPLQILANIAIIVLEE  306
             + +YI   LKG ++F+ IALIG+GWAF+K  L +K+K IF +VIPL +LAN+A IV+E
Sbjct  347  AVMHYITHLLKGALLFITIALIGSGWAFVKYMLSDKEKKIFGIVIPLHVLANVAYIVIES  406

Query  307  TA---------PEVLRLVDIICCGAILVPIIWSIKHLRDAAAIDGKAKRNMEKLKLFREF  357
                        E+L LVD+ICCGAIL P++WSI+HL+DA+  DGK   N+ +LKLFR +
Sbjct  407  REEGASDYGLWKEILFLVDLICCGAILFPVVWSIRHLQDASGTDGKVAVNLARLKLFRHY  466

Query  358  YLLVVTYIYFTRIIVFLLDATLHYQYVWLGEFFTELATLIFWGLTGYKFRPVADNPYLKL  417
            Y++V+ YIYFTRII  LL   + +Q+ WL +   E +TL F+ LTGYKF+P  DNPYL+L
Sbjct  467  YVMVICYIYFTRIIAILLQVAVPFQWQWLYQLLVESSTLAFFVLTGYKFQPAGDNPYLQL  526

Query  418  DDE-EEDAEREEAQRQSRTEAG  438
              E EED + E+    S    G
Sbjct  527  PQEDEEDVQMEQVMTDSGFREG  548
```

Range 2: 32 to 67

Score:33.9 bits(76), Expect:1.4,
Method:Compositional matrix adjust.,
Identities:11/36(31%), Positives:24/36(66%), Gaps:0/36(0%)

```
Query  26  ANGLIHKLSIKNDRRLAFRIETFGFFTGGVMEMAIE  61
            +G IH+L++  ++R   ++ +FGF+T G +E+ +
Sbjct  32  CSGRIHRLTLTGEKRADIQLNSFGFYTNGSLEVELS  67
```

>unnamed protein product, partial [Mus musculus]
Sequence ID: BAC33036.1 Length: 554
Range 1: 220 to 540

Score:248 bits(634), Expect:1e-75,
Method:Compositional matrix adjust.,
Identities:135/322(42%), Positives:191/322(59%), Gaps:16/322(4%)

```
Query  132  ITIDKTRPEGFYTVVFLNCQ---PGTYVSFDLTLTNYNPGP-NYLSAGLTALPTLYAMLF  187
            I I      EG Y++ F NC    PG    FDLT+       P  +L A    L  LY ++
Sbjct  220  IVISSRAEEGQYSLNFHNCHNSIPGQEQPFDLTVMIREKNPEGFLPAAEIPLFKLYLIMS  279

Query  188  VVWTVILGVWLFHFMRGQGKRIFRIHHLVTGIILLKLLTLLFEAIEFHYKKTTGHP-GGW  246
              +       W+    +     +F+IH L+  +   K ++LLF +I +++  + GHP  G
Sbjct  280  ACFLAADIFWVSVLCKNT-YSVFKIHWLMAALAFTKSVSLLFHSINYYFINSQGHPIEGL  338

Query  247  VIAYYIFSGLKGTMMFVVIALIGTGWAFIKPFLGEKDKNIFLVVIPLQILANIAIIVLEE  306
             + +YI   LKG ++F+ IALIG+GWAF+K  L +K+K IF +VIPLQ+LAN+A IV+E
Sbjct  339  AVMHYITHLLKGALLFITIALIGSGWAFVKYMLSDKEKKIFGIVIPLQVLANVAYIVIES  398

Query  307  TA---------PEVLRLVDIICCGAILVPIIWSIKHLRDAAAIDGKAKRNMEKLKLFREF  357
                        E+L LVD+ICCGAIL P++WSI+HL+DA+  DGK   N+ +LKLFR +
Sbjct  399  REEGASDYGLWKEILFLVDLICCGAILFPVVWSIRHLQDASGTDGKVAVNLARLKLFRHY  458
```

```
Query  358  YLLVVTYIYFTRIIVFLLDATLHYQYVWLGEFFTELATLIFWGLTGYKFRPVADNPYLKL  417
            Y++V+ YIYFTRII  LL   + +Q+ WL +   E +TL F+ LTGYKF+P  DNPYL+L
Sbjct  459  YVMVICYIYFTRIIAILLQVAVPFQWQWLYQLLVESSTLAFFVLTGYKFQPAGDNPYLQL  518

Query  418  DDE-EEDAEREEAQRQSRTEAG  438
              E EED + E+    S    G
Sbjct  519  PQEDEEDVQMEQVMTDSGFREG  540
```

Range 2: 24 to 59

Score:33.9 bits(76), Expect:1.3,
Method:Compositional matrix adjust.,
Identities:11/36(31%), Positives:24/36(66%), Gaps:0/36(0%)

```
Query  26  ANGLIHKLSIKNDRRLAFRIETFGFFTGGVMEMAIE  61
            +G IH+L++  ++R   ++ +FGF+T G +E+ +
Sbjct  24  CSGRIHRLTLTGEKRADIQLNSFGFYTNGSLEVELS  59
```

>unnamed protein product [Mus musculus]
Sequence ID: BAE25070.1 Length: 447
Range 1: 33 to 436

Score:207 bits(527), Expect:5e-61,
Method:Compositional matrix adjust.,
Identities:134/405(33%), Positives:198/405(48%), Gaps:88/405(21%)

```
Query  28   GLIHKLSIKNDRRLAFRIETFGFFTGGVMEMAIENFKVVDDKGSLWDDLSAGFIIKHIET  87
            G +H L++K+D R    + TFGFF  G M + + +  V + +G+   D   GF +   +
Sbjct  33   GRVHHLALKDDVRHKVHLNTFGFFKDGYMVVNVSSLSVNEPEGATDKDAEIGFSLDRTKN  92

Query  88   DSGSFIEETDASKC---------------------VSLLDEPERG-----------DTIA  115
            D  S   + D + C                     V +   PE G           + I
Sbjct  93   DGFSSYLDEDVNYCILKKKSMSSVTLVILDISGSIVKVRSPPEAGKQLPEIVFSKDEKIL  152

Query  116  VKVTKPS-----KENEKQ--------------SKVIT--------------------IDK  136
             +  +P+     K++E +              SK IT                    I
Sbjct  153  SQSQEPAVSSNPKDSEARRTLDGFKAGRSTVDSKAITERSFSIHKNDGVVSFQFFFNIST  212

Query  137  TRPEGFYTVVFLNC-----QPGTYVSFDLTLTNYNPGPN-YLSAGLTALPTLYAMLFVVW  190
               EG Y++ F  C     +PG   SF L +      PN YLSAG   LP LY  + + +
Sbjct  213  DDQEGLYSLYFHKCSGNNVKPGEQASFSLNIAITEKNPNSYLSAGEIPLPKLYVSMALFF  272

Query  191  TVILGVWLFHFMRGQGKRIFRIHHLVTGIILLKLLTLLFEAIEFHYKKTTGHP-GGWVIA  249
             +   +W+ H +R +   +F+IH L+  +   K L+L+F AI++HY  + G P  GW +
Sbjct  273  FLSGTIWI-HILRKRRNDVFKIHWLMAALPFTKSLSLVFHAIDYHYISSQGFPIEGWAVV  331

Query  250  YYIFSGLKGTMMFVVIALIGTGWAFIKPFLGEKDKNIFLVVIPLQILANIAIIVLEETA-  308
            YYI   LKG ++F+ IALIGTGWAFIK  L +KDK IF++VIPLQ+LAN+A I++E T
```

```
Sbjct  332  YYITHLLKGALLFITIALIGTGWAFIKHILSDKDKKIFMIVIPLQVLANVAYIIIESTEE  391

Query  309  --------PEVLRLVDIICCGAILVPIIWSIKHLRDAAAIDGKAK  345
                     + L LVD++CCGAIL P++WSI+HL++A+A DGK K
Sbjct  392  GTTEYGLWKDSLFLVDLLCCGAILFPVVWSIRHLQEASATDGKGK  436
```

>unnamed protein product [Mus musculus]
Sequence ID: BAB31509.2 Length: 94
Range 1: 1 to 80

Score:77.8 bits(190), Expect:7e-17,
Method:Composition-based stats.,
Identities:38/80(48%), Positives:50/80(62%), Gaps:1/80(1%)

```
Query  360  LVVTYIYFTRIIVFLLDATLHYQYVWLGEFFTELATLIFWGLTGYKFRPVADNPYLKLDD  419
            +V+ YIYFTRII  LL   + +Q+ WL +   E +TL F+ LTGYKF+P  DNPYL+L
Sbjct  1    MVICYIYFTRIIAILLQVAVPFQWQWLYQLLVESSTLAFFVLTGYKFQPAGDNPYLQLPQ  60

Query  420  E-EEDAEREEAQRQSRTEAG  438
            E EED + E+    S    G
Sbjct  61   EDEEDVQMEQVMTDSGFREG  80
```

>G protein-coupled receptor 108, isoform CRA_b [Mus musculus]
Sequence ID: EDL38249.1 Length: 125
Range 1: 1 to 54

Score:66.6 bits(161), Expect:2e-12,
Method:Composition-based stats.,
Identities:28/54(52%), Positives:37/54(68%), Gaps:0/54(0%)

```
Query  360  LVVTYIYFTRIIVFLLDATLHYQYVWLGEFFTELATLIFWGLTGYKFRPVADNP  413
            +V+ YIYFTRII  LL   + +Q+ WL +   E +TL F+ LTGYKF+P  DNP
Sbjct  1    MVICYIYFTRIIAILLQVAVPFQWQWLYQLLVESSTLAFFVLTGYKFQPAGDNP  54
```
